# The *abaI/abaR* quorum sensing system effects pathogenicity in *Acinetobacter baumannii*

**DOI:** 10.1101/2021.01.19.427366

**Authors:** Xiaoyu Sun, Zhaohui Ni, Jie Tang, Yue Ding, Xinlei Wang, Fan Li

## Abstract

*Acinetobacter baumannii* is a Gram-negative pathogen that has emerged as one of the most troublesome pathogens for health care institutions globally. Bacterial quorum sensing (QS) is a process of cell-to-cell communication that relies on the production, secretion and detection of autoinducer (AI) signals to share information about cell density and regulate gene expression accordingly. In this study, we performed a comprehensive set of experiments show that deletion of quorum sensing genes showed differences in growth characteristics, morphology, biofilm formation and virulence, and increased susceptibility to some antimicrobials and exhibited motility defects. RNA-seq analysis indicated that genes involved in various aspects of energy production and conversion, Valine, leucine and isoleucine degradation and lipid transport and metabolism showed different expression.

**IMPORTANCE:** Previous studies on bacterial quorum sensing mainly focused on biofilm formation and motility and antibiotic resistance. In this study, we focused on detecting the role of the *abaI/abaR* QS system in the virulence of *A. baumannii*. Our work provides a new insight into *abaI/abaR* quorum sensing system effects pathogenicity in *A. baumannii*. We propose that targeting the AHL synthase enzyme *abaI* could provide an effective strategy for attenuating virulence. On the contrary, interdicting the autoinducer synthase–receptor *abaR* elicits unpredictable consequences, which may lead to enhanced bacterial virulence.

## INTRODUCTION

*Acinetobacter baumannii* is a Gram-negative pathogen that has emerged as one of the most troublesome pathogens for health care institutions globally. *A. baumannii* infections are increasingly difficult to treat because of multidrug-resistance in most strains causing infection in intensive care units (ICUs) (1). Virulence factors were used by bacteria that enable successful interaction with and subsequent adhesion and invasion of the human host (2). Previous studies have emphasized the importance of iron acquisition, transports, cell-associated pili, lipopolysaccharides, outer membrane proteins such as *ompA, omp33, surA1* for virulence (3–5), but studies of quorum sensing effects pathogenicity in *A. baumannii* are limited. Bacterial quorum sensing (QS) is a process of cell-to-cell communication that relies on the production, secretion and detection of autoinducer (AIs) signals to share information about cell density and regulate gene expression accordingly (6). AIs are involved in the regulation of varied biological functions, including expression of virulence gene in *Vibrio cholera* (7), *Pseudomonas aeruginosa* PAO1 (8), *Staphylococcus aureus* (9), *Escherichia coli* (10) and other types of bacteria.

*A. baumannii* presenting a typical quorum sensing (QS) system (*abaI*/*abaR*) has been described(11). The *abaI* gene encodes 183 amino acids, and this protein was predicted to function in signal transduction. The *abaR* gene encodes 238 amino acids, this protein is an autoinducer synthase–receptor (11). Previous studies on *A. baumannii* had focused on the role of quorum sensing systems in drug resistance, biofilm formation and motility (12, 13), but studies of quorum sensing effects pathogenicity in *A. baumannii* are poor.

In this study, we used *A. baumannii* strain ATCC 17978 which has been the most frequently used model for scientific studies over the past two decades (2). A previous study isolated and characterized the autoinducer synthase *abaI* from *Acinetobacter baumannii* M2, but failed to create an *abaR* deletion mutant for unknown reasons (14). To explore the role of the *abaI*/*abaR* QS system in drug resistance, biofilm formation and virulence of *A. baumannii*, Δ*aba*I and Δ*abaR* mutants of strain ATCC 17978 were created. We also made double mutant Δ*abaIR*. The transcriptomes of WT, Δ*abaI*, Δ*abaR* and Δ*abaIR* were determined by RNA-sequencing (RNA-seq).

## RESULTS

### The mutants showed differences in growth characteristics and morphology

To test the role of the *abaI*/*abaR* QS system in *A. baumannii* growth curve, we determined by measuring the optical density (OD) of the culture over time. The growth of the Δ*abaR* and Δ*abaIR* mutants did not differ from that of the parent strain (Fig. 1A). The complemented strain Δ*abaR*(pME*abaR*) and overexpressed strains WT(pME*abaI*), WT(pME*abaR*) also showed growth profiles similar to those of the wild type(Fig. 1A), suggesting that the gene *abaR* is not essential for *A. baumannii* growth. In contrast, Δ*abaI* mutant showed visibly slowed growth at the logarithmic phase. Growth of the Δ*abaI* mutant was partly rescued by expression of *abaI* via the pME6032-derived plasmid pME*abaI* (Fig. 1A). The empty vector pME6032, used as a control, did not affect the growth profile of *A. baumannii* ATCC 17978(Fig. 1A). These results demonstrated that *abaI* affect the growth of *A. baumannii*. Subsequently, the morphology of the bacteria was observed by transmission electron microscopy. As shown in Figure 1B, WT was attached with extracellular secretions. Δ*abaI* mutant cell edges were transparent. Δ*abaR* mutant strains cytoplasmic density is lower. There was no obviously change in Δ*abaIR* mutant, but around the cell adhered partial secretions. Morphology of the Δ*abaI* mutant was partly rescued by expression of *abaI* via the pME6032-derived plasmid pME*abaI.* Morphology of the Δ*abaR* mutant was rescued by expression of *abaR* via the pME6032-derived plasmid pME*abaR*.

**Figure 1.**
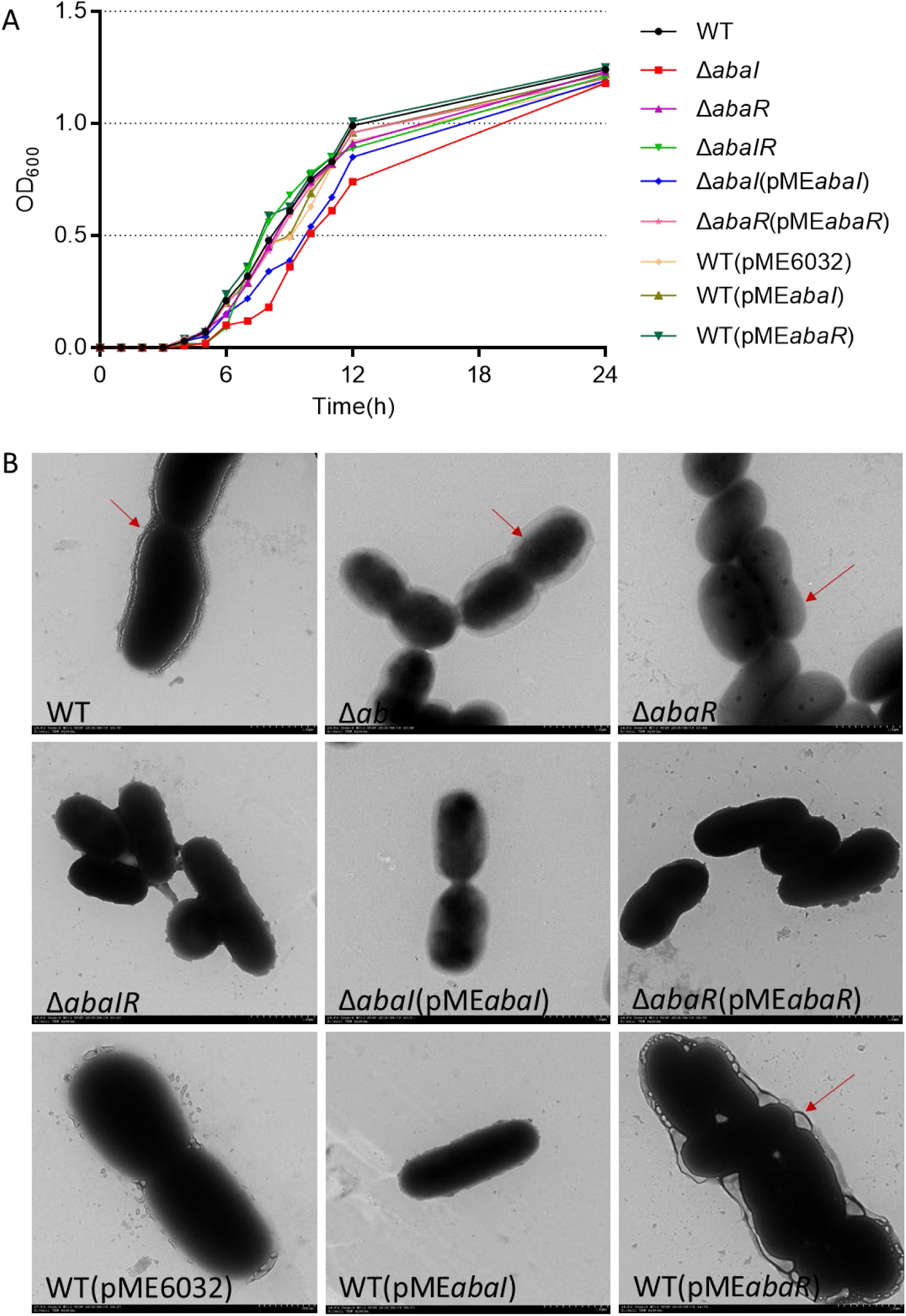
A) Growth curve analysis of *A.baumannii* and mutants in broth. B) Transmission electron microscope images of the targeted bacteria. Cells were observed with a HITACHI H-7650 TEM transmission electron microscope operated at 120 kv. Scale bar = 500 nm.

There was no significant difference between WT and overexpressed strain WT(pME*abaI*). The overexpressed strain WT(pME*abaR*) attached a large number of extracellular secretions. The empty vector pME6032, used as a control, did not affect the morphology of *A. baumannii* ATCC 17978. These results indicated that QS is closely related to cell morphology and extracellular secretions.

### Mutants showed increase susceptibility to antimicrobials

To determine whether deletion of quorum sensing genes changed in the drug resistance, the MICs of commonly used antibiotics for strains was determined. There was a decrease in the MICs of Kanamycin, Gentamicin, Penicillin, Streptomycin, Meropenem, Imipenem and Ampicillin for quorum sensing genes deletions compared to WT. Drug susceptibility of Δ*abaI* and Δ*abaR* mutants were partly rescued by expression of *abaI* and *abaR* via plasmid pME*abaI* and pME*abaR* respectively. Overexpressed strain WT(pME*abaI*) were more resistant in Cefepime and Cefperazone-Sulbactam. WT(pME*abaR*) were more resistant in Kanamycin, Streptomycin, Ceftizoxime, Cefepime, Cefperazone-Sulbactam and Paraxiline-tazobatan. The empty vector pME6032, used as a control, partly affect the drug susceptibility of *A. baumannii* ATCC 17978 (Table 4). These results indicated that QS affect strain resistance.

### Screening of strains for AHLs production

To detect the effect of quorum sensing system on AHL of the strains, two different biosensor strains were used. As a result, no strains were found to produce AHLs based on the development of purple coloration in CV026 reporter strain (Fig. 2A to C). Δ*abaI*, Δ*abaR* and Δ*abaIR* were not observed to produce AHLs based on the development of green coloration in *A. tumefaciens* KYC55 reporter strain. AHLs production of Δ*abaI* and Δ*abaR* mutants were rescued by expression of *abaI* and *abaR* via plasmid pME*abaI* and pME*abaR*, respectively. The empty vector pME6032, used as a control, did not affect the AHLs production of *A. baumannii* ATCC 17978 (Fig. 2D to F). These results indicate that *abaI* and *abaR* both can affect AHLs production and *A. baumannii* may produce only long-chain signal molecules, but not short-chain AHLs.

**Figure 2.**
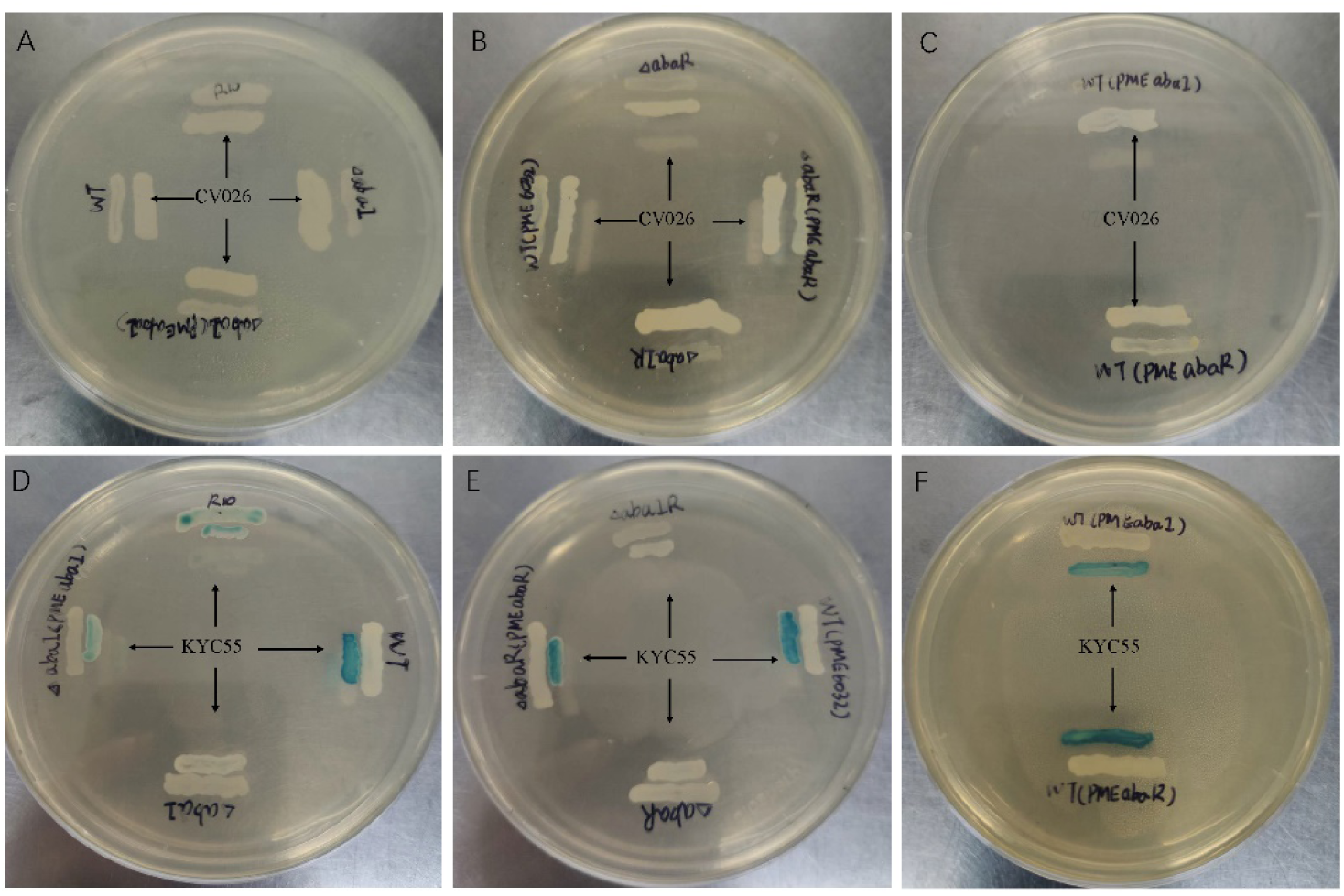
Screening of WT and mutants for AHLs production using agar-plate well diffusion assay with *C. violaceum* CV026 and *A. tumefaciens* KYC55 reporter strain.

### Surface-associated motility relies on quorum sensing

To explore the role of quorum sensing in motility, we tested these strains surface motility phenotype on LB media supplemented with 0.3% agar. As a result, Δ*abaI*, Δ*abaR* and Δ*abaIR* mutants compared with wild-type strain showed no obvious motility, motility of Δ*abaI* mutants were not rescued by expression of *abaI* via the plasmid pME*abaI*, motility of Δ*abaR* mutants were partly rescued by expression of *abaR* via the plasmid pME*abaR*. The empty vector pME6032, used as a control, has an inhibitory effect on the motility of *A. baumannii* ATCC 17978. Overexpressed strain WT(pME*abaR*) compared with wild-type strain showed significantly increased motility. There was no significant difference between overexpressed strain WT(pME*abaI*) and WT in the motility(Fig. 3A, B). These results indicated that quorum sensing system affects the motility of bacteria and *abaR* regulates bacterial motility.

**Figure 3.**
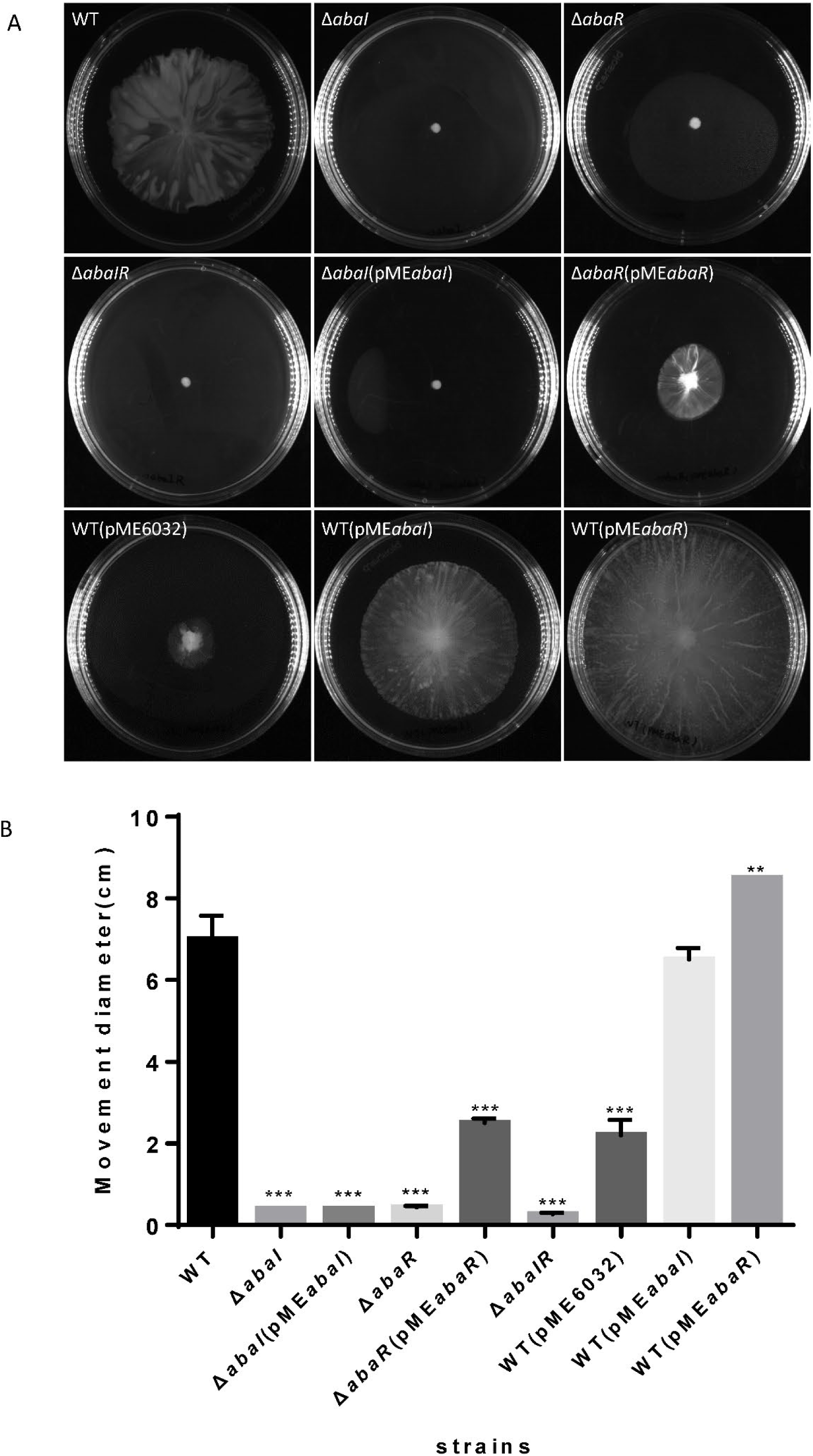
A) WT and mutants were inoculated on the surface of a semisolid agarose plate (0.3%) and incubated for 12–14 h at 30°C. *abaI*, *abaR*, and *abaIR* mutant strains demonstrated defects in surface-associated motility compared to the parental strain. B) The distance migrated (diameter) is shown for each strain and Error bars represent SD for three biological replicates.

### Biofilm formation

To investigate the role of quorum sensing in biofilm formation on an abiotic surface, we cultivated the strains mutants in 96-well plates for 24 h at 37°C. As a result, Filaments were formed in the culture medium of wild strain, mutant strains Δ*abaI*, Δ*abaR* had dot-like biofilm formation on the surface of liquid, Δ*abaIR* mutant strain had no biofilm on the liquid surface, there were dot-like biofilms on the liquid surface of Δ*abaI*(pME*abaI*) and Δ*abaR*(pME*abaR*)and they were connected into pieces. The biofilm on the liquid surface of the overexpressed strain WT(pME*abaI*) was lamellar, while the biofilm on the liquid surface of WT(pMEabaR) was spot-like but thick (Fig. 4A). The absorbance of bacterial solution was detected, and the results were shown in the Figure 4B, there were significant difference between the absorbance of the parental strain and mutants Δ*abaI*, Δ*abaR,* however, for mutant Δ*abaIR* showed no difference. The absorbance of Δ*abaI* mutants were not rescued by expression of *abaI* via the plasmid pME*abaI*, absorbance of Δ*abaR* mutants were rescued by expression of *abaR* via the plasmid pME*abaR*. The absorbance of overexpressed strain WT(pME*abaR*) was significantly higher than that of wild strain. The biofilm forming ability of the strains were determined by crystal violet biofilm assay, and the differences between the wild-type and mutant strains were calculated. Compared with wild strains, the biofilm formation of all strains was significantly decreased, except for the overexpressed strain WT(pME*abaR*), which was significantly higher than that of wild strain (Fig. 4C). The results indicated that quorum sensing system affects the biofilm formation of bacteria.

**Figure 4.**
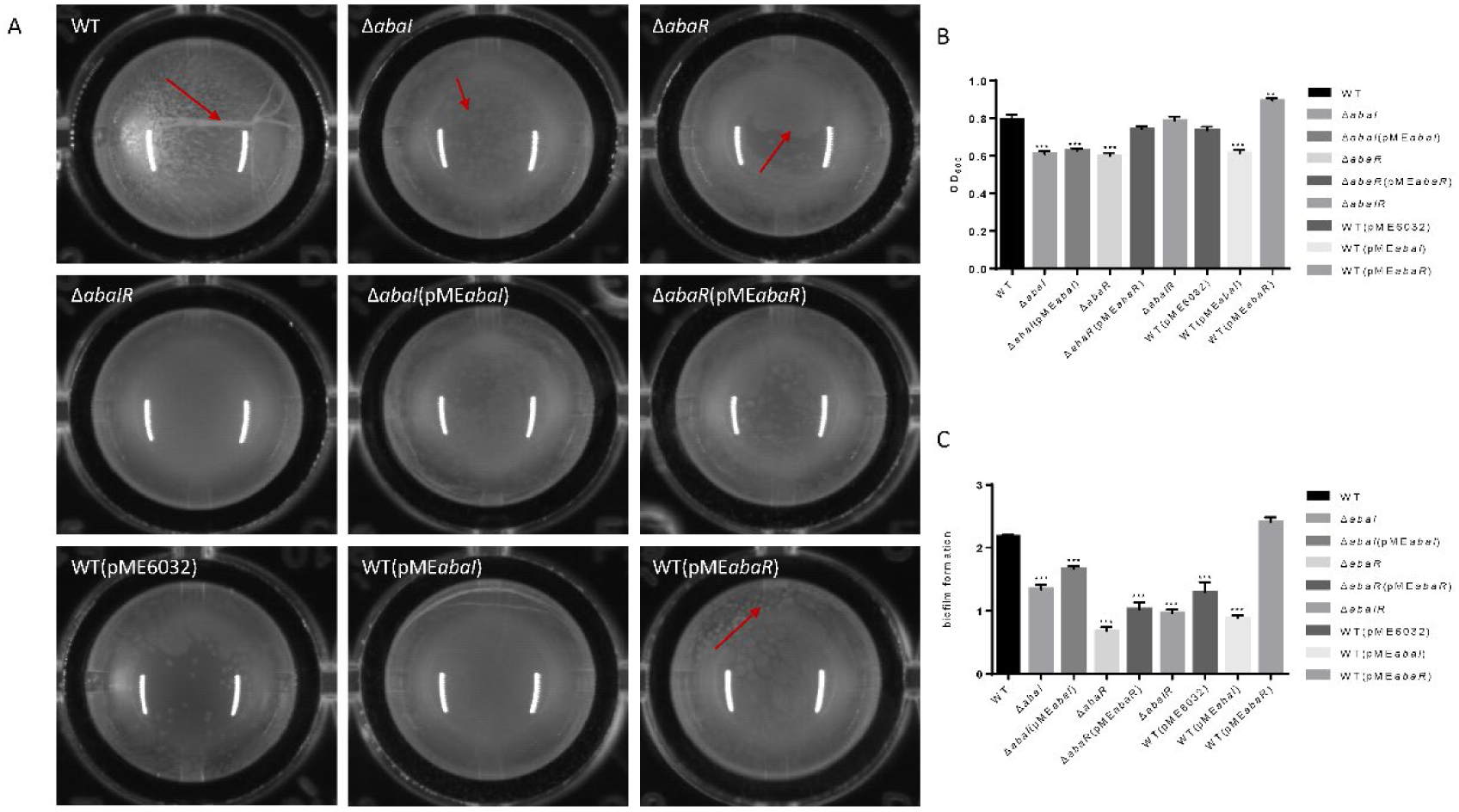
A) Visual changes in biofilm formation in 96-well plates for 24 h at 37°C, images were taken after 24 of growth. B) The absorbance of bacterial solution was detected by OD595. C) Biofilm formation on plastic at 37°C as determined by crystal violet staining. Markers show the OD595 compared with WT in individual biological replicates.

### Serum killing

Serum sensitivity has been involved in the toxic mechanisms of *A. baumannii*, to elucidate the virulence of strains, we compared the serum sensitivity of mutants to WT. As shown in Fig 5 A, WT, *abaR* and WT(pME*abaR*) survived after incubation in serum, whereas Δ*abaI*, Δ*abaI*(pME*abaI*)and Δ*abaIR* were entirely killed after 1 h at 37°C. The serum sensitivity of Δ*abaI* mutants were not rescued by expression of *abaI* via the plasmid pME*abaI*, serum sensitivity of Δ*abaR* mutants were rescued by expression of *abaR* via the plasmid pME*abaR*. The empty vector pME6032, used as a control, did not affect the serum sensitivity of *A. baumannii* ATCC 17978. These results indicated that WT and Δ*abaR* were highly resistant to the killing action of serum, in contrast, Δ*abaI* and Δ*abaIR* mutants were much more serum sensitive, showing a significant difference.

**Figure 5.**
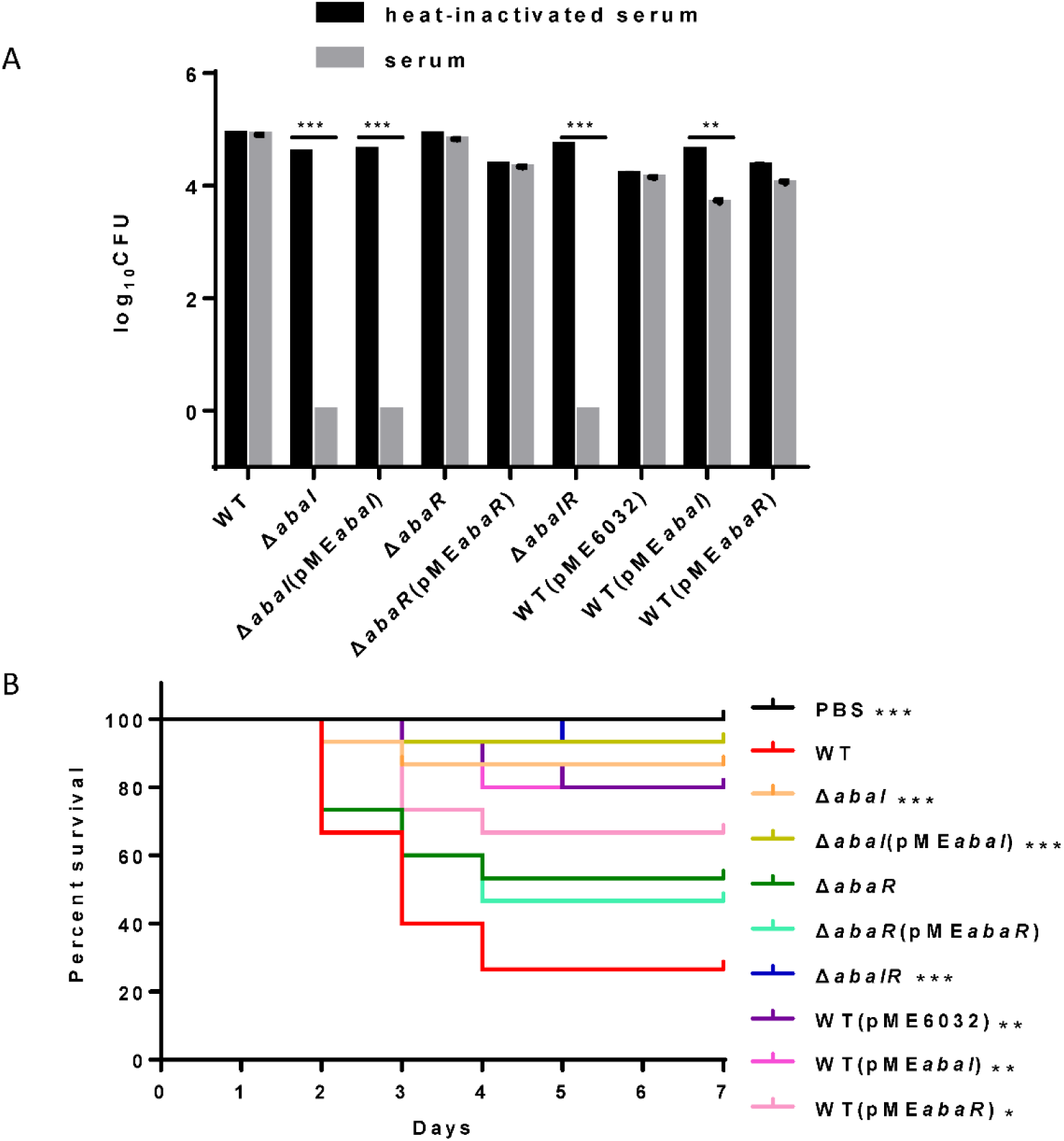
A) Sensitivity of strains to normal human serum (NHS). Viable bacterial counts were determined after 1 h of incubation at 37°C with rotation. Data are presented as percent survival, with 100% being the number of viable bacteria grown in heat-inactivated serum. All values are from triplicate samples and are representative of three independent experiments. B) Kaplan-Meier survival curve showing the virulence of PBS, WT and mutants in *G. mellonella*. The data show the percent survival (n=30) of *G. mellonella* after inoculation with 10^6^ CFU of bacteria. Survival curves were compared using the log-rank (Mantel-Cox) test. *p < 0.05 **;p < 0.005; ***p < 0.001.

### Quorum sensing plays a role in virulence in *G. mellonella* infection models

Quorum sensing controls the production of virulence factors in many bacterial species and is regarded as an attractive target to combat bacterial pathogenicity (15). To explore whether quorum-sensing genes are an important virulence factor determinant for *A. baumannii* in *G. mellonella,* we assessed the virulence of the quorum-sensing mutant strains compared to the isogenic parent strain ATCC 17978. Δ*abaI* and Δ*abaIR* mutants were completely avirulent in this assay, while Δ*abaR* mutant remained fully virulent, killing *G. mellonella* larva as toxic as the wild type (Fig. 5B). Consistent with this finding, Laura Fernandez-Garcia found that injection of *G. mellonella* larvae with the *A. baumannii* ATCC 17978 strain caused higher mortality than injection with mutant *A. baumannii* ATCC 17978 Δ*abaI* (16). The virulence of Δ*abaI* mutant was not rescued by expression of *abaI* via the plasmid pMEabaI, virulence of Δ*abaR* mutant was rescued by expression of *abaR* via the plasmid pME*abaR*. The empty vector pME6032, used as a control, affect the virulence of *A. baumannii* ATCC 17978. The virulence of overexpressed strain WT(pME*abaI*) and WT(pME*abaR*) were significantly reduced compared to the parent strain ATCC 17978. The results indicated that quorum sensing system plays a role in virulence in *G. mellonella* infection models.

### Quorum sensing plays a role in virulence in mouse infection models

Whether these quorum-sensing genes of *A. baumannii* are important for virulence to a mammalian system is currently unknown. To evaluate the virulence of strains, we established a bacteremia model in mice by intraperitoneal injection of *A. baumannii*. In the experiments we analyzed the survival of infected mice with this model, found that a dose of approximately 1.8×10^8^ CFU of Δ*abaI* and Δ*abaIR* mutants were unable to cause lethality, but only one mouse (10/group) survived in the WT group, and all the mice in Δ*abaR* group died (date not shown) after 48 h. Subsequently, we used a dose of approximately 1.2×10^8^ CFU of bacteria infected with 10 mice per group for survival studies. We observed that Δ*abaI* and Δ*abaIR* mutants were unable to cause lethality; WT exhibited a low fatality rate, but Δ*abaR* and complemented strain Δ*abaR*(pME*abaR*) can cause more deaths (Fig. 6A). These results indicated that deletion of *abaI* results in weaker toxicity in mouse models, while deletion of *abaR* results in enhanced virulence. To explore the virulence of mutants in host resistance against *A. baumannii* infection, the blood, lungs and spleens from BALB/C were collected at various time points after injected with 1.2×10^8^ CFU of *A. baumannii*. We analyzed bacterial burdens in the blood, lung and spleen of mice infected for 4 h, 24 h and 72 h. As a result, WT, Δ*abaR* mutant and complemented strain Δ*abaR*(pME*abaR*) resulted in an increase in bacterial counts, Δ*abaI* and Δ*abaIR* mutants exhibited a remarkable reduction in the burden in blood, lung, and spleen at 4 h postinoculation. More specifically, as shown, Δ*abaI* and Δ*abaIR* mutants displayed a high serum clearance rate and resulted in a significant decrease in bacteremia at 4 h postinoculation. There were no differences at 24 h (Fig. 6B). The bacteria were completely eliminated at 72 hours (data not shown). These results indicated that quorum sensing system plays a role in virulence in mouse infection models.

**Figure 6.**
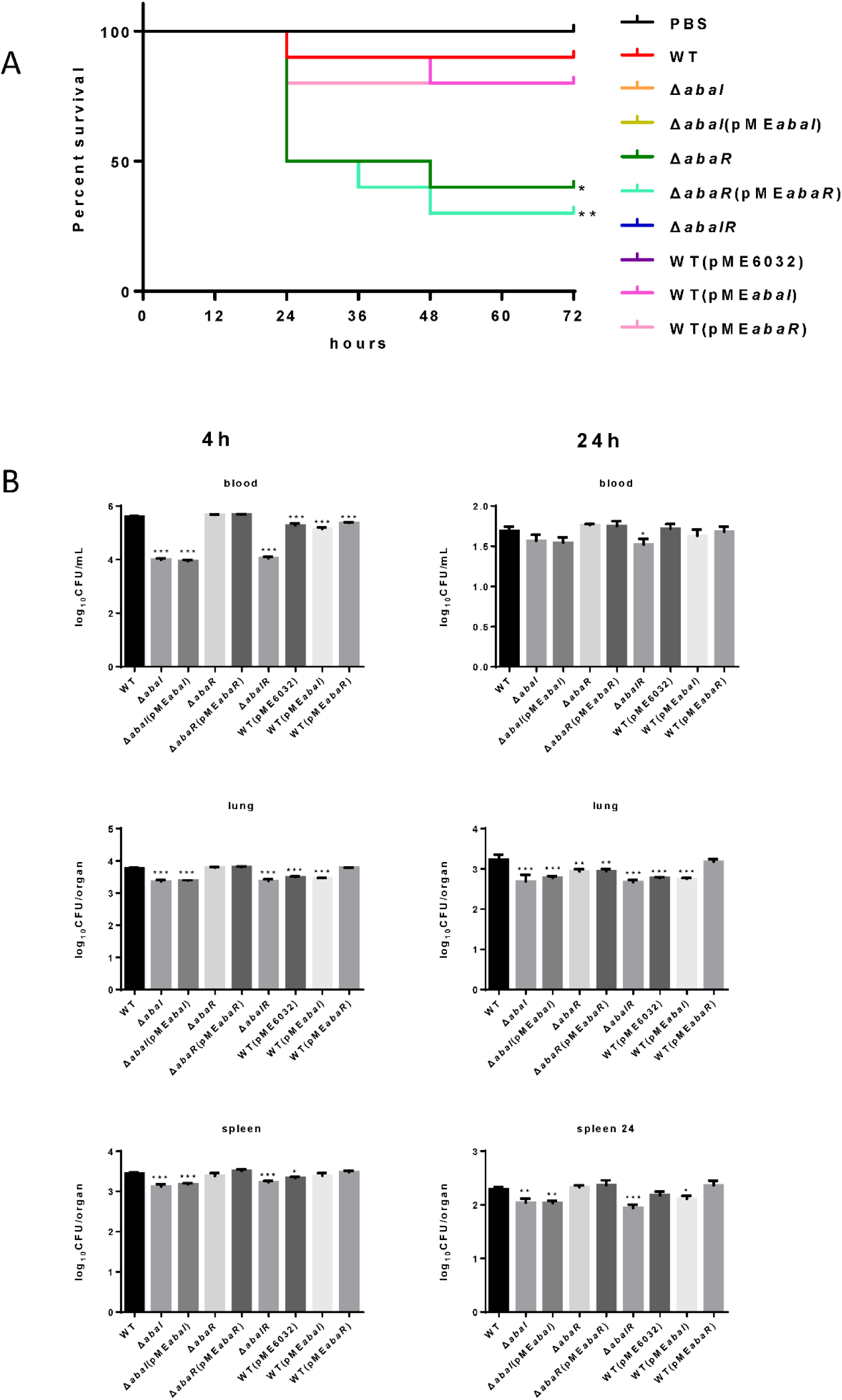
A) Kaplan-Meier survival curve of mouse inoculated with strains at a dose of 1.2×108 CFUs (n=10). Survival curves were compared using the log-rank (Mantel-Cox) test, *p < 0.05. B) *A. baumannii* bacterial burdens in the lungs, spleen, and blood. Bacterial burdens in the blood and respective organs were determined by quantitative bacteriology at 4 and 24 h postinoculation. An unpaired t test was used to validate the experimental data. *p < 0.05; **p < 0.005; ***P< 0.001.

### Mutants causes differential gene expression in *A. baumannii* ATCC 17978

To identify transcriptional activity dependent on *abaI*/*abaR* function, RNA-seq analysis was performed on *abaI*/*abaR* mutants and WT strain. A total of 463 genes were classified as differentially expressed in mutants relative to WT. Compared with WT, in Δ*abaI* mutant, a total of 159 protein-coding genes (out of 3848) were identified as differentially expressed by a log2-fold change greater than 1 or less than −1 (P <= 0.05) (126 with increased expression, 33 with decreased expression). Deletion of *abaR* had a larger impact on the transcriptome of strain ATCC 17978, with the differential expression of 324 genes (211, 113 up down regulated). Δ*abaIR* mutant had a total of 123 differentially expressed genes (79 genes with increased expression, 44 with decreased expression) (Fig. 7A to C) (Supplementary table).

**Figure 7.**
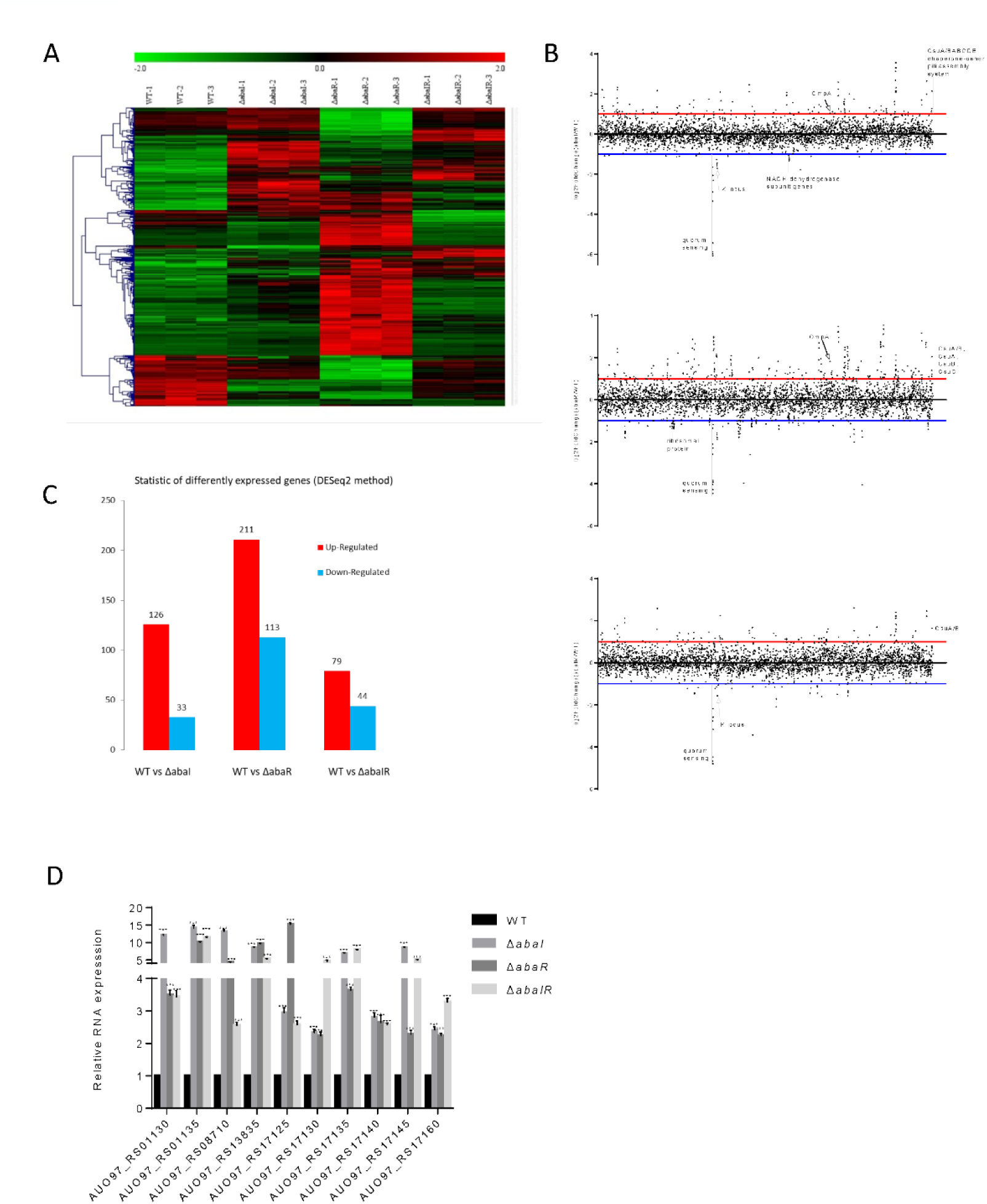
A) Heat map representation and hierarchical clustering of gene expression changes. The x-coordinate represents the different experimental groups. Data represents differential gene expression profiles (PFKM) for genes listed along ordinate. Red indicates increased expression and green indicates decreased expression; color intensity indicates the magnitude of difference in expression according to horizontal scale at top. Black depicts genes with no significant difference in expression. B) The genome wide transcriptomic profile, genome-wide differential gene expression locus map, Plot of differential gene expression mutant compared with WT with respect to gene locus tag number. C) The vertical axis represents the number of significantly differentially expressed genes. The log2-fold change in expression for each gene meeting the study threshold (log2 fold change > 1, false discovery rate <0.001). The number of significantly differentially expressed genes can be divided into the number of significantly up-regulated differentially expressed genes and the number of significantly down-regulated differentially expressed genes. D) Relative expression of 10 up-regulated genes, results are presented relative to WT which was normalized to 1. The results are expressed as mean ± SEM for at least 3 biological replicates. ***P < .001, t test.

These results revealed that partial changes in gene expression occur with changes in *abaI/abaR*. To validate our RNA-seq analysis, quantitative reverse transcription qRT-PCR was used to validate the 10 upregulated genes (Fig. 7D). The results were consistent with the high-throughput sequencing data.

### COGs annotation

To link transcriptional reprogramming by *abaI/abaR* to function, differentially expressed genes were categorized into COGs (17, 18). The COG enrichment analysis identified that 7% of the genes belonging to the COG category [C] Energy production and conversion, were significantly repressed, and 10% of the genes were associated with the COG category [Q] Secondary metabolites biosynthesis, transport and catabolism, were highly regulated in Δ*abaI* mutant. We observed a high proportion of genes with increased expression in Δ*abaR* mutant that encoded proteins involved in [I] Lipid transport and metabolism (21%), [Q] Secondary metabolites biosynthesis, transport and catabolism (18%), [G] Carbohydrate transport and metabolism (12%), [C] Energy production and conversion (11%) and [E] Amino acid transport and metabolism (10%), while a strong decrease in [J] Translation, ribosomal structure and biogenesis(24%). In Δ*abaIR* mutant, 4% of the genes belonging to the COG category [C] Energy production and conversion, and [I] Lipid transport and metabolism were down-regulated, while 5% of the genes belonging to the COG category [G] Carbohydrate transport and metabolism were up-regulated (Fig. 8).

**Figure 8.**
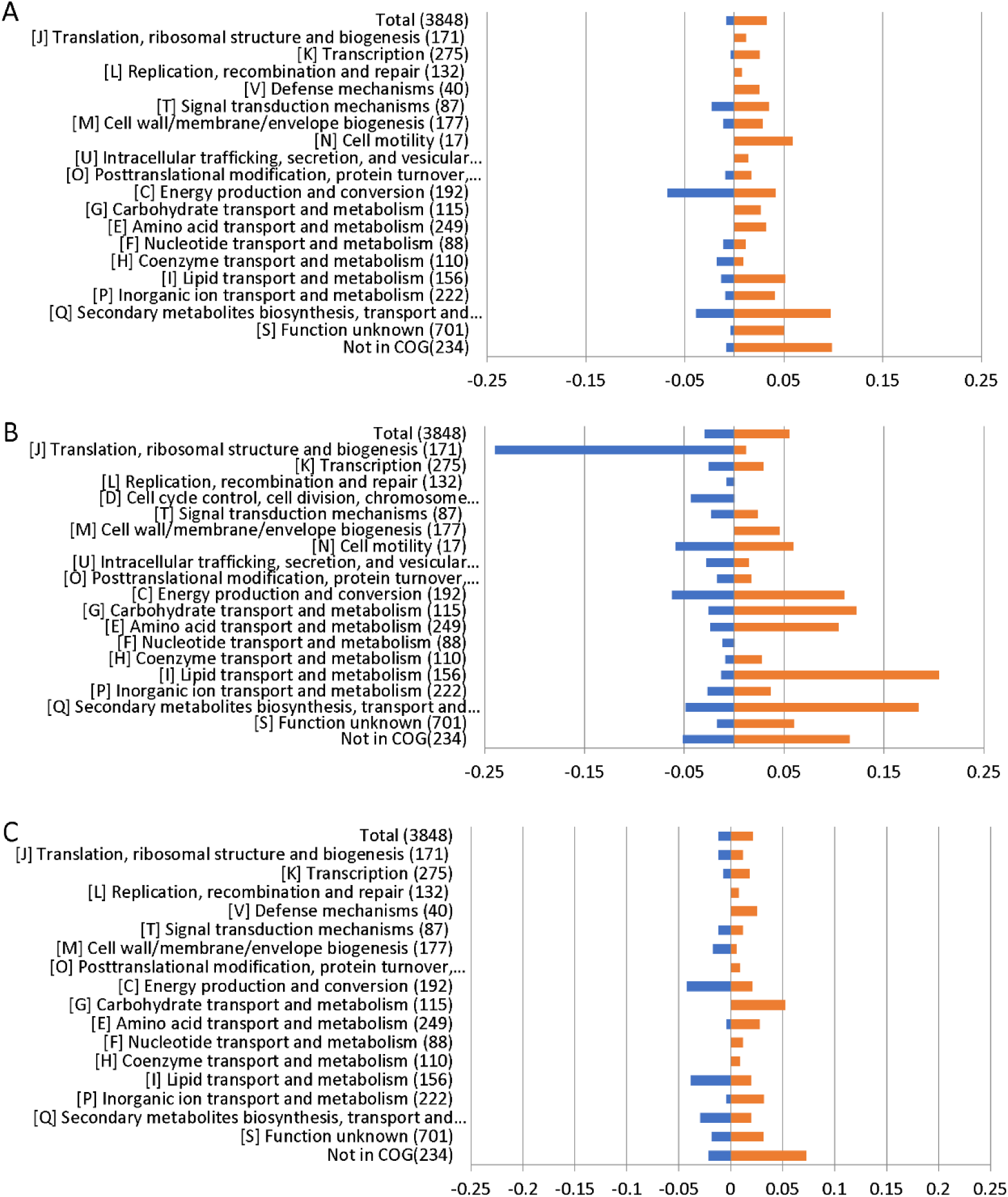
RNA-Seq results displayed by COG enrichment for differentially regulated genes. Cluster of orthologous groups (COG) enrichment analysis was performed by dividing the percentage of genes up-or down-regulated for each category by the percentage of genes in that category across the whole genome. Panels: A, Δ*abaI*; B, Δ*abaR*; C, Δ*abaIR*.

### GO and KEGG pathways enrichment analysis of DEGs

The functions of the differentially expressed genes were annotated and classified gene ontology (GO) and Kyoto Encyclopedia of Genes and Genomes (KEGG) pathway enrichment analyses. The top 30 of significant GO terms were divided into two major categories—biological process and cellular component in Δ*abaI* mutant, as shown in Figure 9A. Among the biological processes, the DEGs were distributed to xenobiotic metabolic process, toxin metabolic/catabolic process, auxin metabolic /catabolic process, phenylacetate catabolic process and so on. In the cellular component field, DEGs were belonged to respiratory chain complex I, plasma membrane respiratory chain complex I, NADH dehydrogenase complex and respiratory chain. The top 30 significant GO terms were divided into three major categories—biological process, cellular component and molecular function in Δ*abaR* mutant, as shown in Figure 9C. Among the biological processes, the DEGs were distributed to ribosome assembly, translation, ribonucleoprotein complex assembly, organelle assembly, ribonucleoprotein complex subunit organization and peptide biosynthetic process. In the cellular component field, DEGs were belonged to cytosolic ribosome, ribosome, ribonucleoprotein complex, organelle part and cytosolic part. In molecular function, DEGs were responsible for structural constituent of ribosome, structural molecule activity and rRNA binding. The top 30 significant GO terms were divided into three major categories—biological process, cellular component and molecular function in Δ*abaIR* mutant, as shown in Figure 9E. Among the biological processes, the DEGs were distributed to response to metal ion, response to organonitrogen compound, cellular response to stress and anion transport. In the cellular component field, DEGs were belonged to outer membrane−bounded periplasmic space, periplasmic space, cell envelope and envelope. In molecular function, DEGs were responsible for copper ion binding, cation/amino acid/organic acid/ organic anion transmembrane transporter activity.

**Figure 9.**
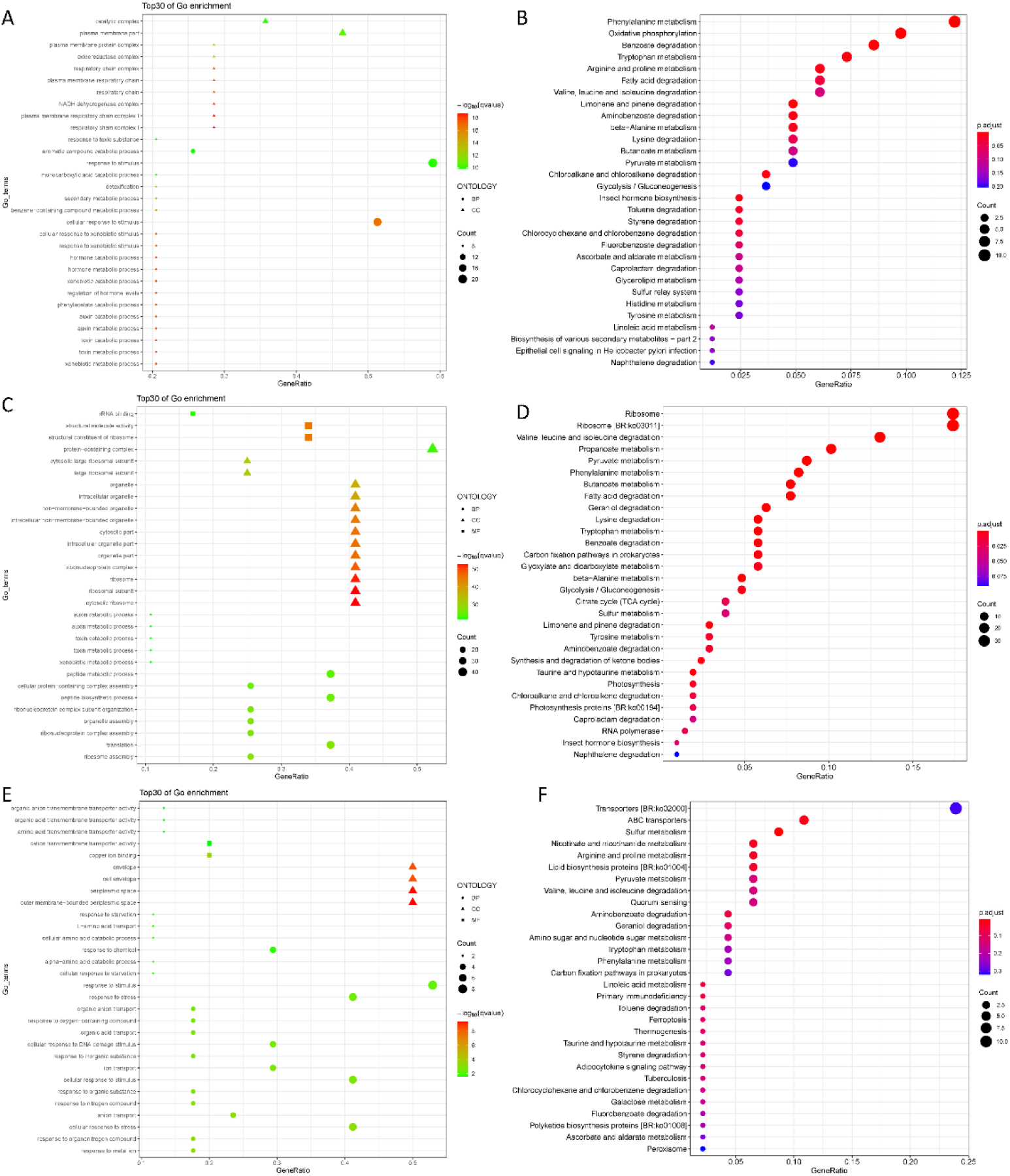
A) Top 30 of GO enrichment of DEGs in Δ*abaI*. The shape of the point indicates the different ontology. The enrichment q value of each GO term was normalized as negative log10(q value) and is shown as a color gradient. The number of genes enriched in each GO term is represented by the size of the points. B) The significantly enriched KEGG pathway of DEGs in Δ*abaI*. The enrichment P adjustvalue of each pathway was shown as a color gradient. GeneRatio, number of differentially expressed genes/total number of genes in this KEGG pathway. The number of genes enriched in each pathway is represented by the size of the points. C) Top 30 of GO enrichment of DEGs in Δ*abaR*. The shape of the point indicates the different ontology. The enrichment q value of each GO term was normalized as negative log10(q value) and is shown as a color gradient. The number of genes enriched in each GO term is represented by the size of the points. D) The significantly enriched KEGG pathway of DEGs in Δ*abaR*. The enrichment P adjustvalue of each pathway was shown as a color gradient. GeneRatio, number of differentially expressed genes/total number of genes in this KEGG pathway. The number of genes enriched in each pathway is represented by the size of the points. E) Top 30 of GO enrichment of DEGs in Δ*abaIR*. The shape of the point indicates the different ontology. The enrichment q value of each GO term was normalized as negative log10(q value) and is shown as a color gradient. The number of genes enriched in each GO term is represented by the size of the points. F) The significantly enriched KEGG pathway of DEGs in Δ*abaIR*. The enrichment P adjustvalue of each pathway was shown as a color gradient. GeneRatio, number of differentially expressed genes/total number of genes in this KEGG pathway. The number of genes enriched in each pathway is represented by the size of the points.

KEGG pathway enrichment suggested that DEGs of Δ*abaI* mutant were significantly enriched in the pathways related to Phenylalanine metabolism, Oxidative phosphorylation, Benzoate degradation, Tryptophan metabolism, Arginine and proline metabolism, Fatty acid degradation and so on (Fig. 9 B). DEGs of Δ*abaR* mutant were mainly involved in the pathways related to Ribosome, Valine, leucine and isoleucine degradation, Propanoate metabolism, Pyruvate metabolism, Phenylalanine metabolism, Fatty acid degradation and so on (Fig. 9D). DEGs of Δ*abaIR* mutant were mainly involved in the pathways related to ABC transporters, Sulfur metabolism Nicotinate and nicotinamide metabolism and so on (Fig.9 F).

Given the different phenotypes of Δ*abaI*, Δ*abaR* and Δ*abaIR* mutants, the RNA-seq analysis revealed a subset of the genes that was most highly activated or suppressed by quorum sensing systerm, as shown in Figure 7B. We centralized our analysis on the gene subsets whose transcription was down in Δ*abaI* and Δ*abaR* and Δ*abaIR* mutants, the quorum sensing gene (*abaI* AUO97_RS06645) and nearby locus (AUO97_RS06600–06630) showed strongly reduced transcription. One study found that in *A. baumannii* M2 strain quorum sensing mediated by the *abaI* is required for motility (19). We assessed the surface motility of the mutants, as shown in Figure 3. We observed that wild-type strain exhibited a robust surface motility phenotype and the mutants did not exhibit any signs of motility.

We mainly analyzed the gene expression of each group. Hemerythrin-like proteins have an effect on oxidation-reduction regulation and antibiotic resistance (20). AUO97_RS11650 (hemerythrin) was down-regulated in Δ*abaI* mutant, and other antimicrobial resistance genes including AUO97_RS07490 (*mexK*), AUO97_RS07485 (*mexJ*), AUO97_RS07485 (efflux RND transporter periplasmic adaptor subunit) and AUO97_RS05665 (beta-lactamase domain protein), were down regulated. One gene, AUO97_RS16540 (*AdeA*/*AdeI* family multidrug efflux RND transporter periplasmic adaptor subunit) had 1.2-fold increased expression in Δ*abaR* mutant, while the expression of this gene was not changed in the Δ*abaIR* mutant. We then assessed the antimicrobial susceptibility of the mutants, as shown in Table 4, susceptibility of mutants towards a part of antimicrobials increased.

The *csu* operon is composed of 6 genes (*csuA*/*BABCDE*) and play a central role in initial bacterial attachment, and biofilm formation on abiotic surfaces (21). In Δ*abaI* mutant, the *CsuA*/*BABCDE* chaperone-usher pili assembly system was showed high expression, except for the *CsuA* (AUO97_RS19210) gene. In Δ*abaR* mutant, *CsuA/B* (AUO97_RS19215), *CsuA* (AUO97_RS19210), *CsuB* (AUO97_RS19205) and *CsuC* (AUO97_RS19200) were highly expressed, whereas in the Δ*abaIR* mutant, only *CsuA/B* (AUO97_RS19215) was highly expressed. *CsuA/B*, which is predicted to form part of the type I pili rod that was up-regulated in all mutants. Of particular interest is their regulator genes *bfmR*-*bfmS*, which did not change in all mutants. Biofilm formation was observed on the liquid surface of the Δ*abaI* and Δ*abaR* mutant (Fig. 4A). Previously study found that *CsuC* and *CsuE* are required in the early steps of biofilm formation (21). In some *A. baumannii* strains biofilms are not essential for virulence (22). Apart from the *csu* operon, other genes controlled by quorum sensing that may have relations with biofilm formation. A1S_0644 (AUO97_RS08180), a hypothetical protein involved in biofilm formation was repressed in the Δ*abaR* mutant.

Secreted bacterial proteins can mediate serum resistance, a secreted serine protease which termed PKF is required for serum resistance and inhibits biofilm formation in *A. baumannii* (23). In Δ*abaR* mutant, *PKF* (AUO97_RS01525) was highly expressed, and no difference was observed in the Δ*abaI* and Δ*abaIR* mutants. We assessed the serum sensitivity test and the ability of mutant to form biofilms on abiotic surface. As shown in (Fig. 5A and Fig.4), WT and Δ*abaR* survived after incubation in serum, whereas Δ*abaI* and Δ*abaIR* were entirely killed after 1 h at 37°C. There was a remarkable decrease in the biofilm formation by Δ*abaR* mutant. Apart from this gene, other genes that may be associated with serum resistance and biofilm formation may be regulated by quorum sensing.

NADH is mainly involved in material and energy metabolism in cells, which is transferring energy to ATP synthesis through oxidative phosphorylation on the mitochondrial inner membrane (24, 25). In the respiratory chain of *A. baumannii,* there are 14 NADH-quinone oxidoreductase subunits involved in NADH dehydrogenase, which includes *NuoA-NuoN*. In Δ*abaI* mutant, a subset of genes including AUO97_RS10980-11015 (*NuoG*, *NuoH*, *NuoI*, *NuoJ*, *NouK*, *NuoL*, *NuoM* and *NuoN*) were all down-regulated (Fig. 10A). In Δ*abaR* mutant AUO97_RS11000 (*NouK*) and AUO97_RS11015 (*NuoN*) were down-regulated, and there was no change in Δ*abaIR* mutant. One study found that these genes are essential affect growth in LB medium (26). We then assessed the growth characteristics and morphology of the mutants, as shown in Figure 1A, 1B, Δ*abaI* mutant showed slightly slowed growth at the logarithmic phase, the cytoplasm of the Δ*abaI* mutant appeared to be transparent, and the cytoplasmic density of Δ*abaI* mutant is relatively low. These genes play a significant role in mediating cell growth and energy metabolism in *A. baumannii*. Apart from NADH dehydrogenase, other genes that may be associated with cell growth and energy metabolism may be regulated by quorum sensing. In Δ*abaI* mutant, a subset of genes involved in benzoate degradation (AUO97_RS17065-17085) and tryptophan metabolism and limonene and pinene degradation were up-regulated, this strain may utilize the beta-ketoadipate pathway and tryptophan for energy supply (Fig. 10A).

**Figure 10.**
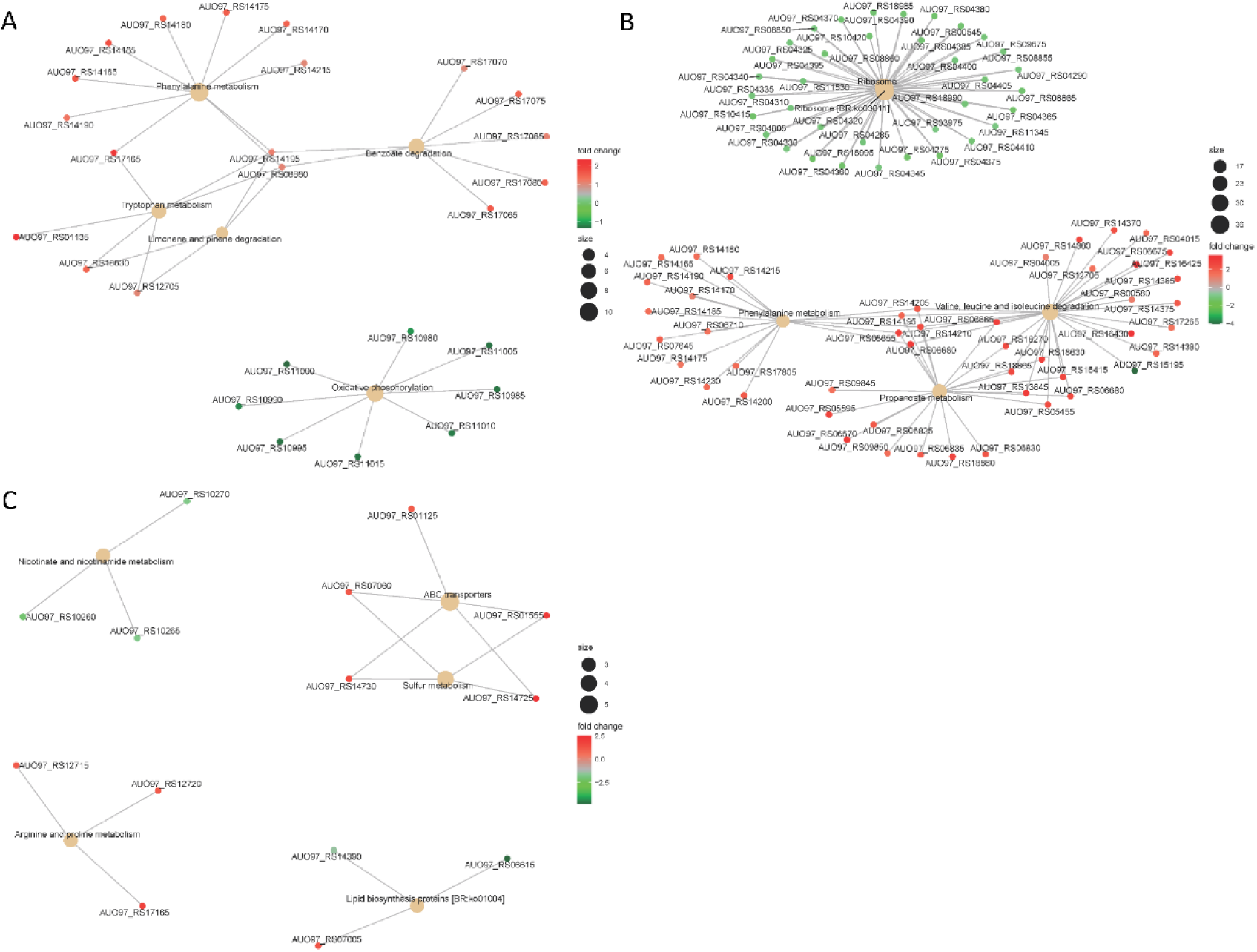
A) The top 5 of significantly enriched KEGG pathway of DEGs in Δ*abaI*. The size of the nodes shows the number of genes enriched in each pathway. The color gradient shows log2-fold change of gene expression. B) The top 5 of significantly enriched KEGG pathway of DEGs in Δ*abaR*. The size of the nodes shows the number of genes enriched in each pathway. The color gradient shows log2-fold change of gene expression. C) The top 5 of significantly enriched KEGG pathway of DEGs in Δ*abaIR*. The size of the nodes shows the number of genes enriched in each pathway. The color gradient shows log2-fold change of gene expression.

The phenylacetic acid (PAA) catabolic pathway encoded by *paa* operon is a key route in the catabolism of the Krebs cycle and this pathway is thought to contribute to bacterial virulence (27, 28). The cluster is composed of 15 coding sequences (*paaZ*, *paaA*, *paaB*, *paaC*, *paaD*, *paaE*, *paaF*, *paaG*, *paaH*, *paaJ*, *paaK1*, *paaK2*, *paaX*, *paaY* and *paaI*). In Δ*abaR* mutant, the *paa* operon was highly expressed (Fig. 10B). In Δ*abaI* mutant, AUO97_RS14165 (*paaZ*), AUO97_RS14170 (*paaA*), AUO97_RS14175 (*paaB*), AUO97_RS14180 (*paaC*), AUO97_RS14185 (*paaD*), AUO97_RS14190 (*paaE*), AUO97_RS14195 (*paaF*), AUO97_RS14215 (*paaK1*) were highly expressed (Fig. 10A), and the expression of this operon was not changed in the Δ*abaIR* mutant. Branched chain amino acids (BCAAs), including leucine (Leu), isoleucine (Ile), and valine (Val), is vital to both growth and virulence in bacteria (29, 30). In Δ*abaR* mutant, the Valine, leucine and isoleucine degradation pathway was up-regulated (Fig. 10B). The BCAAs serve as precursors for branched-chain fatty acids (BCFAs), which are predominant membrane fatty acids, the BCAAs are key co-regulators of virulence factors. In Δ*abaR* mutant, there are 27 genes involved in the propanoate metabolism pathway was up-regulated (Fig. 10B). In Δ*abaI* mutant, AUO97_RS14195, AUO97_RS16430, AUO97_RS06660, AUO97_RS18630 and AUO97_RS12705 involved in the propanoate metabolism pathway was up-regulated. In Δ*abaIR* mutant, AUO97_RS14380, AUO97_RS14375, AUO97_RS14370 involved in the propanoate metabolism pathway was up-regulated. Lipids play an important role in both the physiology and pathophysiology of living systems (31). Membrane phospholipids play a key role in the defense against antimicrobials, including host fatty acids (32–34).The COG enrichment analysis indicated that a large amount of genes with predicted functions in [I] Lipid transport and metabolism (21%), were up-regulated in Δ*abaR* mutant (Fig. 8). PAA, BCAAs and Lipid transport and metabolism may play a protective role against the host and serum. The K locus regulates the production of complex polysaccharides to protect against killing by host serum and enhance virulence in animal models of infection (35). In the K locus (O-glycosylation and wzy-dependent capsule synthesis locus) (AUO97_RS06870–06965), the gene AUO97_RS06875 (UDP-glucose 4-epimerase GalE) showed weakened transcription in the Δ*abaI* and Δ*abaIR* mutants by a 1.7-fold decrease and a 1.6-fold decrease, respectively, while there was no change in Δ*abaR* mutant. AUO97_RS13365 (*OmpA* family protein), a naturally glycosylated protein in *A. baumannii* ATCC 17978, was expressed with a 1.8-fold increase in Δ*abaR* strain, and no difference in Δ*abaI* and Δ*abaIR* mutants was observed. Ribosomal proteins (RPs) are well-known for their role in mediating protein synthesis and maintaining the stability of the ribosomal complex, which includes small and large subunits. There were 36 genes encoding for ribosomal proteins that exhibited reduced expression in Δ*abaR* mutant strain (Fig. 10B). 23 (L1-6, L9-13, L15-20, L22-23, L25, L28-29, L35) of them were associated with the large subunit while the remaining 13 (S2-8, S10-11, S14, S17-19) were associated with the small subunit. This change may enable *A. baumannii* to ‘fine-tune’ their proteomes to regulate the pathogenicity of bacteria. There was no change in Δ*abaI* and Δ*abaIR* mutants. We assessed the virulence of the mutant in *G. mellonella* and mouse and serum sensitivity tests. As shown in Figure 5B and Figure 6 and Figure 5A, Δ*abaR* mutant enhances more virulence and serum resistance, while Δ*abaI* and Δ*abaIR* mutants markedly attenuated the virulence of *A. baumannii*. The selected genes (*paaG*, *paaH*, *paaJ*, *paaK2*, *paaX*, *paaY* and *paaI*) encode proteins in the PAA catabolic pathway, BCAAs, the capsule synthesis locus *GalE* and *OmpA* may contribute to virulence.

## DISCUSSION

*A. baumannii* has become a very important hospital-acquired pathogen. Bacterial virulence is the prime determinant for the deterioration of an infected patient’s health. Quorum sensing (QS) is a cell-to-cell communication system utilized by bacteria to promote collective behaviors. Many bacteria use quorum sensing (QS) to control virulence.

In the present study, we focused on detecting the role of the *abaI/abaR* QS system in the virulence of *A. baumannii*. The mutant lacking *abaI* is believed to be less virulent than wild-type strain. In contrast, *abaR* mutants were significantly more pathogenic than wild-type strain. This result was confirmed in our study by injection of *G. mellonella* larvae and a mouse model of infection and serum killing test. Our transcriptomic analysis results revealed that deletion of *abaI* leads to the significant repression of energy production and conversion genes. The connection between energy metabolism and virulence has been reported in a multitude of bacteria. In *Vibrio cholerae*, the expression of virulence regulatory protein *ToxT* is affected by the NADH via respiration activity (36). In *Pseudomonas savastanoi*, *RhpR* directly regulate multiple metabolic pathways and phosphorylation to specifically control virulence (37). Therefore, *abaI* may indirectly control bacterial virulence by inducing the differential expression of some key genes involved in NADH dehydrogenase in the respiratory chain. Deletion of *abaR* enhances more cytotoxicity and immune evasion. RNA-Seq analysis indicated that deletion of *abaR* leads to the significant overexpression of lipid transport and metabolism, carbohydrate transport and metabolism and amino acid transport and metabolism genes. Lipid metabolism plays a key role in the pathogenicity of some intracellular bacteria (38). It has been observed that lipids are the main carbon and energy source of *M. tuberculosis*, which switches from carbohydrate utilization to the fatty acid utilization pathway for the establishment of a successful infection(39). *A. baumannii* is a ubiquitous, facultative intracellular bacterial pathogen (40). Δ*abaR* mutant may enhance lipid transport and metabolism to plays a protective role against the host. Δ*abaIR* mutant was slightly more pathogenic than Δ*abaI* mutants but less pathogenic than the Δ*abaR* mutants. This result was verified in our study by injection bacteria into *G. mellonella* larvae. Virulence through the intermediated phenylacetate catabolism pathway has been found in *A.baumannii,* and deletion of *paaE* attenuated *A. baumannii* virulence in mouse septicemia model (27). In *Burkholderia cenocepacia*, *paaA* and *paaE* insertional mutants showed reduced virulence, and interruption of *paaZ* and *paaF* slightly increased virulence in the *Caenorhabditis elegans* model of infection(41). Therefore, deletion of the *abaR* gene may indirectly enhance bacterial virulence via triggering the differential expression of a lot of key genes involved in the phenylacetate catabolism pathway. The selected genes (*paaG*, *paaH*, *paaJ*, *paaK2*, *paaX*,*paaY* and *paaI*) encode proteins in the PAA catabolic pathway that may contribute to virulence.

*abaI* is a protein that synthesized acyl-homoserine lactones (AHLs), *abaR* is a LuxR homolog transcription factor/receptor for AHLs (42). In this study, only WT strain was observed to produce AHLs based on *A. tumefaciens* KYC55 reporter strains, while the green pigment was not observed in Δ*abaI*, Δ*abaR* and Δ*abaIR* mutants. No purple pigment was observed in all strains based on *C. violaceum* CV026. In previous study, the absence of purple pigment may be attributed to low rate of the production of short chains homoserine lactone and fast degradation in the strains (43). Our transcriptomic analysis results revealed that deletion of *abaR* leads to the significant repression of *abaI.* Therefore, *abaR* may be a repressor such that repression is relieved when AHLs are bound. The results suggest that *abaR* generally represses its regulon of genes until it binds AHLs. When AHLs are bound, then repression is relieved. This may explain why deletion of abaI is substantially different from deletion of *abaR*. When *abaI* is mutated, then *abaR* still represses many different genes expression.

Interfering with quorum sensing is known as ‘quorum quenching’ and it will attenuate the virulence of the organisms (44). Many strategies for quorum-quenching have been proposed including targeting AHL synthase enzyme, the AHL binding receptor and the AHL itself (44). But previous studies on bacterial quorum quenching mainly focused on the influences on biofilm formation and motility and antibiotic resistance (12, 13). Our work provides a new insight into *abaI/abaR* quorum sensing system effects pathogenicity in *A. baumannii*. We propose that targeting the AHL synthase enzyme *abaI* could provide an effective strategy for attenuating virulence. On the contrary, interdicting the autoinducer synthase– receptor *abaR* elicits unpredictable consequences, which may lead to enhanced bacterial virulence.

## MATERIALS AND METHODS

### Bacterial strains, plasmids and culture conditions

Bacterial strains and plasmids used in this study are listed in Table 1. *A. baumannii* strains were grown in lysogeny broth (LB). Antibiotics were used at the following concentrations for *Escherichia coli*: kanamycin, 10 mg/L; ampicillin 25µg/mL and tellurite, 6 mg/L. For *A. baumannii*: tellurite, 30 mg/L.

**Table 1.**
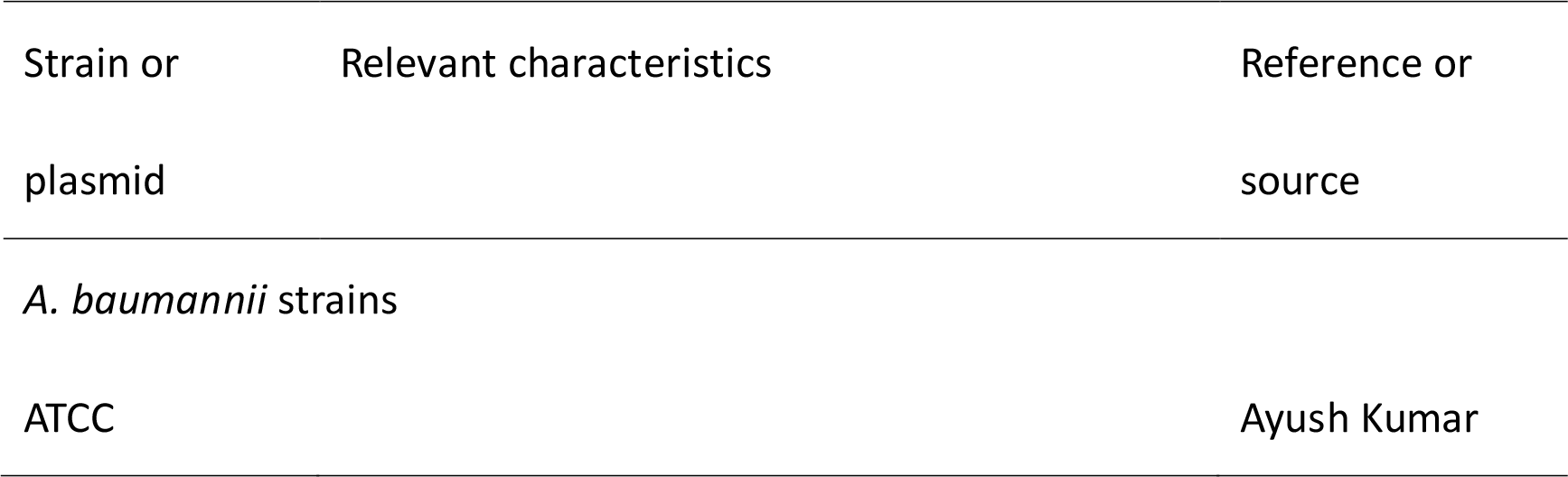
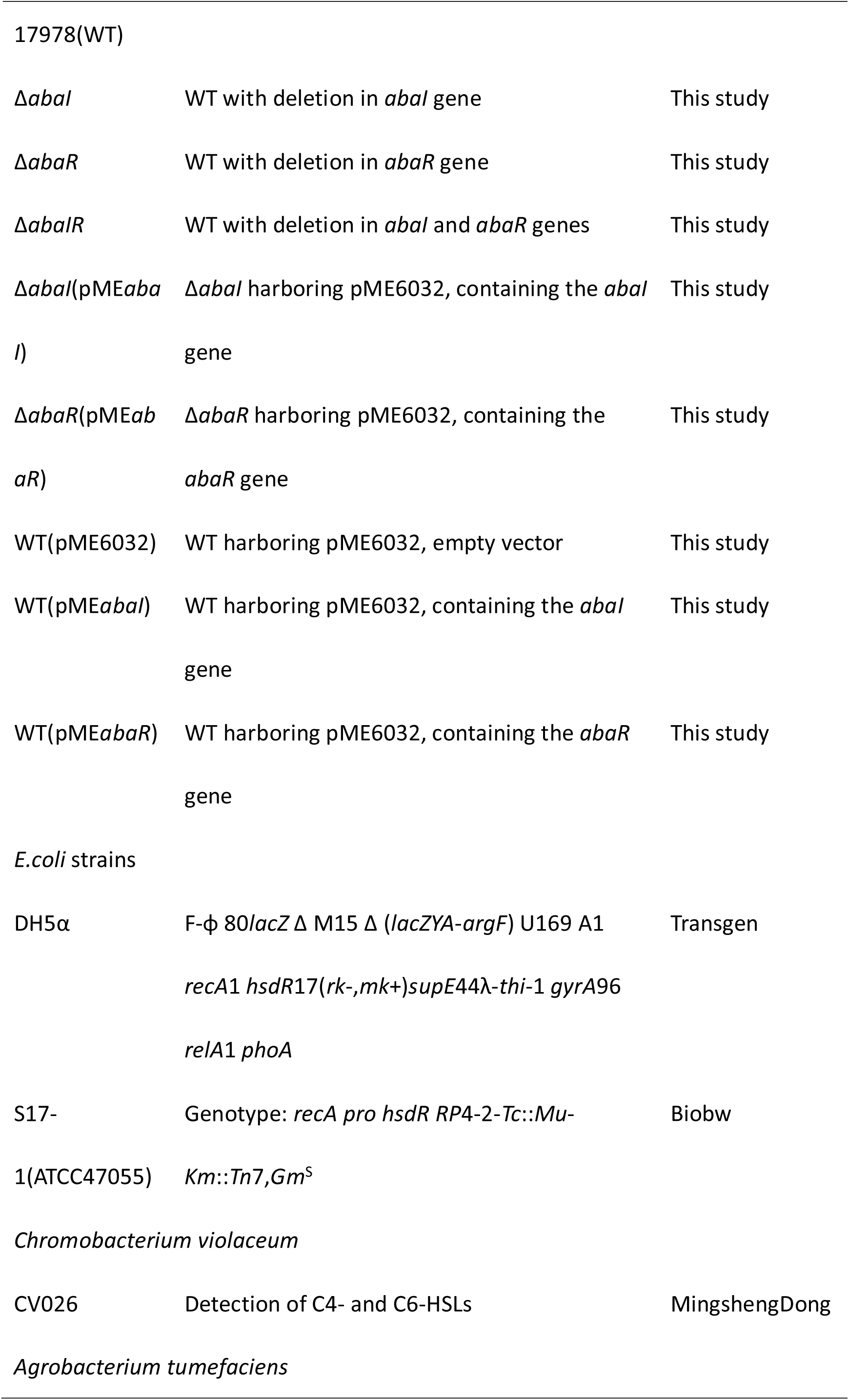
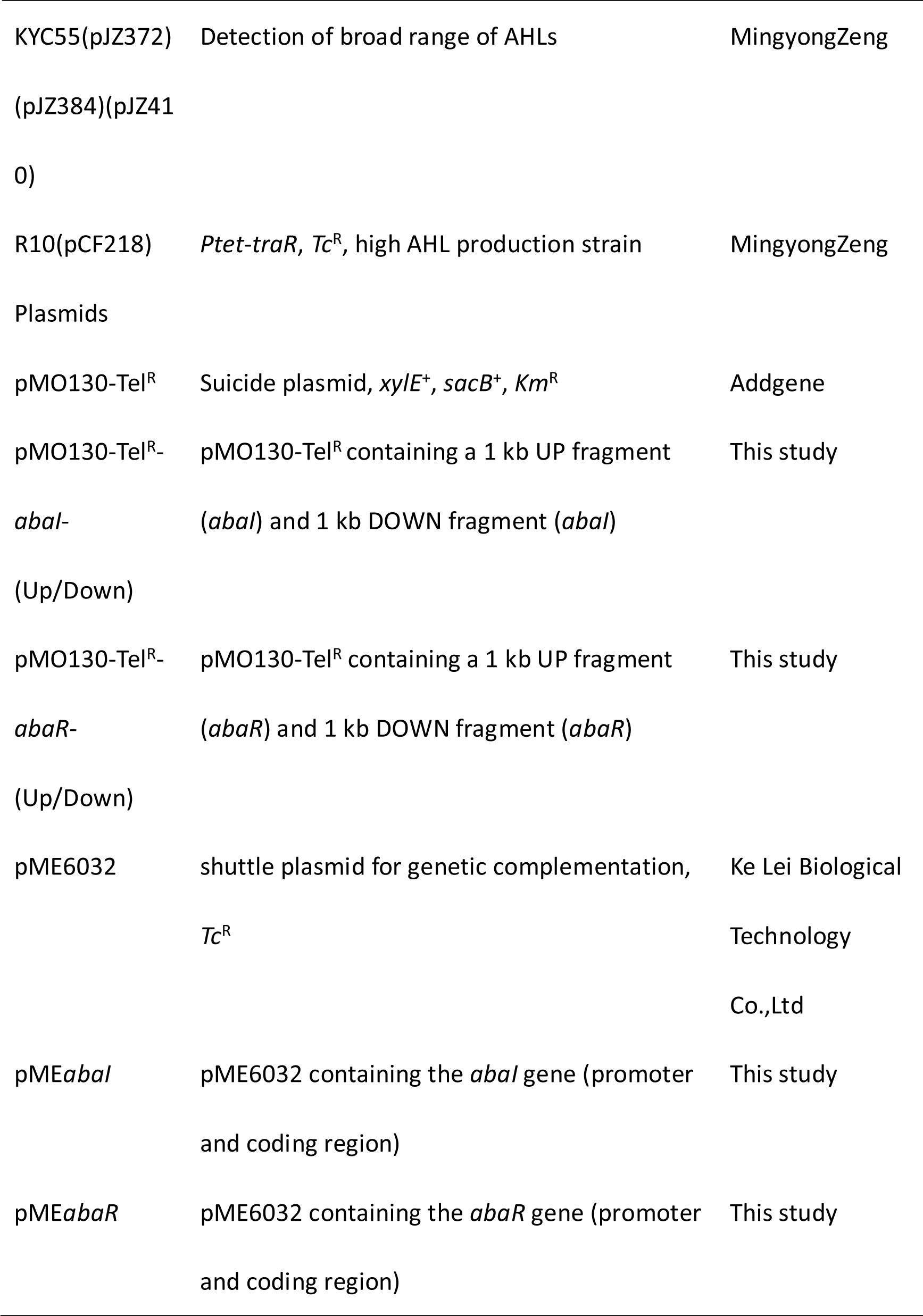
Bacterial strains and plasmids used in this study

### Strain construction

Strains Δ*abaI*, Δ*abaR*, Δ*abaIR* were unmarked deletion mutants created by a previous described method for acquiring marker-less deletions in *A. baumannii* with minor modifications (45). The primers used in this study are listed in Table 2. Briefly, the upstream and downstream homologous arms of the target gene were amplified and fused and then ligated into a tellurite-resistant suicide vector with the T4 ligase, pMO130-Tel^R^ (a generous gift from Addgene). The plasmid constructs were first introduced into *Escherichia coli* DH5α and subsequently selected on LB agar containing 30 µg/mL kanamycin. The kanamycin resistant colonies which carry a insertion of pMO130-Tel^R^ and a 2 kb amplimer corresponding to the size of the ligated upstream and downstream homologous arms of the target gene were tested by the corresponding designed primers. The resulting plasmids were used to transform into *E.coli* S17-1 and subsequently conjugate into *A. baumannii* ATCC 17978 via biparental conjugation. Exconjugants were selected on LB containing 30 µg/mL tellurite and 25 µg/mL ampicillin. *A. baumannii* ATCC17978 harboring the inserted pMO130-Tel^R^-Gene-(Up/Down) construct was cultured in LB broth containing 10% sucrose and passaged seven days to select for stabilized deletion of gene and loss of the sacB gene by a second cross-over and allelic replacement. If the target gene has been deleted, PCR of genomic DNA from these bacteria would not produce any amplimer using a primer pair that anneals to the DNA that has been deleted. Mutants were complemented with the pME*abaI* and pME*abaR* plasmids, generated by cloning the *abaI* and *abaR* genes and ligating into a shuttle plasmid vector with the T4 ligase, pME6032(46). Overexpressed strains were transformed by the pME*abaI* and pME*abaR* plasmids. The complemented strains and Overexpressed strains were confirmed by PCR and restriction analysis of plasmids extracted from *A. baumannii* cells grown in LB medium containing 50 μg/ml tetracycline.

**Table 2.**
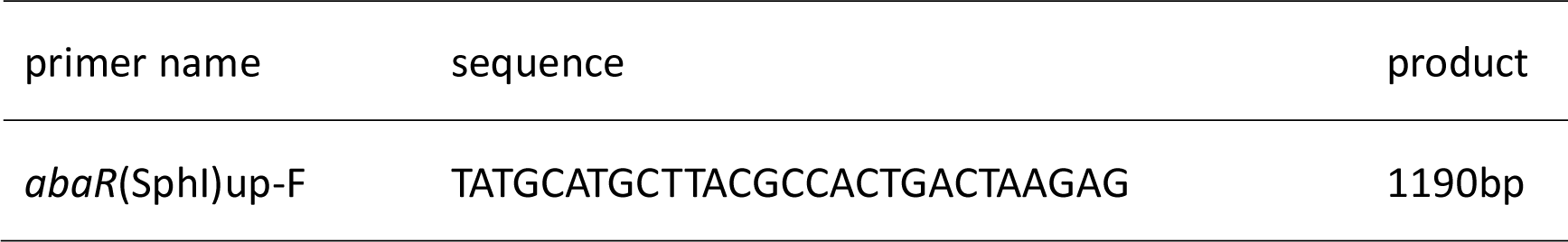
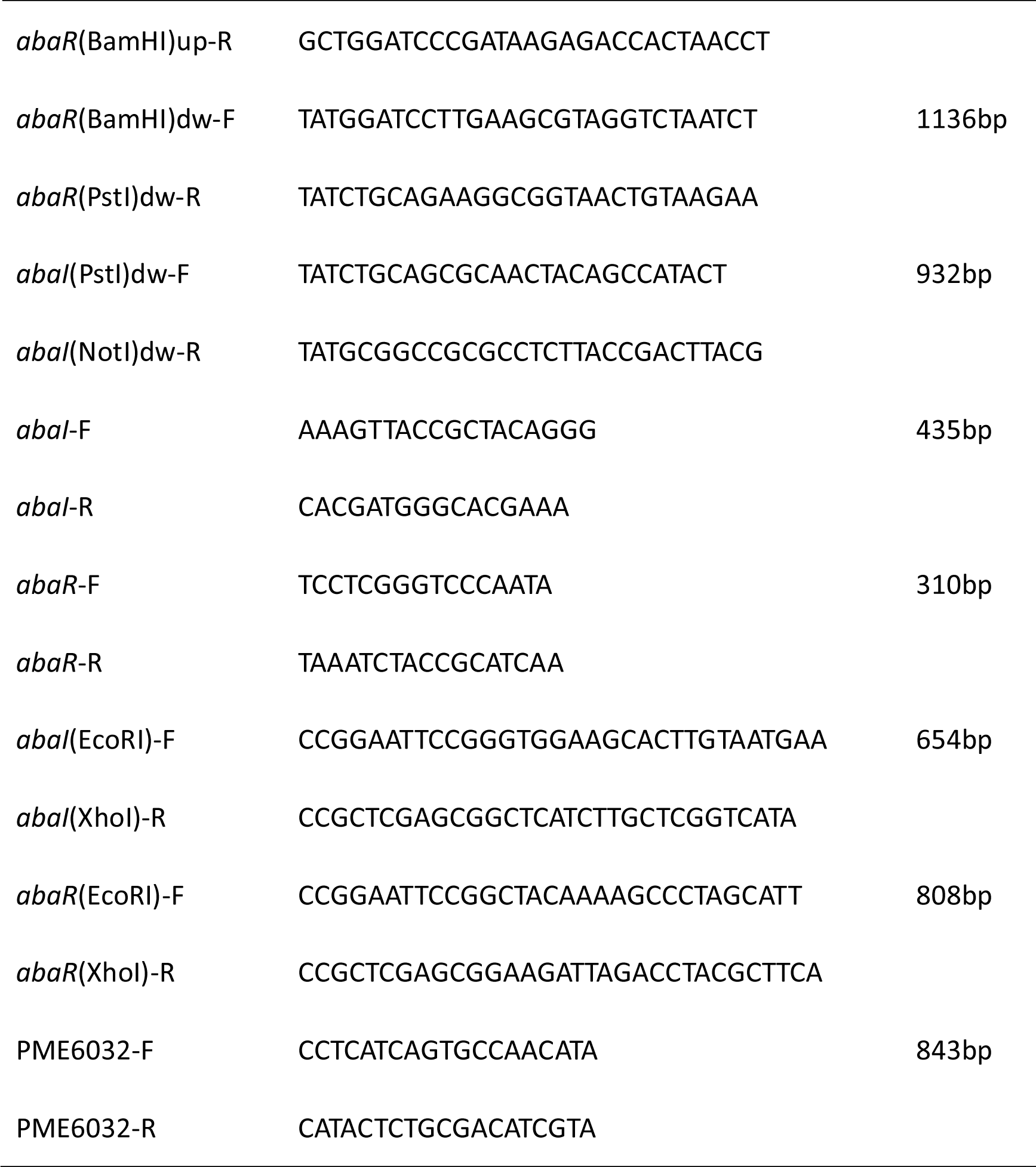
Primer used in this study

### Growth curve measurement

A single colony of strain was inoculated into 2 ml of LB broth, and cultured with shaking (200 rpm) overnight at 37°C.The bacteria were collected by centrifugation at 4000rpm for 5min and suspended in sterile saline to turbidity comparable to a 0.5 McFarland standard. 20 μL was pipetted into a 96-well microtiter plate containing 180 μL of LB broth and incubated at 37°C. The OD_600_ of cultures was measured at hourly intervals for up to 48 h to draw the growth curve. Tests were performed on eight individual biological replicates, in triplicates.

### Transmission electron microscopy (TEM)

Bacterial cells (OD450nm of 1.0) for SEM observation were harvested by centrifugation and washed three times with ddH_2_O. Bacteria were prefixed with 2.5% glutaraldehyde in 0.1 Mphosphate buffer (pH 7.4). Images were captured at 120 kv with a HITACHI H-7650 TEM.

### Antimicrobial susceptibility

*A. baumannii* strains were cultured in LB liquid medium at 37°C with shaking overnight. Minimum inhibitory concentrations (MICs) for Kanamycin, Penicillin, Streptomycin, Meropenem Imipenem, Ceftizoxime, Cefepime, Cefoperazone-Sulbactam, Piperacillin-tazobactam, Ampicillin, Tetracycline and Spectinomycin were determined on 96-well plates by the broth microdilution protocols of the Clinical and Laboratory Standards Institute, and the results of MICs testing were interpreted according to the criteria of the CLSI 2013 guidelines. All experiments were carried out a minimum of three times.

### Screening of AHL by *C. violaceum* CV026 and *A. tumefaciens* KYC55

*A. baumannii* can produce AHLs signal molecules with different chain lengths (43), it is essential to use biosensors that detect a broad range of AHLs. *C. violaceum* CV026 specific for short-chain AHLs (C4-C6 AHL molecules) and *A. tumefaciens* KYC55 specific for long-chain AHLs (C8-C14 AHL molecules) were utilized for screening AHLs producing bacterial strains. By the presence of short-chain AHLs, CV026 produces purple pigments. By the presence of long-chain AHLs, a green color is observed for *A. tumefaciens* KYC55 (43). Screening of the *A. baumannii* for production AHLs was carried out by agar plate diffusion assay with minor modifications (43, 47). Briefly, QS reporter strains were cultured in LB agar plates containing the antibiotics kanamycin 20 μg/mL for *C. violaceum* CV026, spectinomycin 50 μg/mL and tetracycline 4.5 μg/mL for *A. tumefaciens* KYC55. The plates were incubated at 28 °C for 24 h. 40 μg/mL of X-gal as a visualizing agent was incorporated into LB medium used for *A. tumefaciens* KYC55. For determining the production of Acyl homoserine signal molecules, *A. baumannii* and mutants and biosensor *C. violaceum* CV026 and *A. tumefaciens* KYC55 strain were inoculated side by side in a way that they had a 0.5 cm gap between them. *C. violaceum* CV026 and *A. tumefaciens* KYC55 were assessed positive or negative according to the color changes in biosensor strain.

### Surface motility assay

The motility test was performed according to the method as describted previously with minor modifications (19). Briefly, the media used for surface motility assay was tryptone broth [10 g/liter tryptone (OXOID) and 5 g/liter NaCl] supplemented with 0.3% (wt/vol) Noble agar (BD). Plates were prepared and inoculated with bacteria from an overnight culture in LB agar (1.5%, wt/vol) plates at 37°C with a sterile toothpick. All assays were carried out in triplicate in a minimum of three independent experiments. After incubation at 30°C for 12–14 h, the zone of motility at the agar/Petri dish interface was observed.

### Crystal violet biofilm assay

The biofilm forming ability test was performed in accordance with the method as described previously with minor modifications (48). Briefly, a few single colonies were suspended in sterile saline to turbidity comparable to a 0.5 McFarland standard. The suspension was under vortex movement for 1 min; 20 μL was pipetted into a 96-well microtiter plate containing 180 μL of LB broth and incubated for 24 h at 37°C. For crystal violet staining, the wells were rinsed with PBS to exclude loosely adherent cells and then stained for 30 min with 200 µl of 1% crystal violet. The wells were then rinsed with water and dried at room temperature. The amount of biofilm was quantitated by destaining the wells with 200 µl of 33% acetic acid and then measuring the optical density (OD) of the solution in a microplate spectrophotometer set at 595 nm. Tests were performed on ten individual biological replicates, in triplicates. The differences between parent and mutant strains were calculated, and values returning a P value of <0.05 from a Student’s t test were taken as significant.

### Serum bactericidal assay

The serum resistance experiment was performed in accordance with the method as previously described with minor modifications (49). Briefly, 100 µl mid-log-phase *A. baumannii* culture (a bacterial titer of approximately 1×10^5^ CFU) was mixed with 900 µl of either normal human serum or heat-inactivated serum (heated at 56°C for 30 min). The mixtures were incubated at 37°C, and aliquots of 100 ml were removed from the culture at 1 h for the determination of bacterial counts. The number of surviving CFU was determined by plating in triplicate. The results were expressed as percent survival, with 100% being the number of viable bacteria grown on Brain Heart Infusion agar plates.

### Virulence in Galleria mellonella

*Galleria mellonella* has been known as a good model system to study *A. baumannii* pathogenesis (50). Survival of ATCC 17978 and mutants in *G. mellonella* was measured as previously described (50). An inoculum of 10^6^ CFU bacteria was injected into *G. mellonella* larvae. The experiment was performed on ten individual biological replicates, in triplicate. Statistical analysis was carried out with GraphPad Prism 6 to produce Kaplan-Meier survival curves. The statistical significance of differences between parent and Δ*abaI*, Δ*abaR* and Δ*abaIR* mutant strains survival curves were calculated with a log rank test. P values of <0.05 were considered significant.

### Murine model of pneumonia

8-10 week old BALB/C mice were obtained from Jilin University. Mice were kept in a sterile environment at Jilin University and maintained according to standard procedures. All research was conducted in complying with the institutional guidelines. The principles in the ARRIVE guidelines and the Basel declaration (http://www.basel.declaration.org) have been considered when planning the experiments. Models of pulmonary infection were performed as previously described (35). Briefly, infections were initiated by intraperitoneal injection of approximately 1.2×10^8^ CFU of bacteria suspended in 100 µl of PBS into groups of mice (10 mice per group for survival studies; 5 per group for analyses of bacterial counts).

### Quantitative bacteriology

To assess bacterial burden, the lung and spleen were aseptically operated and homogenized in 1 ml sterile of PBS using tissue homogenizers. 100 µl of the homogenates were cultured on Brain Heart Infusion agar plates to quantify the bacterial load of *A. baumannii* in the respective organs. **RNA sequencing and analysis**

RNA was extracted from three biological replicates of each strain with a GeneMark Total RNA Purification Kit. RNA libraries were prepared and sequenced with Illumina HiSeq2000 at the Beijing Genomics Institute (BGI). Sequences were mapped to the ATCC 17978 genome (accession No.cp018664.1) using Bowtie2. Differentially expressed genes were identified using the DESeq2 package (Bioconductor). Genes were deemed as differentially expressed if they presented a log2-fold change greater than 1 or less than −1 and P value (P-adj) was less than 0.05 in mutant strain compared to WT strain. Cluster of orthologous groups (COG) enrichment analysis was performed by dividing the percentage of genes up-or down-regulated for each category by the percentage of genes in that category across the whole genome (17, 51). Multiexperiment Viewer-version 4.9.0 was used to perform hierarchical clustering and heat map visualization(52). Gene Ontology (GO) and Kyoto Encyclopedia of Genes and Genomes (KEGG) pathway enrichment analysis, based on R software, were applied for the identification of pathways in which DEGs significantly enriched. The GO and KEGG pathway analysis of the DEGs was conducted through the ClusterProfiler package in R software (53). A P-value of <0.05 was considered to have statistical significance and to achieve significant enrichment. qRT-PCR was performed on the 7300 Plus Real-Time PCR System (Applied Biosystems) using a standard protocol from the FastStart Universal SYBR Green Master (Roche, Basel, Switzerland). Gene expression levels were quantified by using the 2^-ΔΔCt^ method with endogenous controls (16S). The Primers used for qRT-PCR assays are described in Table 3. All qRT-PCR assays were repeated 3×. The RNA-Seq data obtained in this study was submitted to Sequence Read Archive (SRA), BioProject ID PRJNA600672.

**Table 3.**
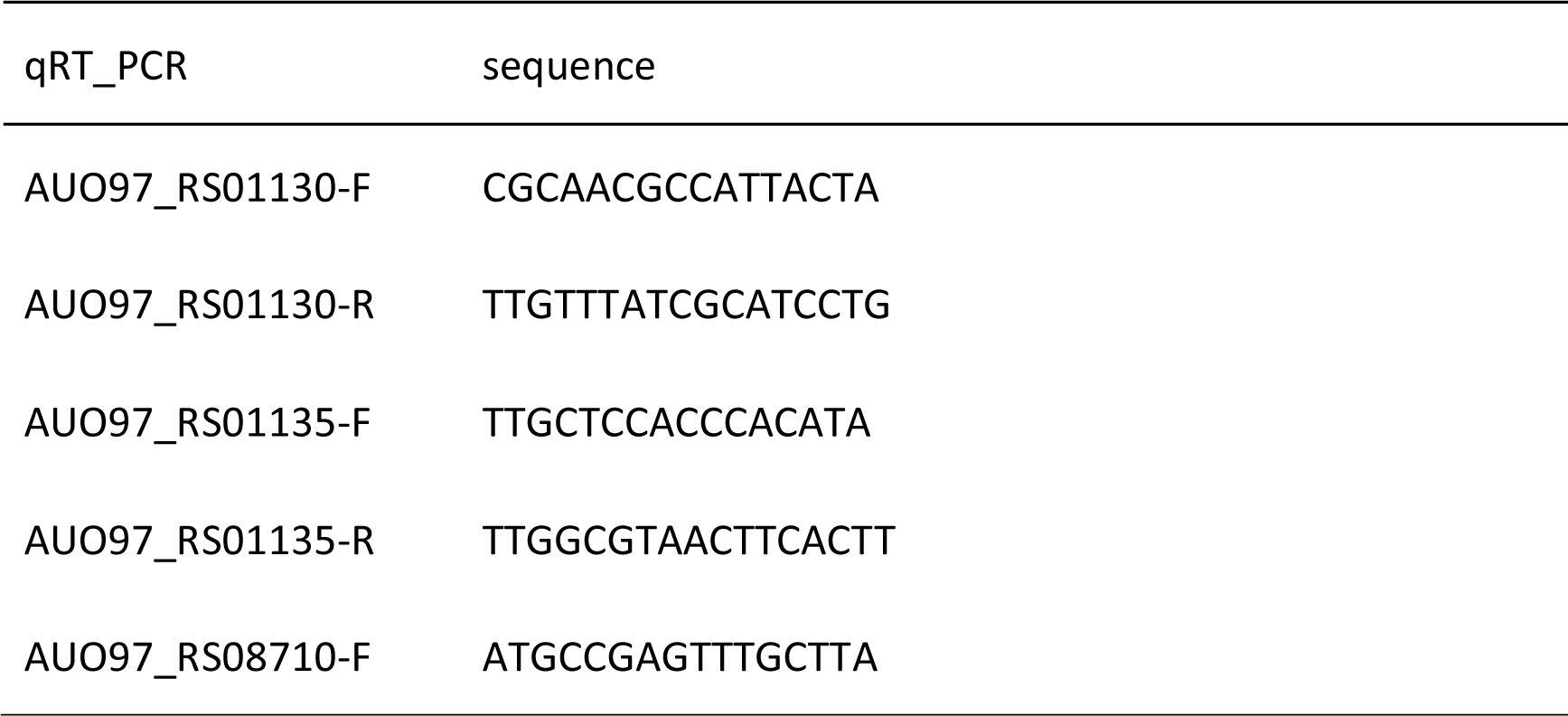
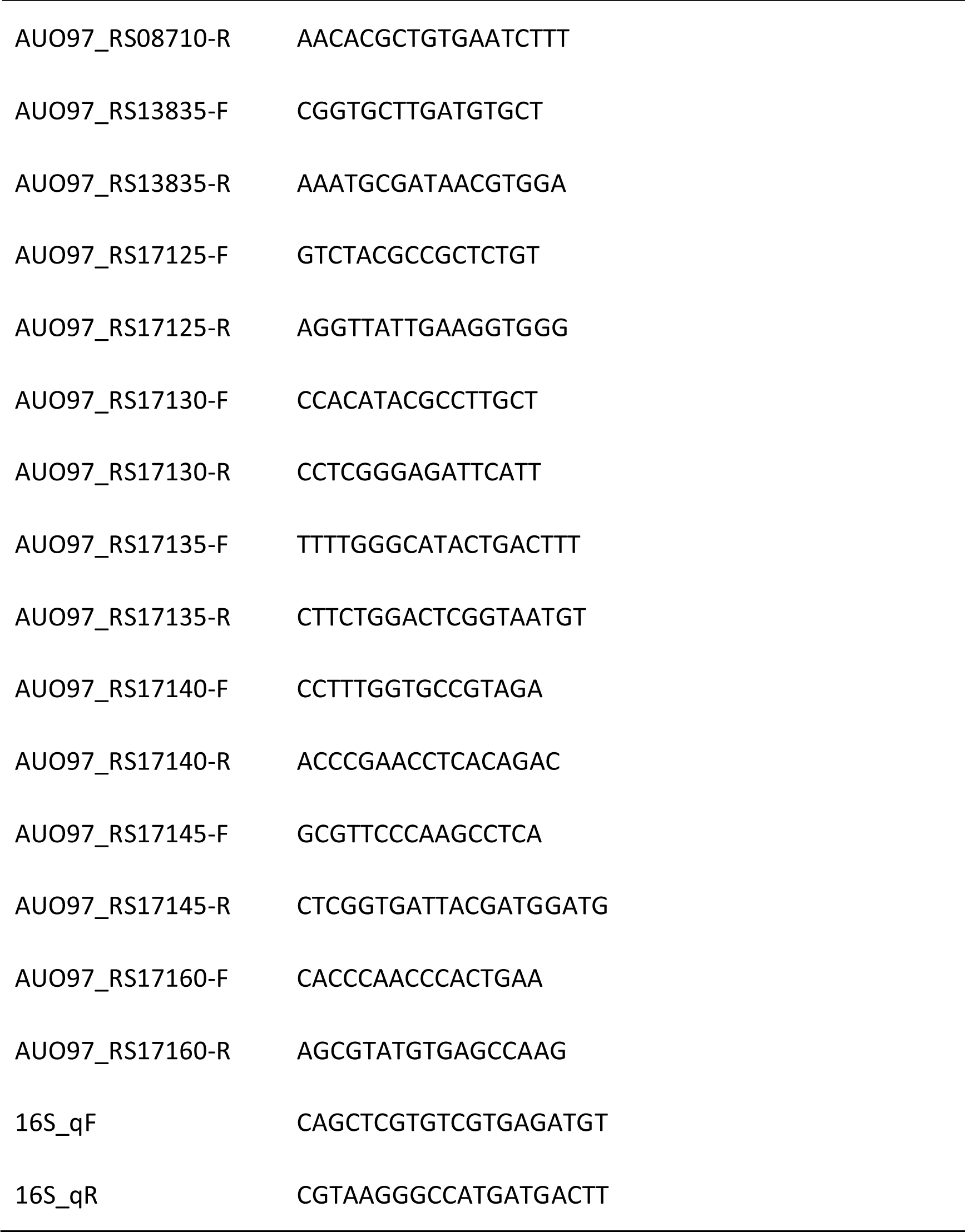
Real-time quantitative PCR primers

**Table 4.**
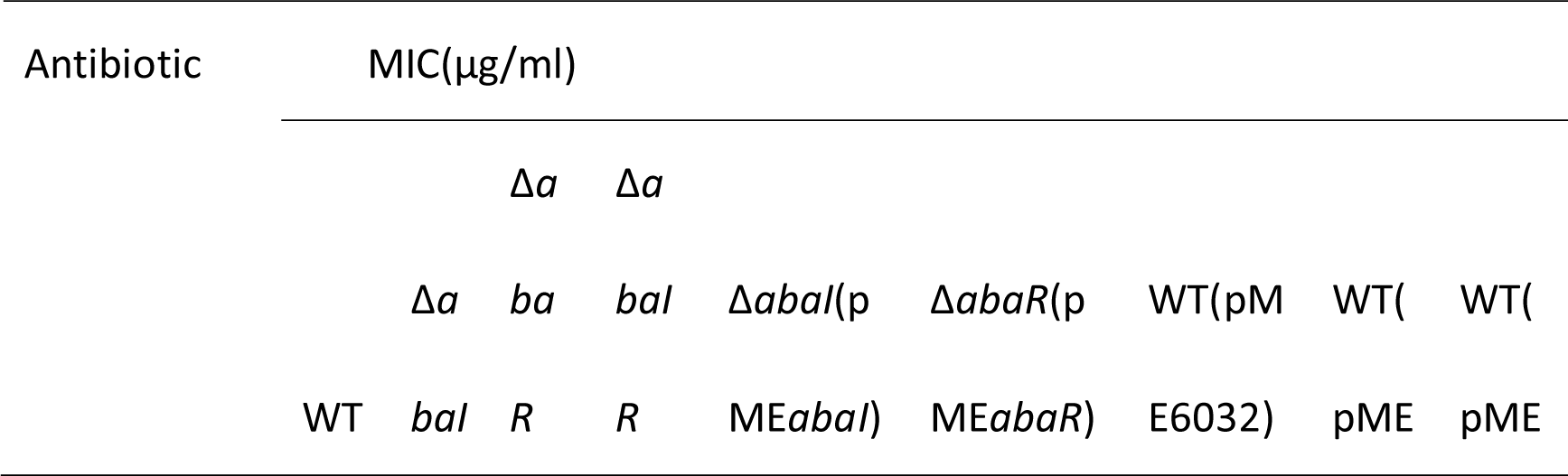
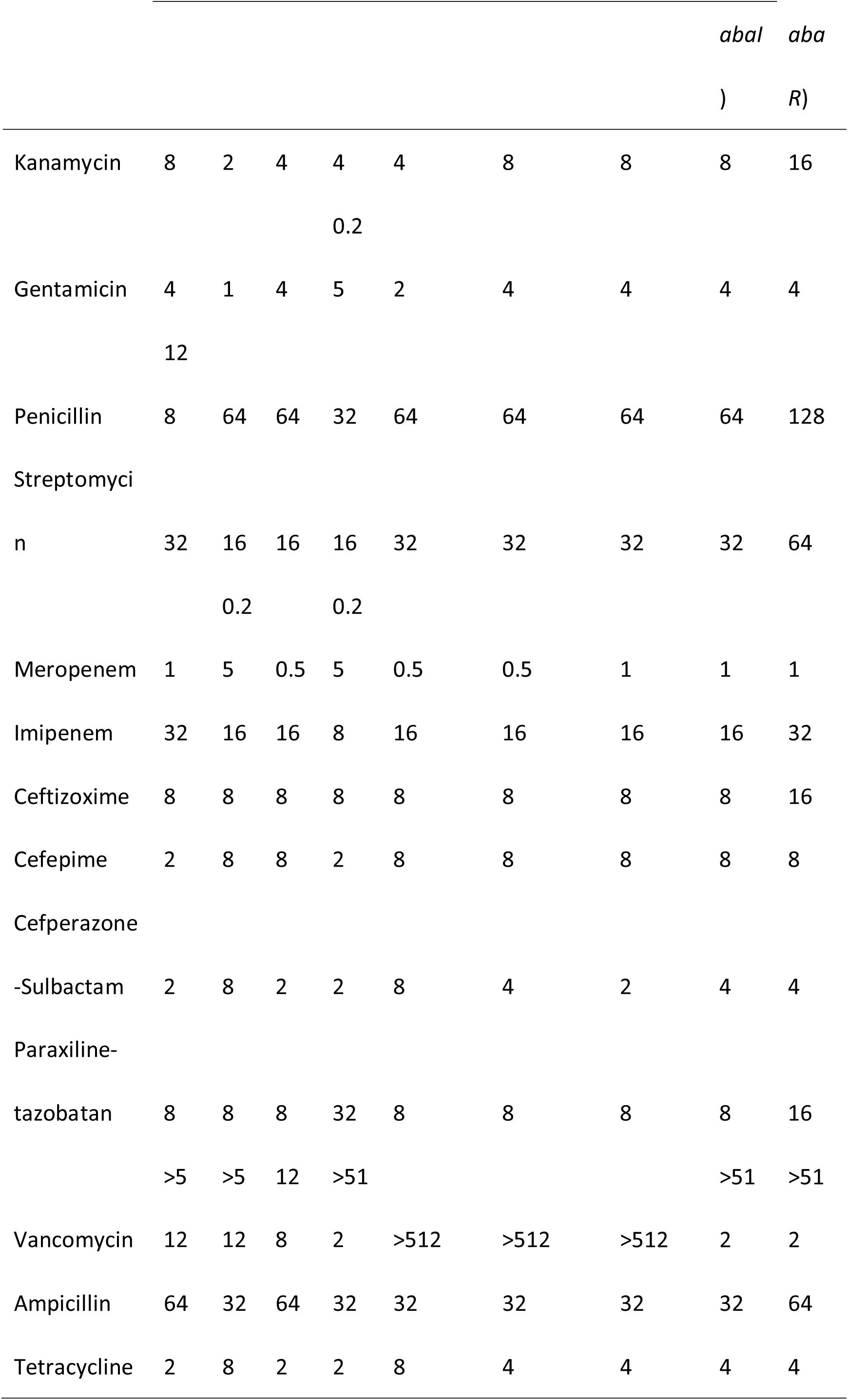
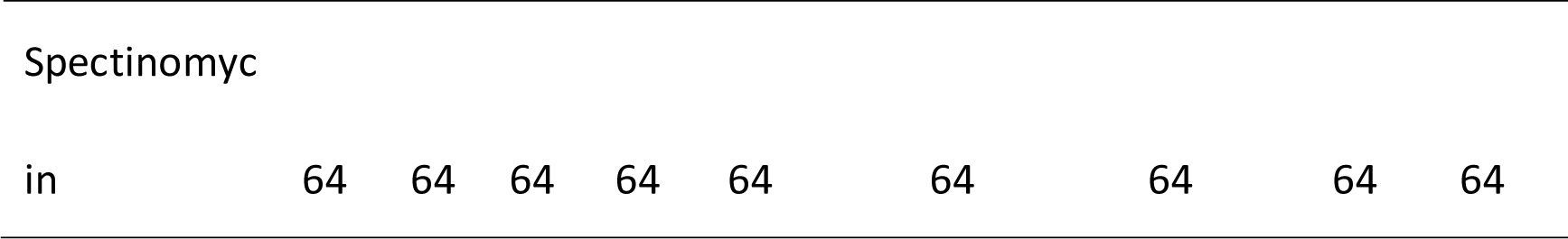
MICs of antibiotics used in this study

## ACKNOWLEDGMENTS

We thank Beijing Genomics Institute (BGI) for assistance with bioinformatics analysis. We appreciate Ayush Kumar, PhD professor from Department of Microbiology, University of Manitoba, Winnipeg, Canada, for *A. baumannii* ATCC 17978 strain; We appreciate MingshengDong, PhD professor from Nanjing Agricultural University for *Chromobacterium violaceum CV026* strain; We appreciate MingyongZeng, PhD professor from Ocean University of China for *Agrobacterium tumefaciens KYC55* and R10 strain; We thank the proofreading work of American Journal Experts (AJE) for this manuscript. This study was partially supported by grants from the National Natural Science Foundation of China (81601817 and 81672109), Jilin Province Development and Reform Commission (2015Y031-5), the Education Department of Jilin Province (JJKH20170820KJ, JJKH20170852KJ), and grants from Department of Finance of Jilin Province.

## Supplemental Material

**Table 5.**
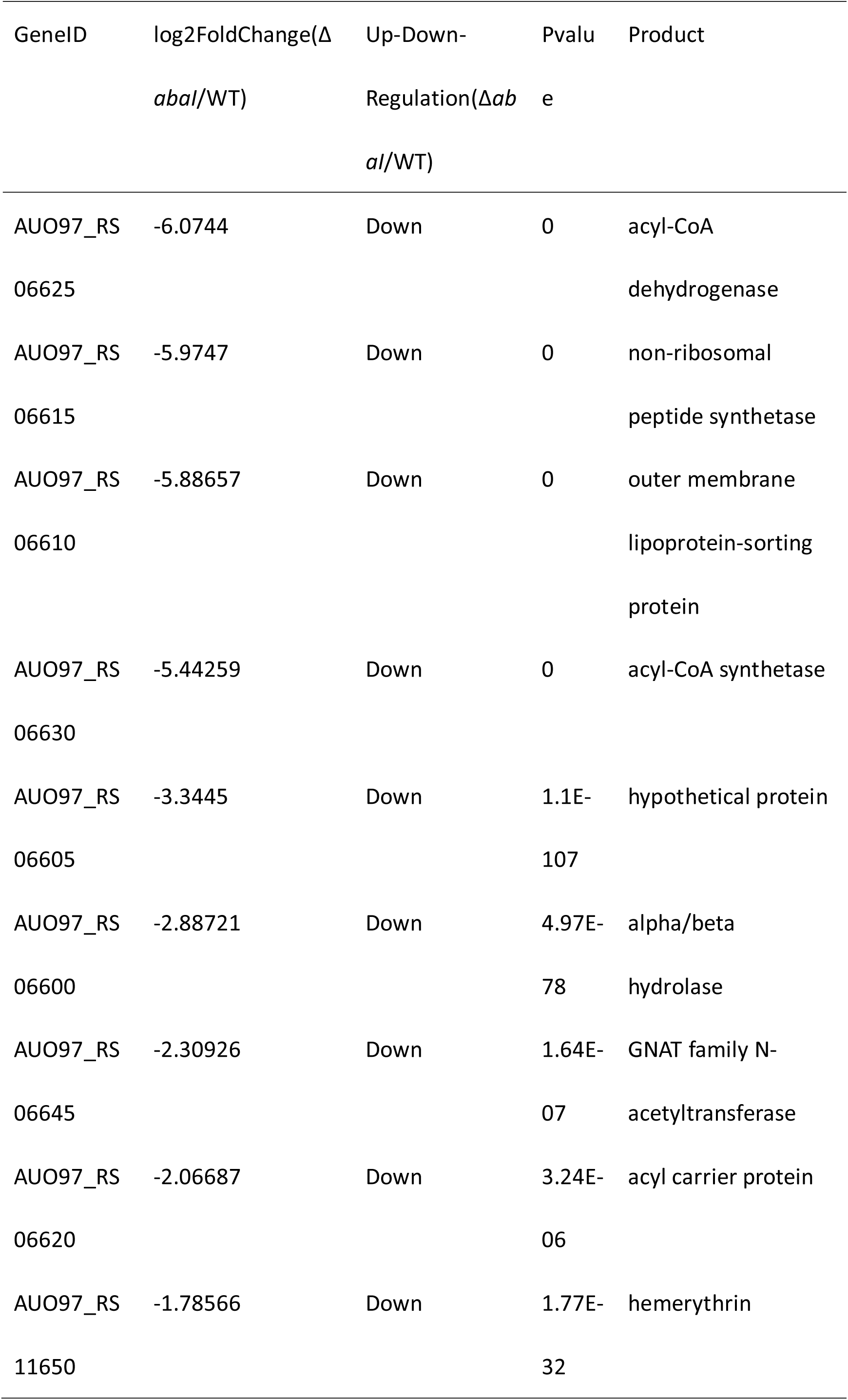
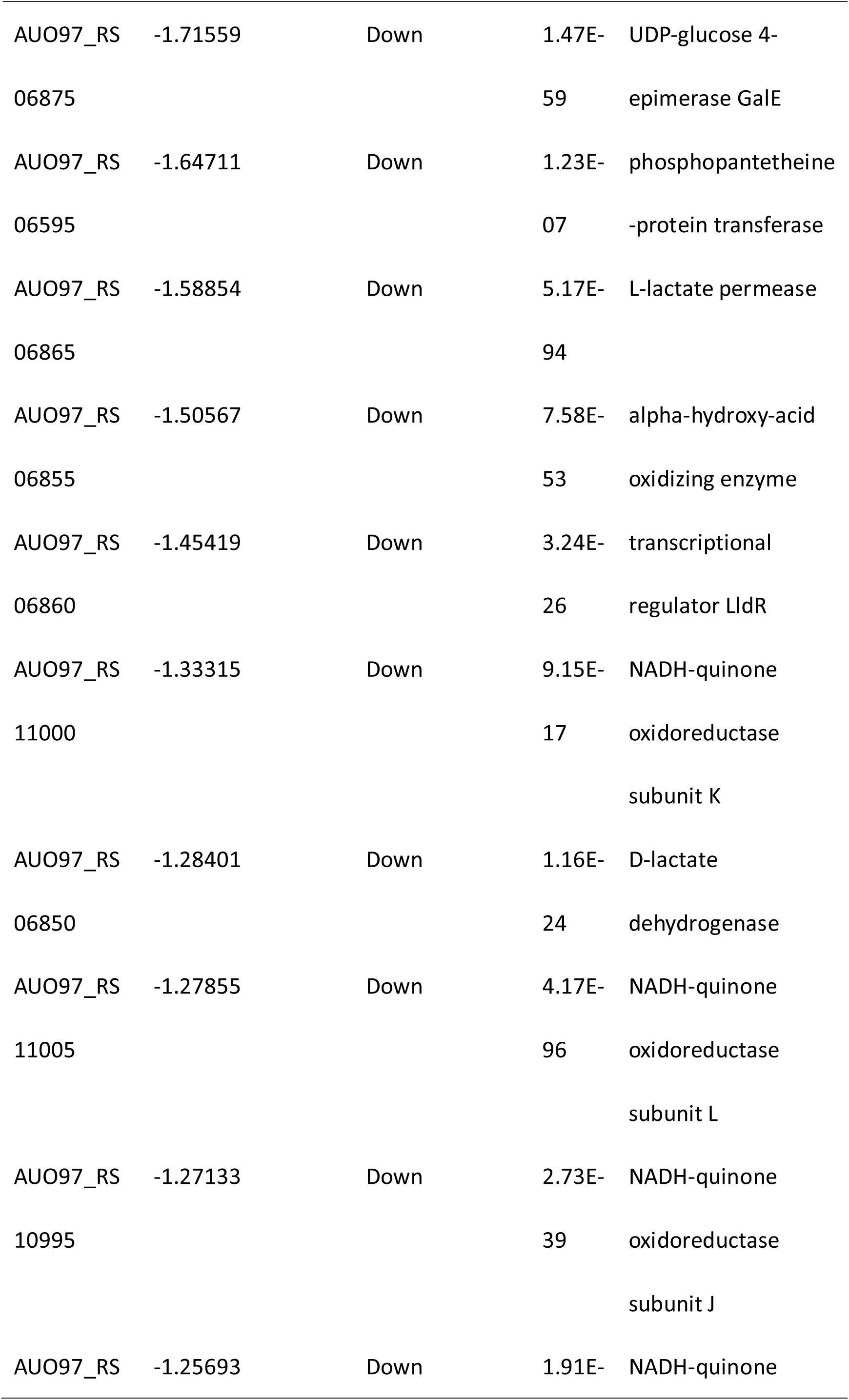
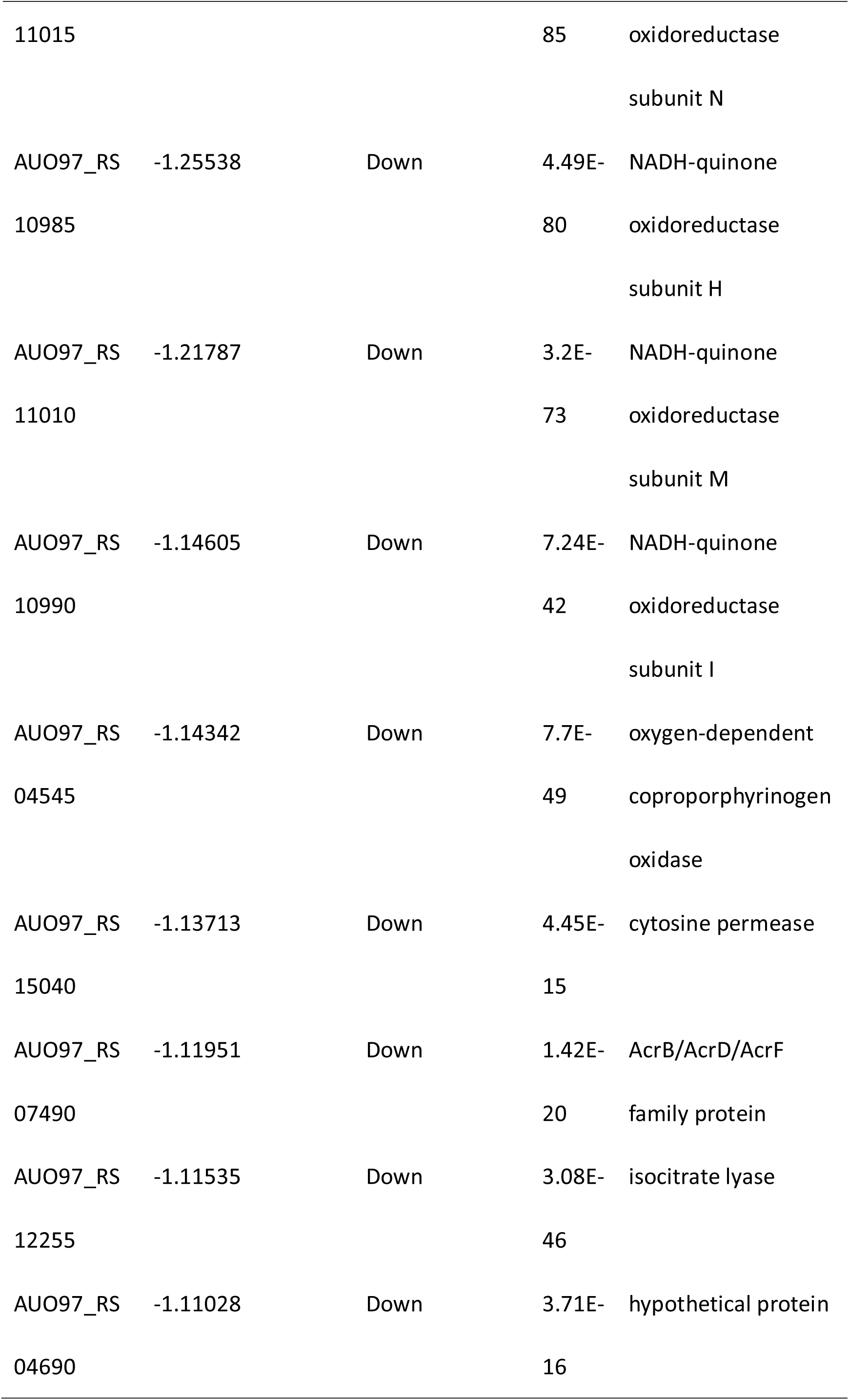
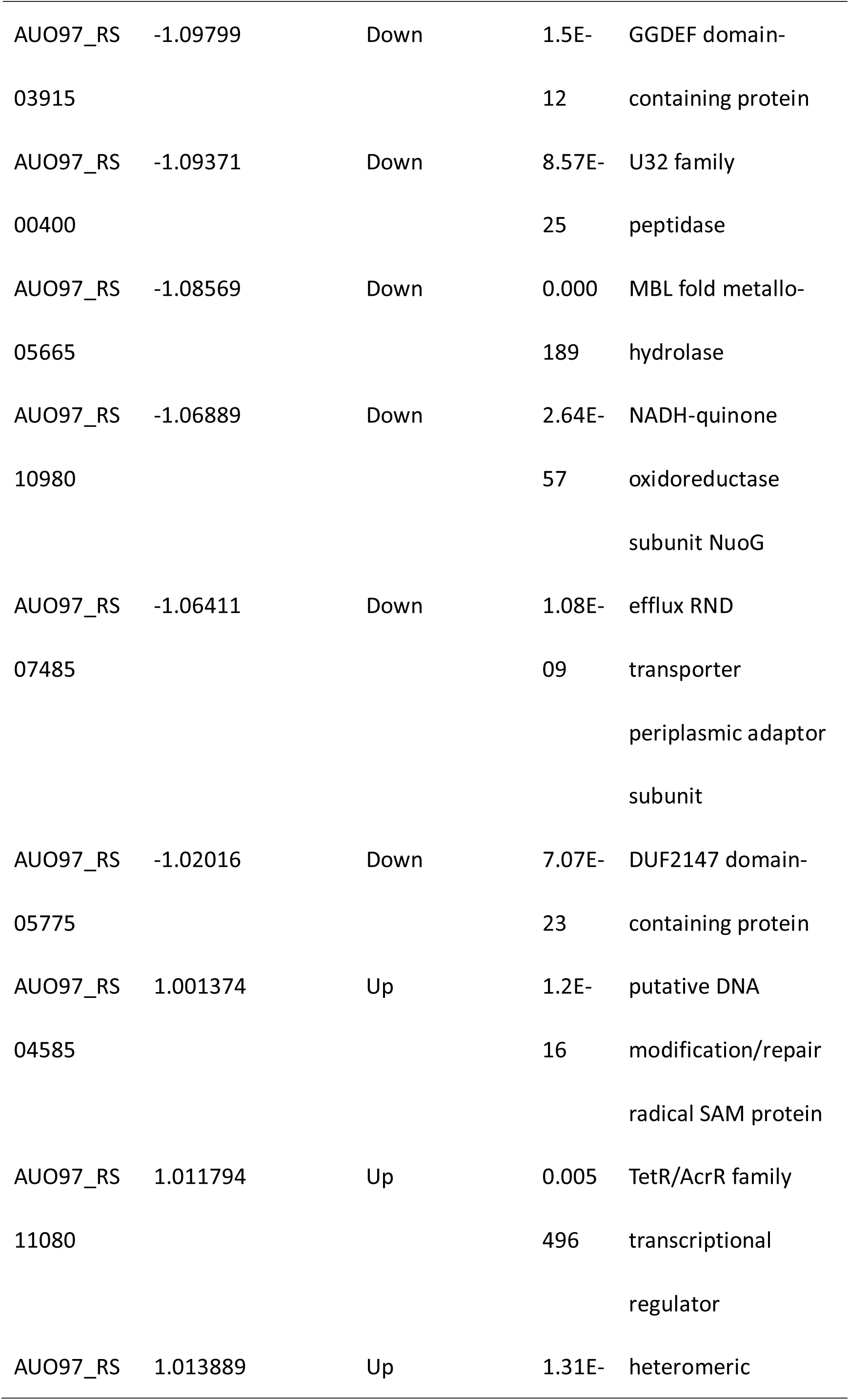
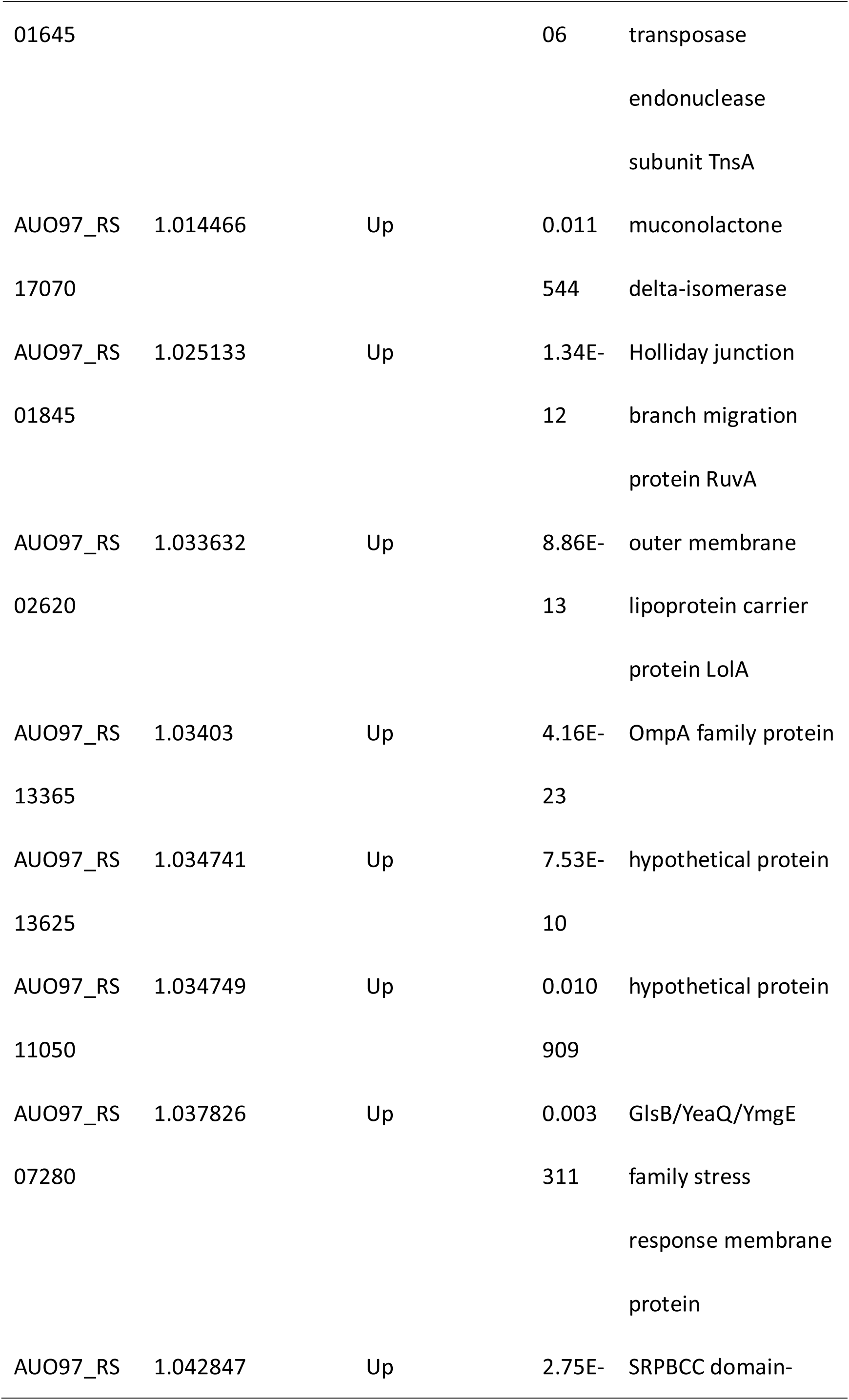
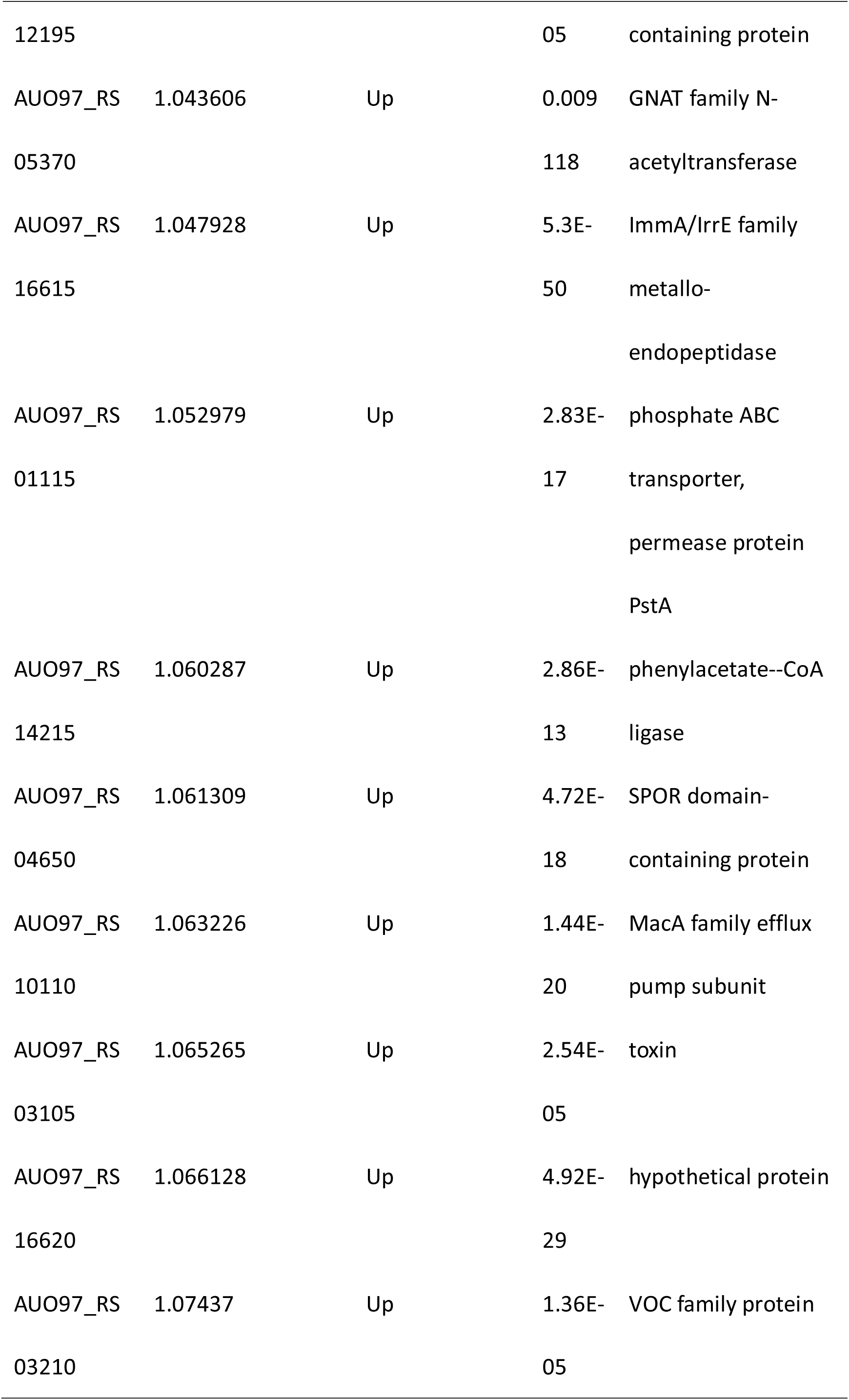
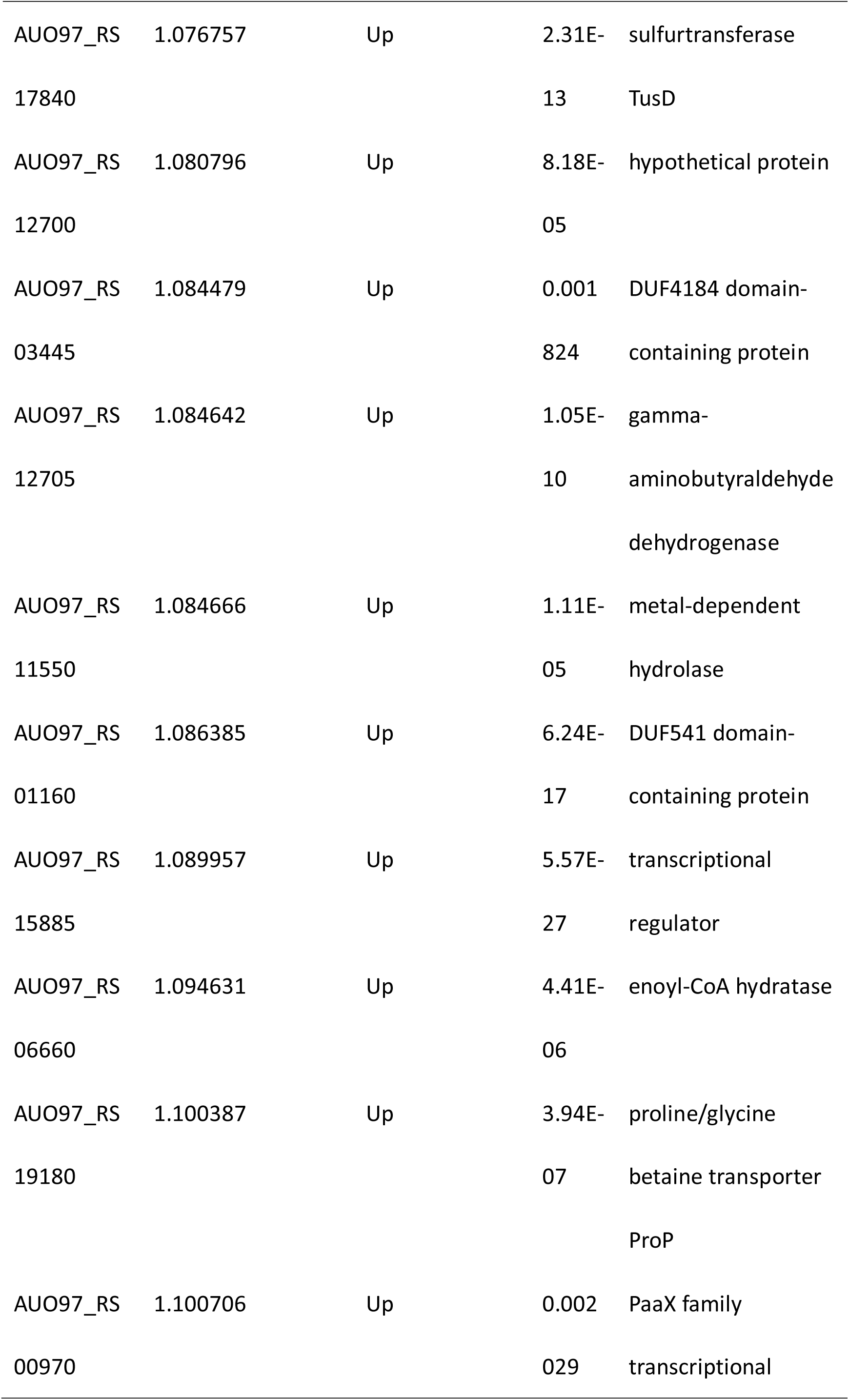
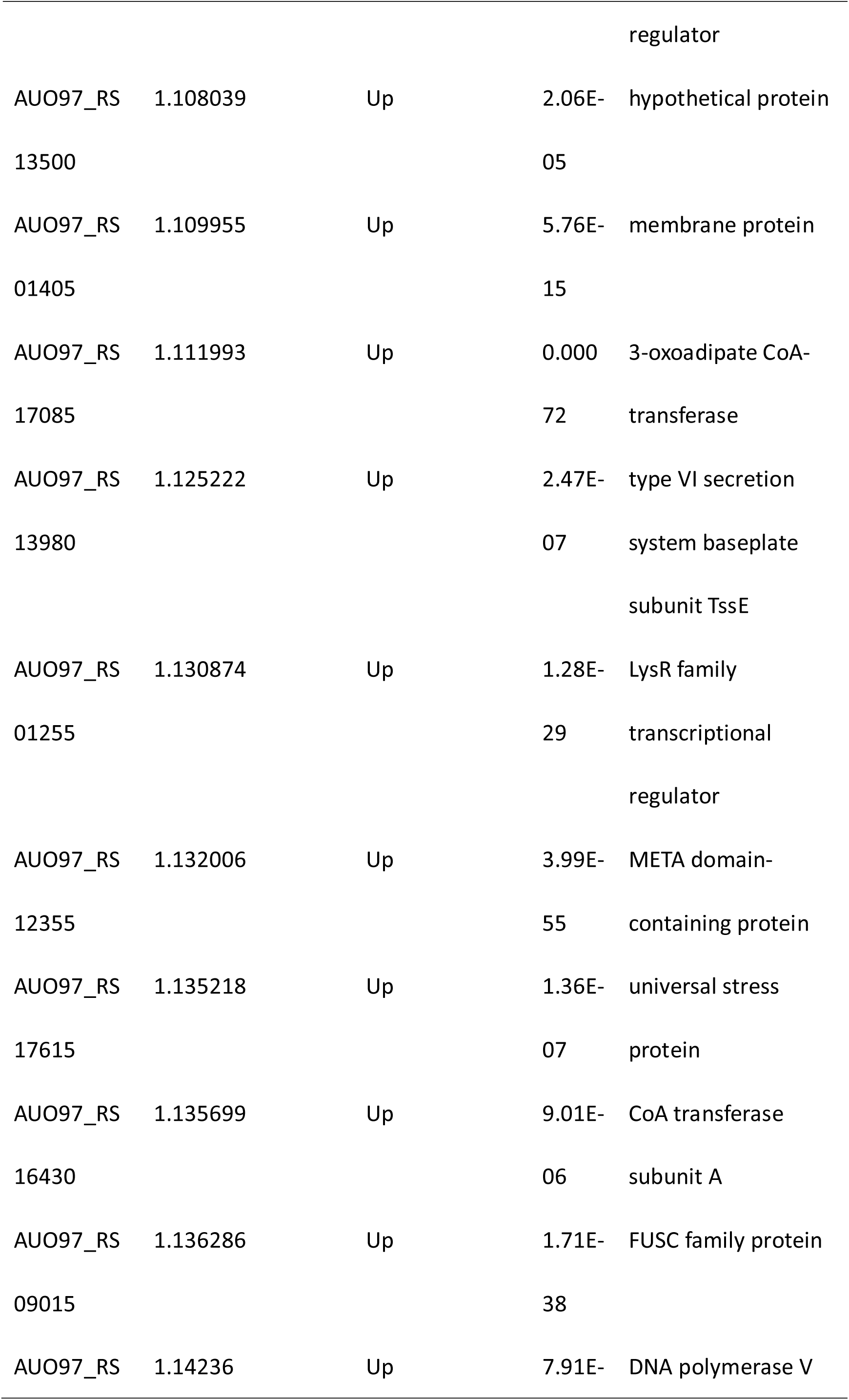
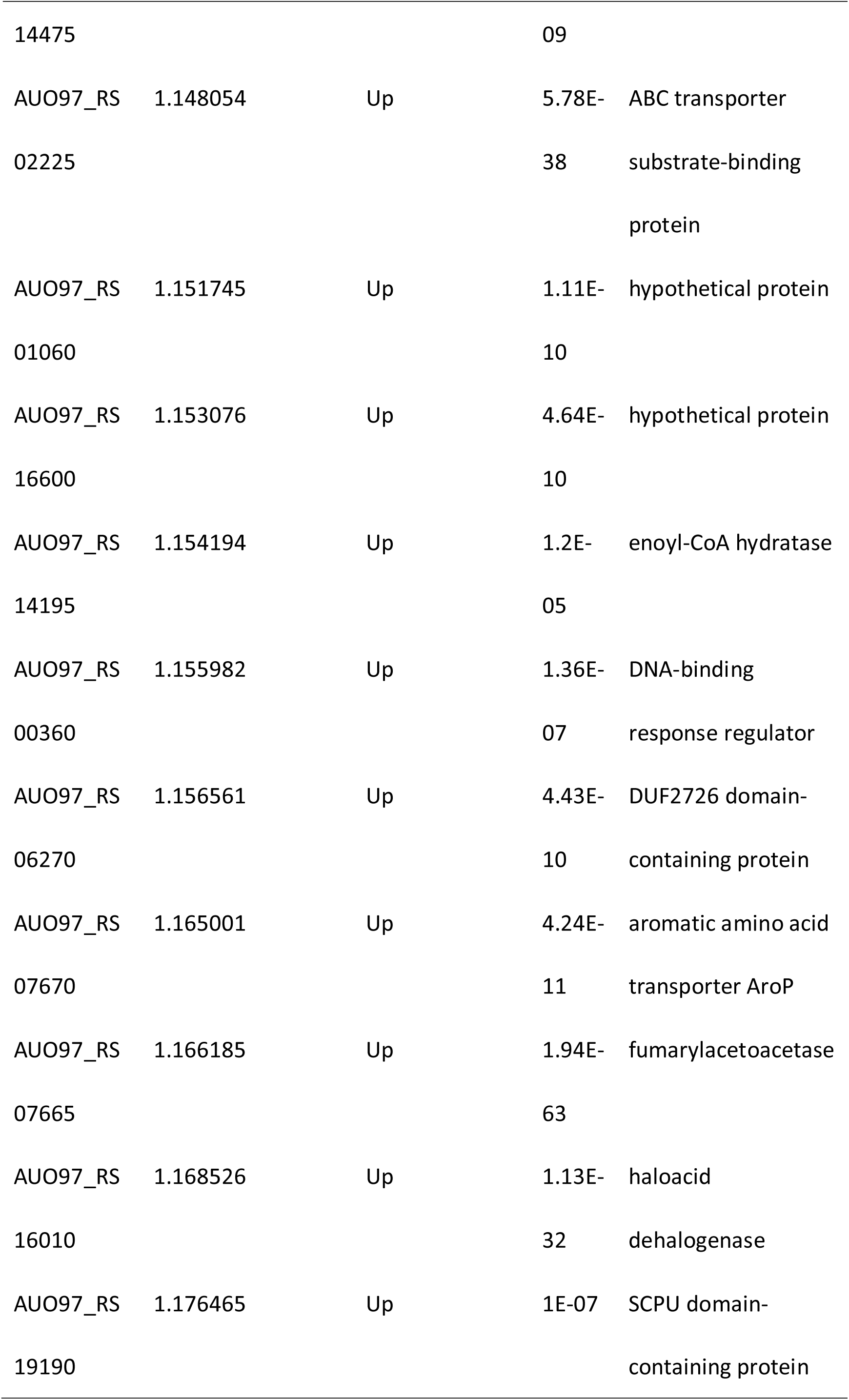
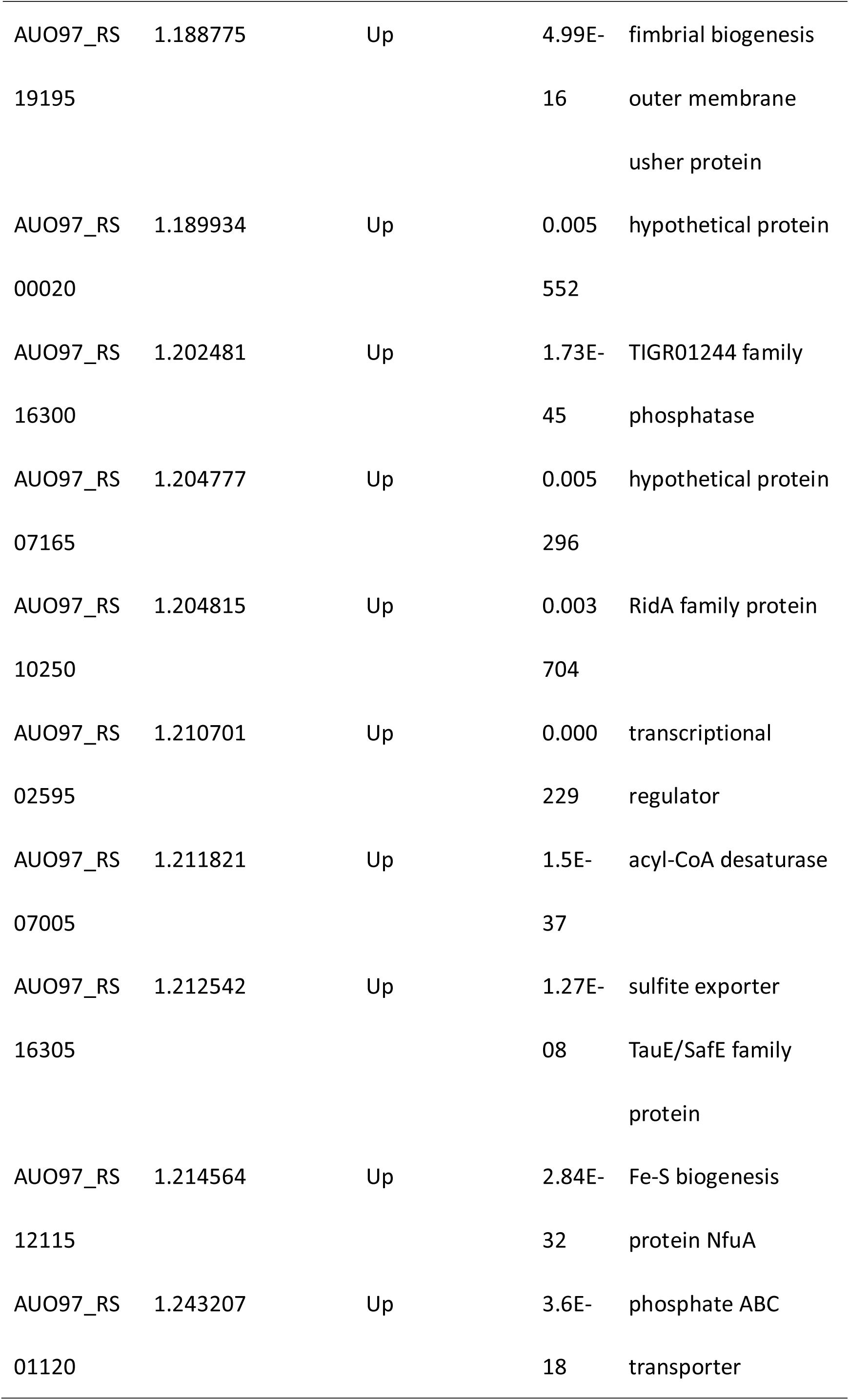
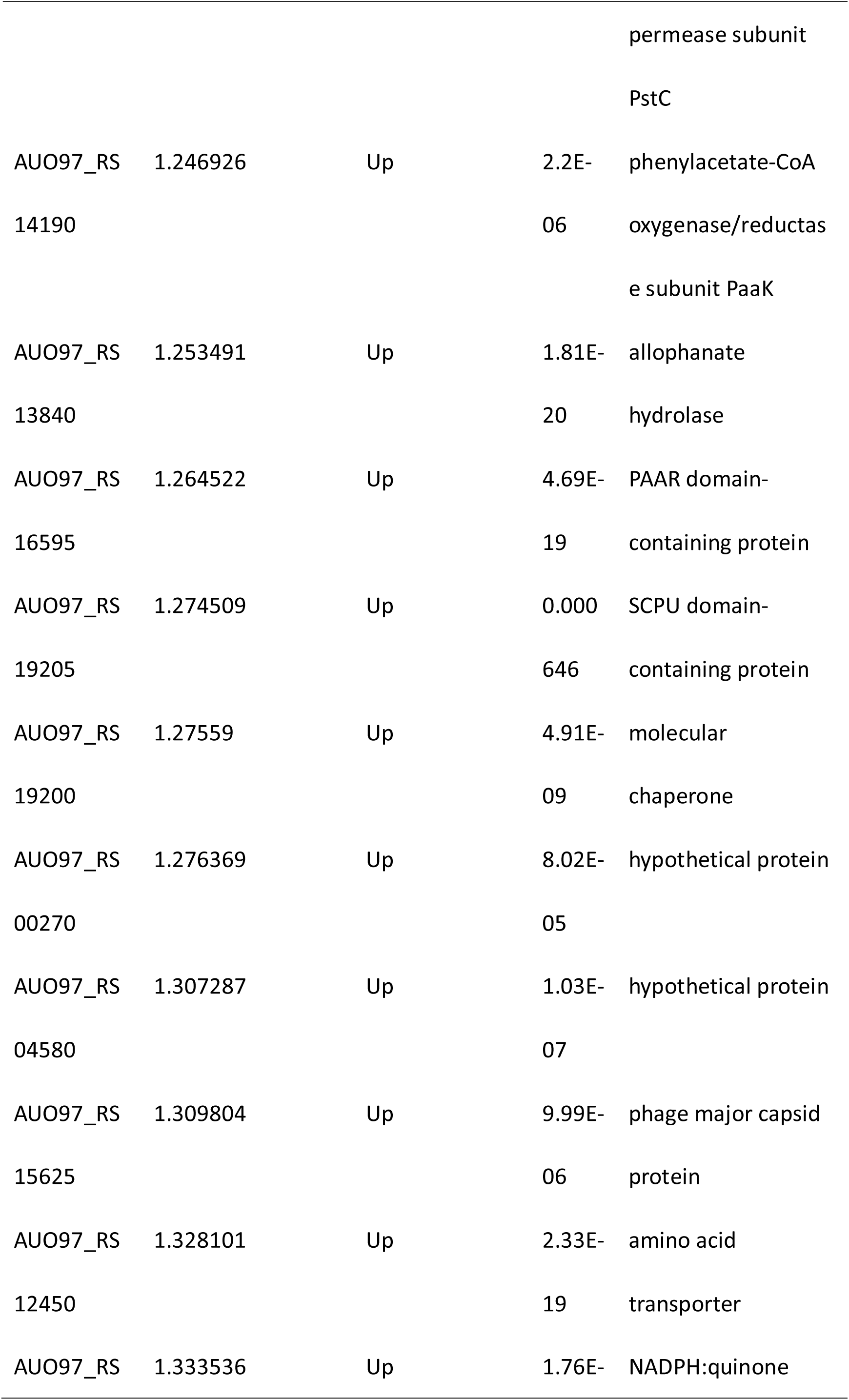
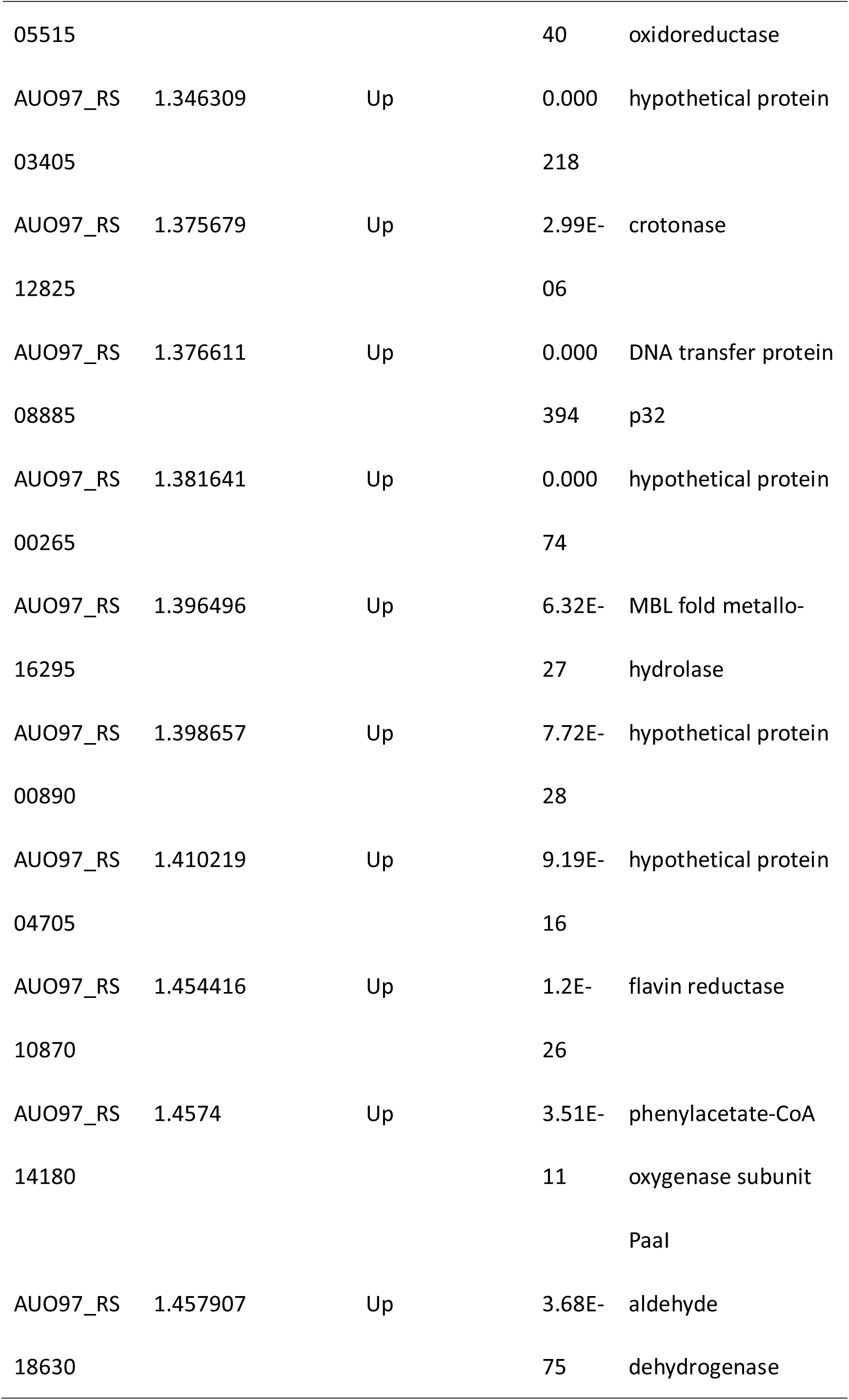
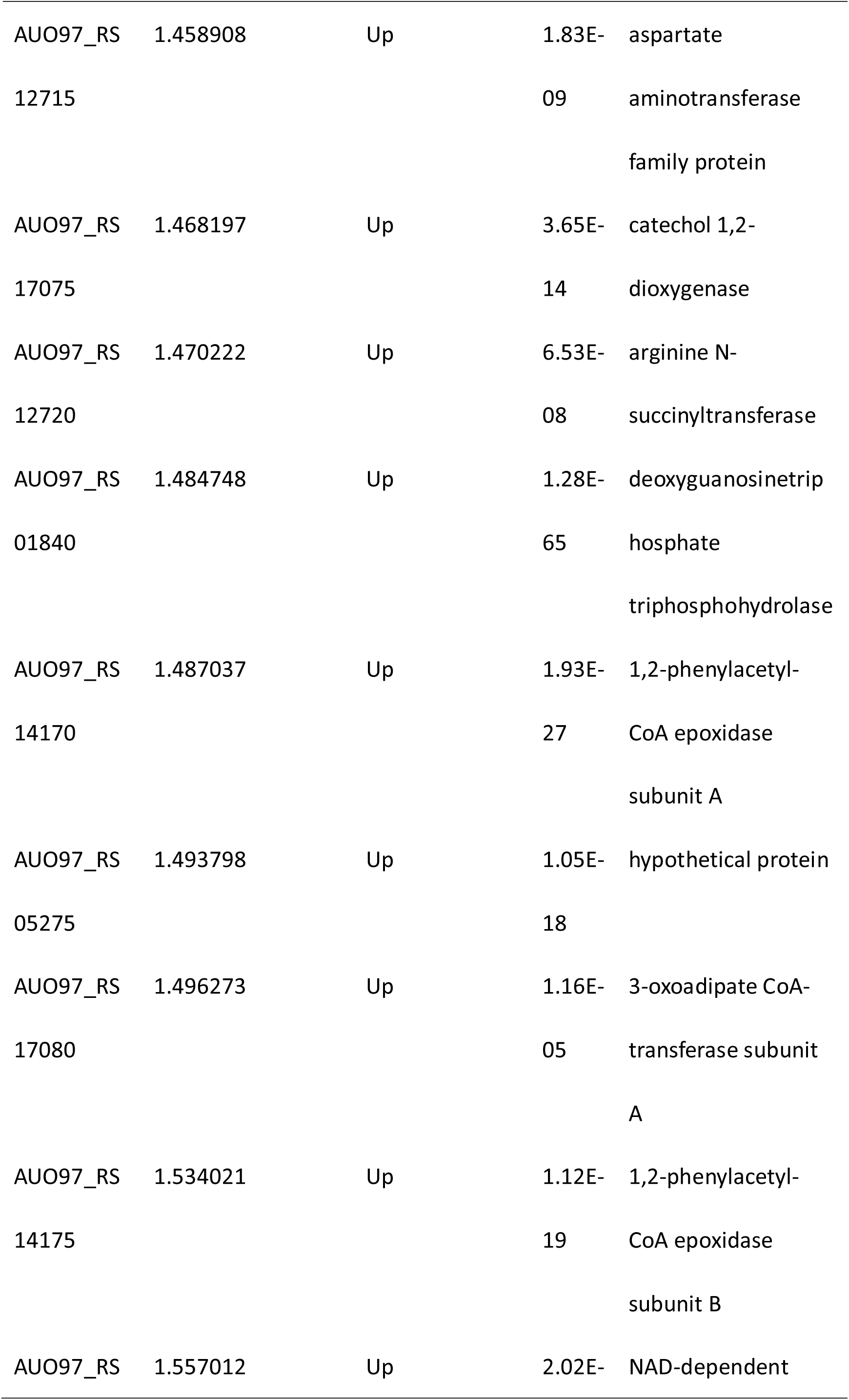
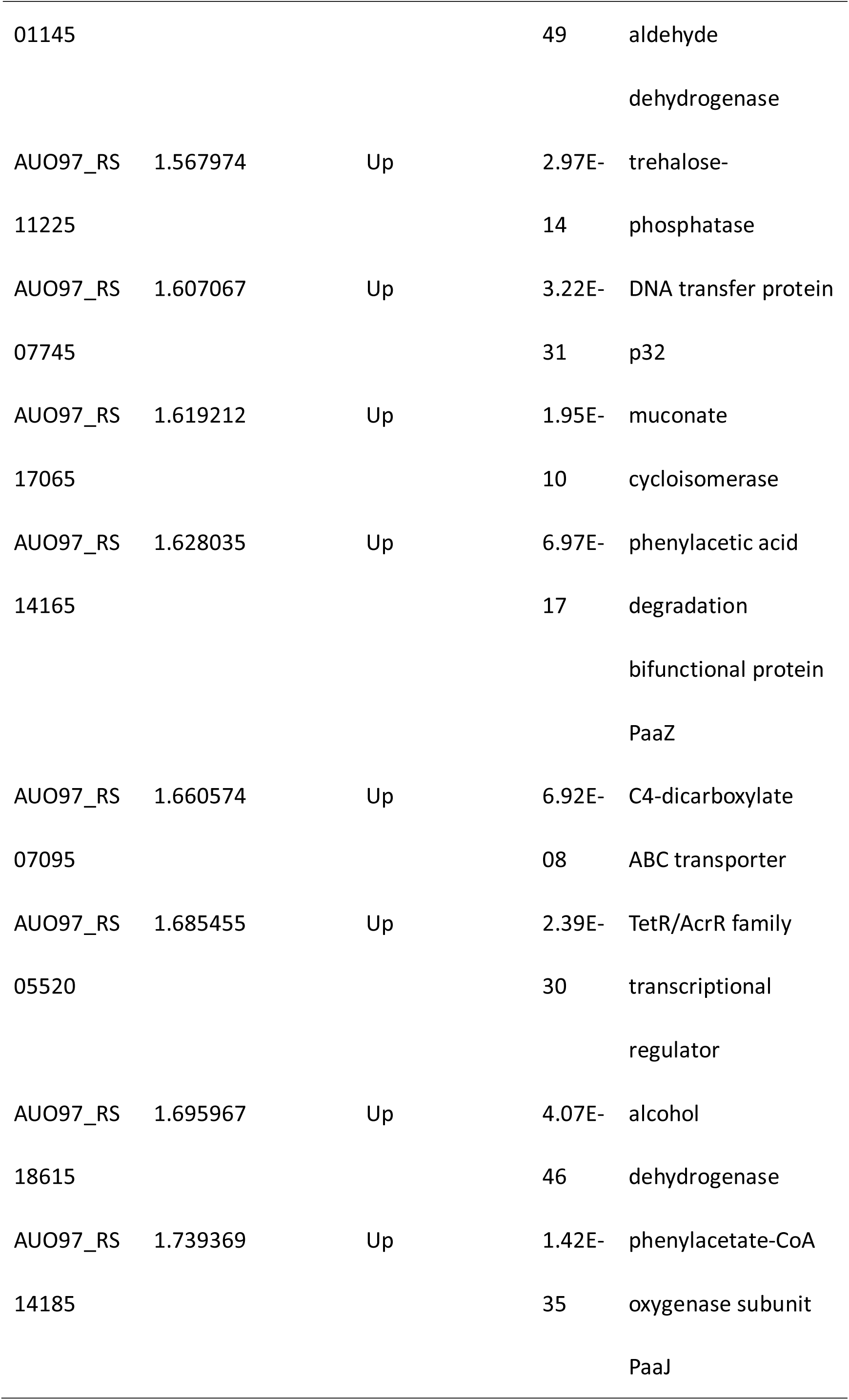
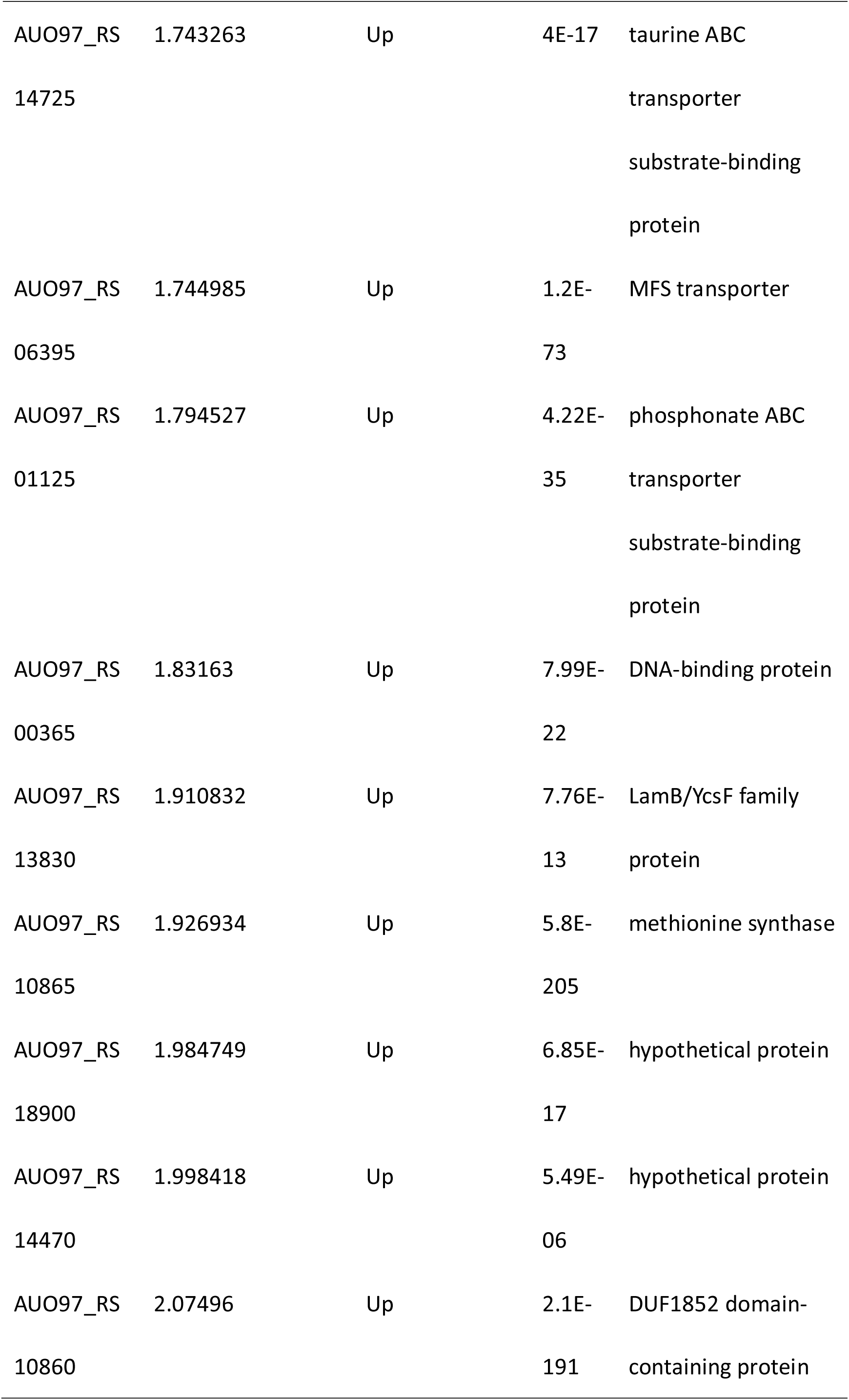
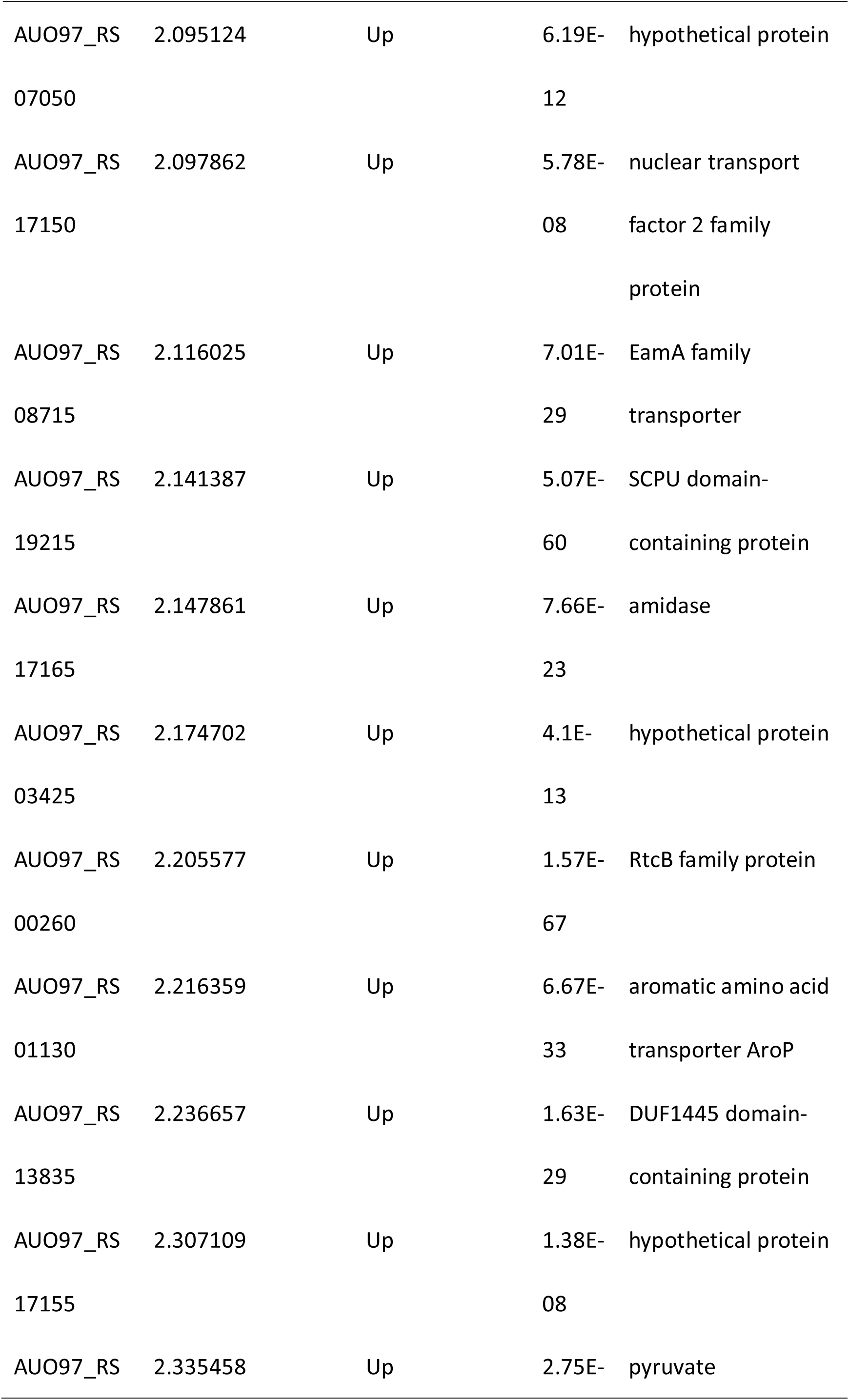
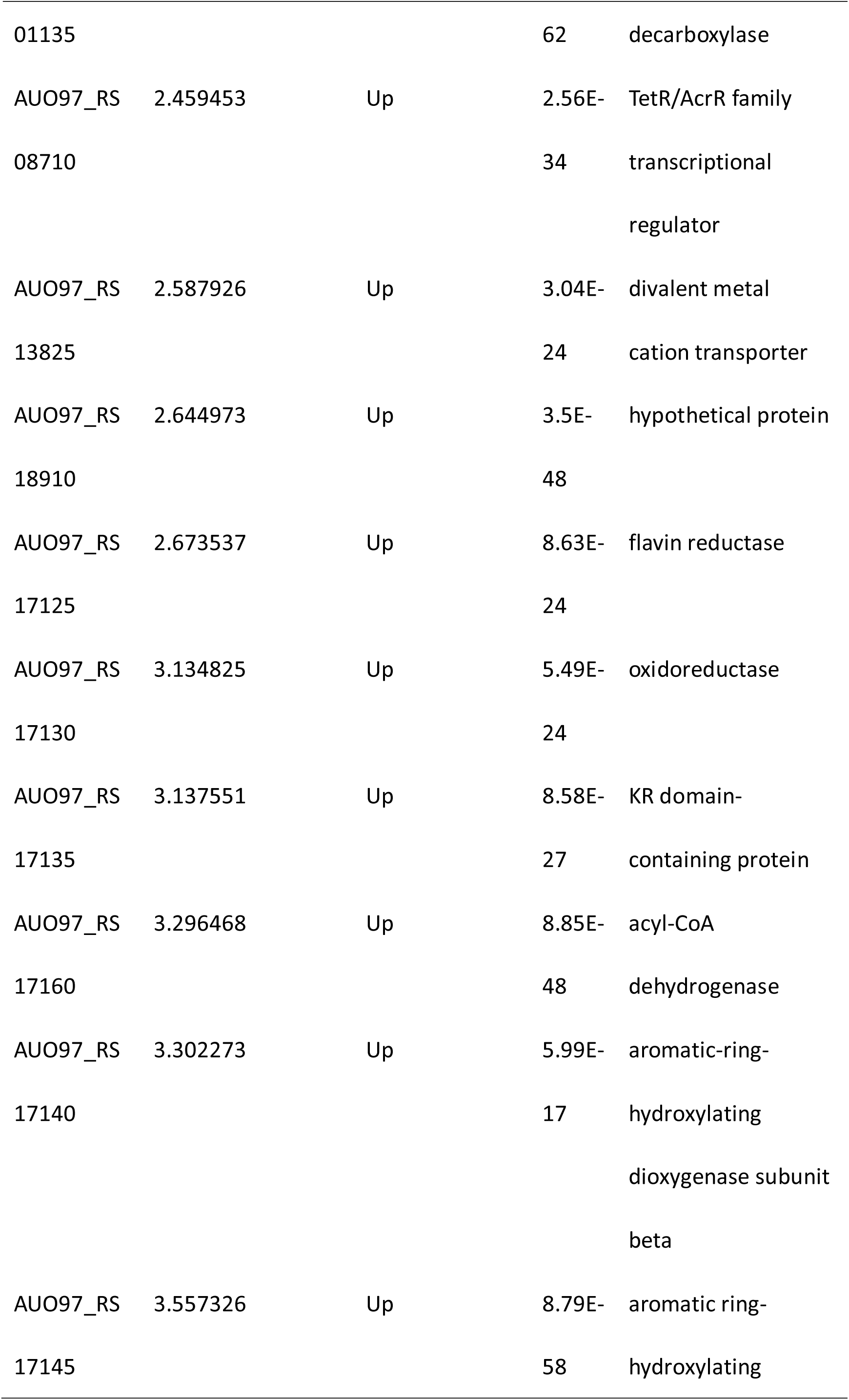
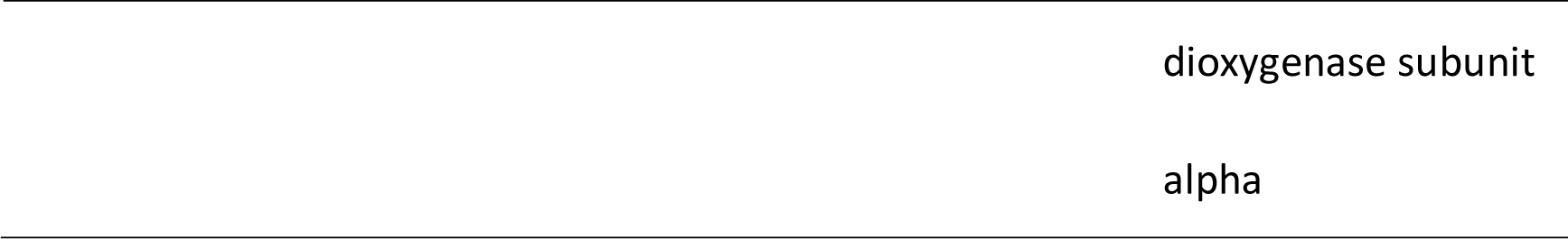
Differential expressed genes in Δ*abaI* strain

**Table 6.**
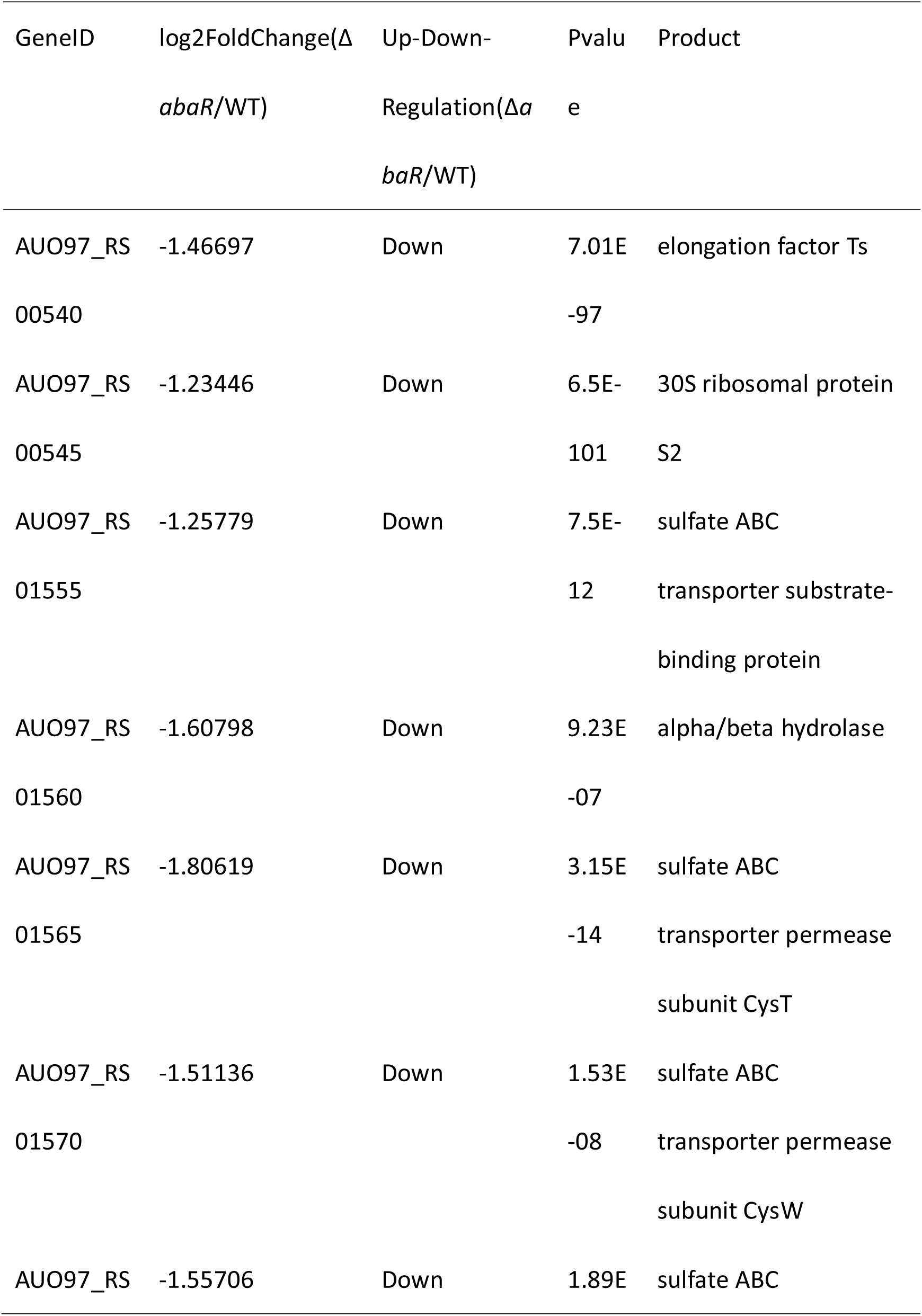
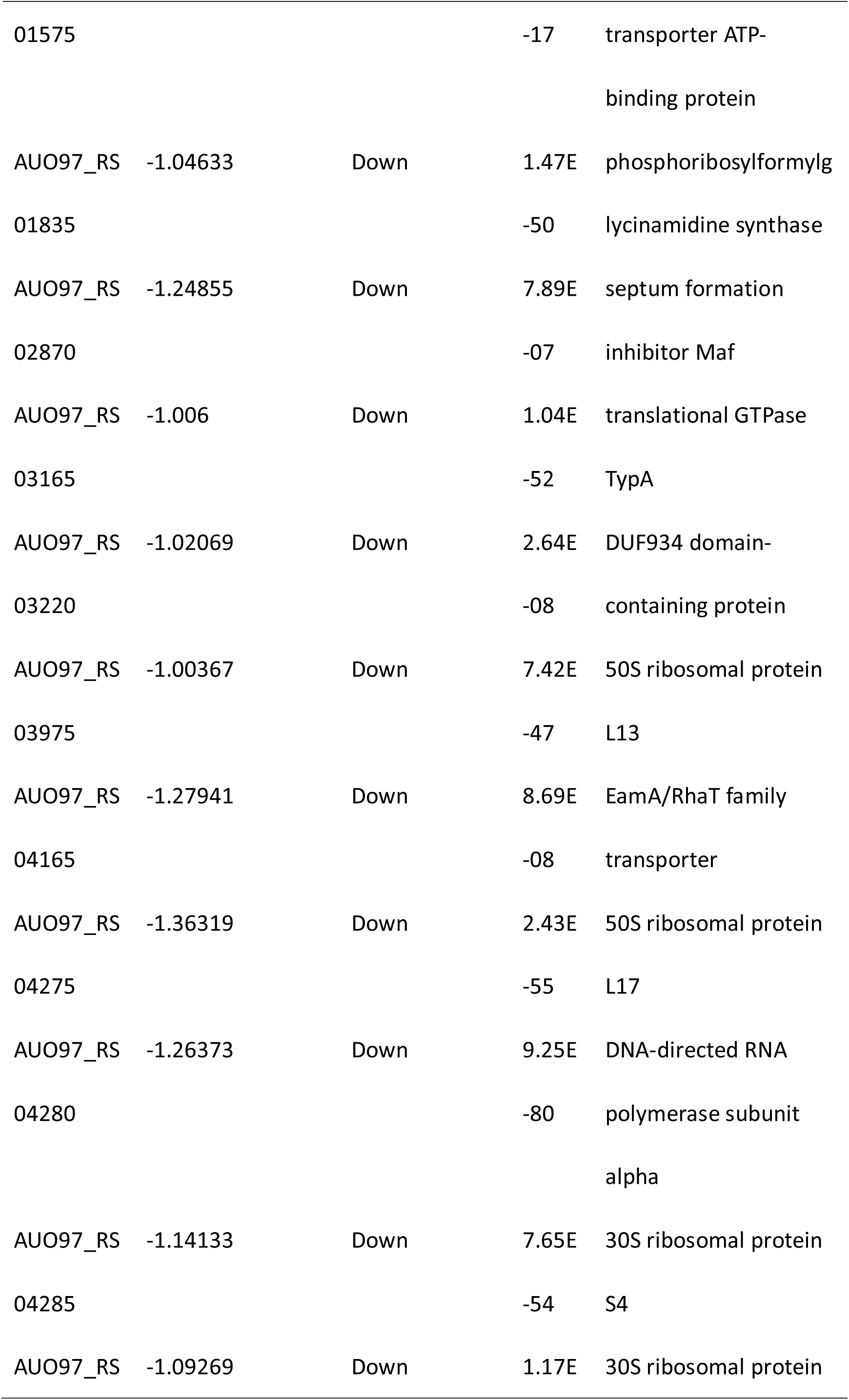
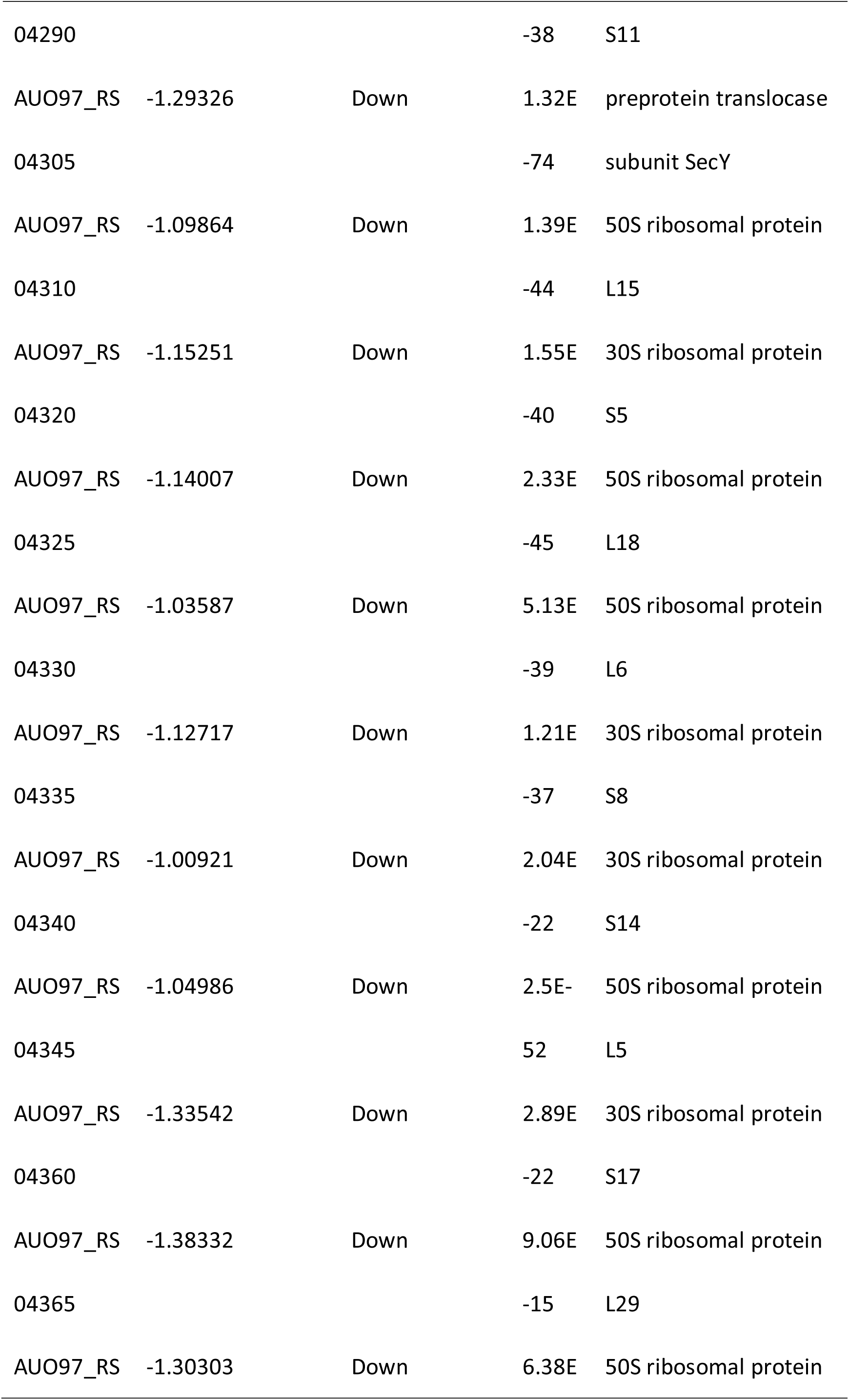
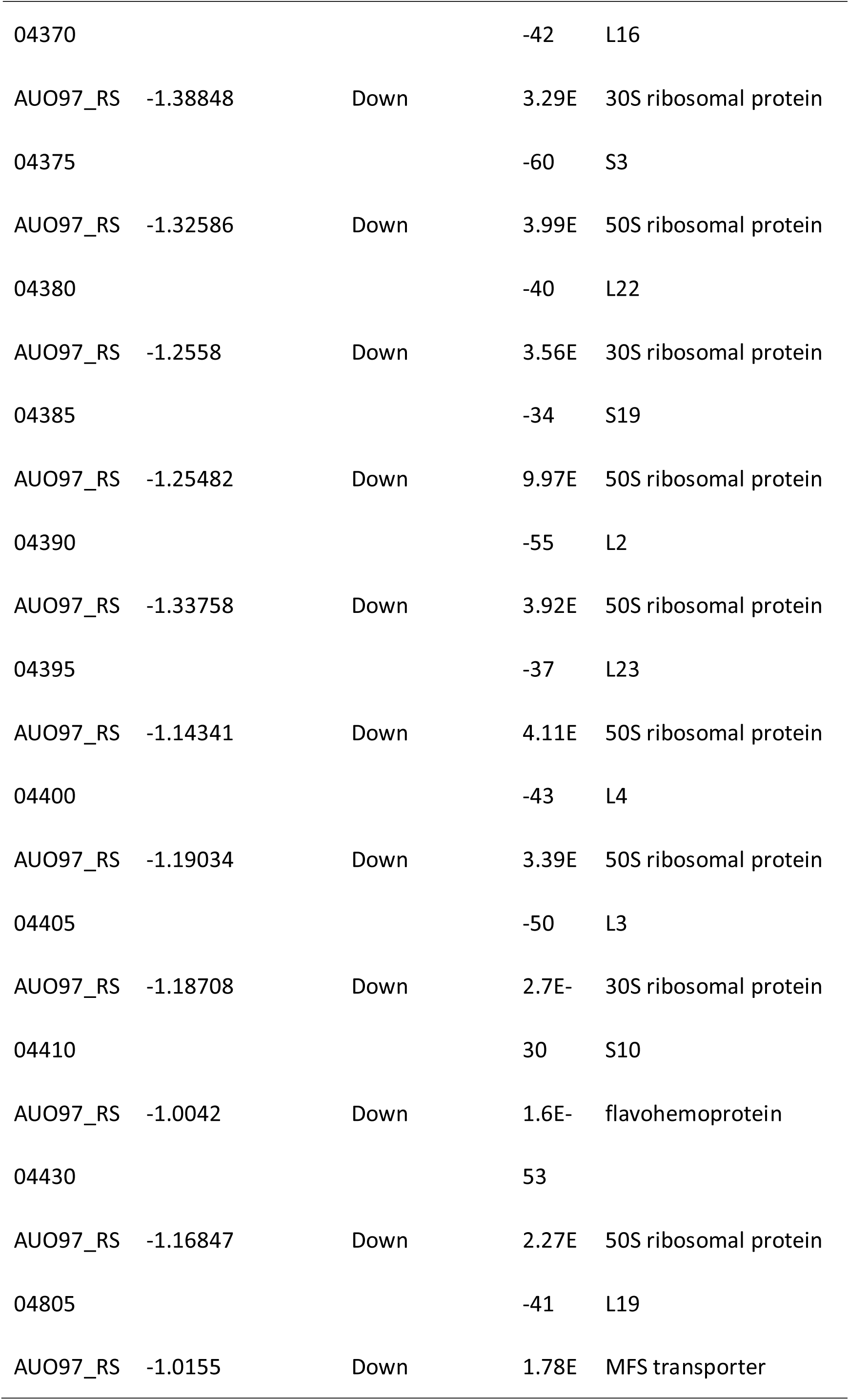
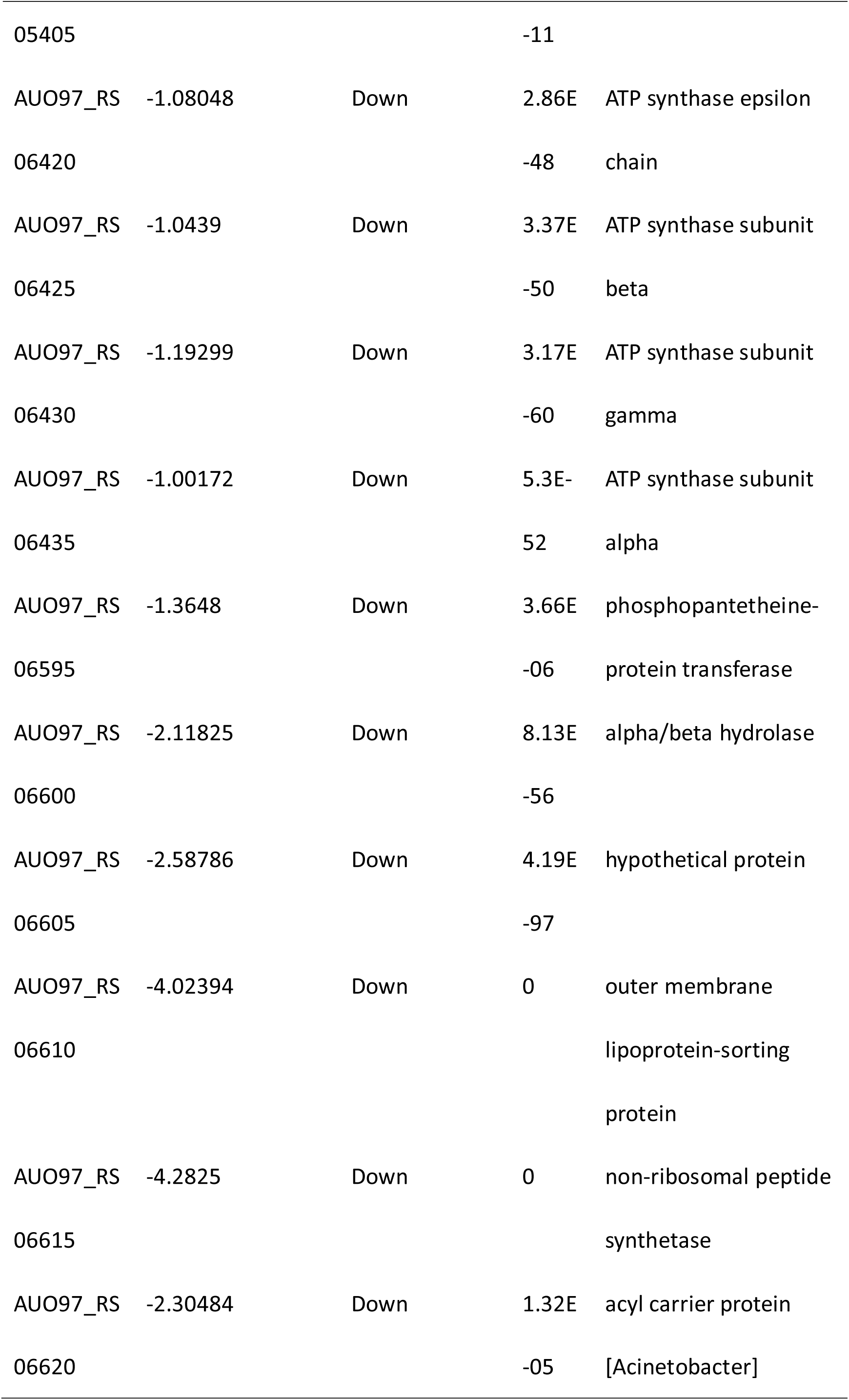
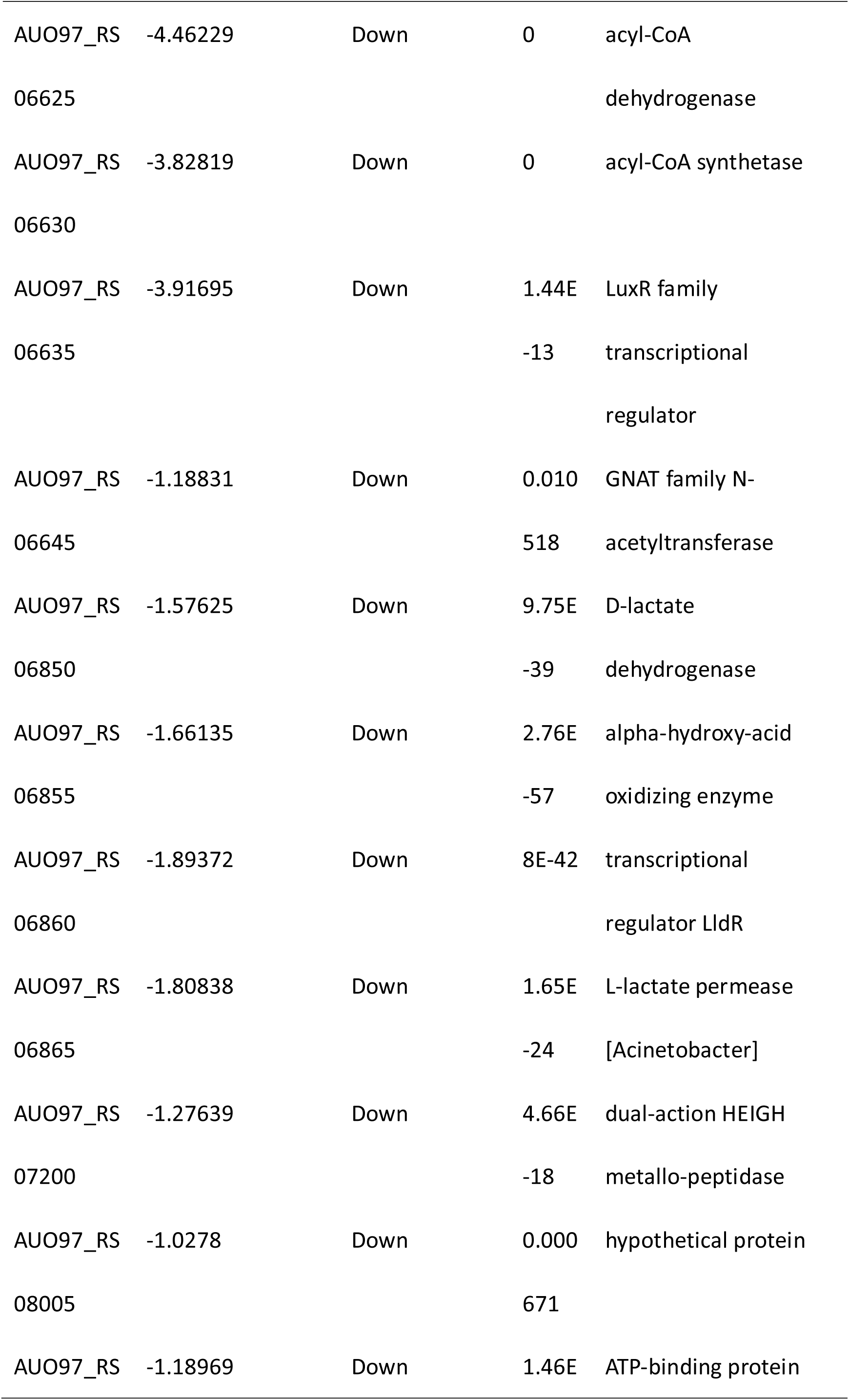
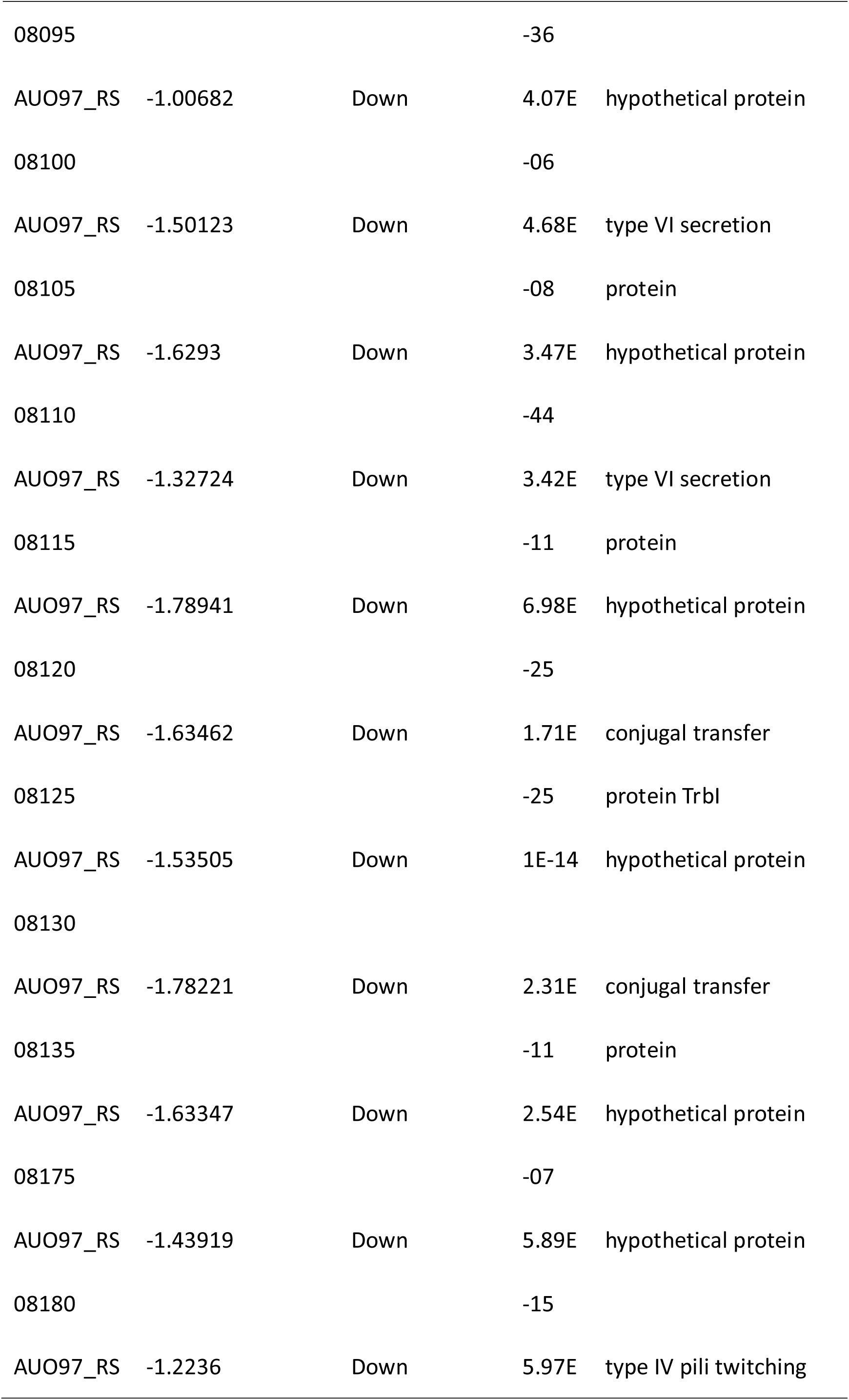
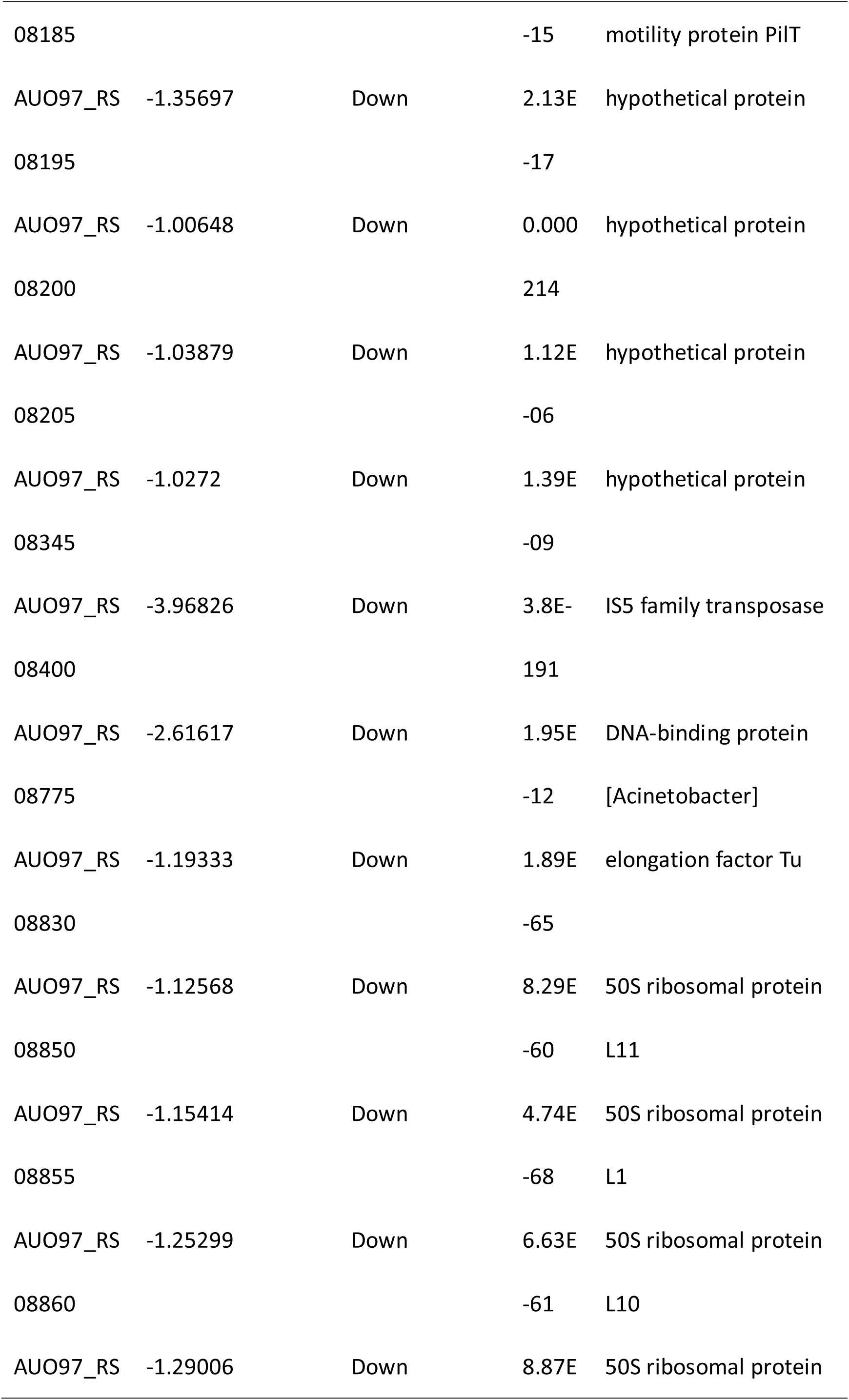
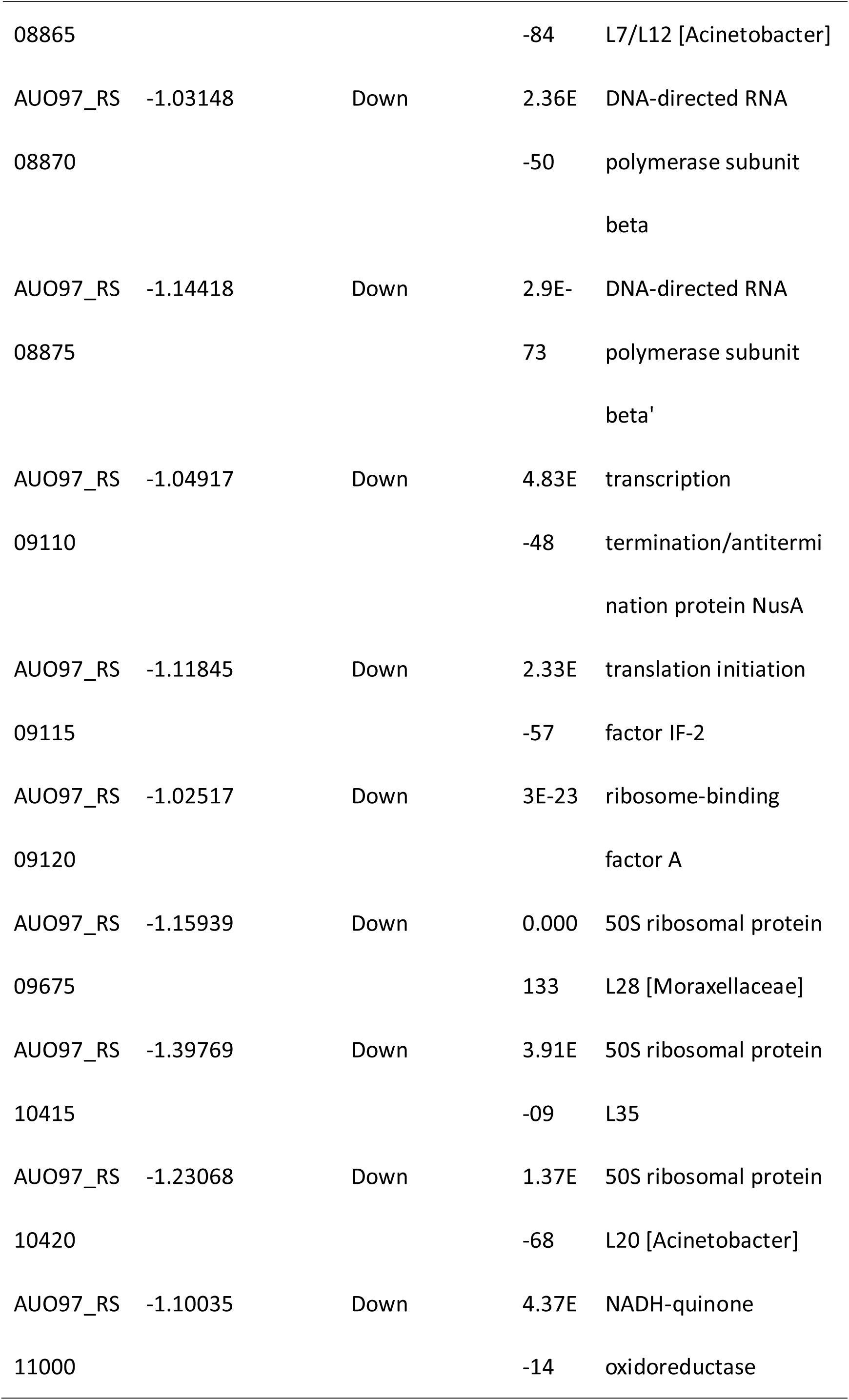
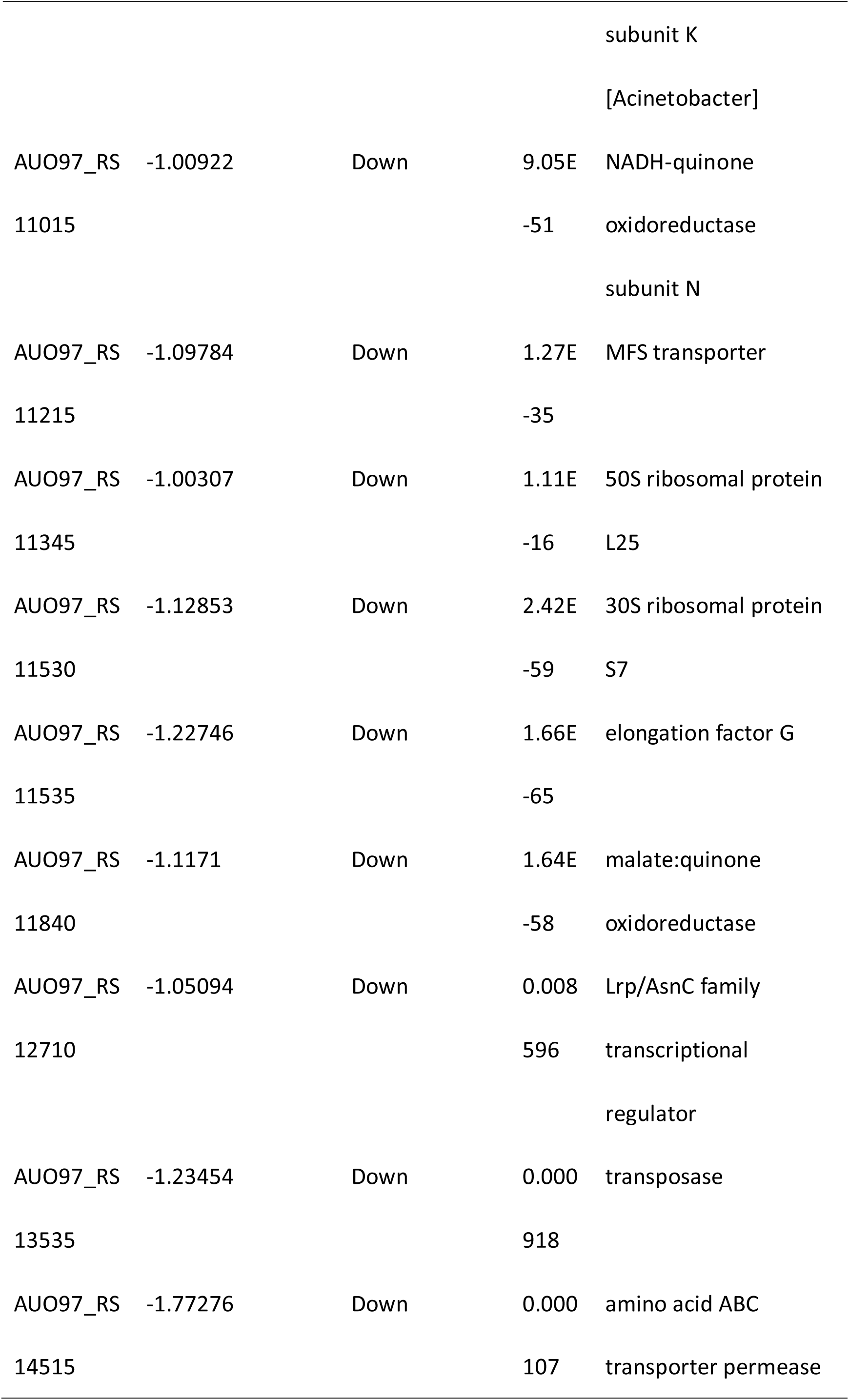
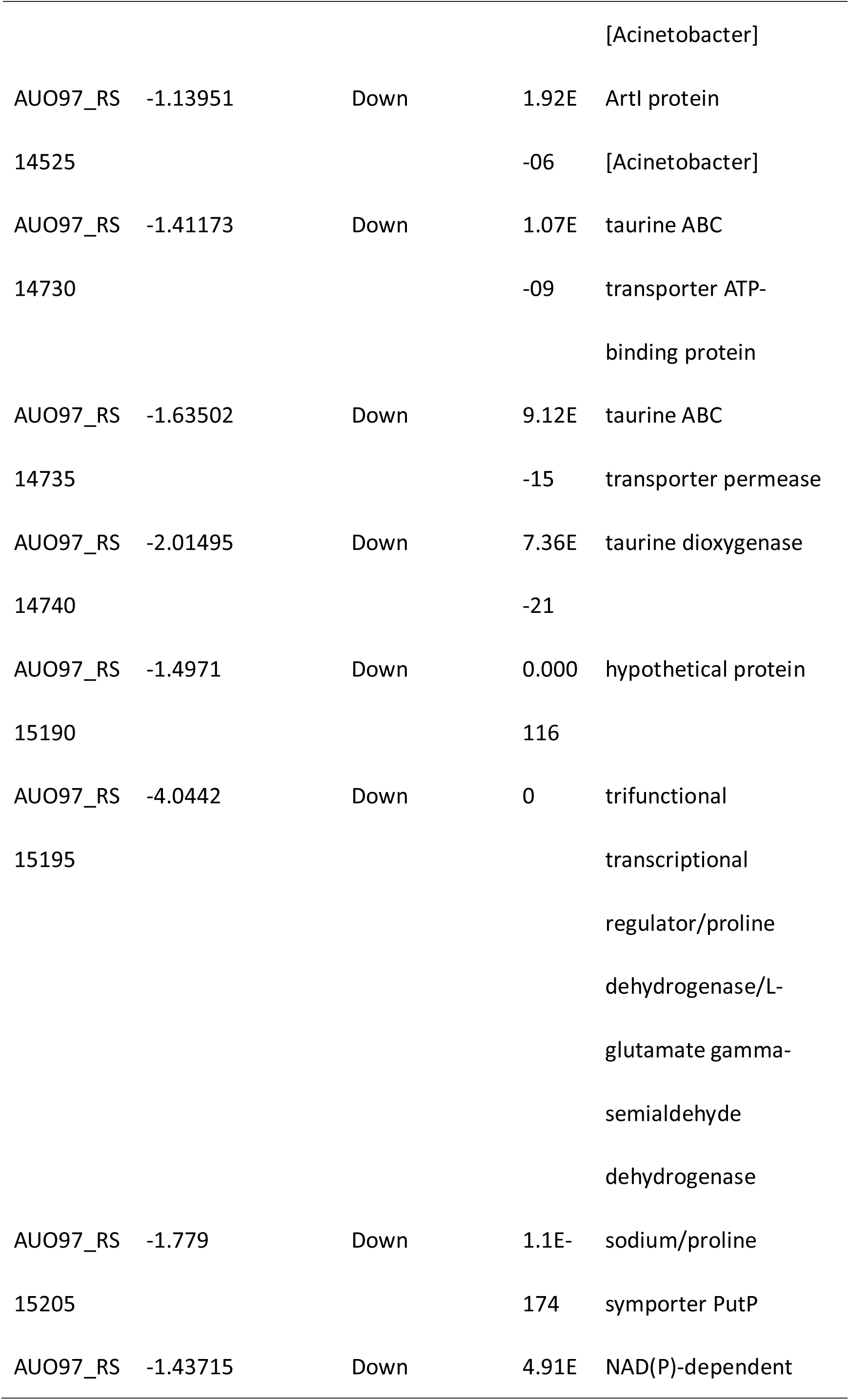
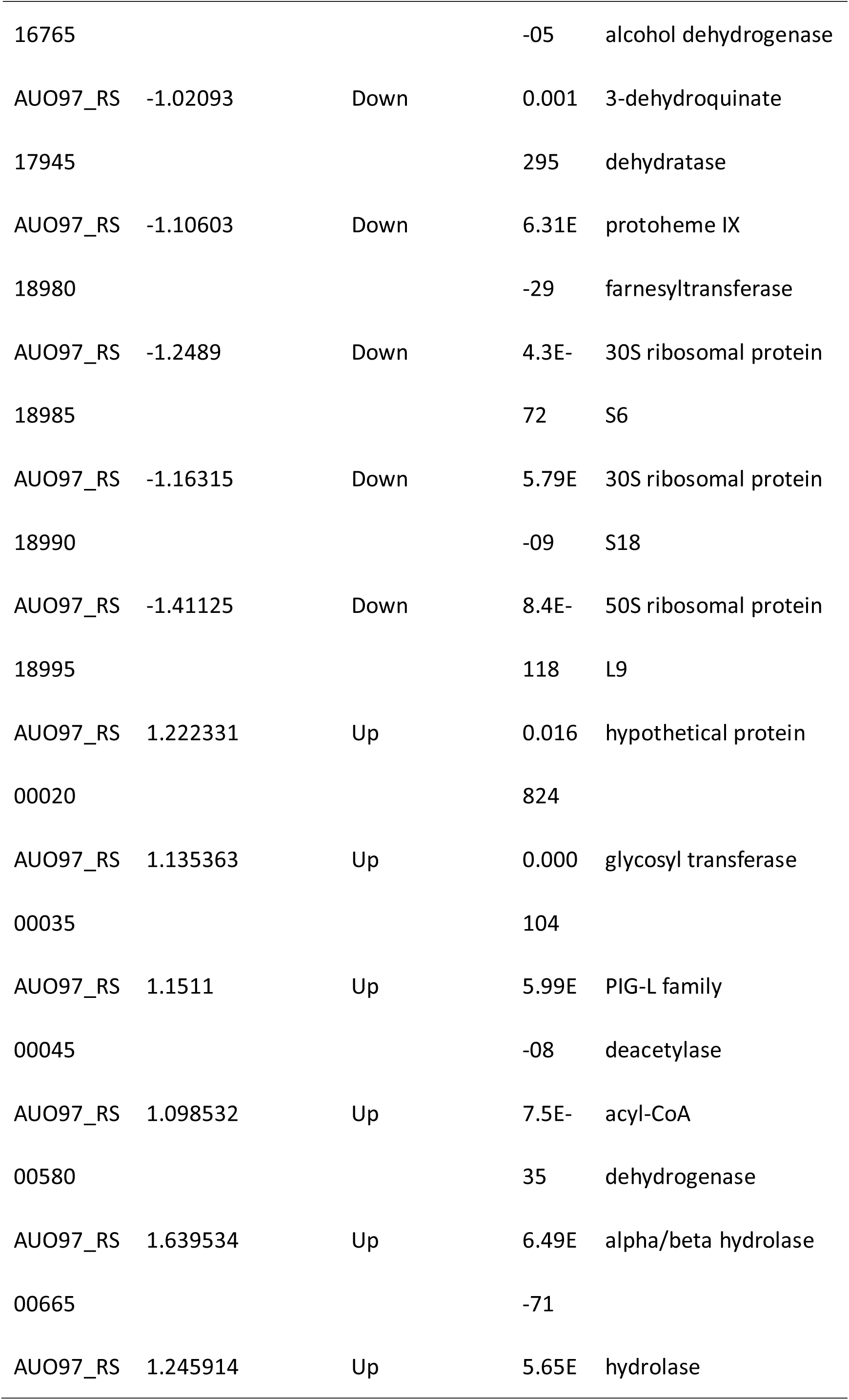
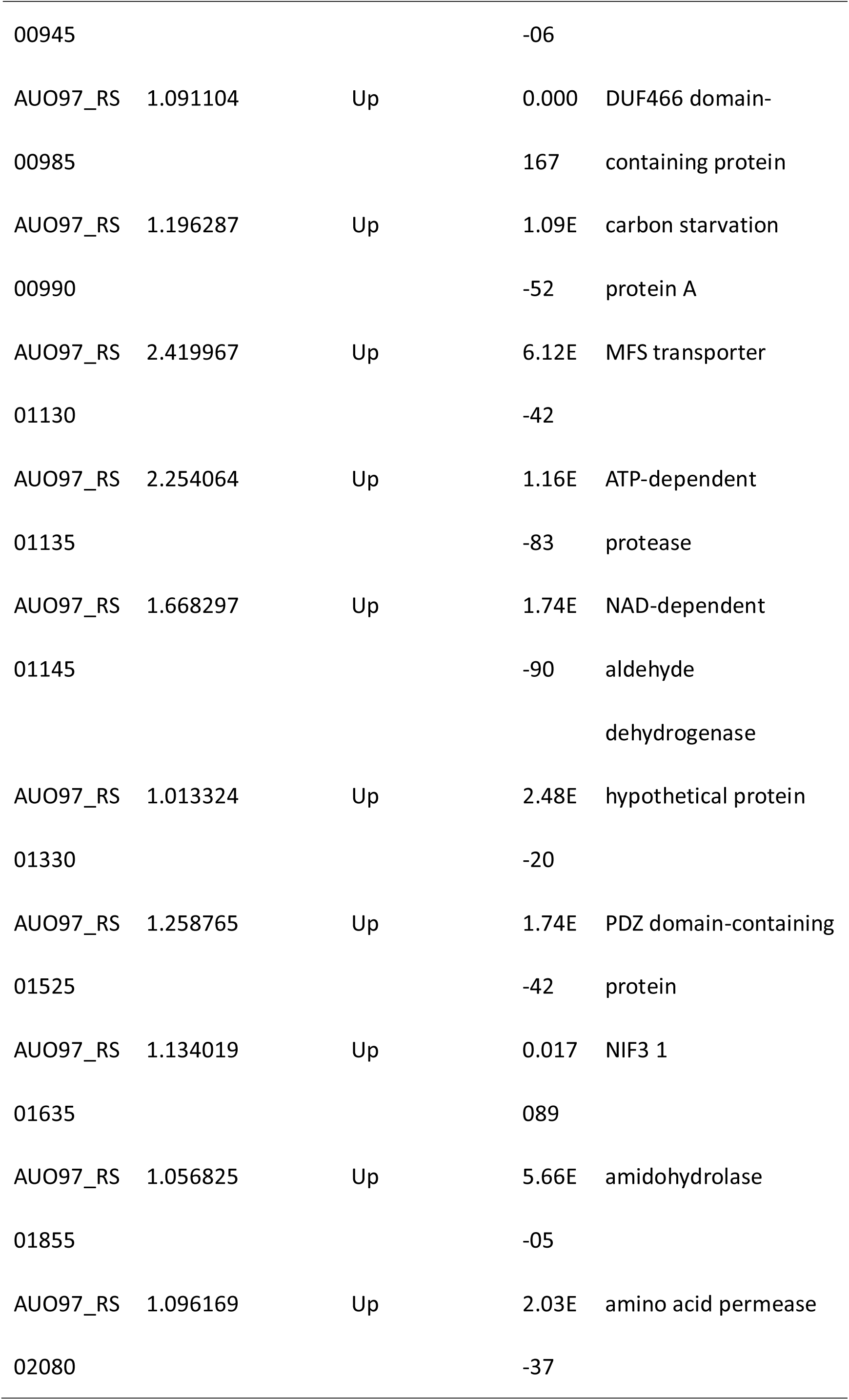
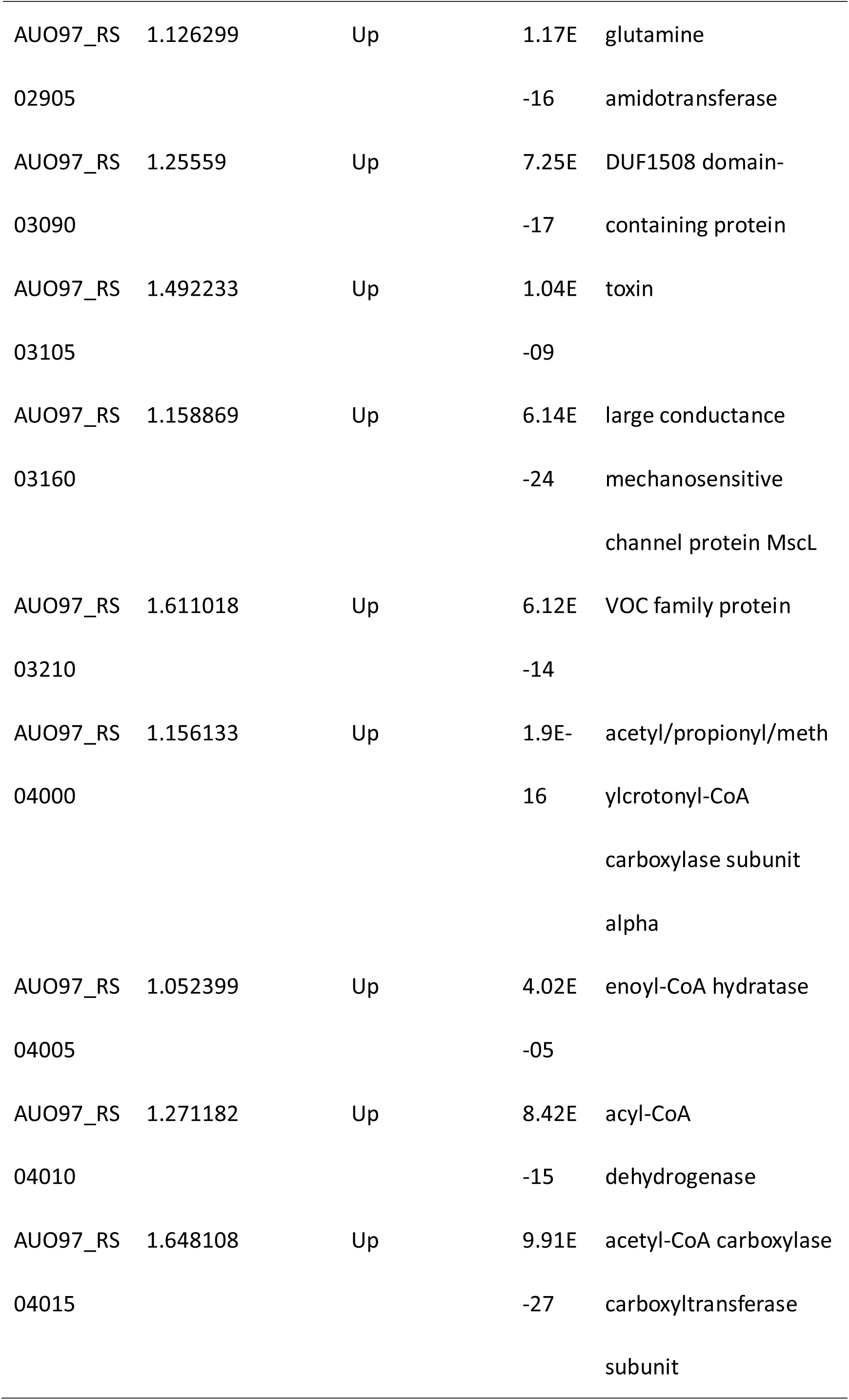
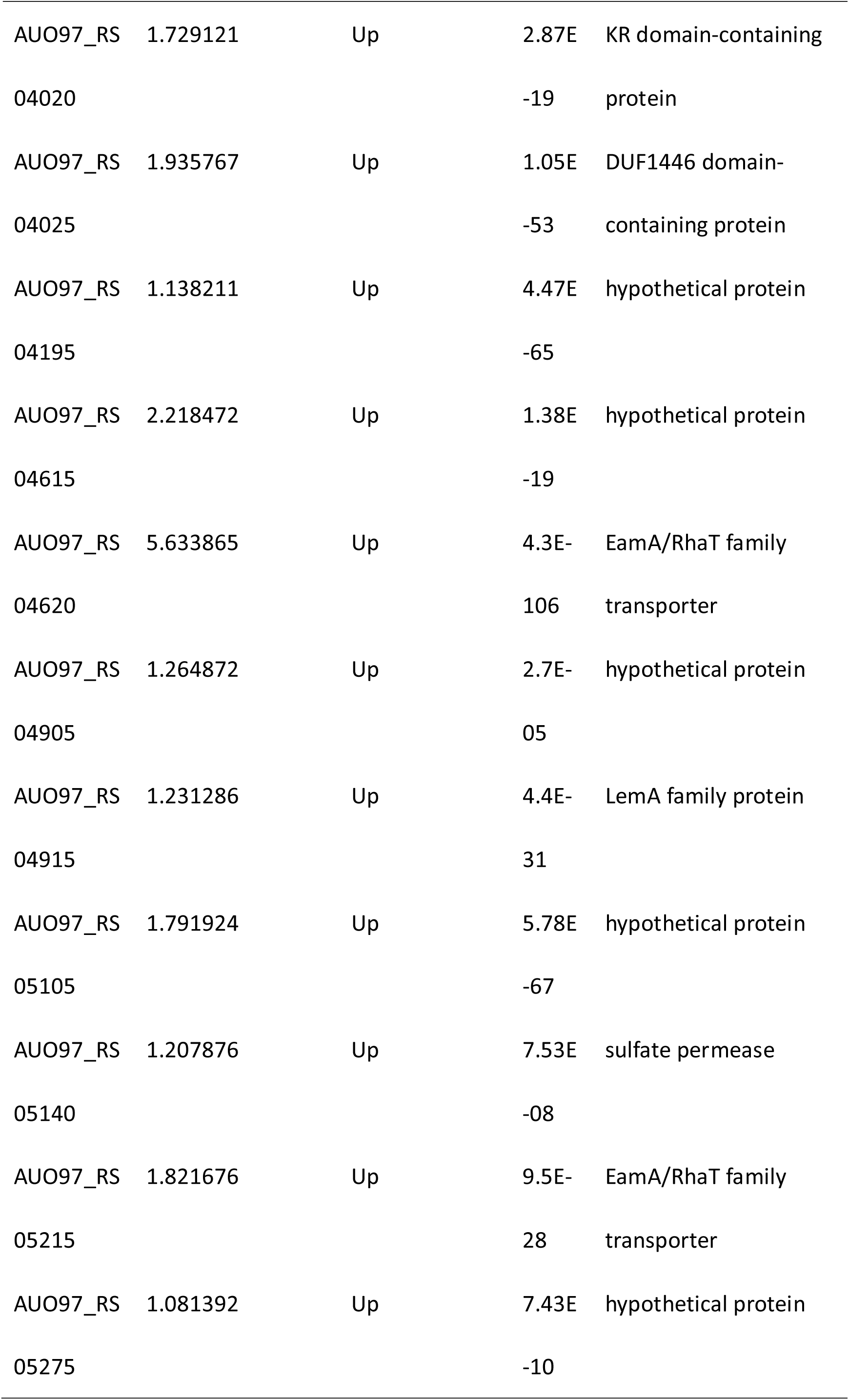
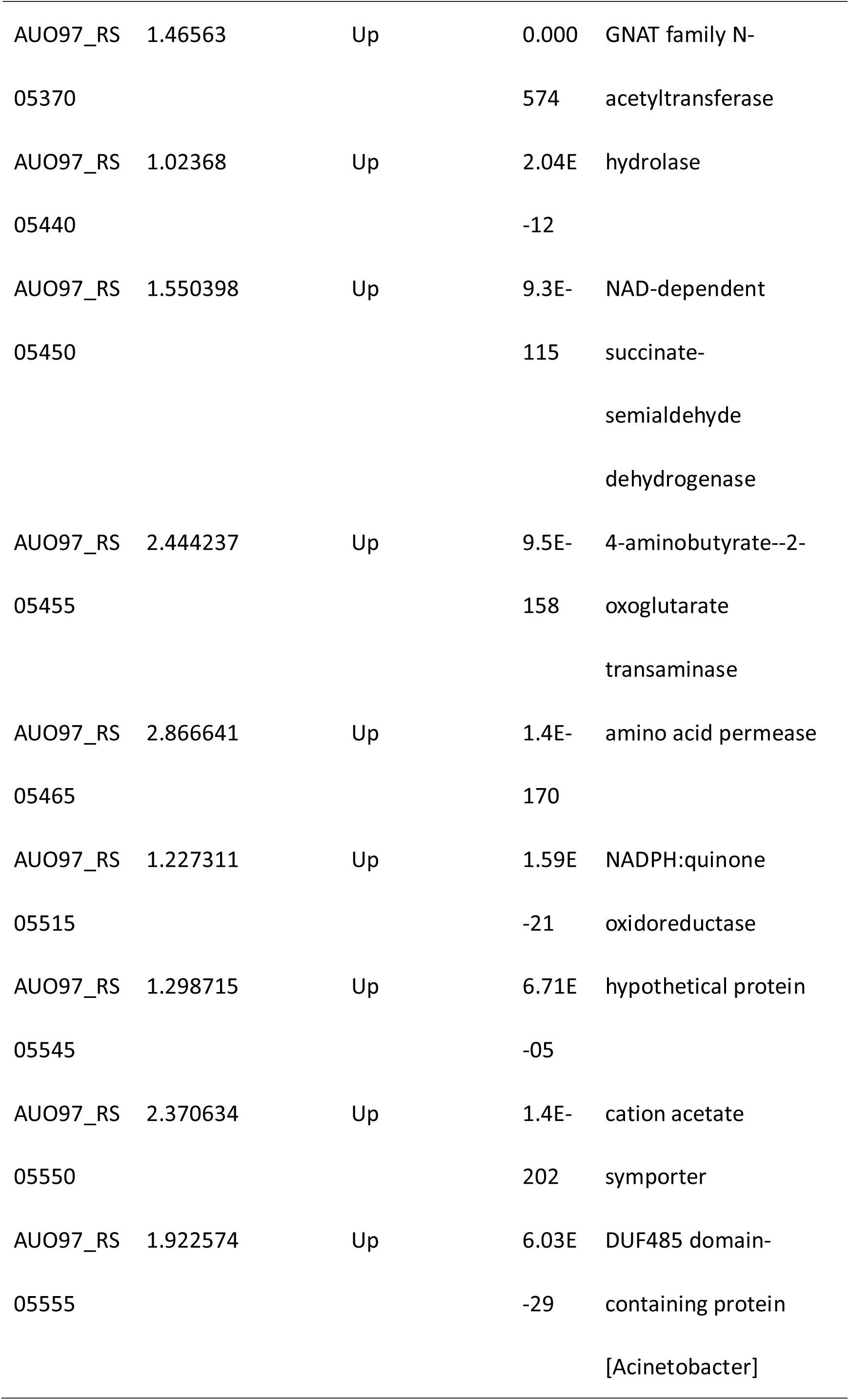
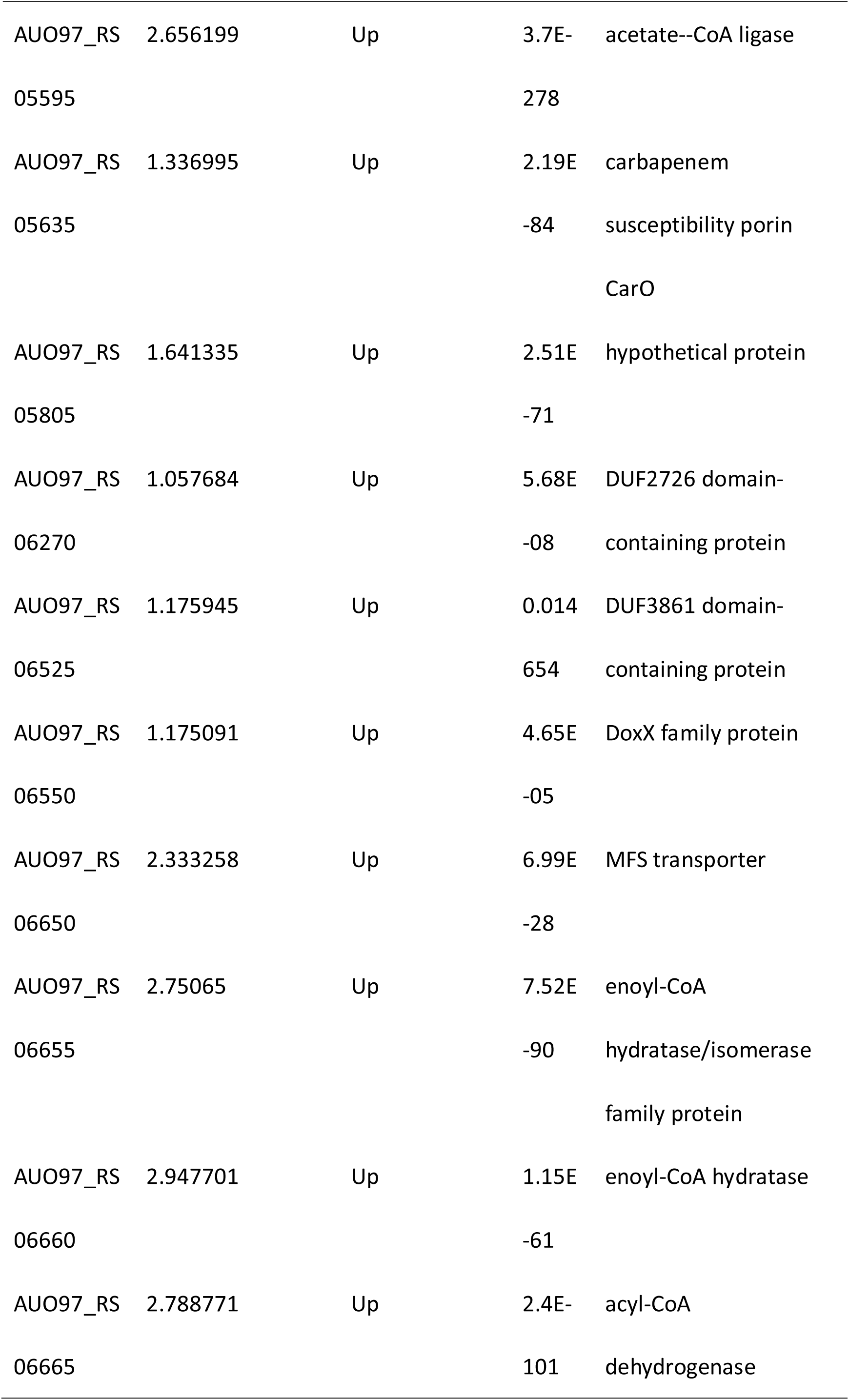
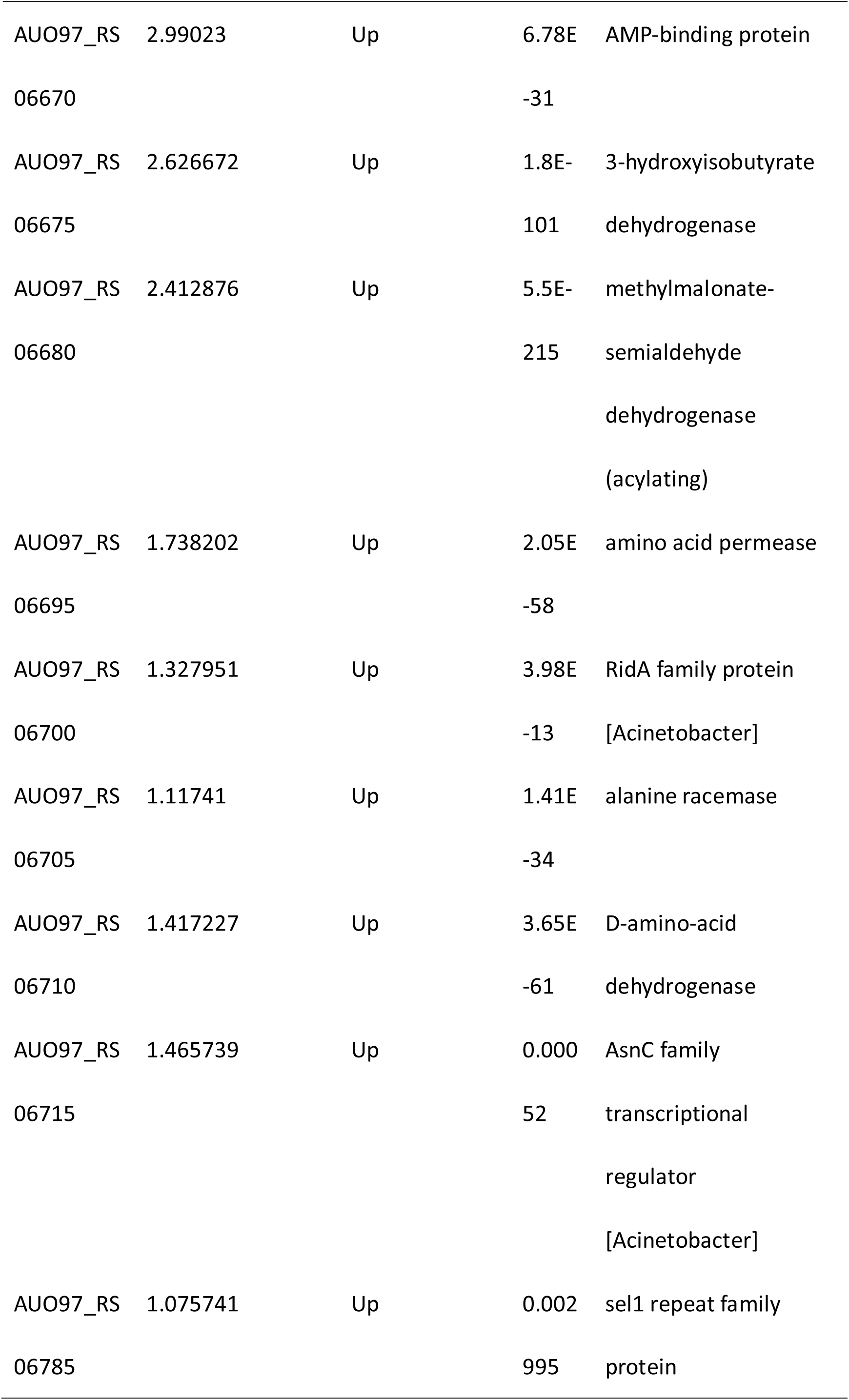
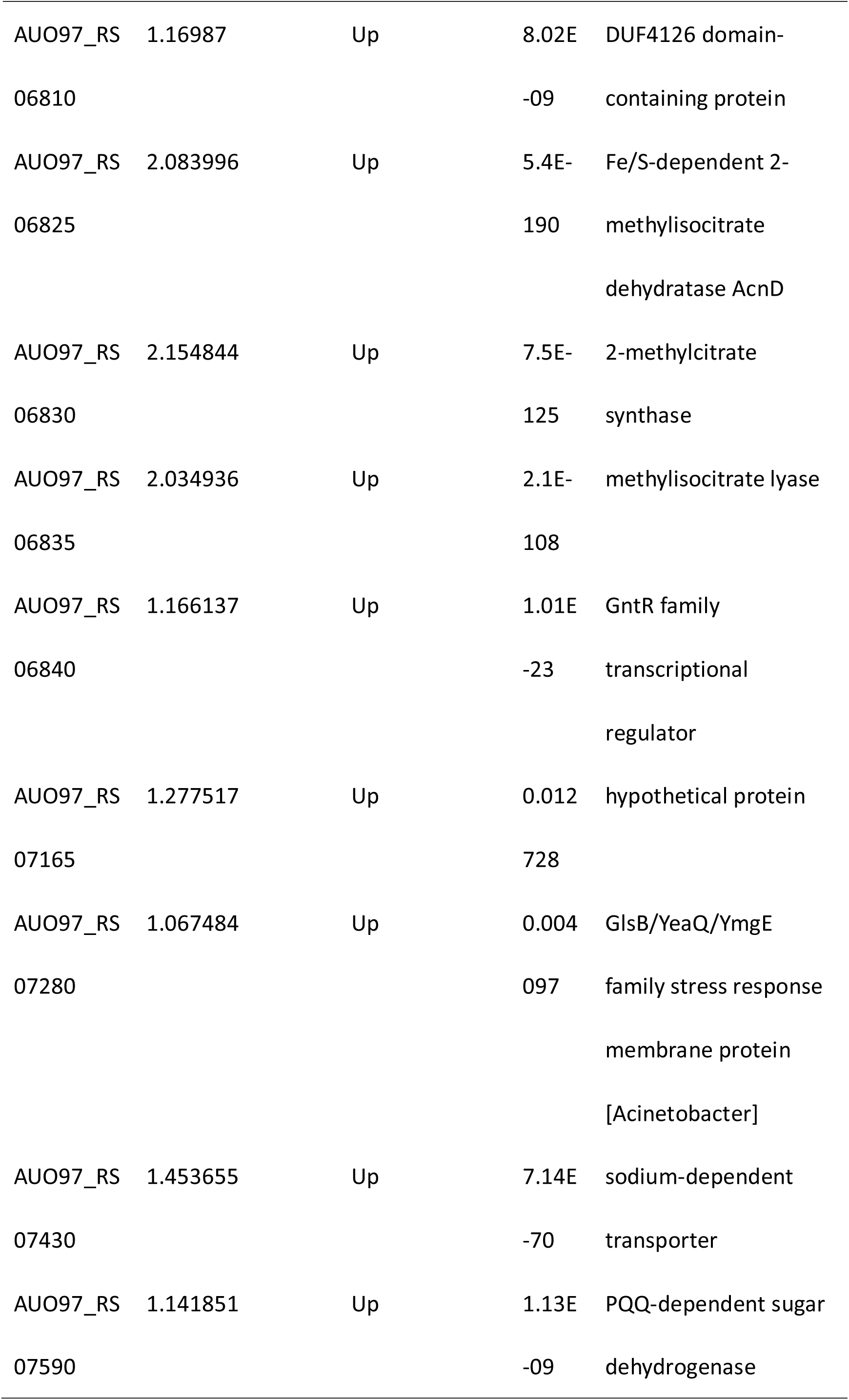
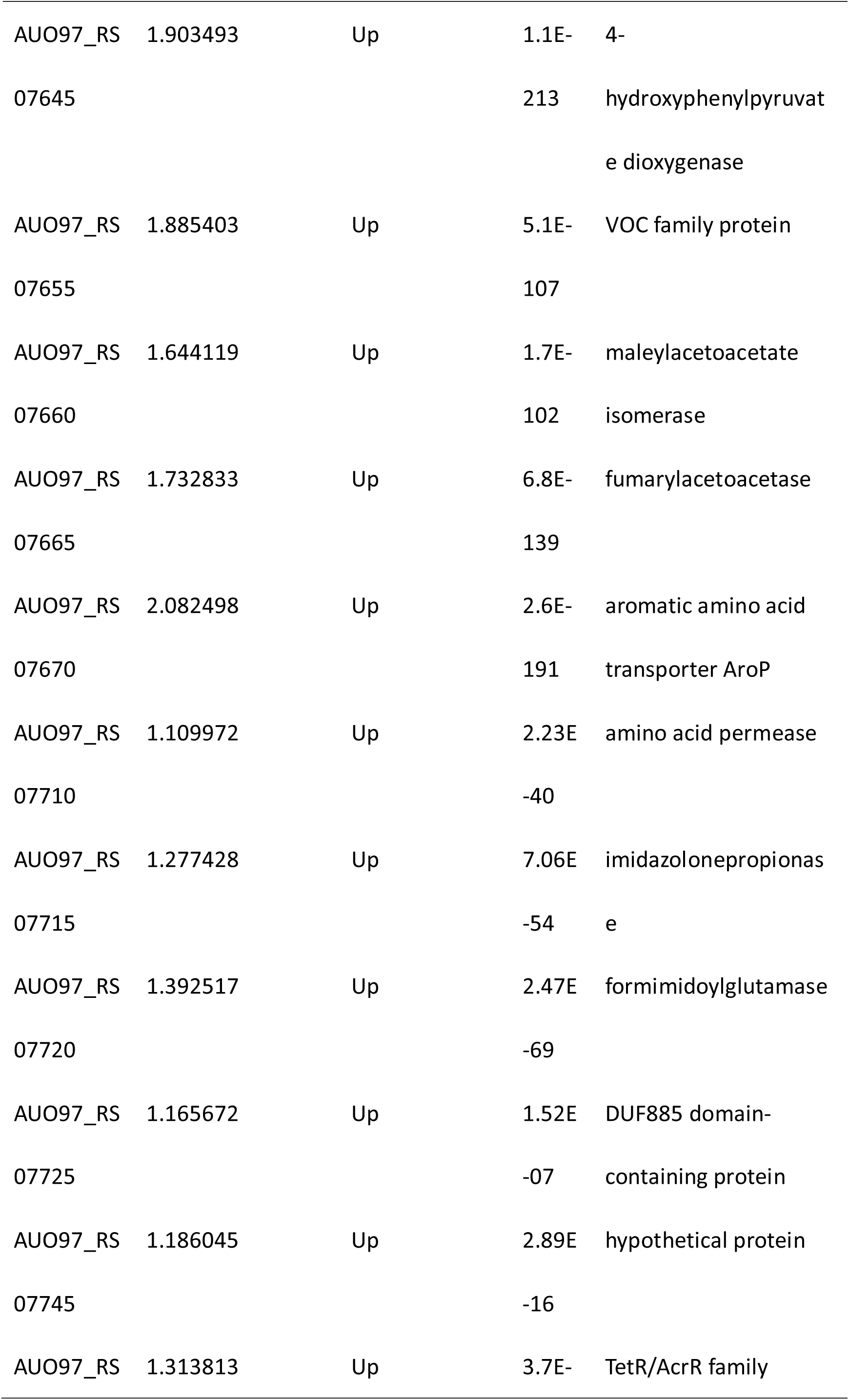
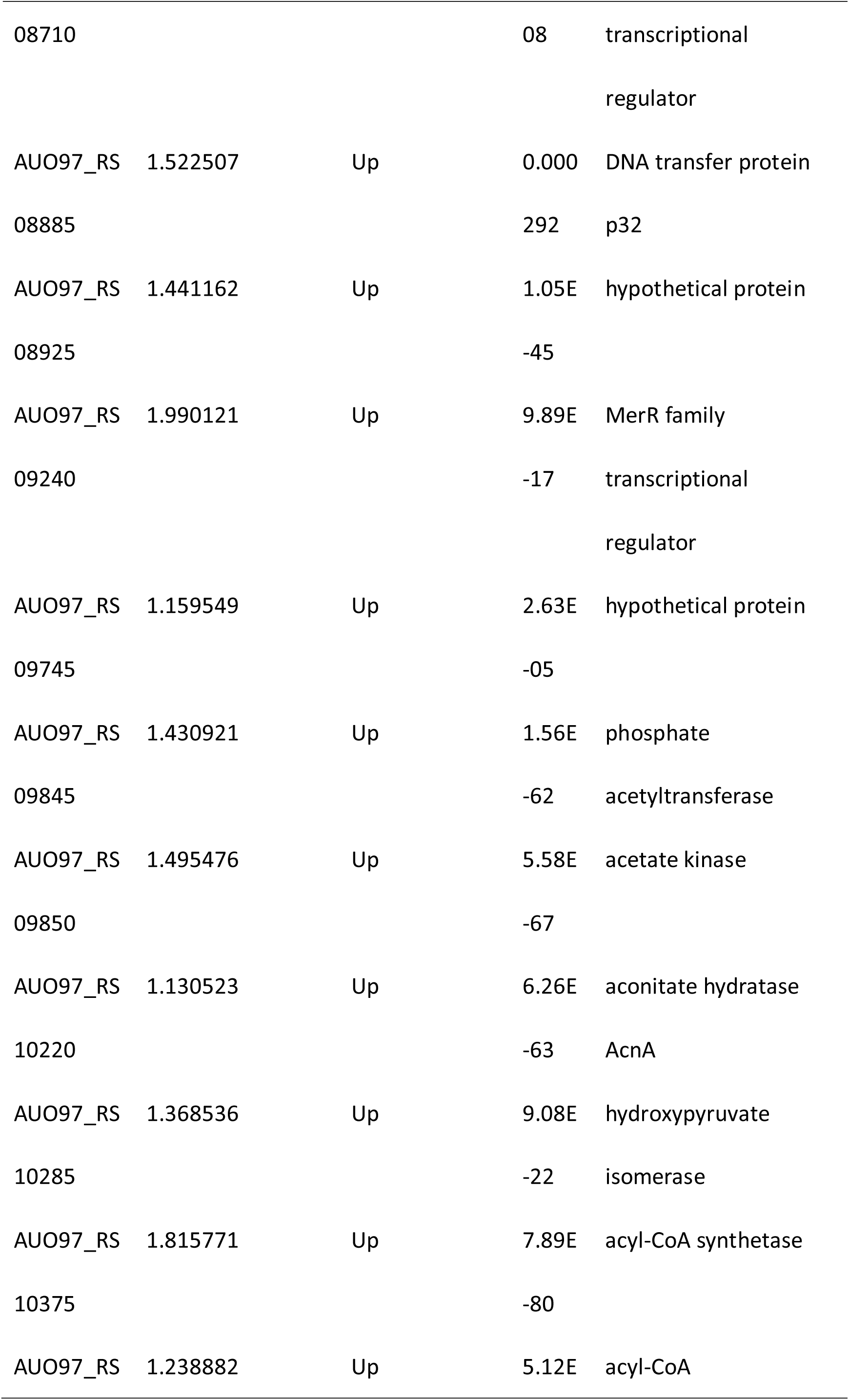
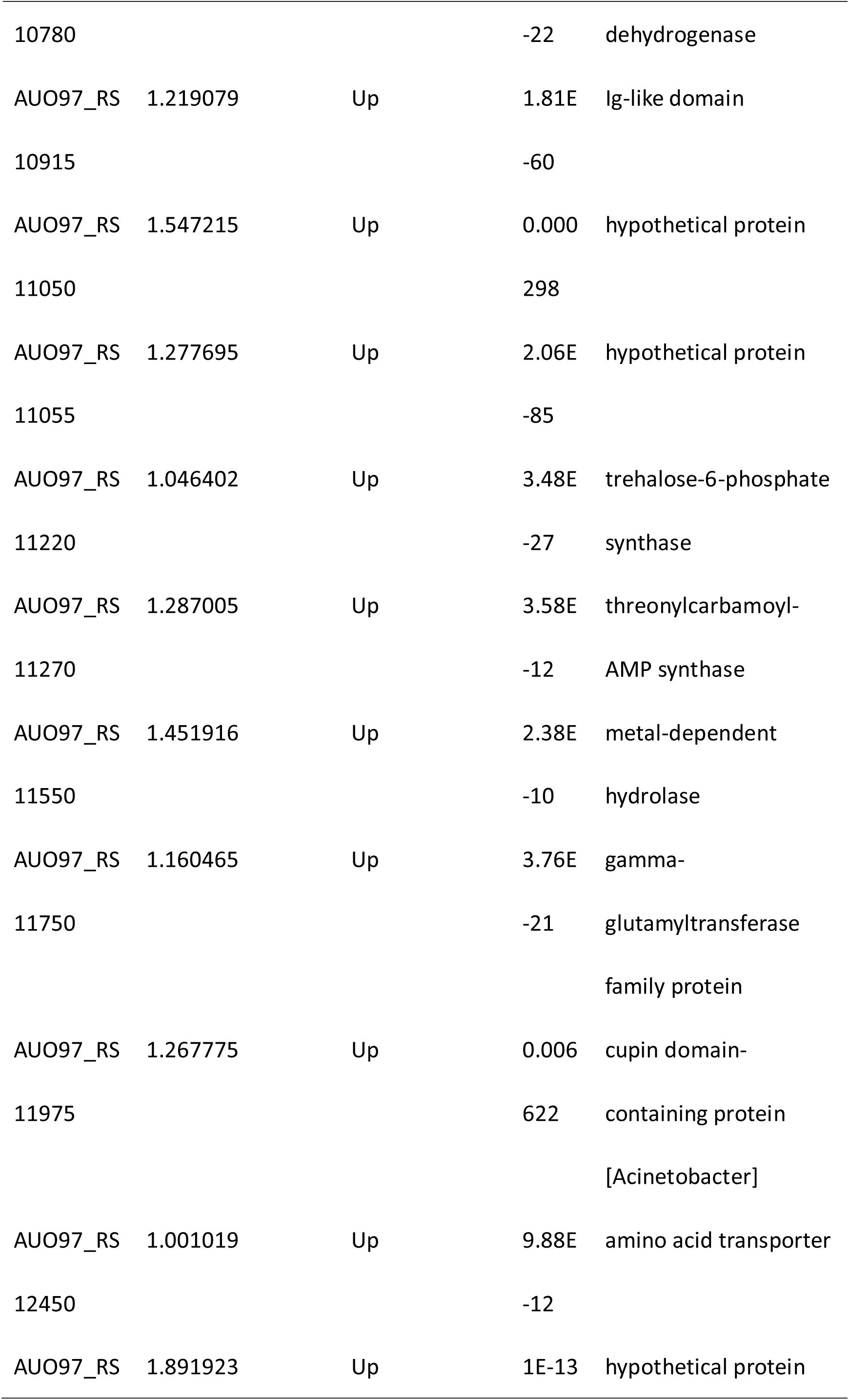
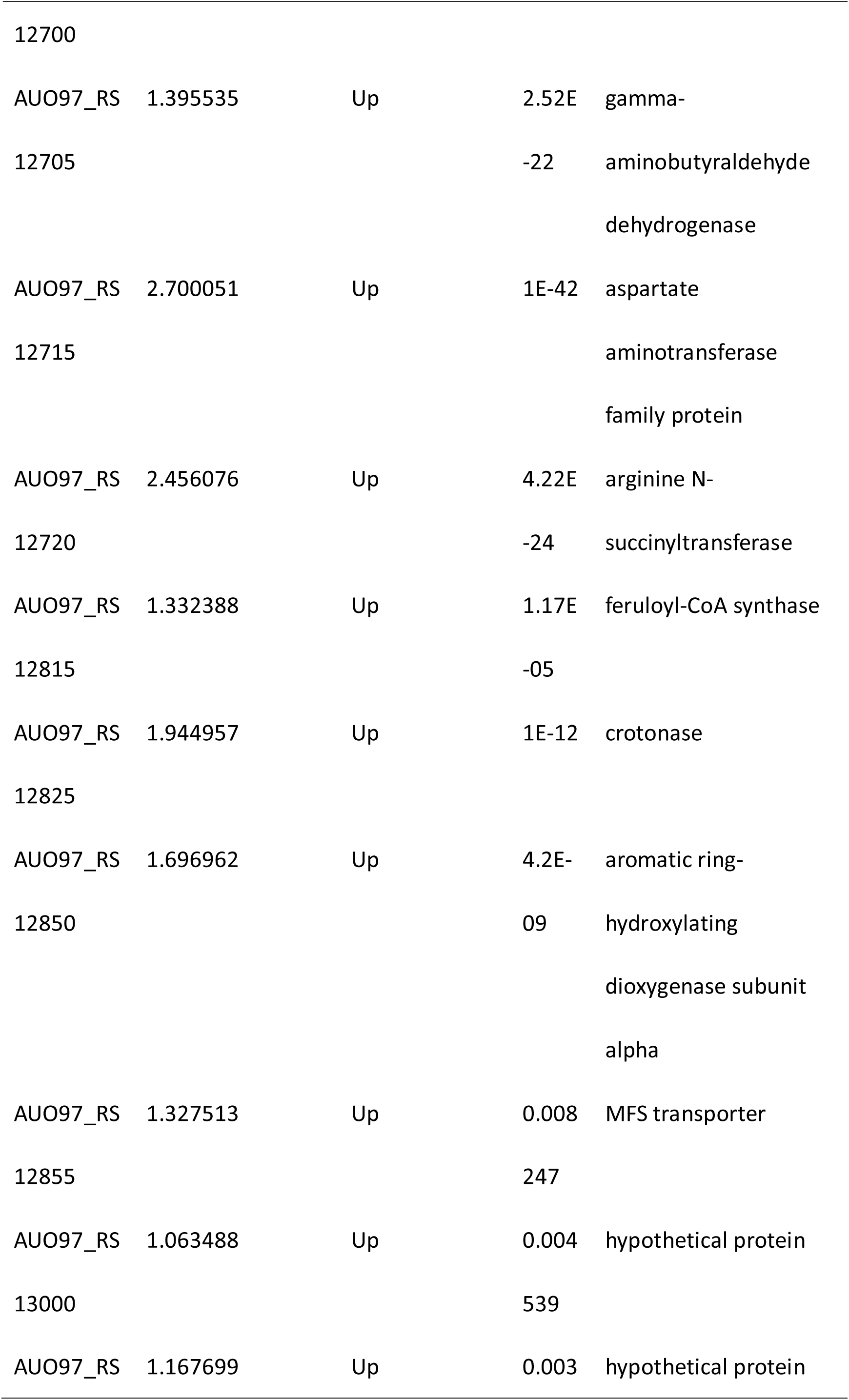
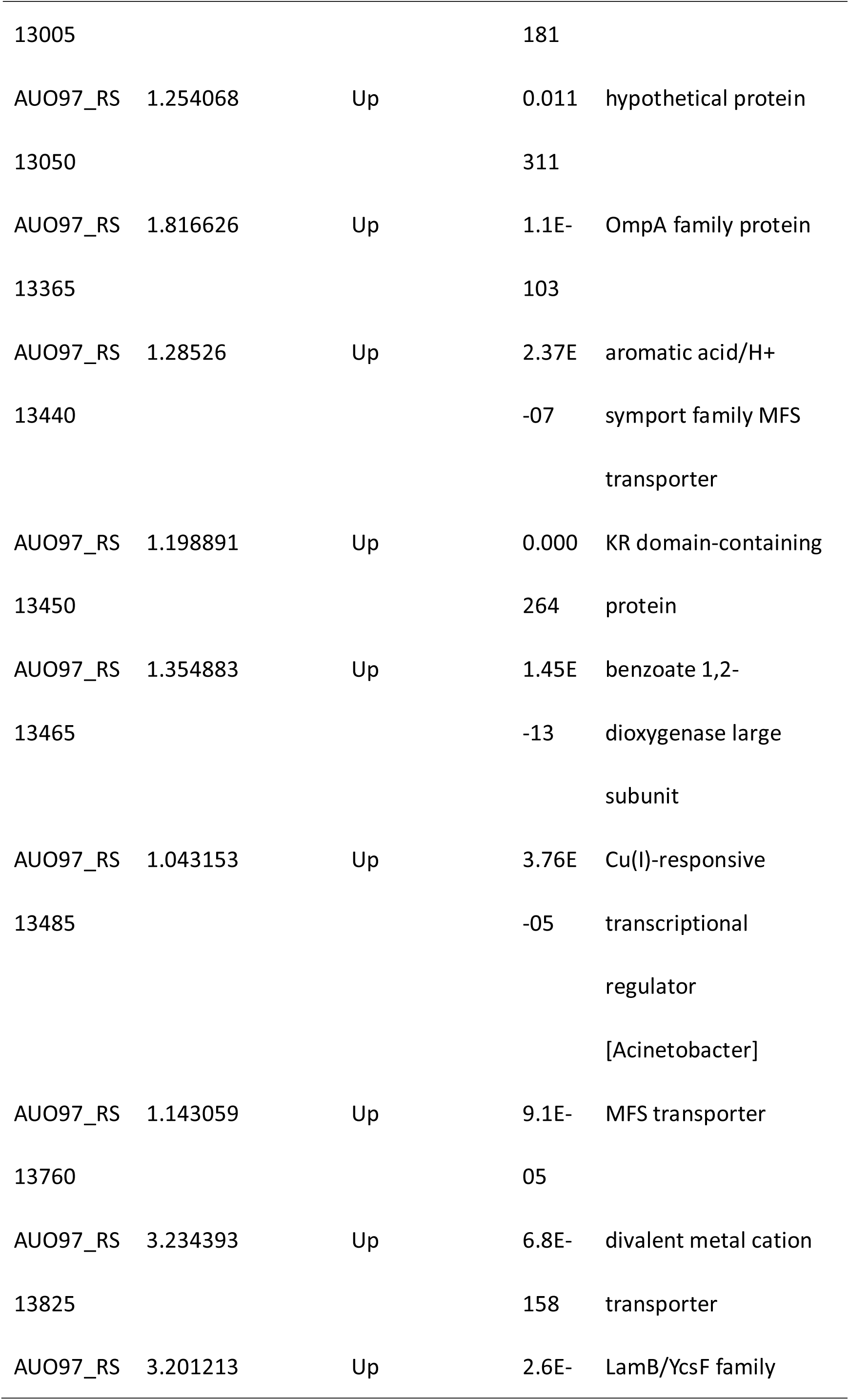
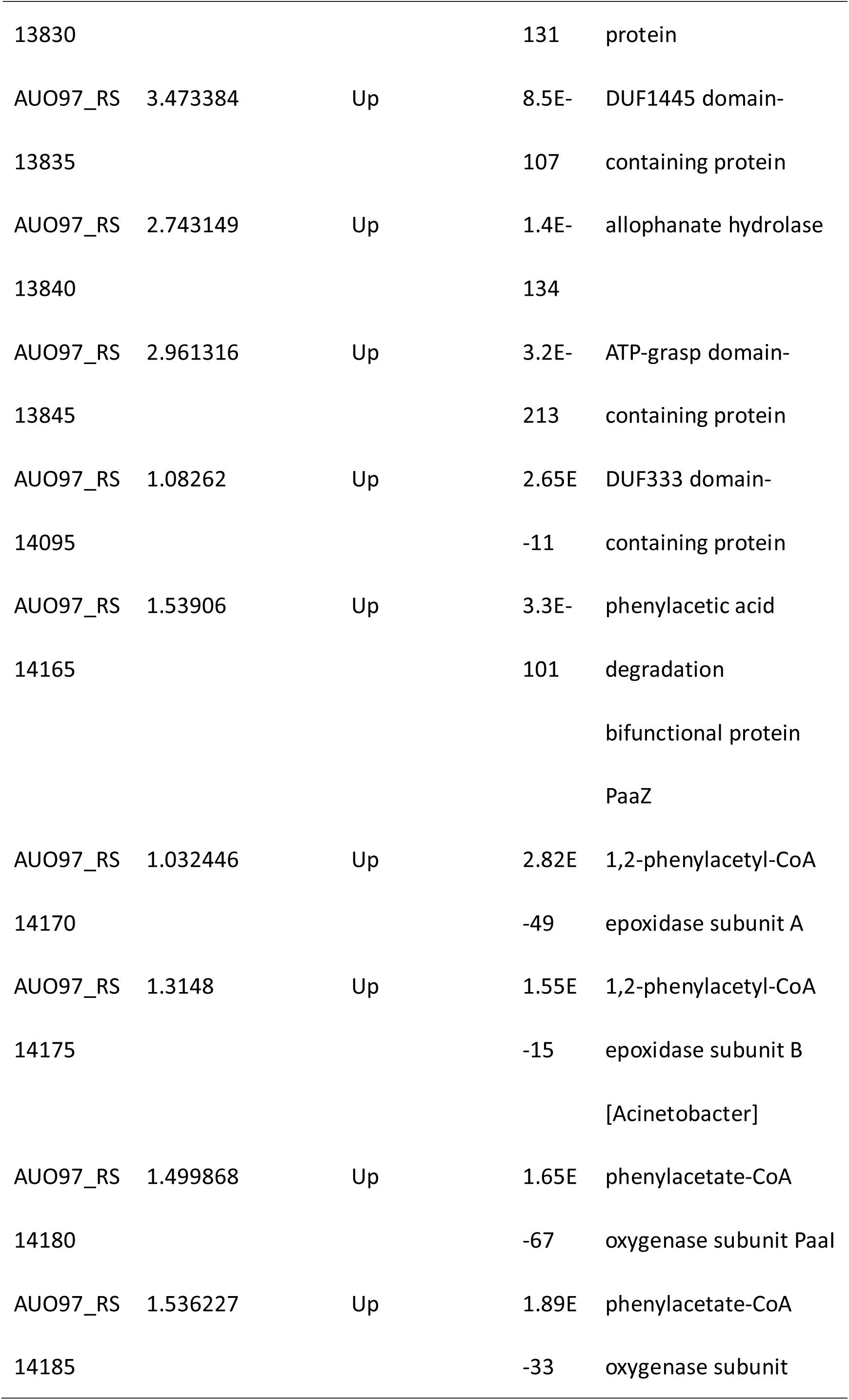
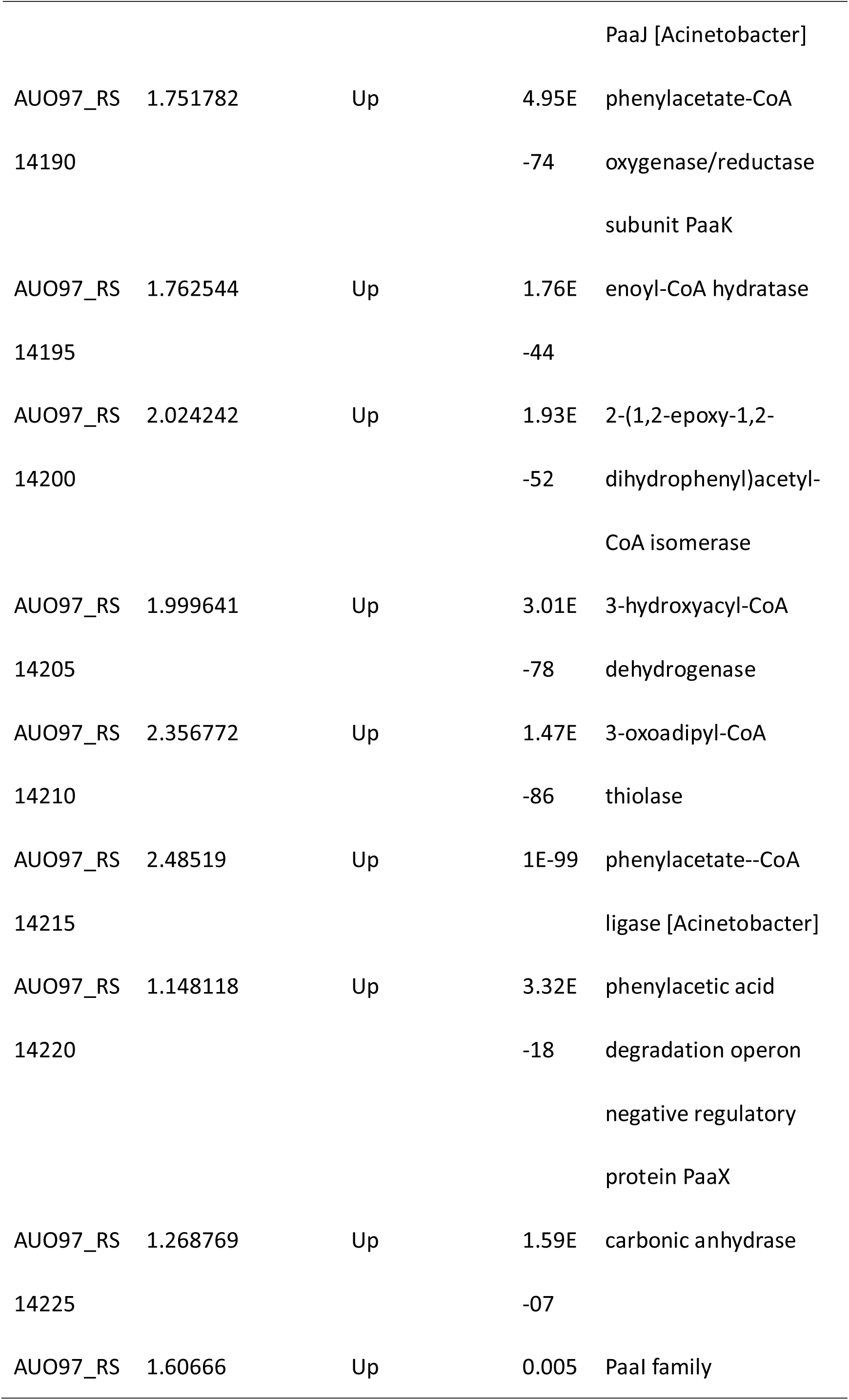
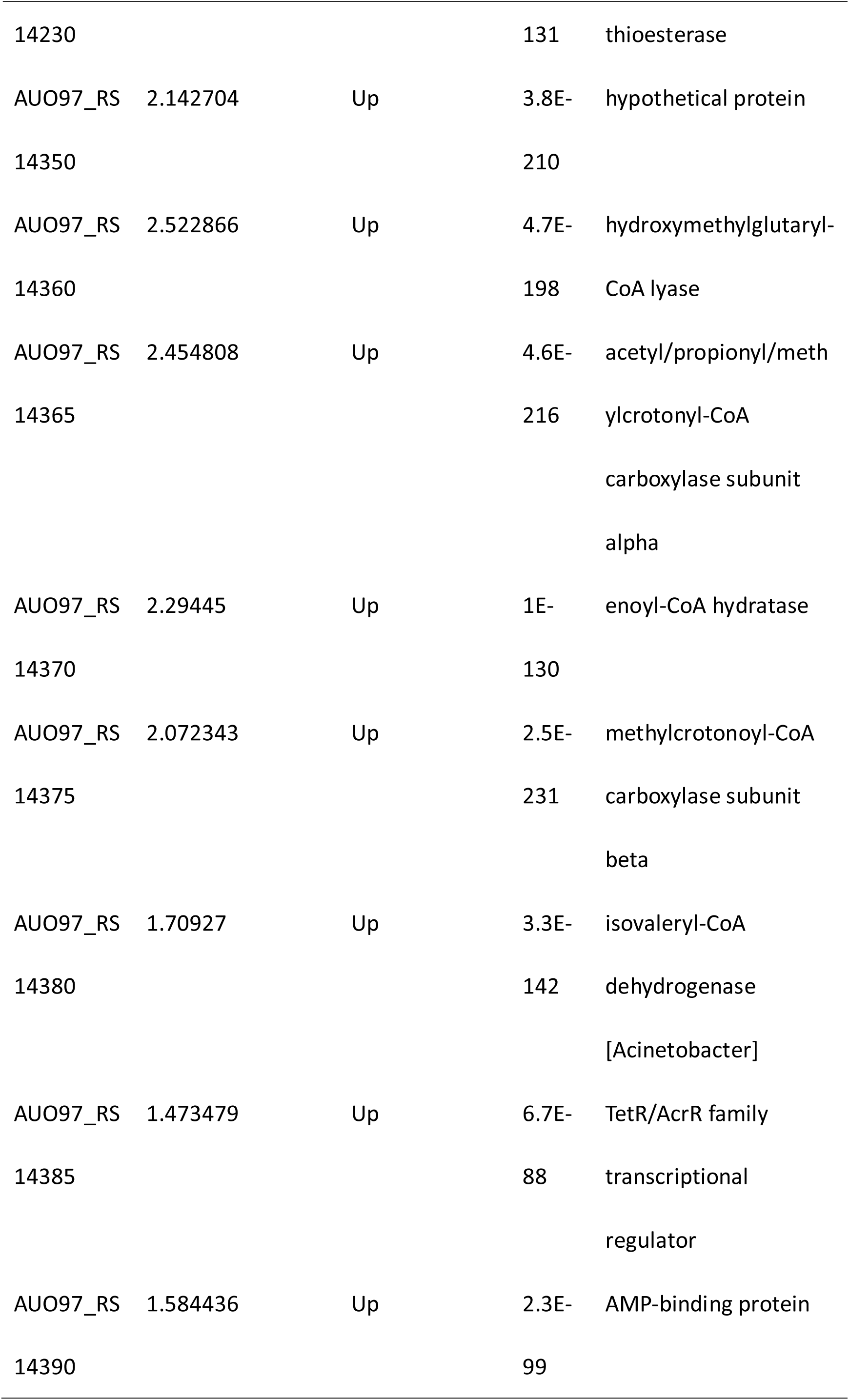
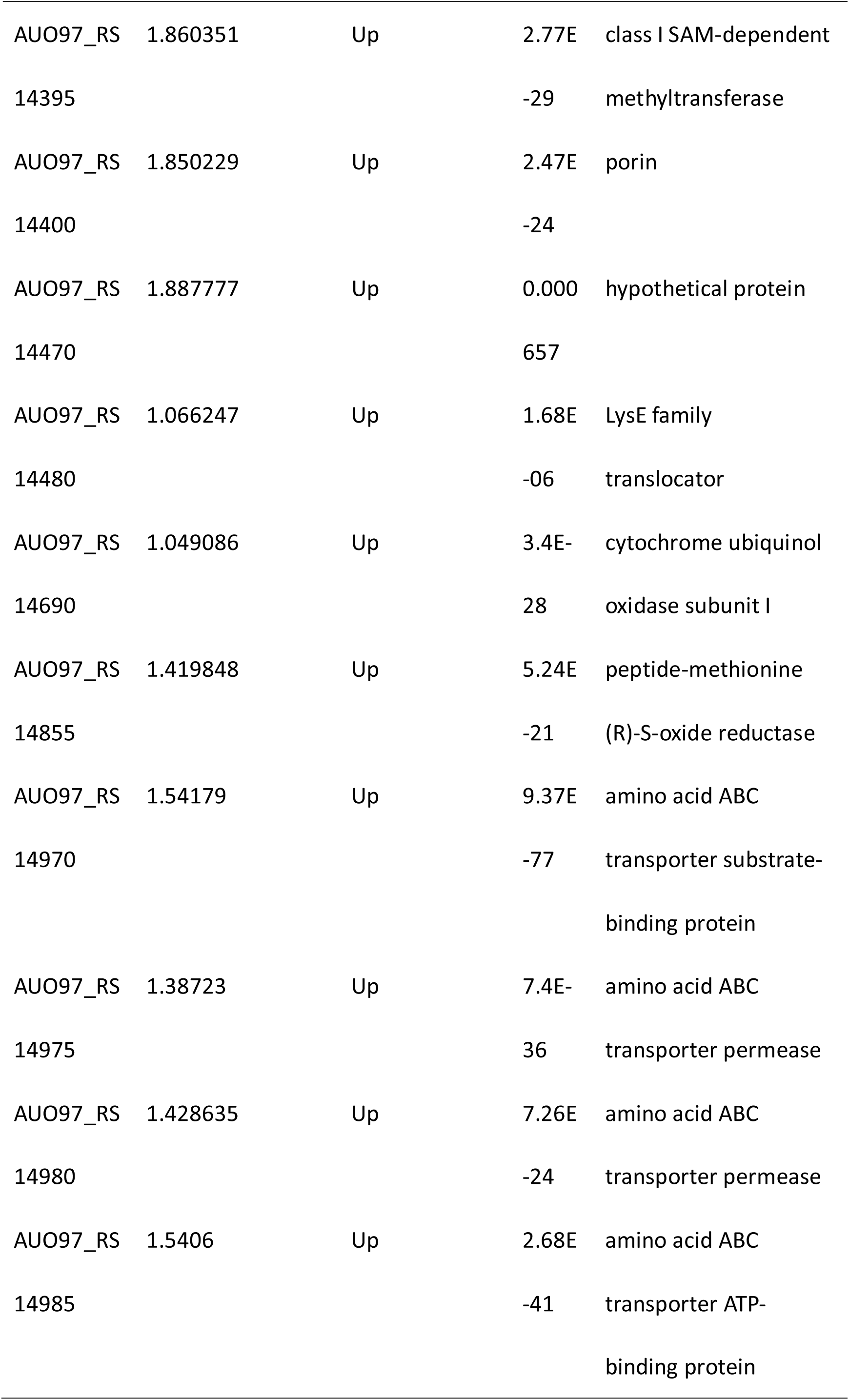
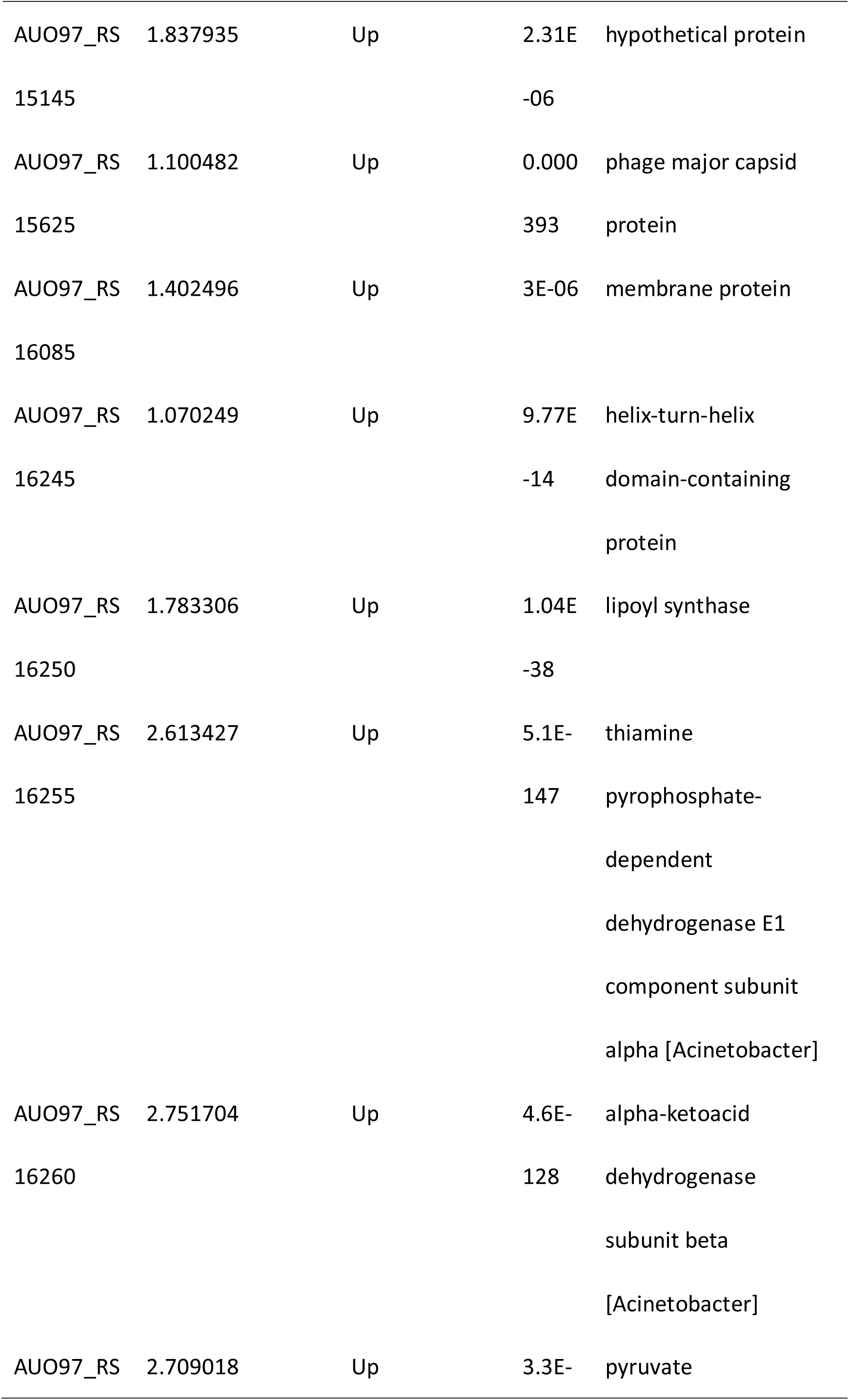
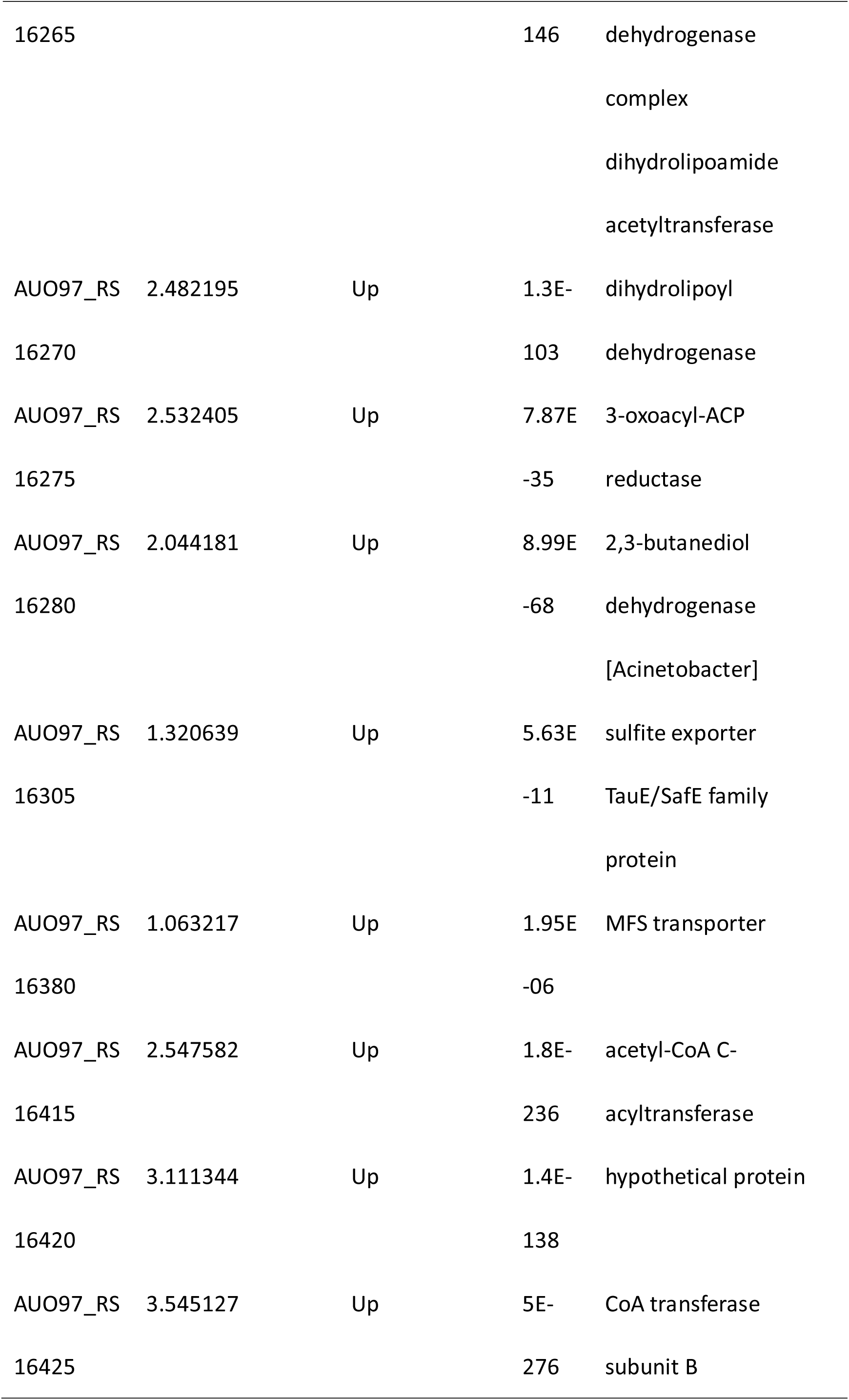
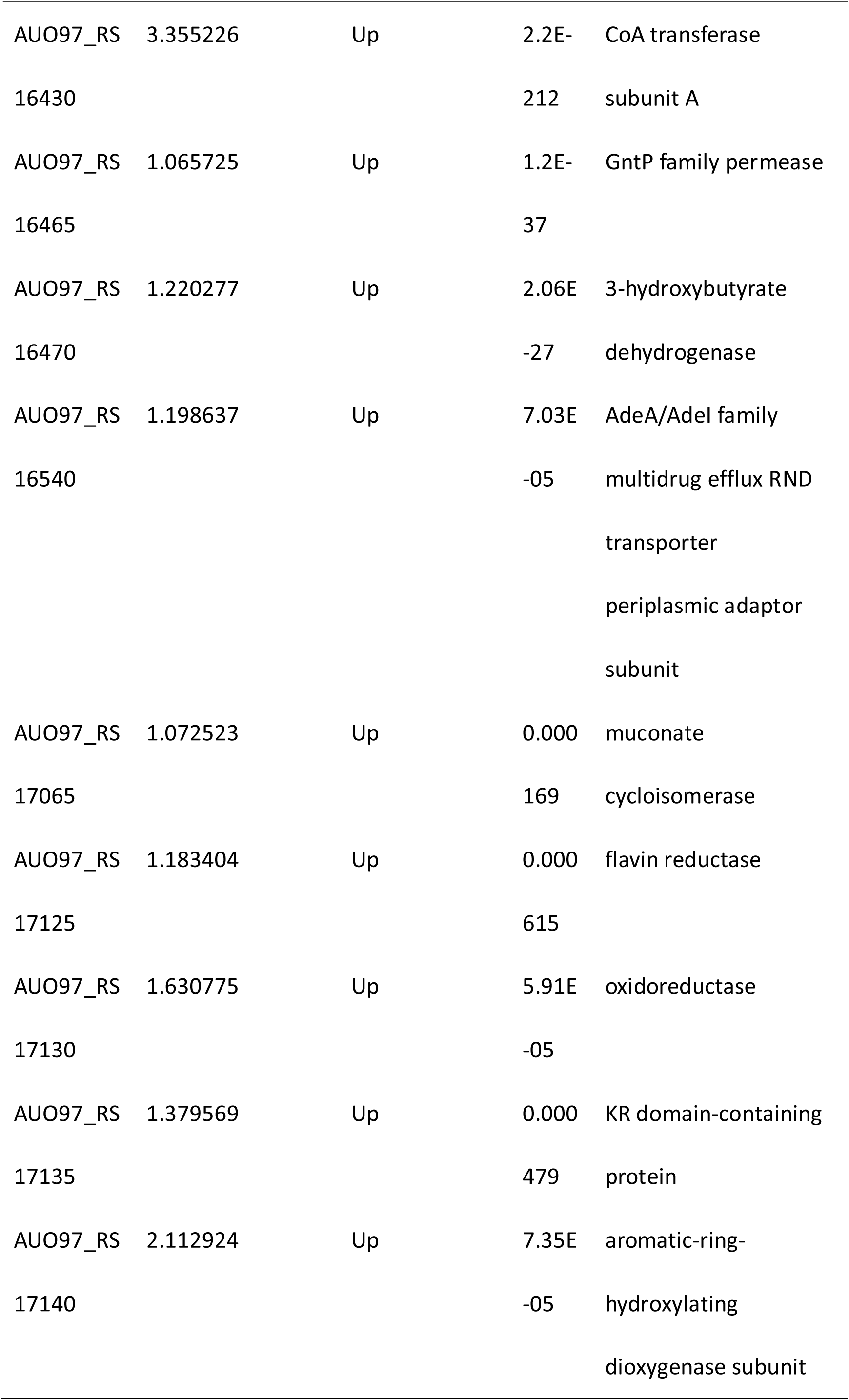
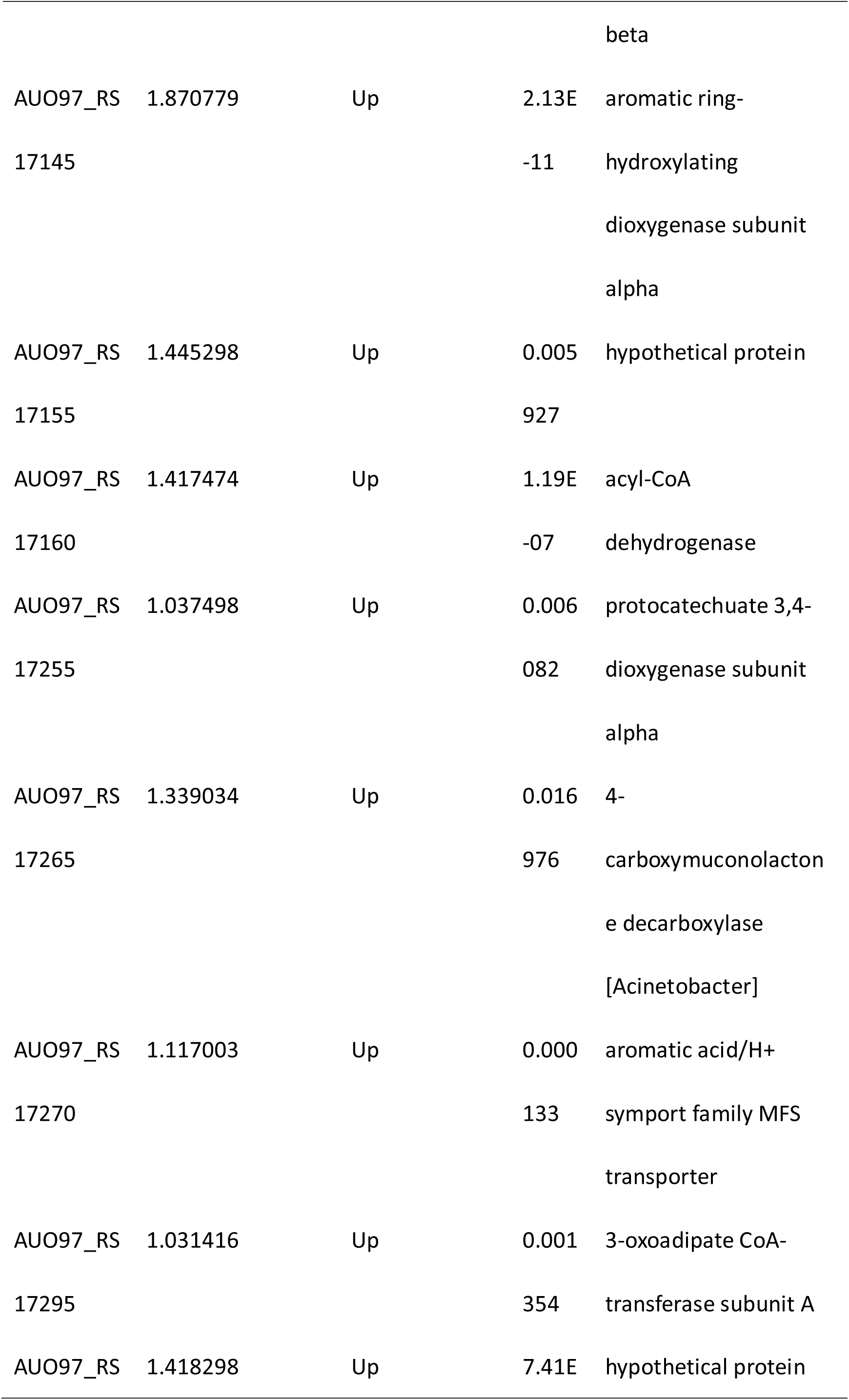
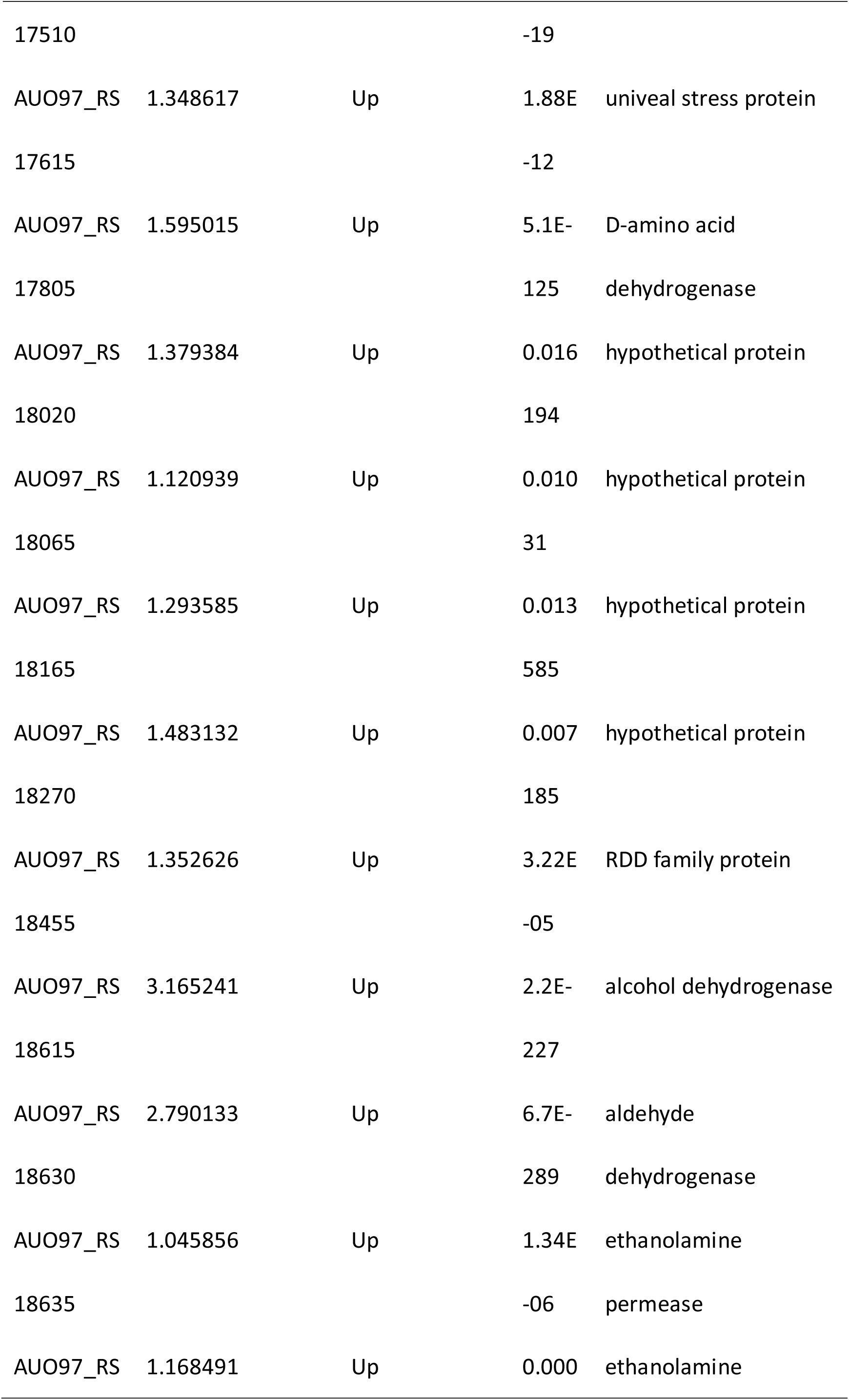
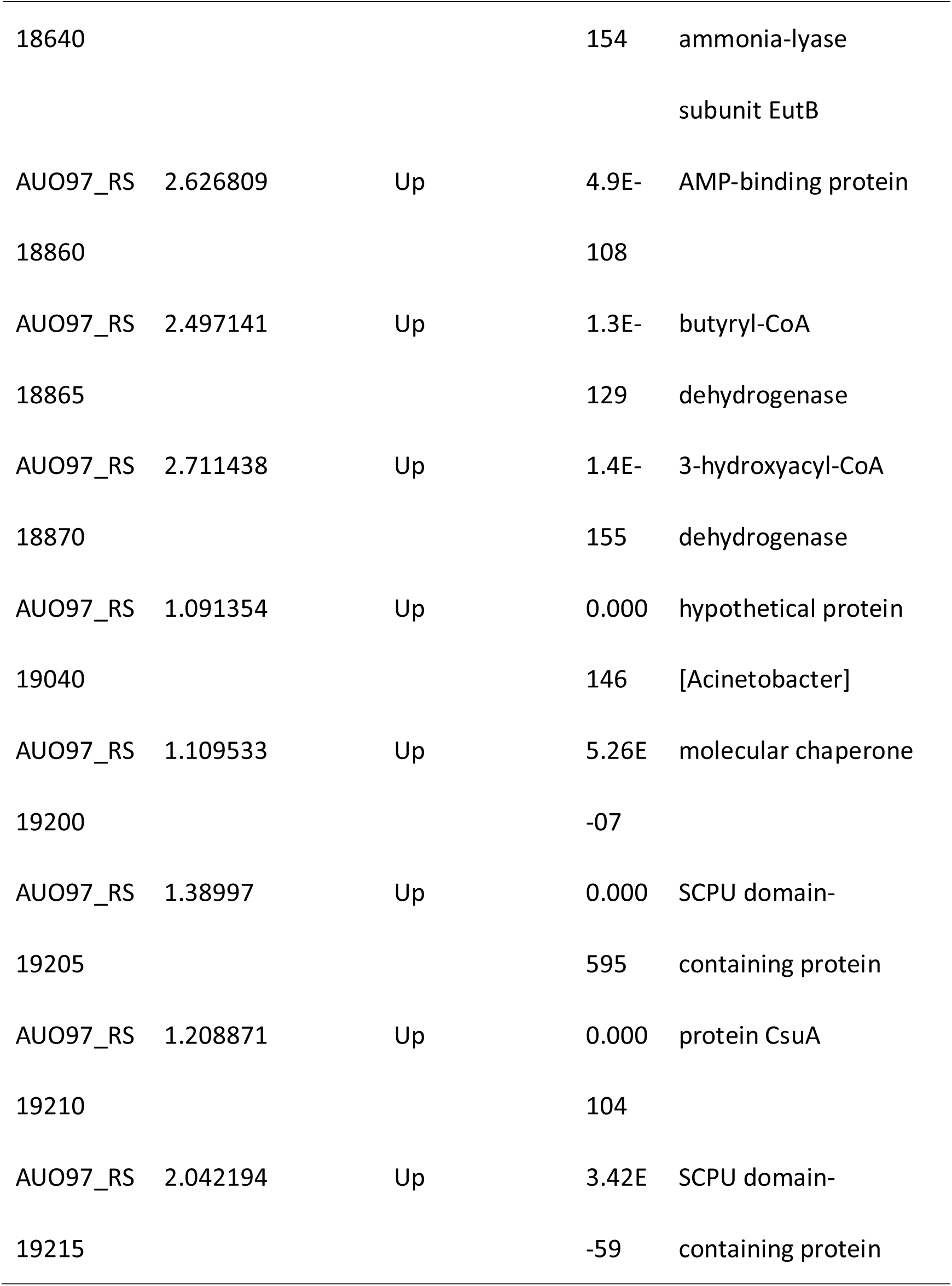
Differential expressed genes in Δ*abaR* strain

**Table 7.**
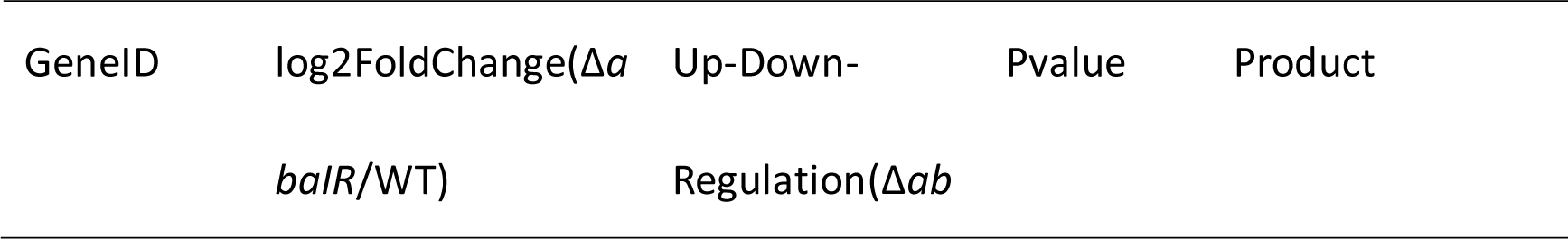
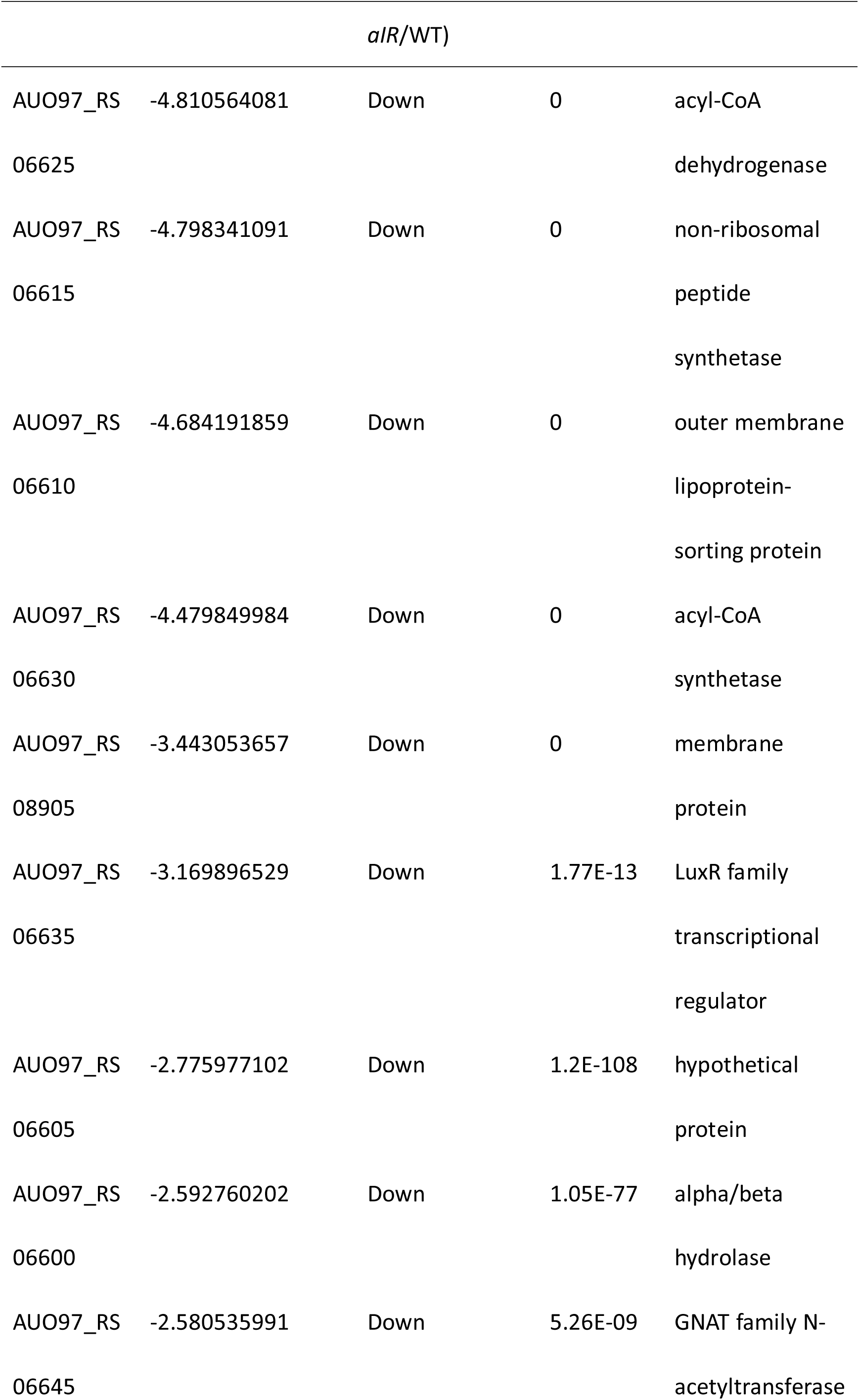
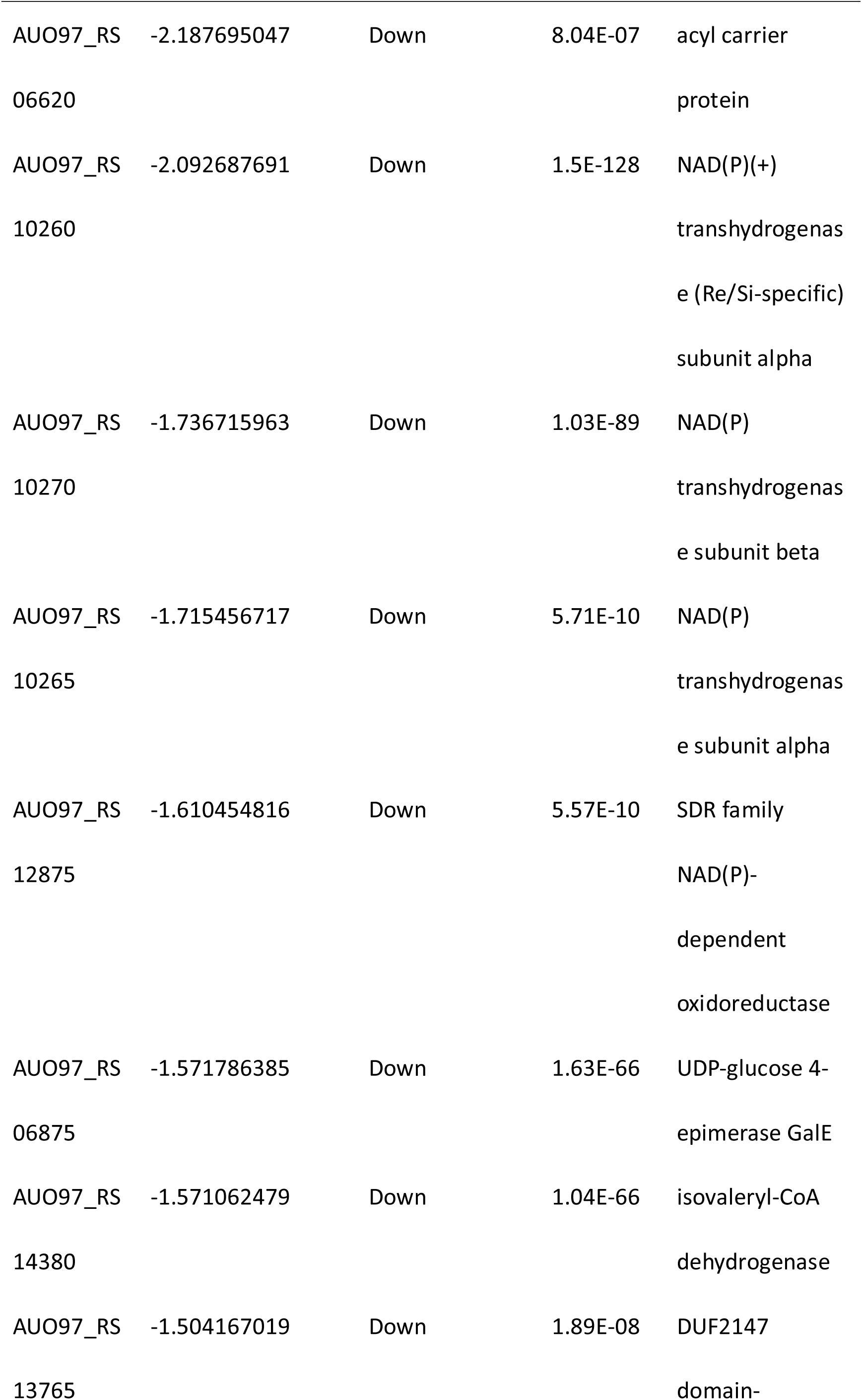
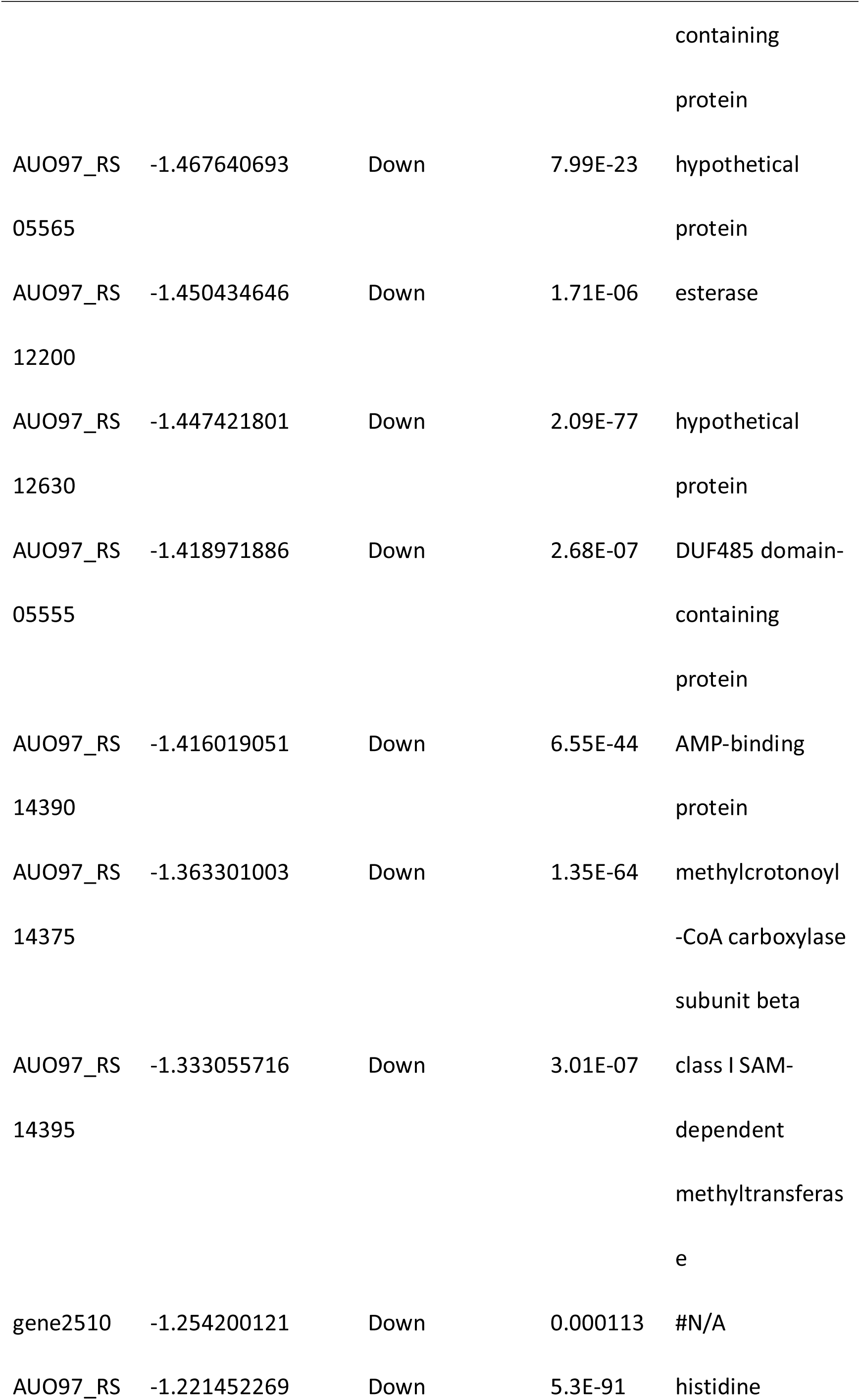
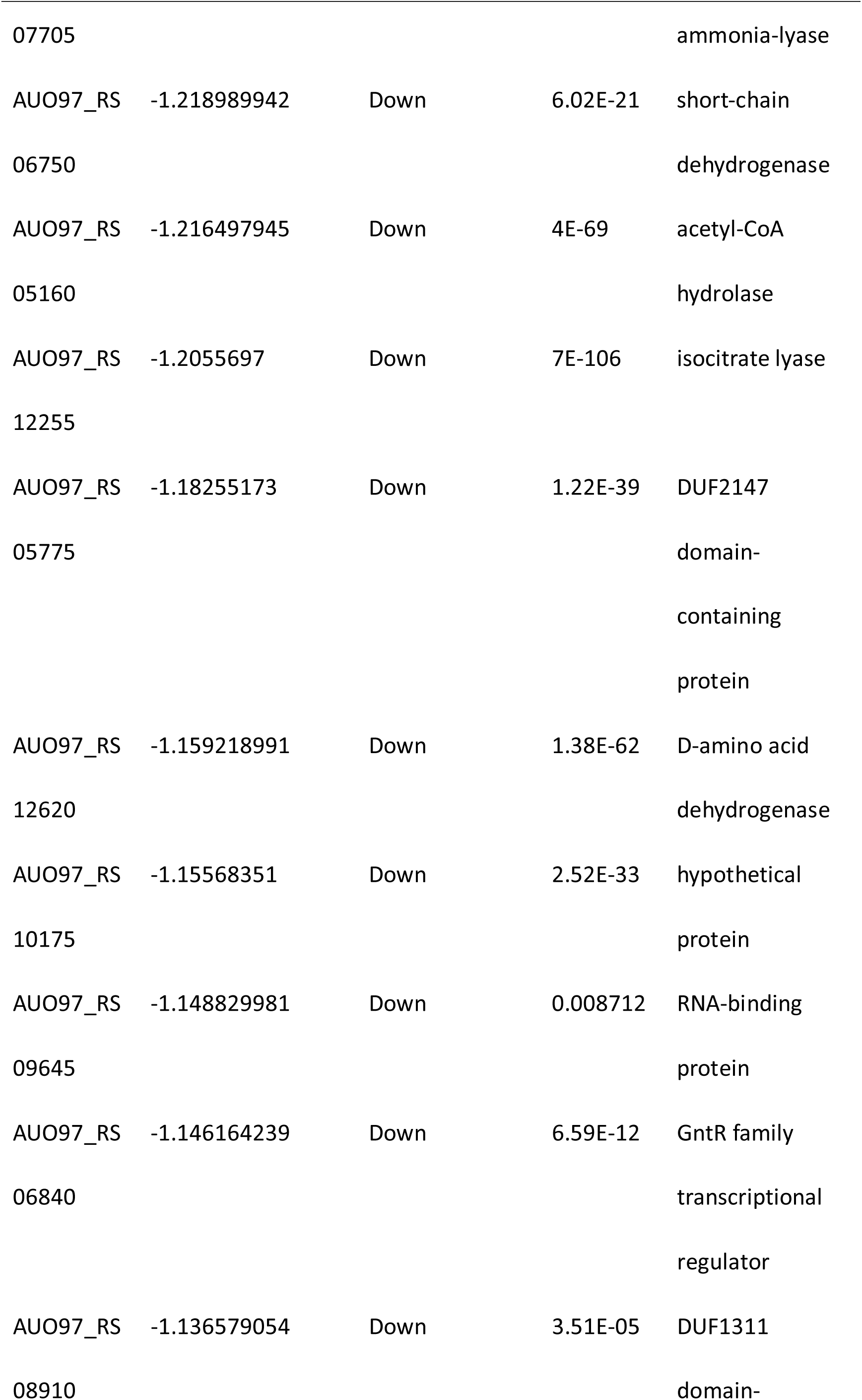
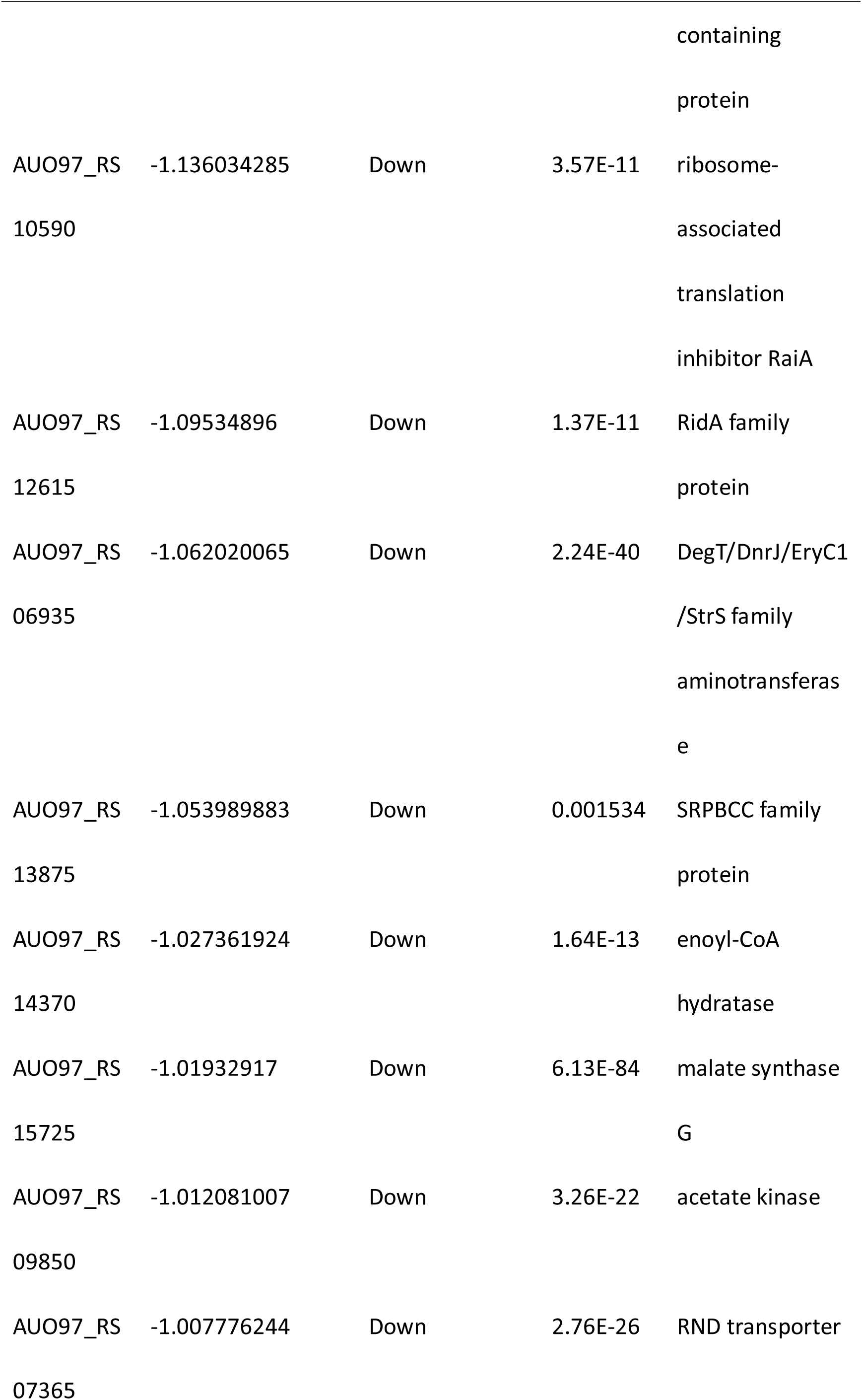
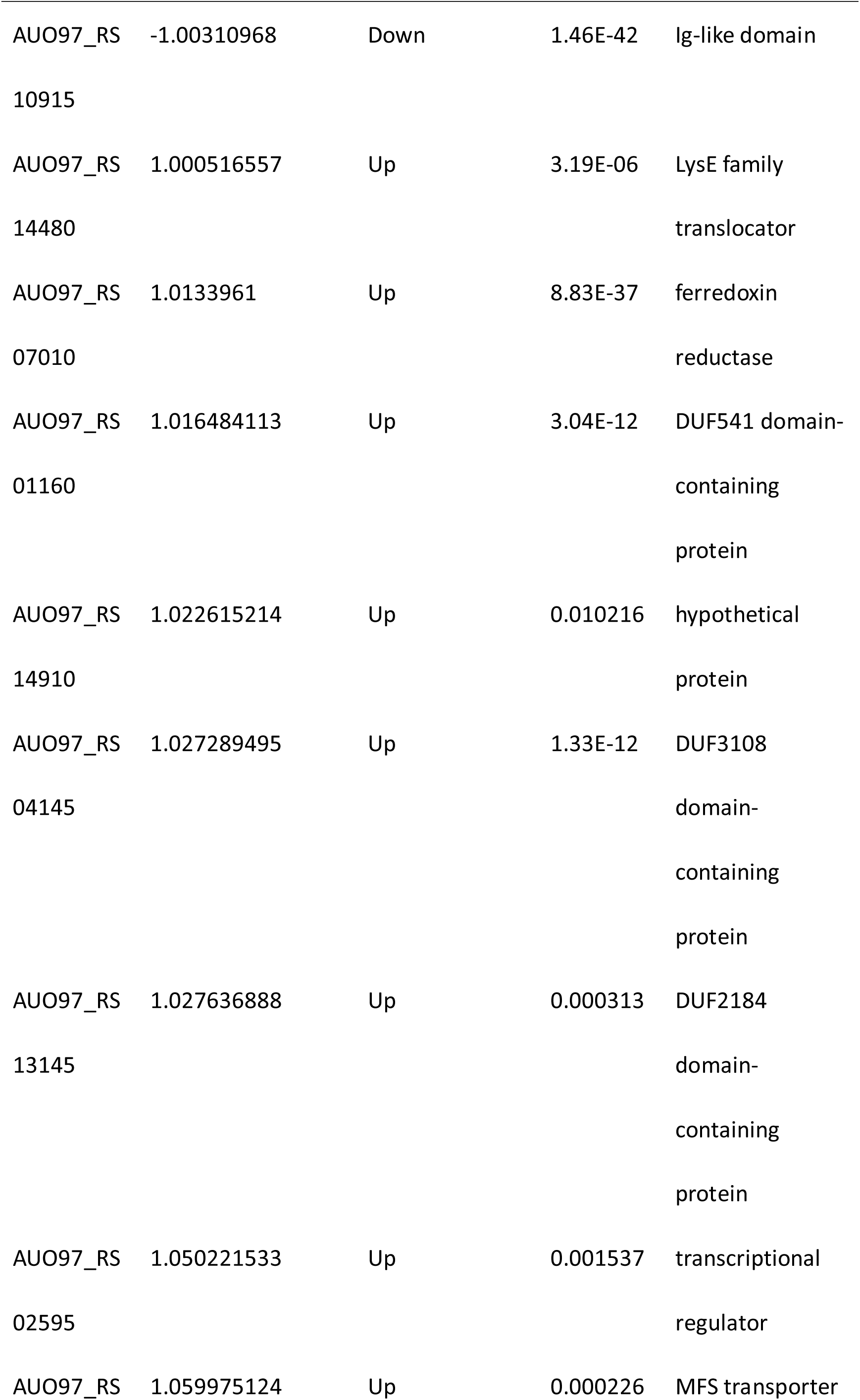
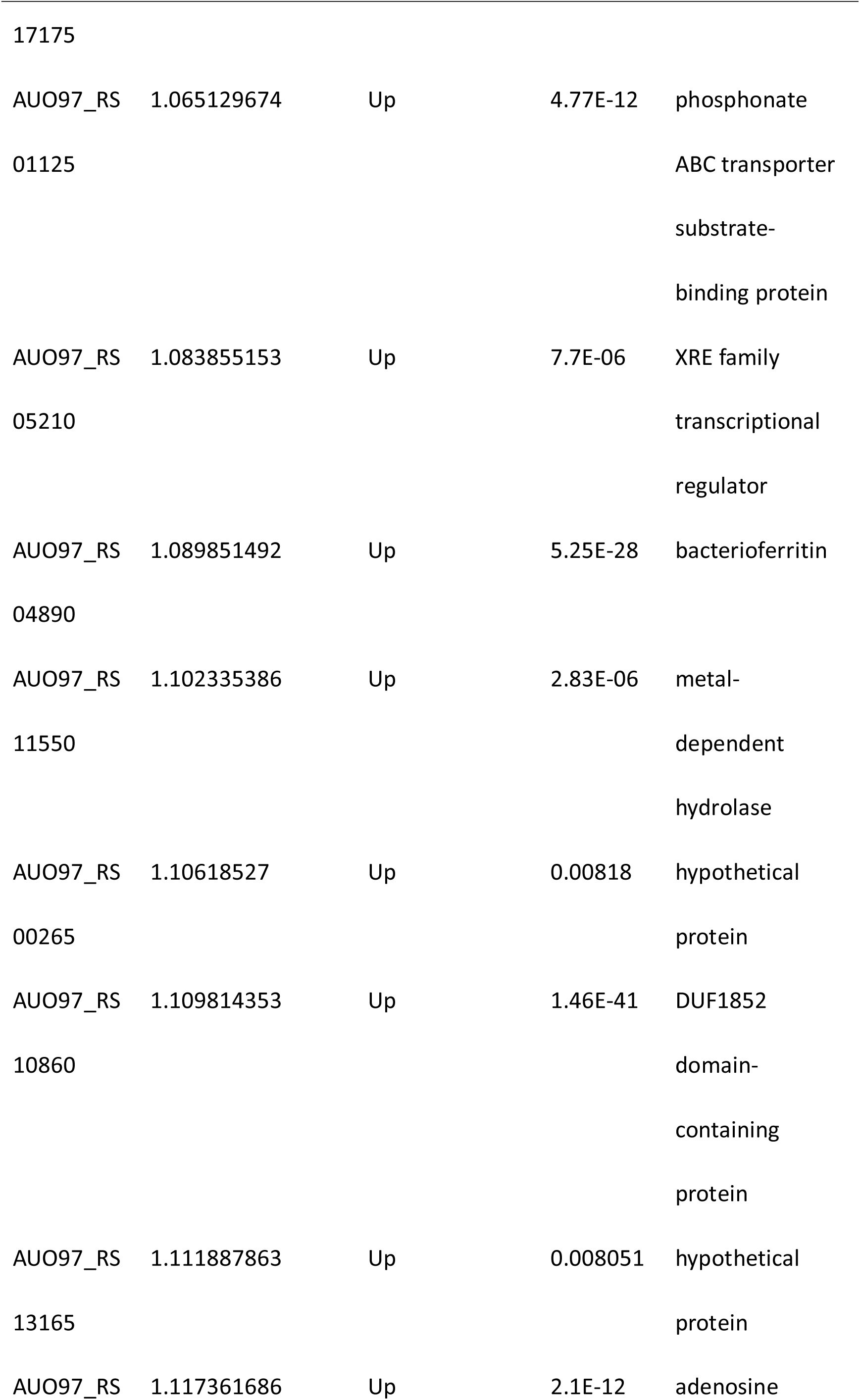
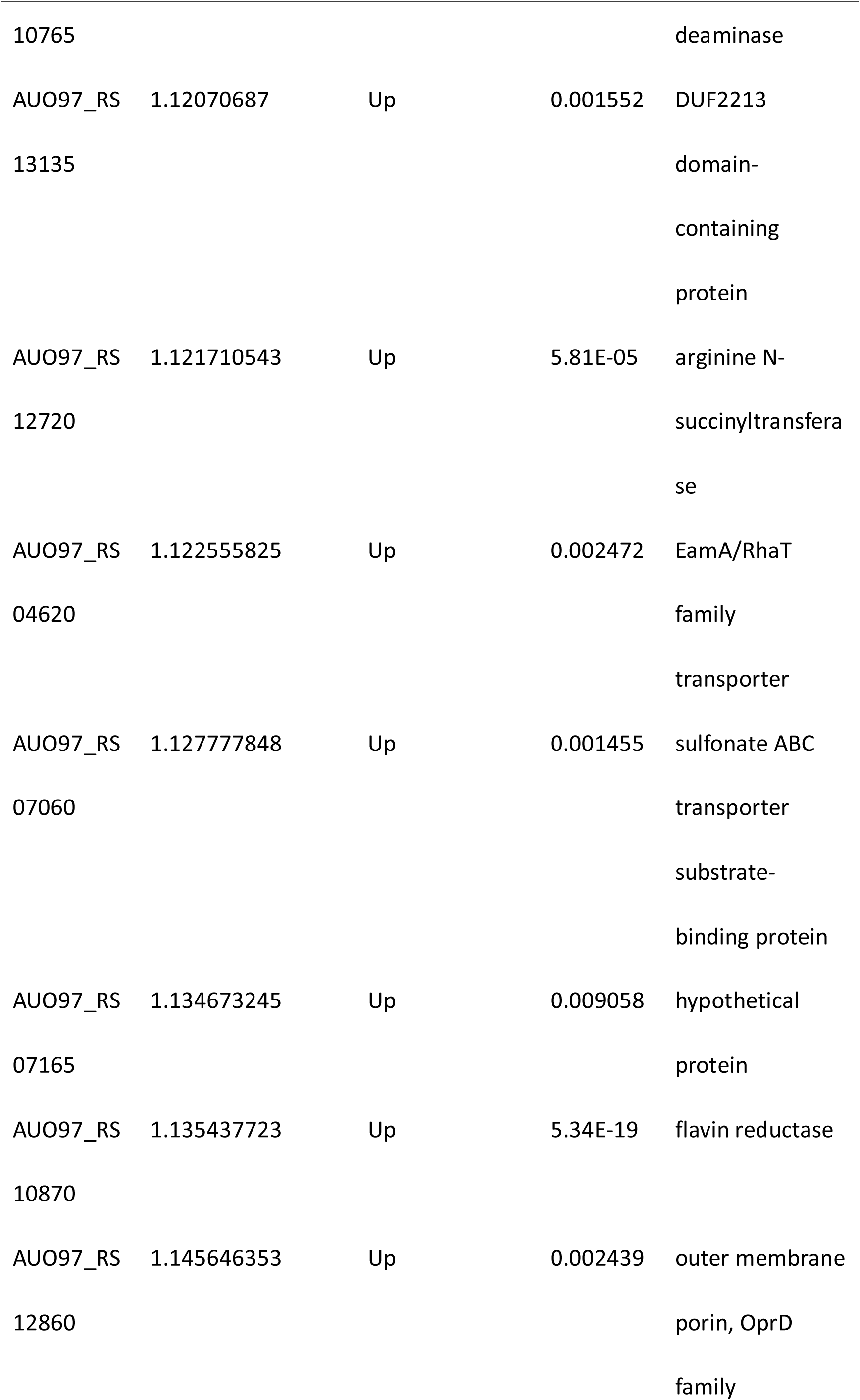
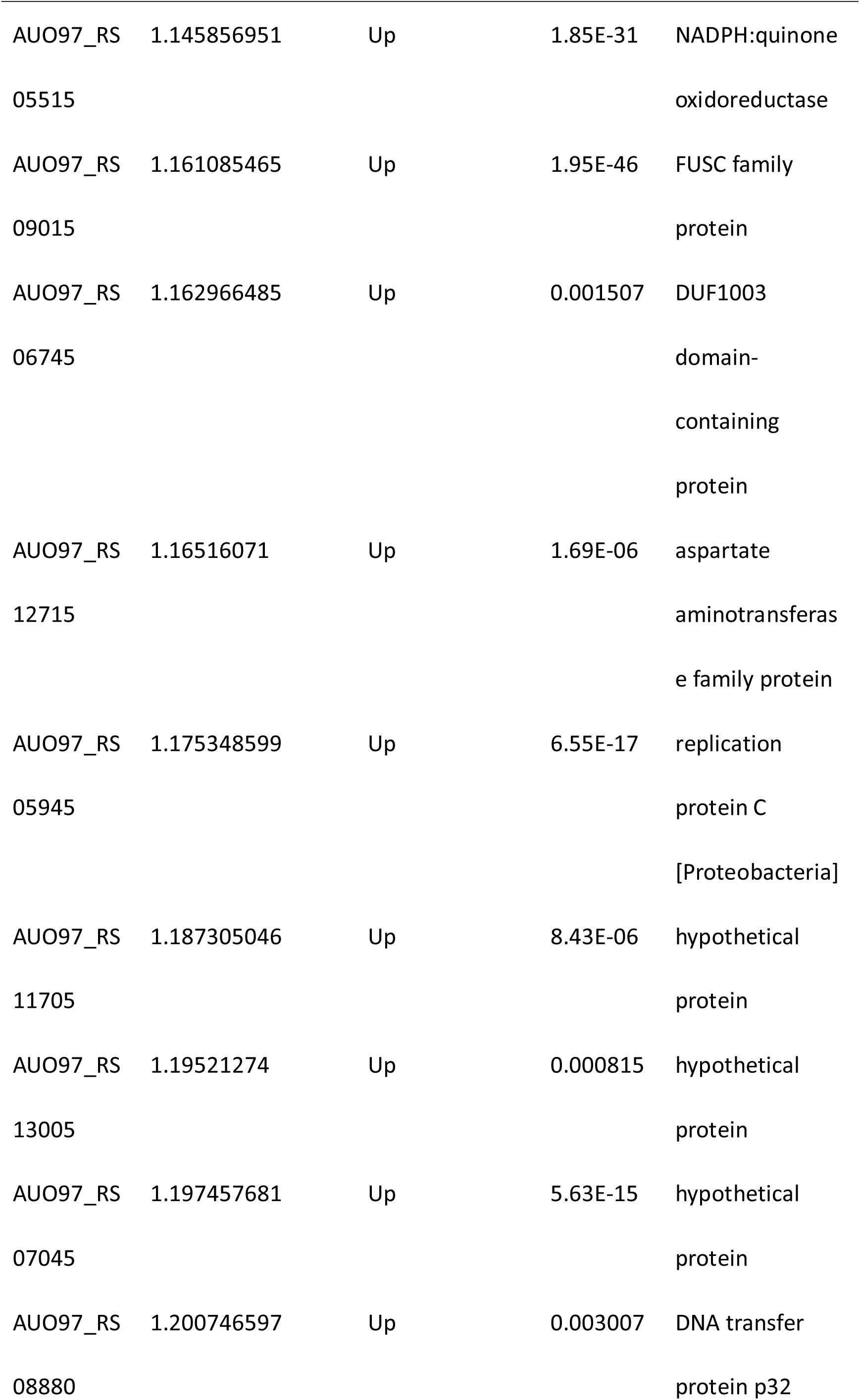
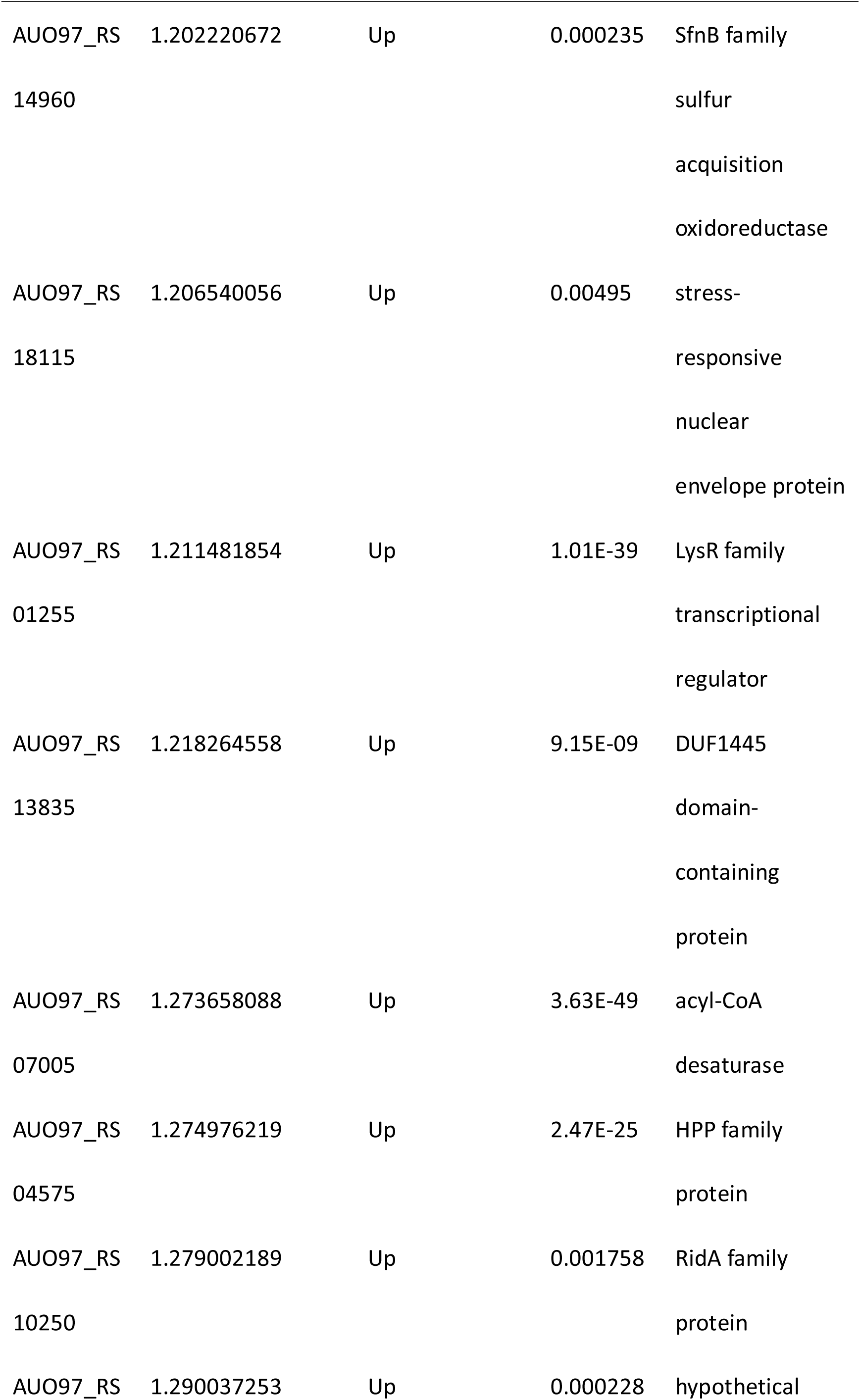
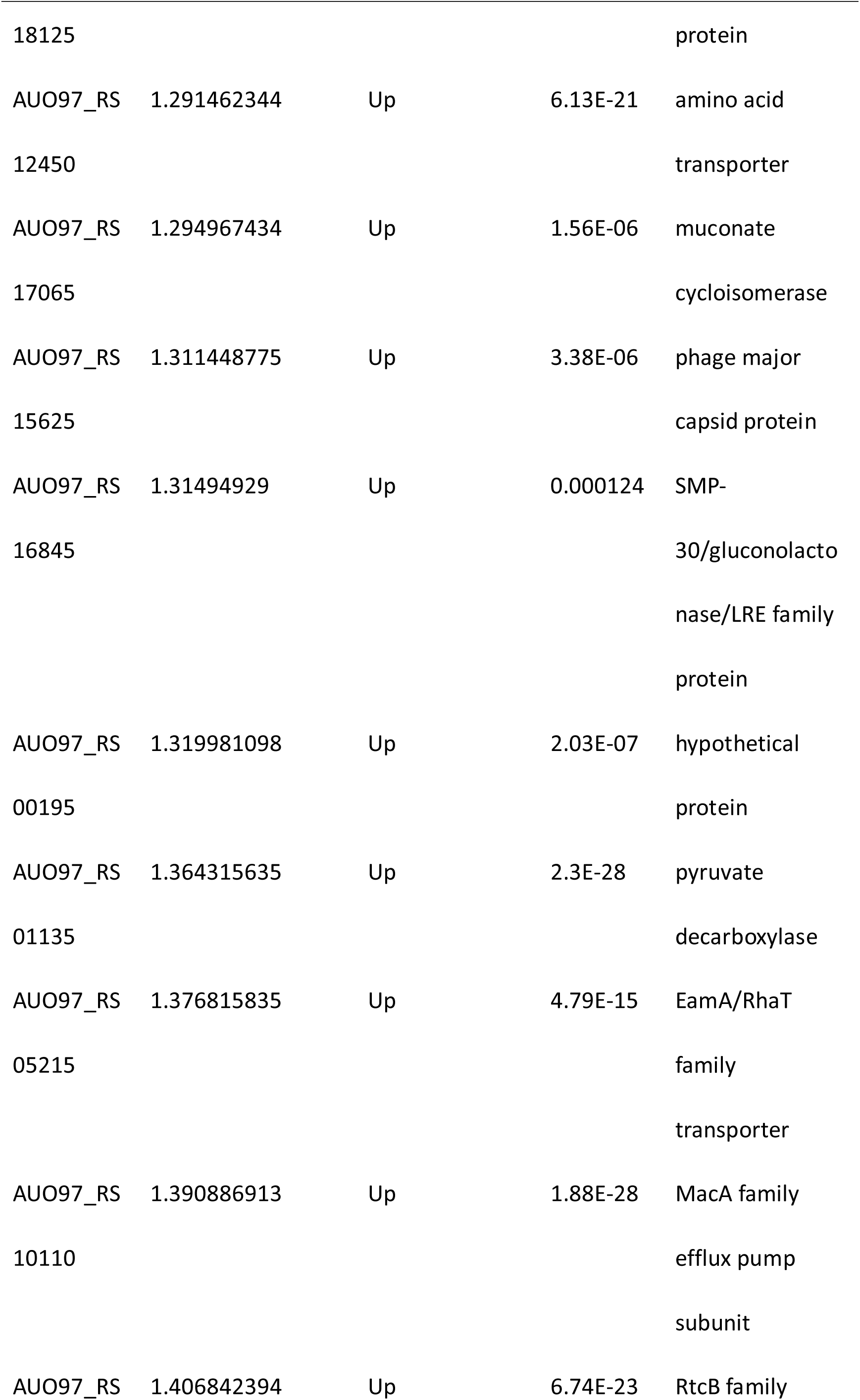
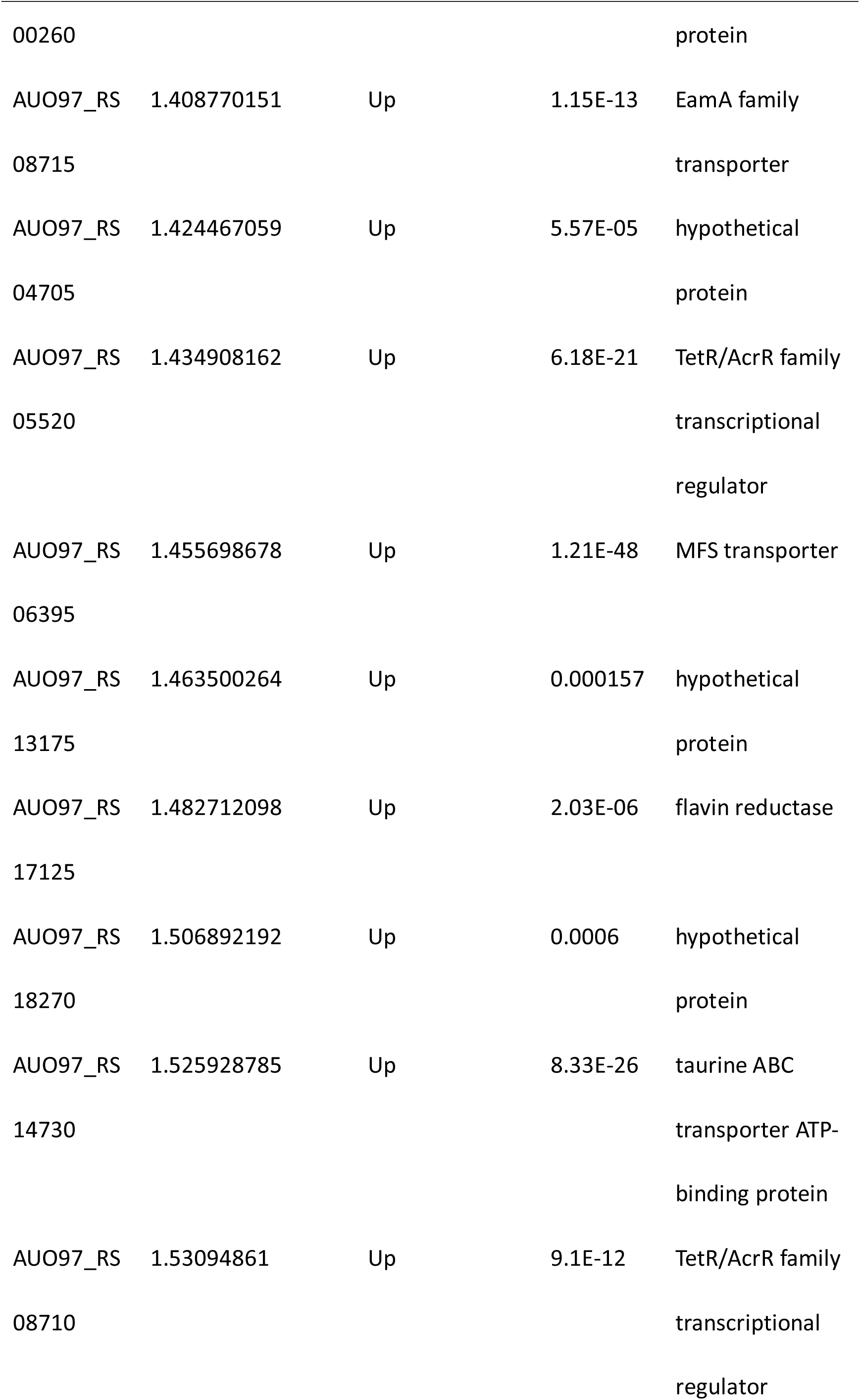
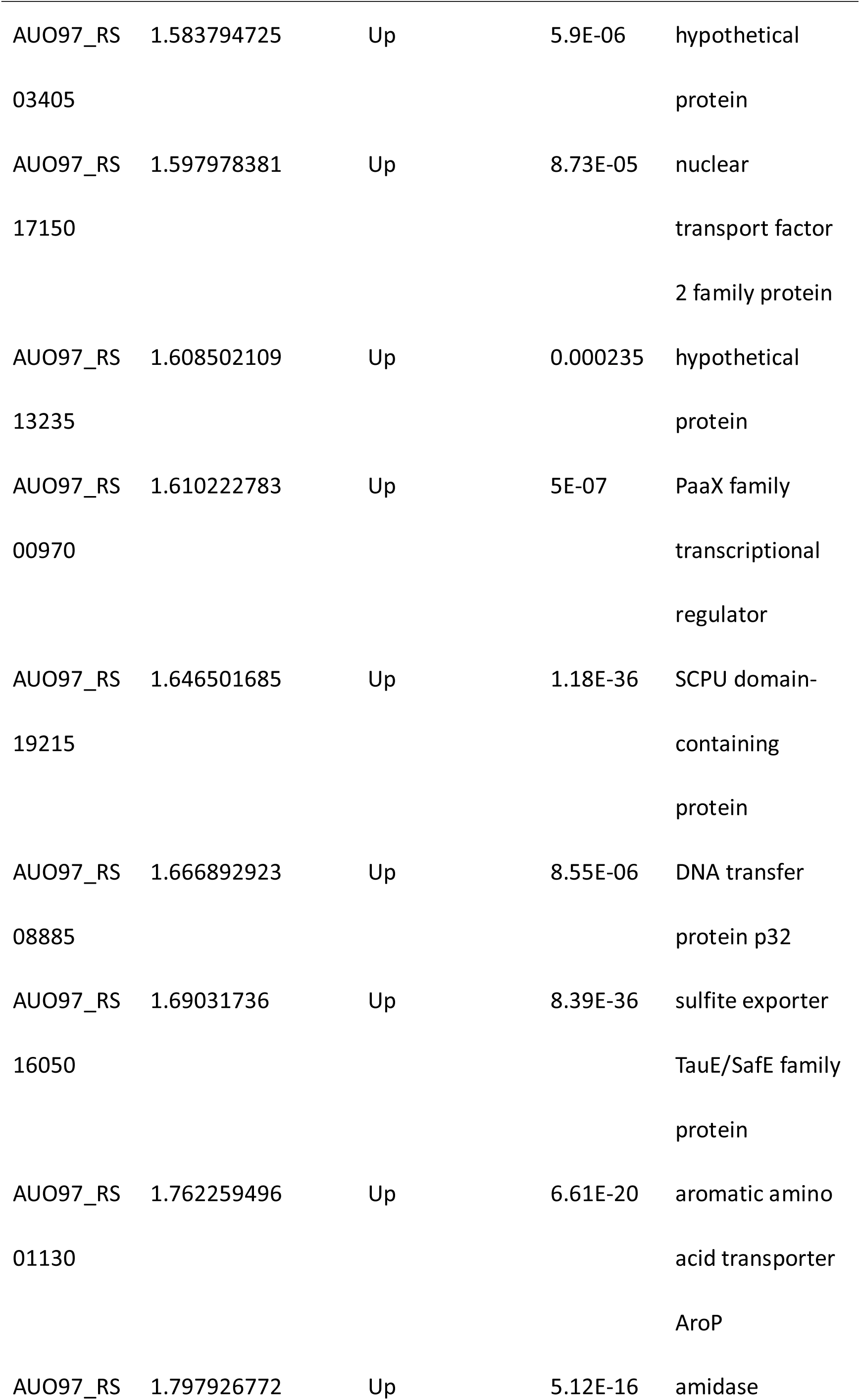
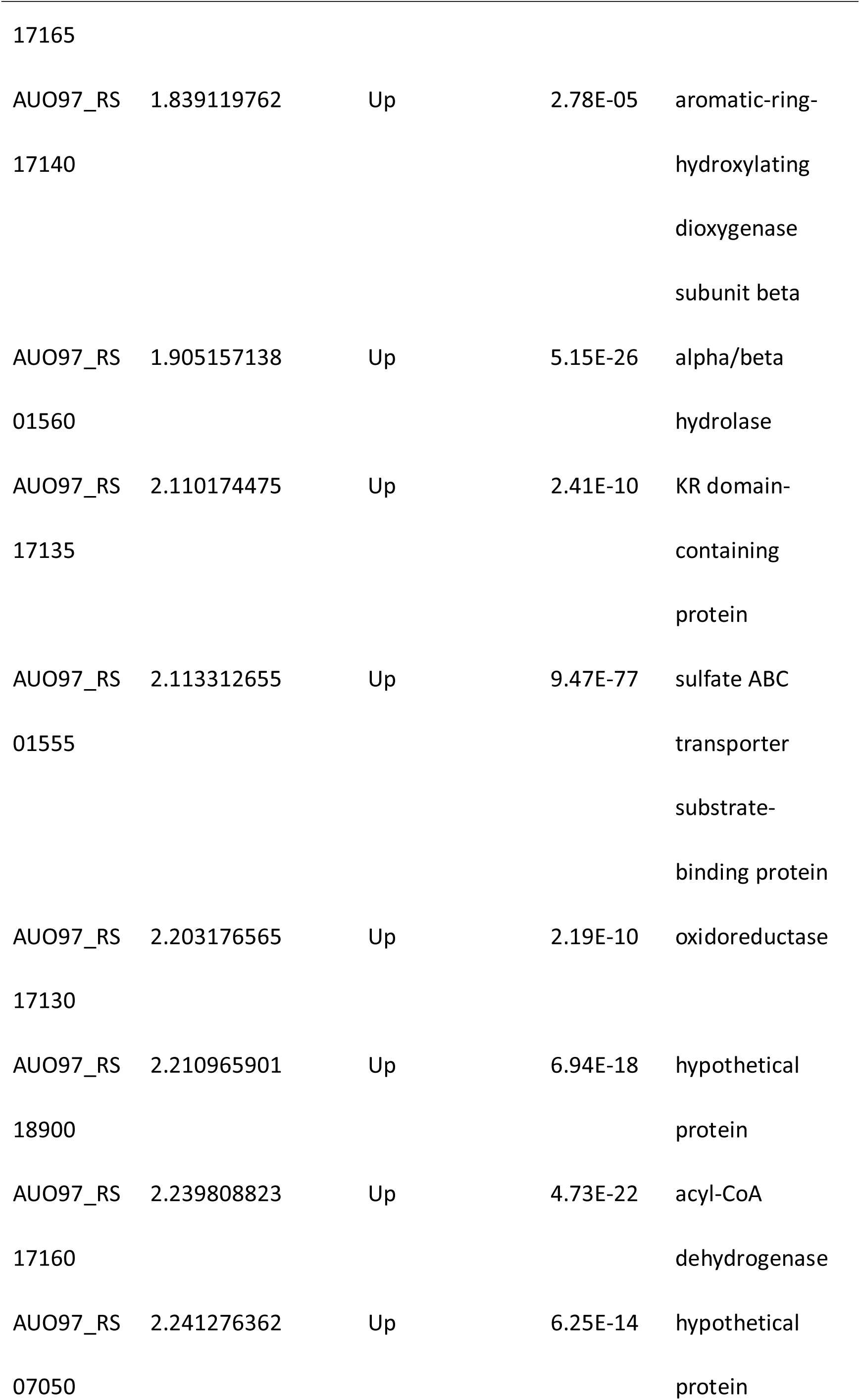
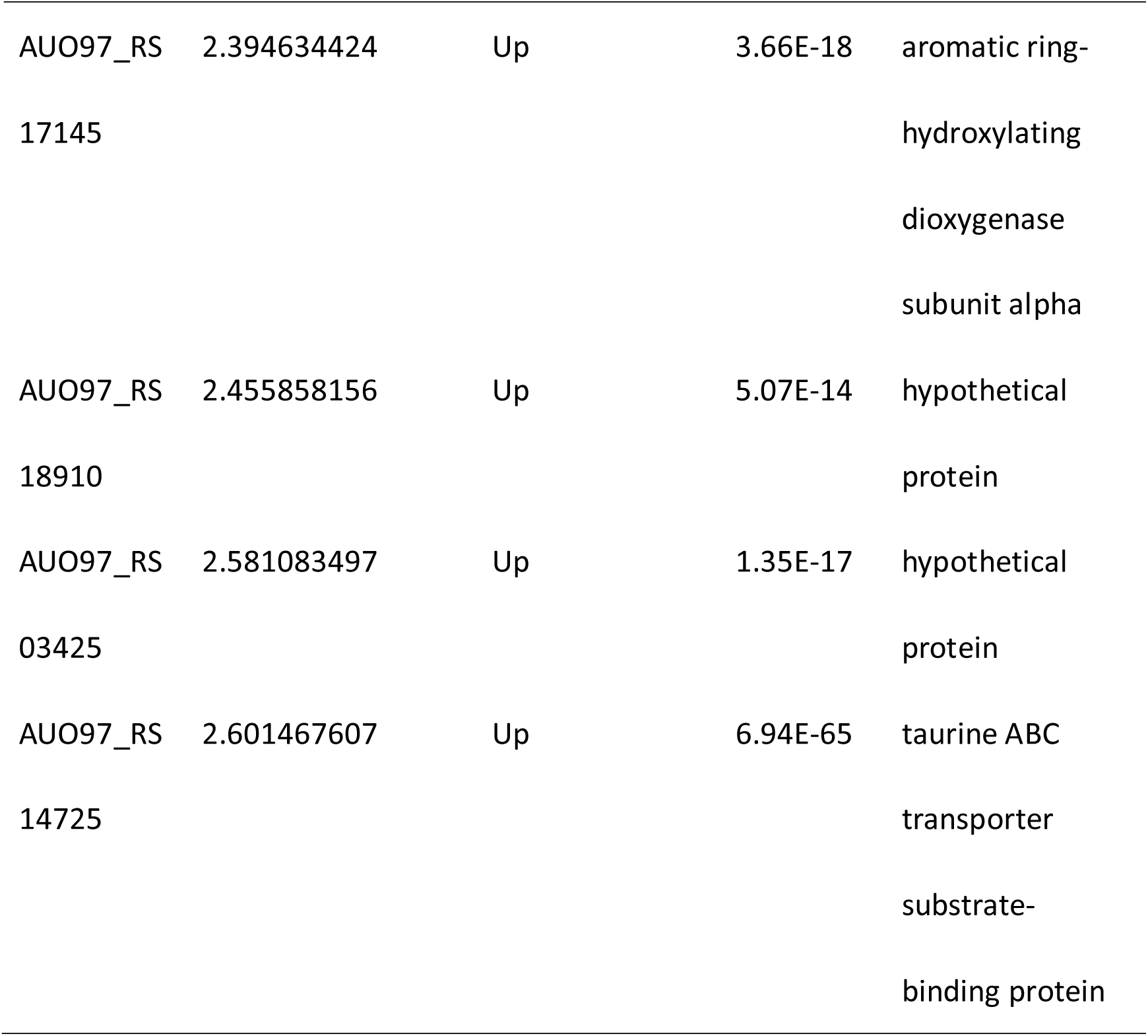
Differential expressed genes in Δ*abaIR* strain

**Table 4.**
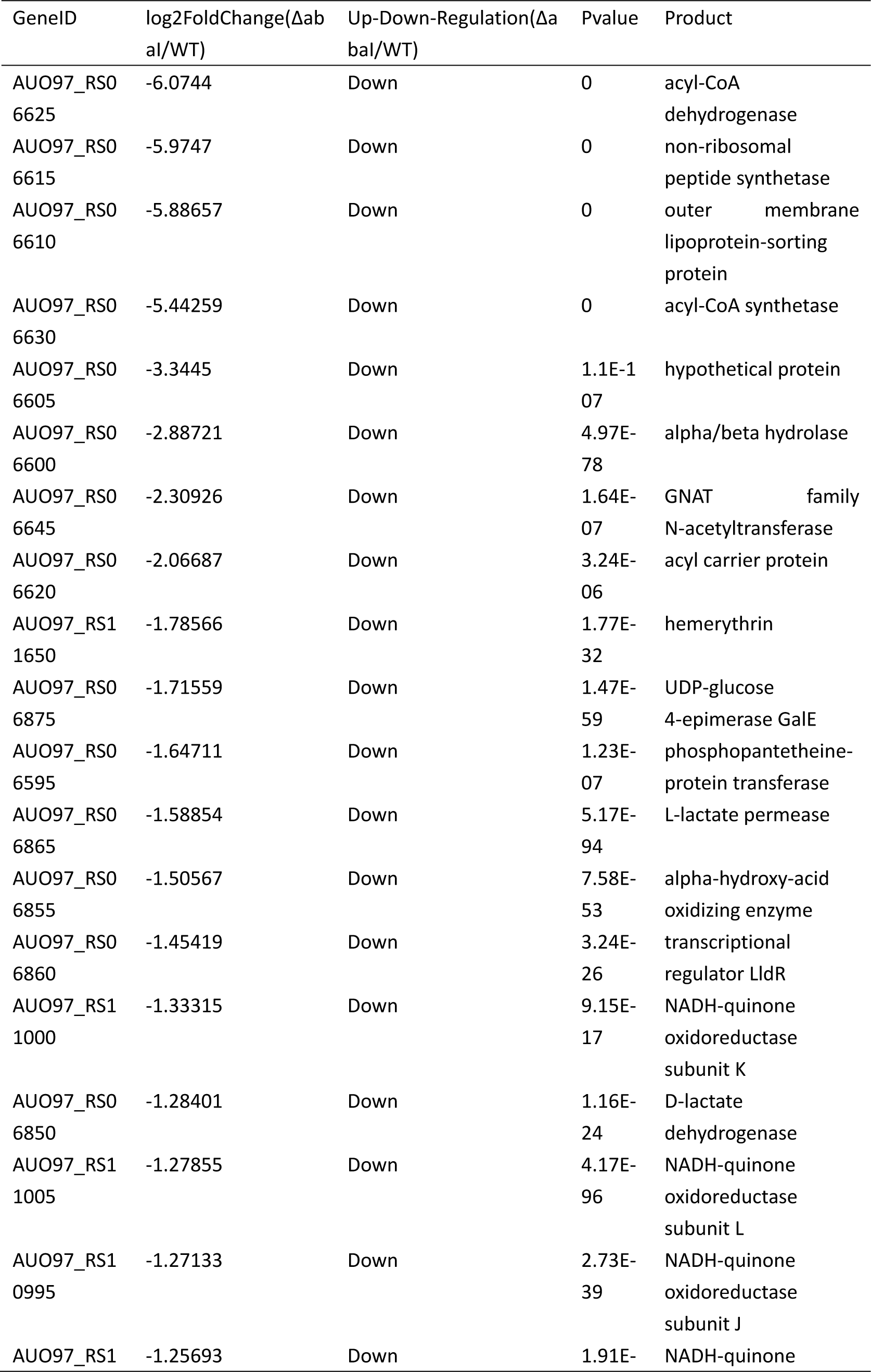
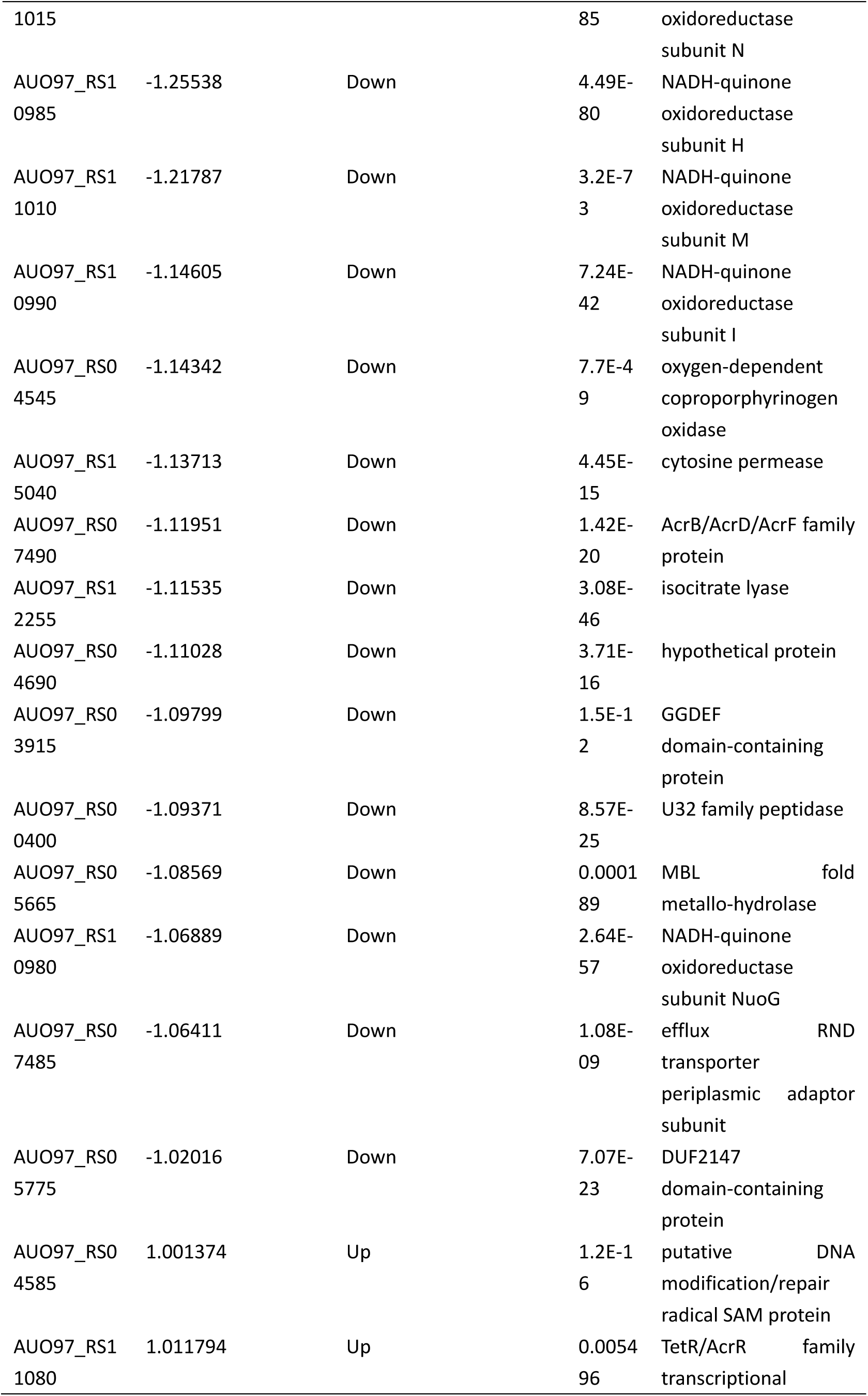
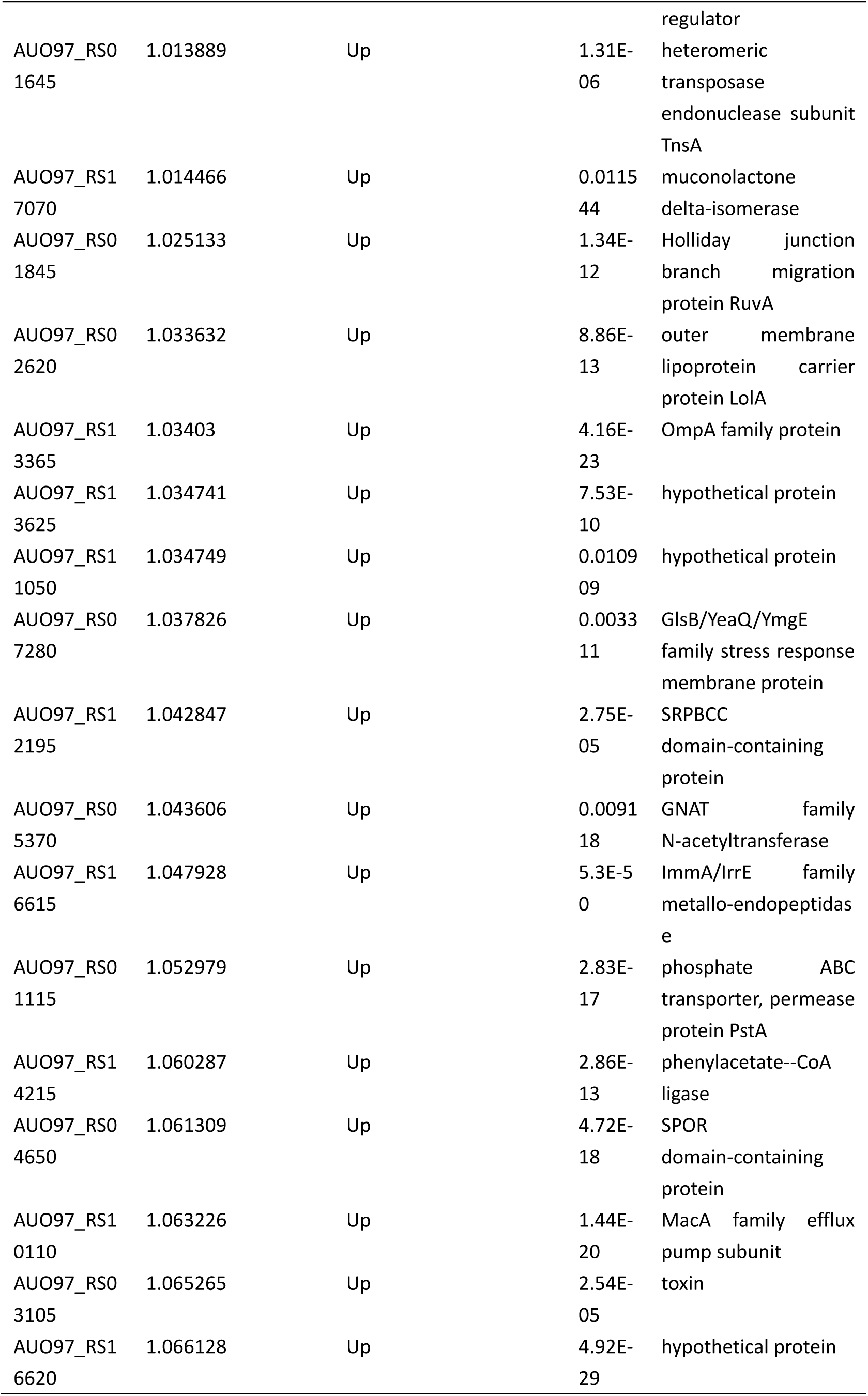
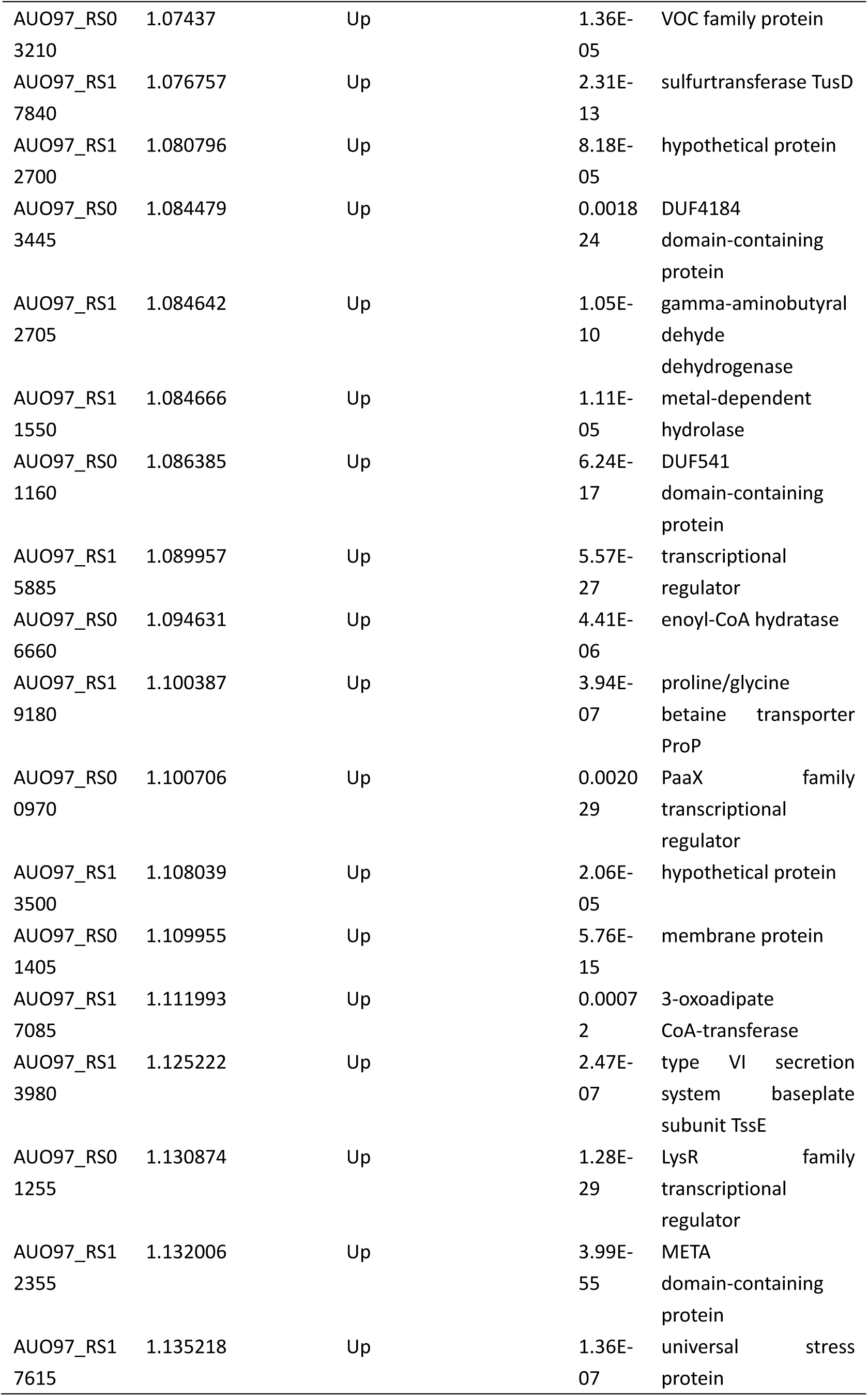
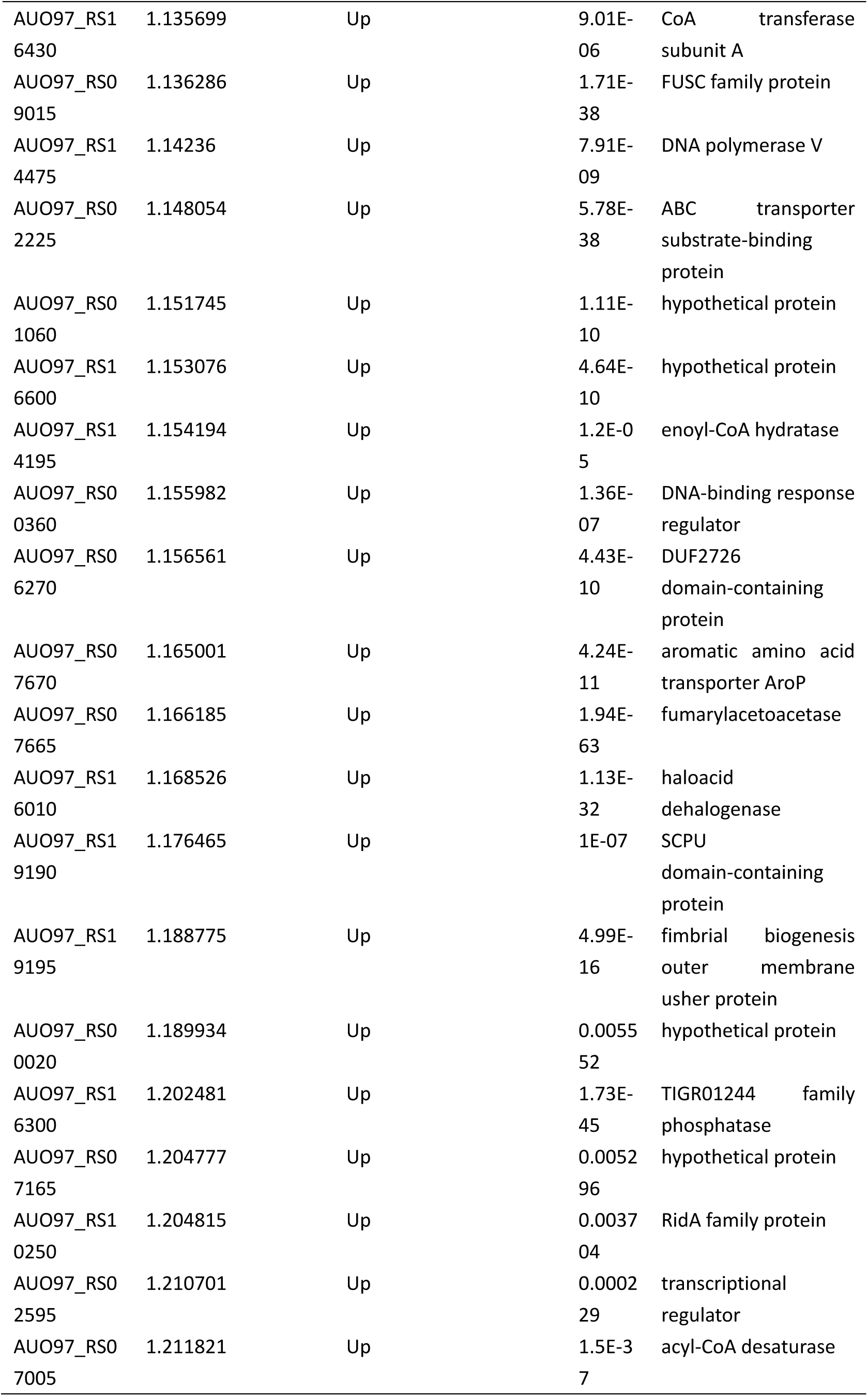
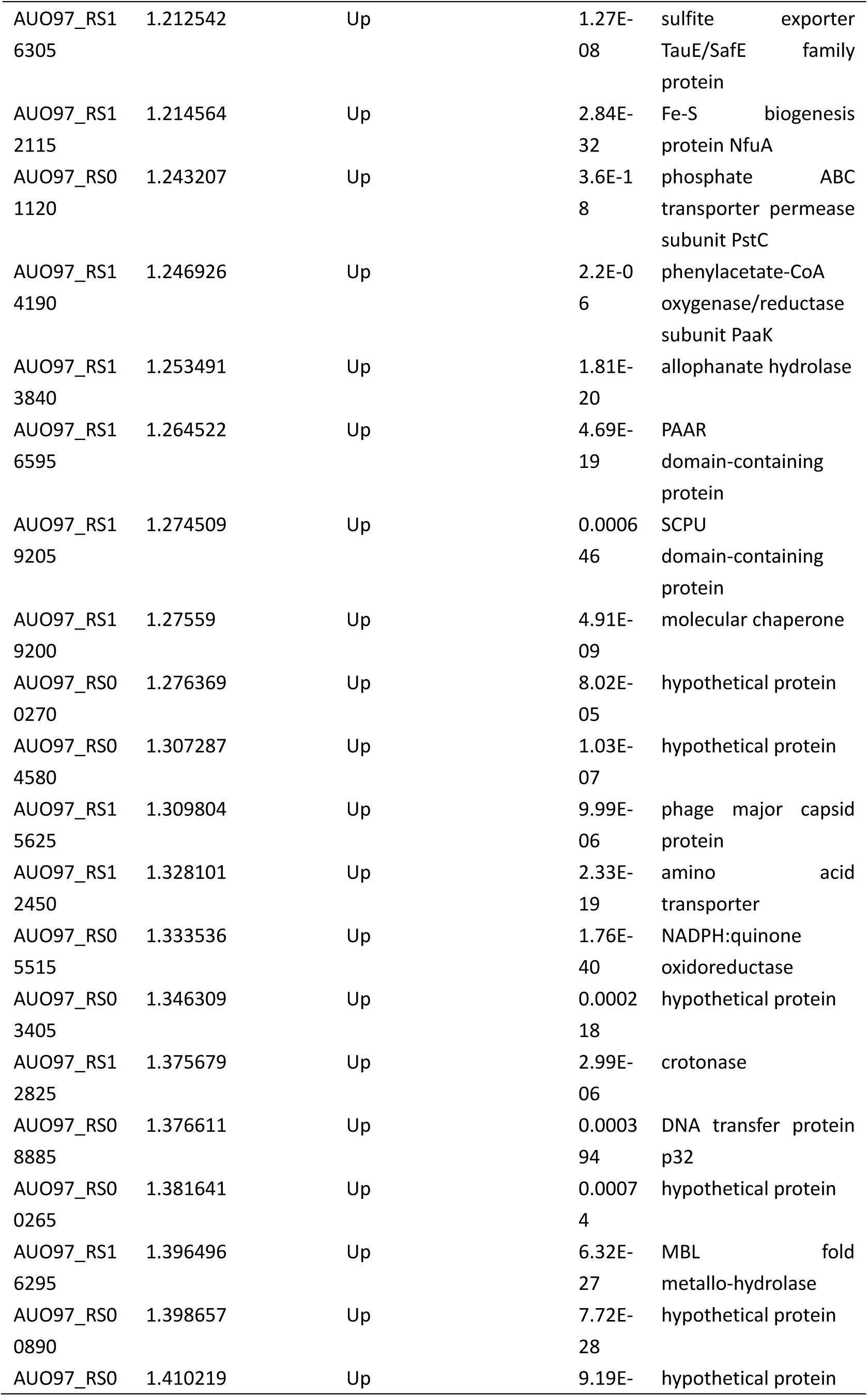
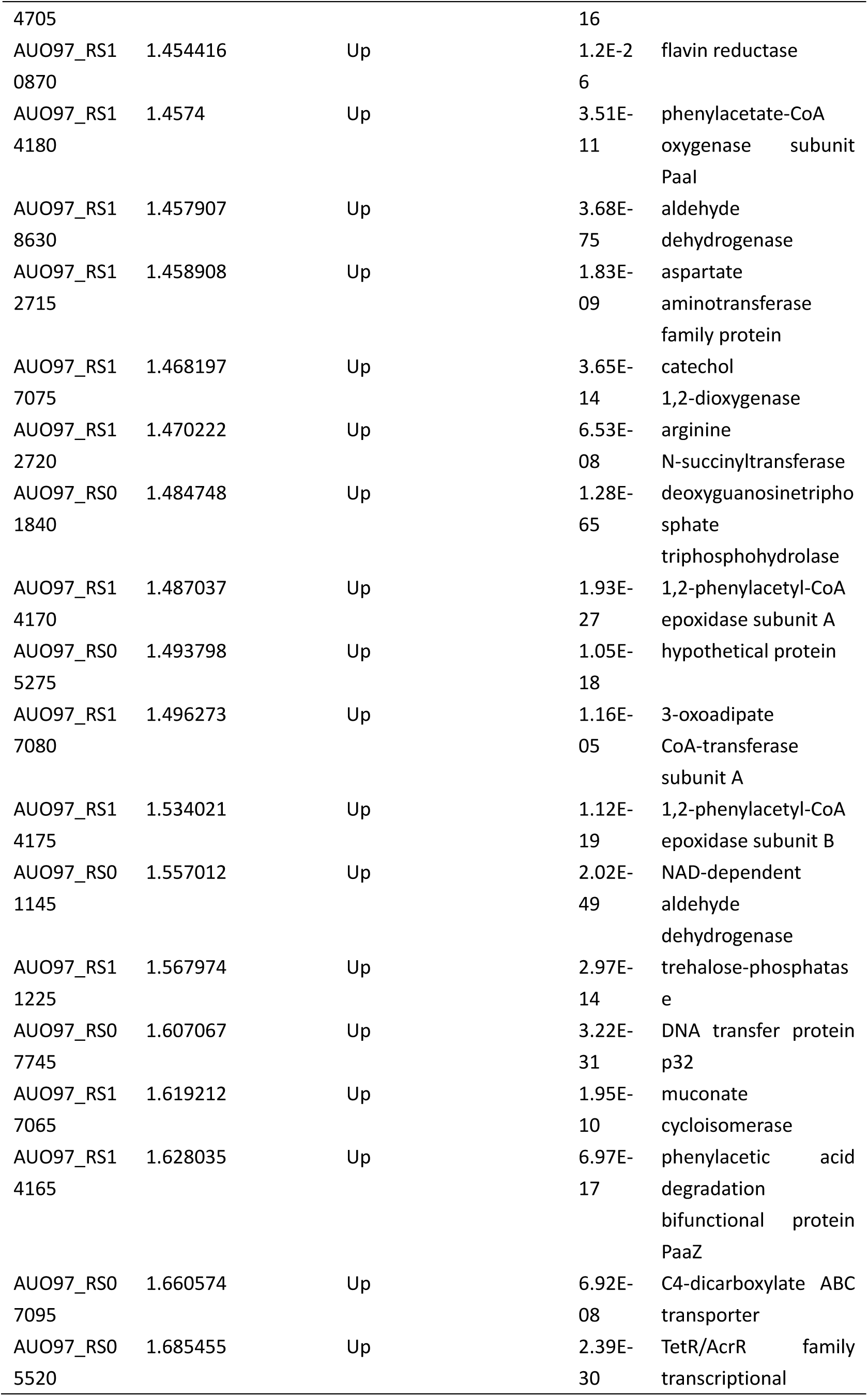
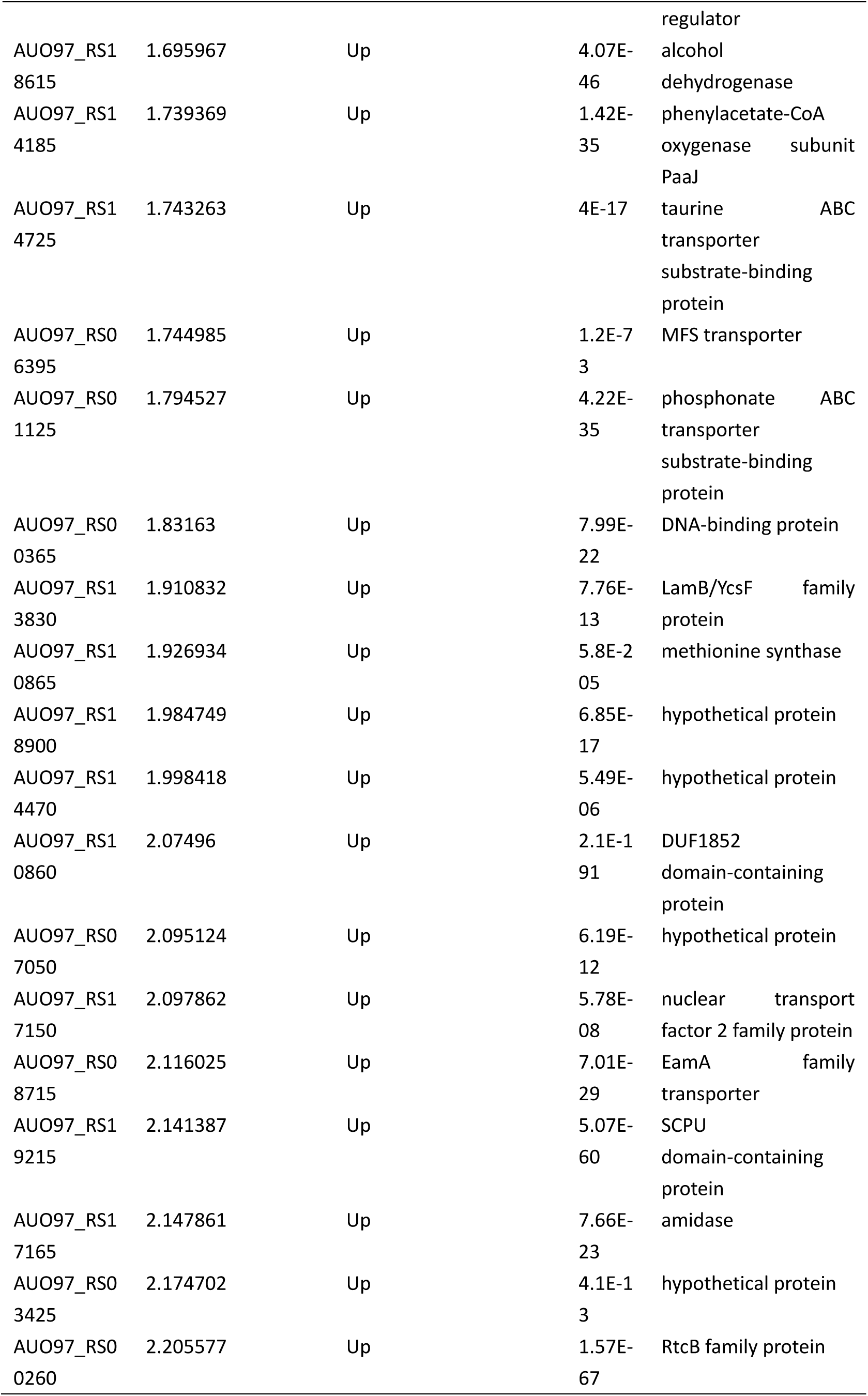
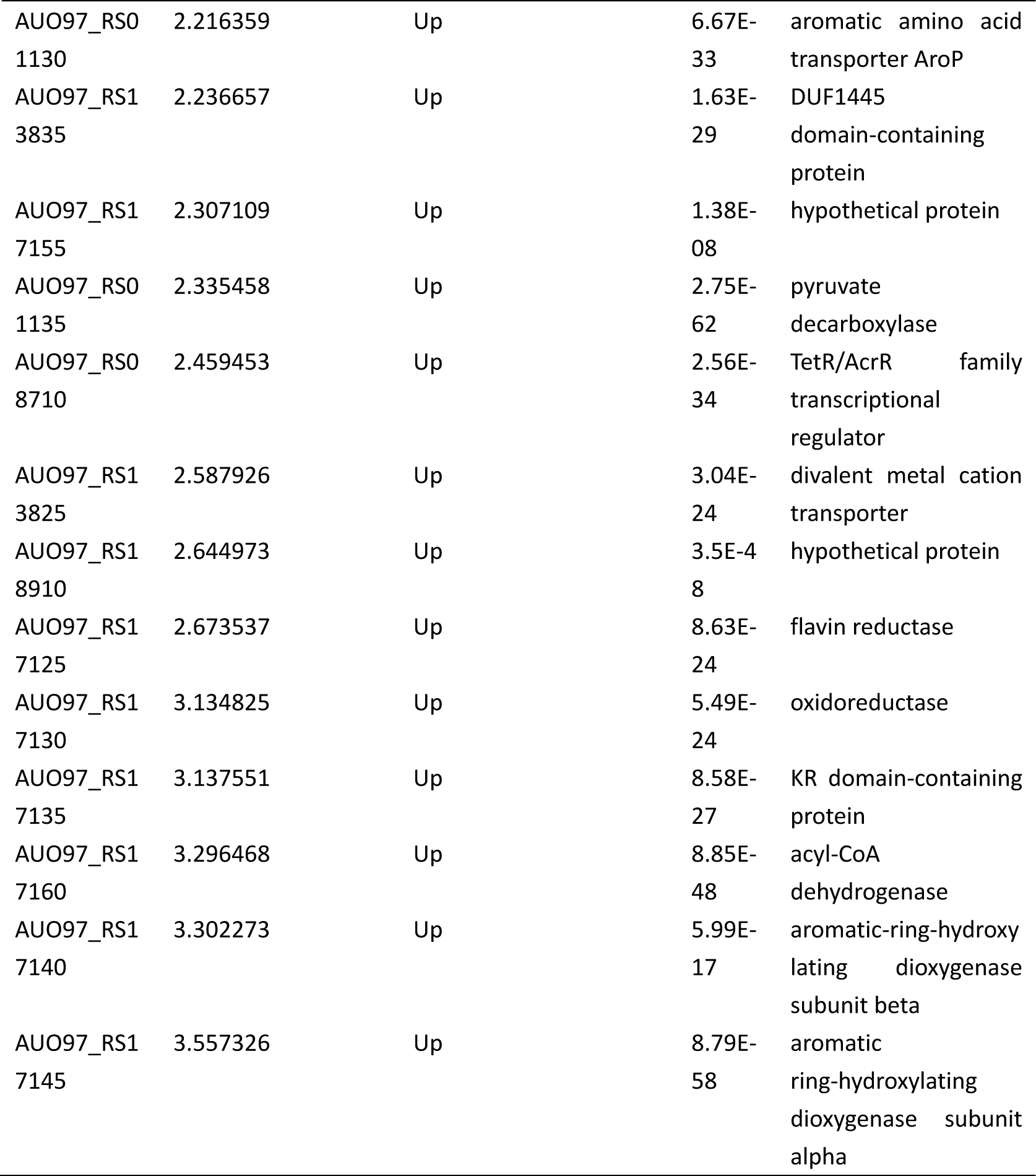
Differential expressed genes in ΔabaI strain

**Table 5.**
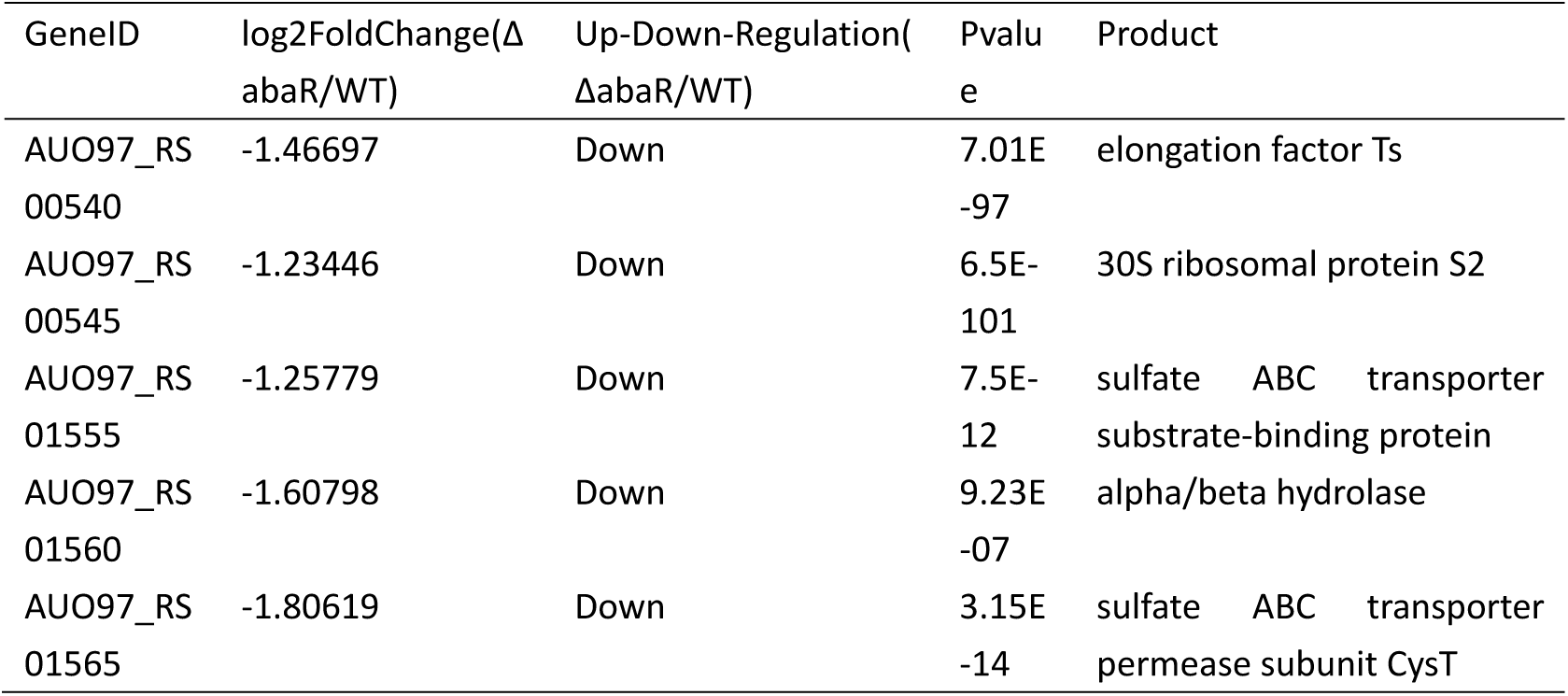
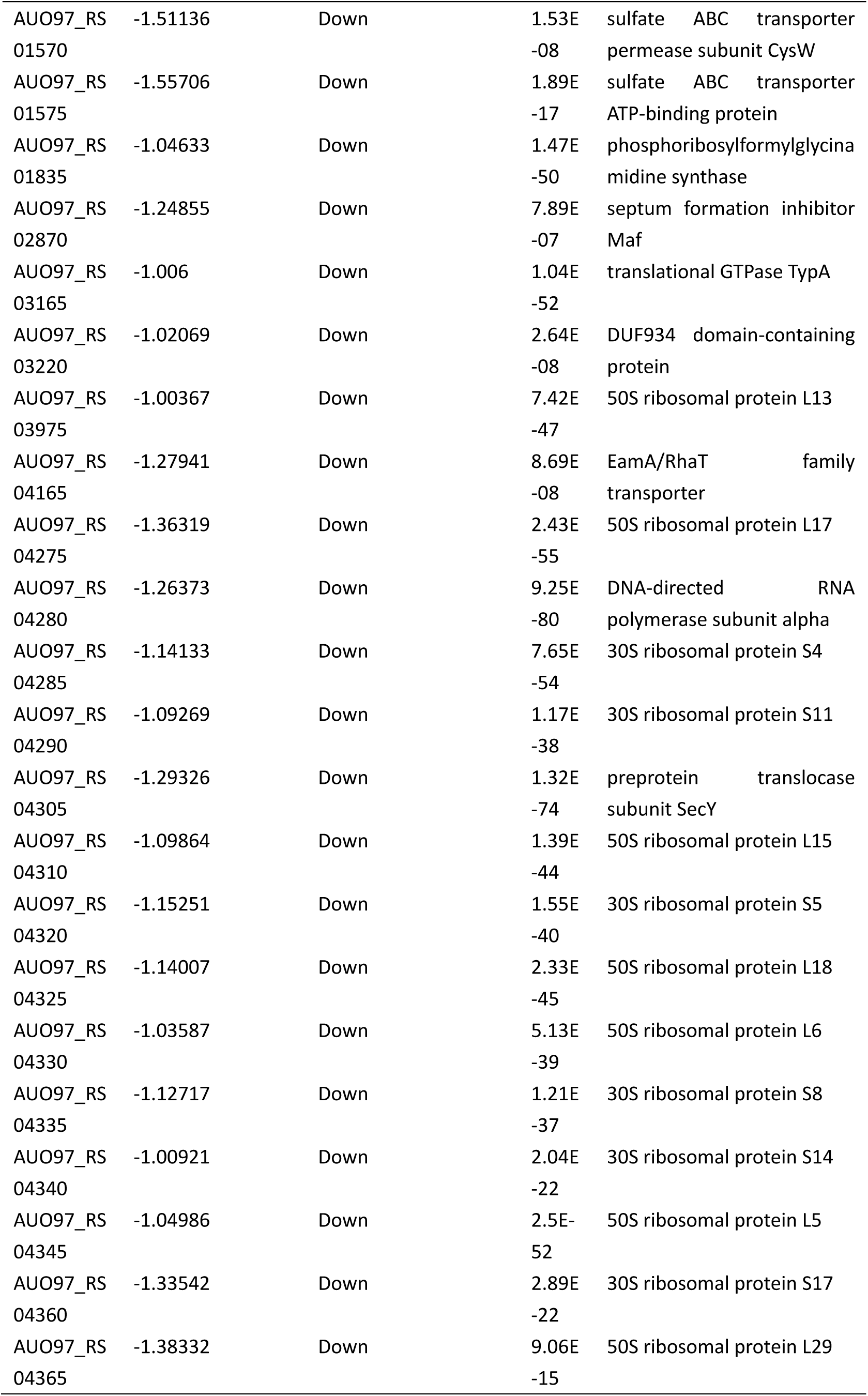
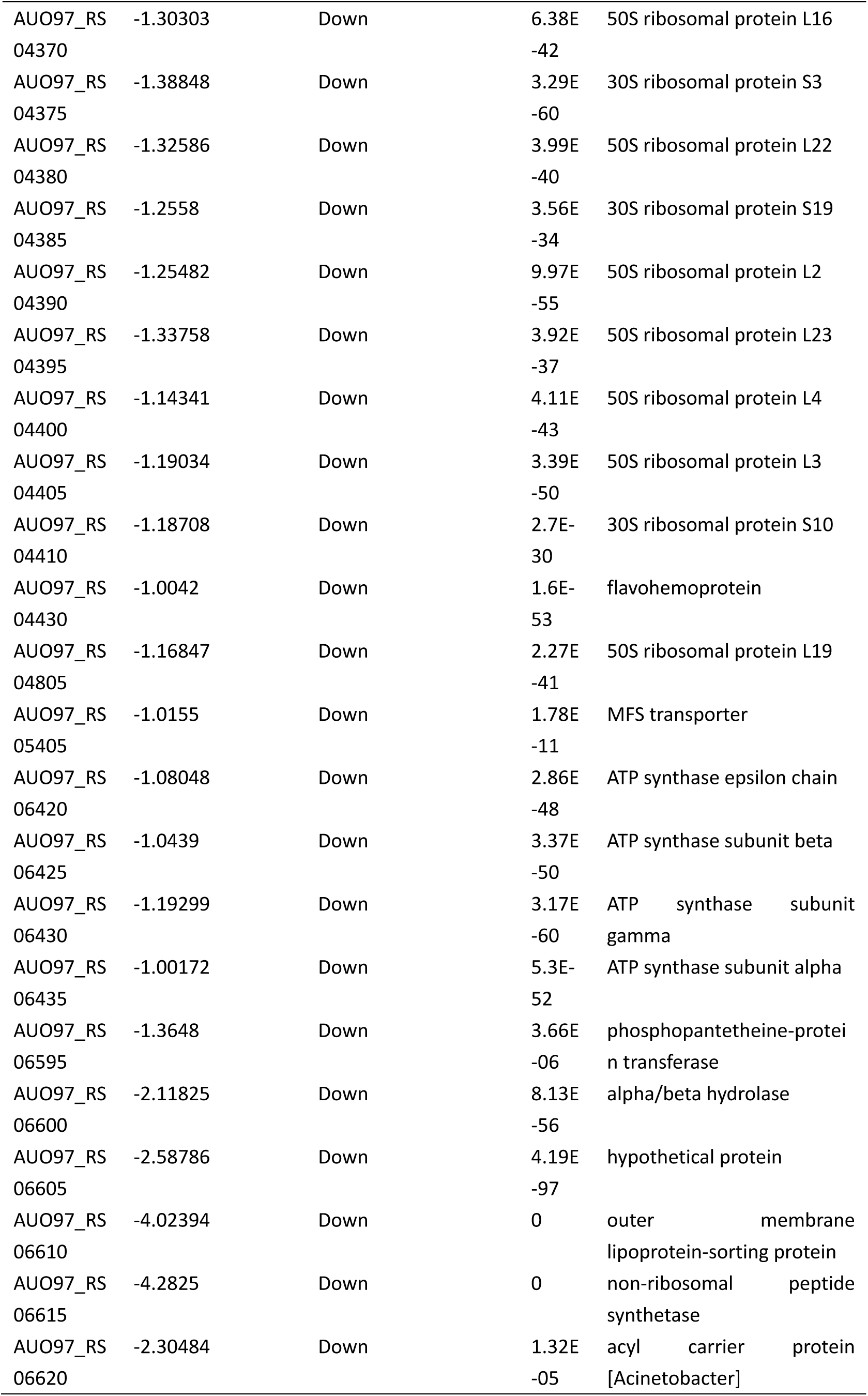
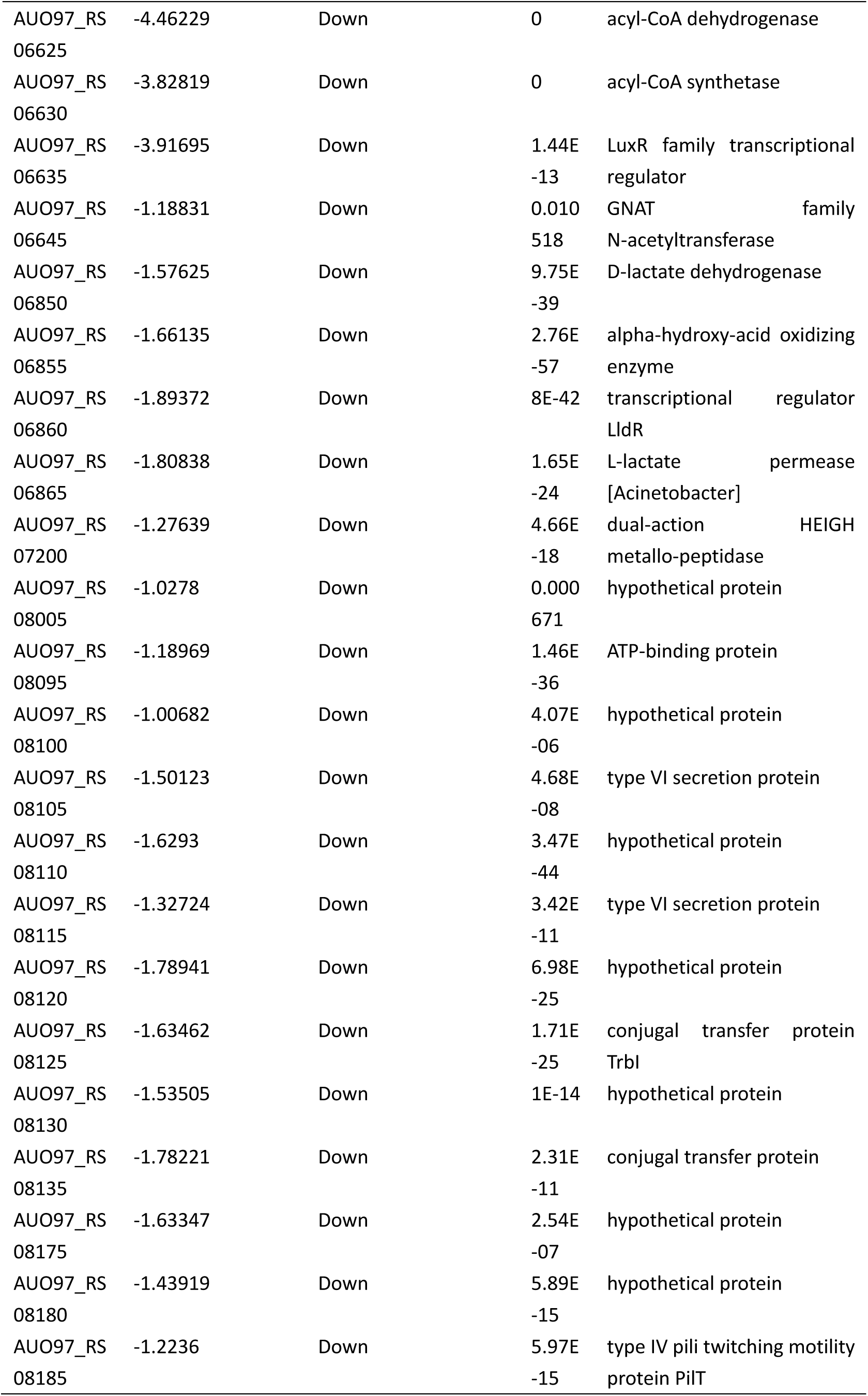
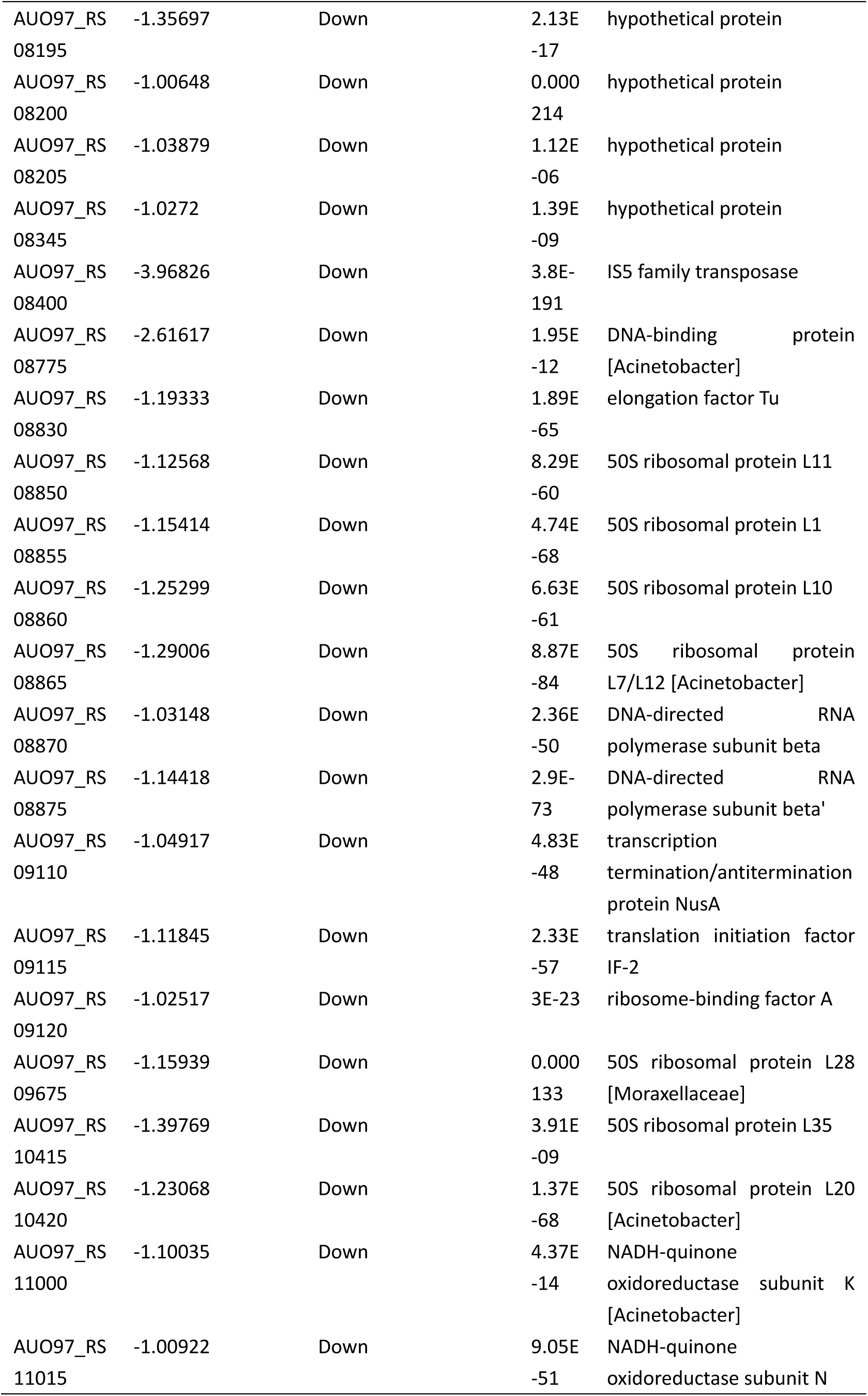
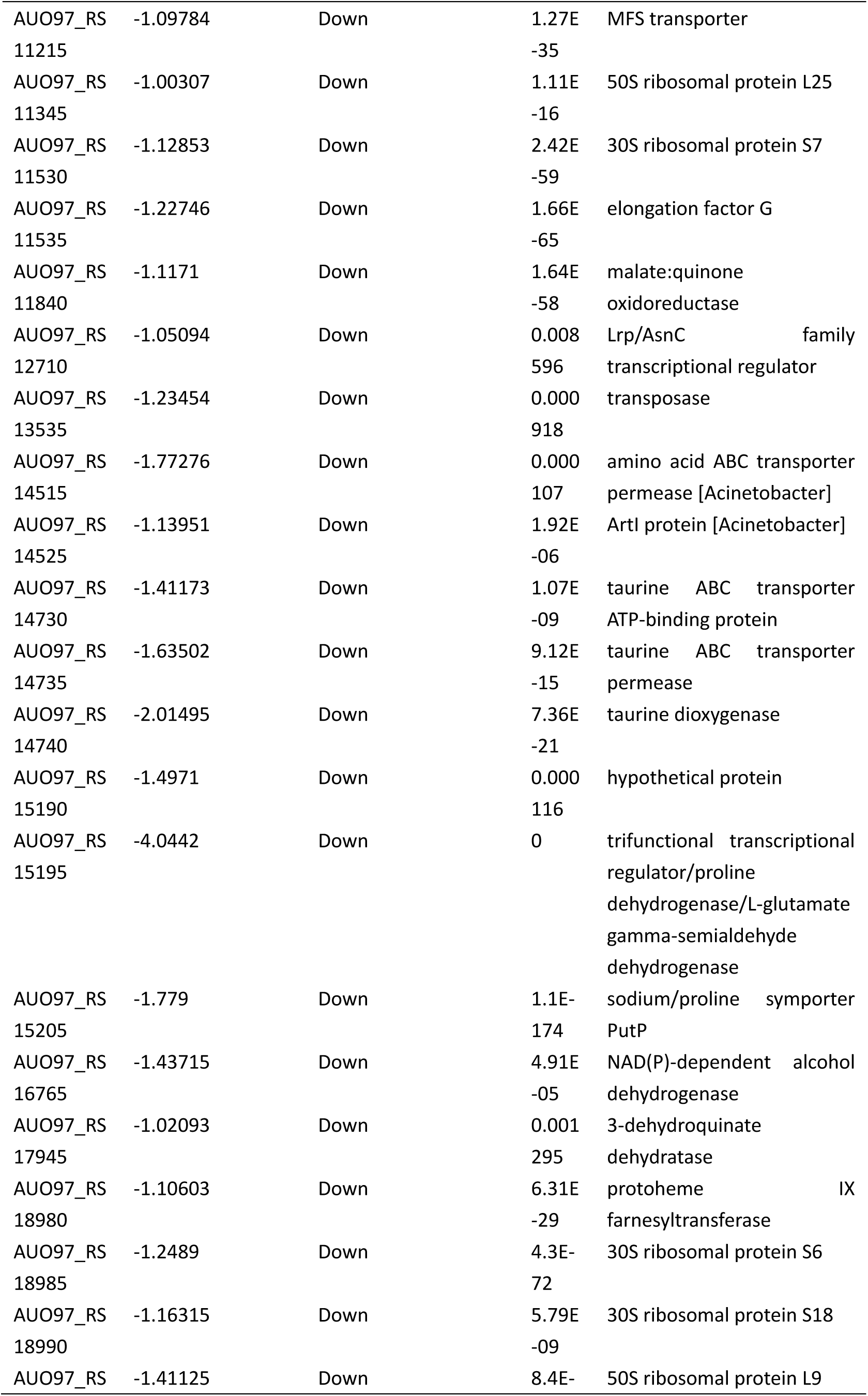
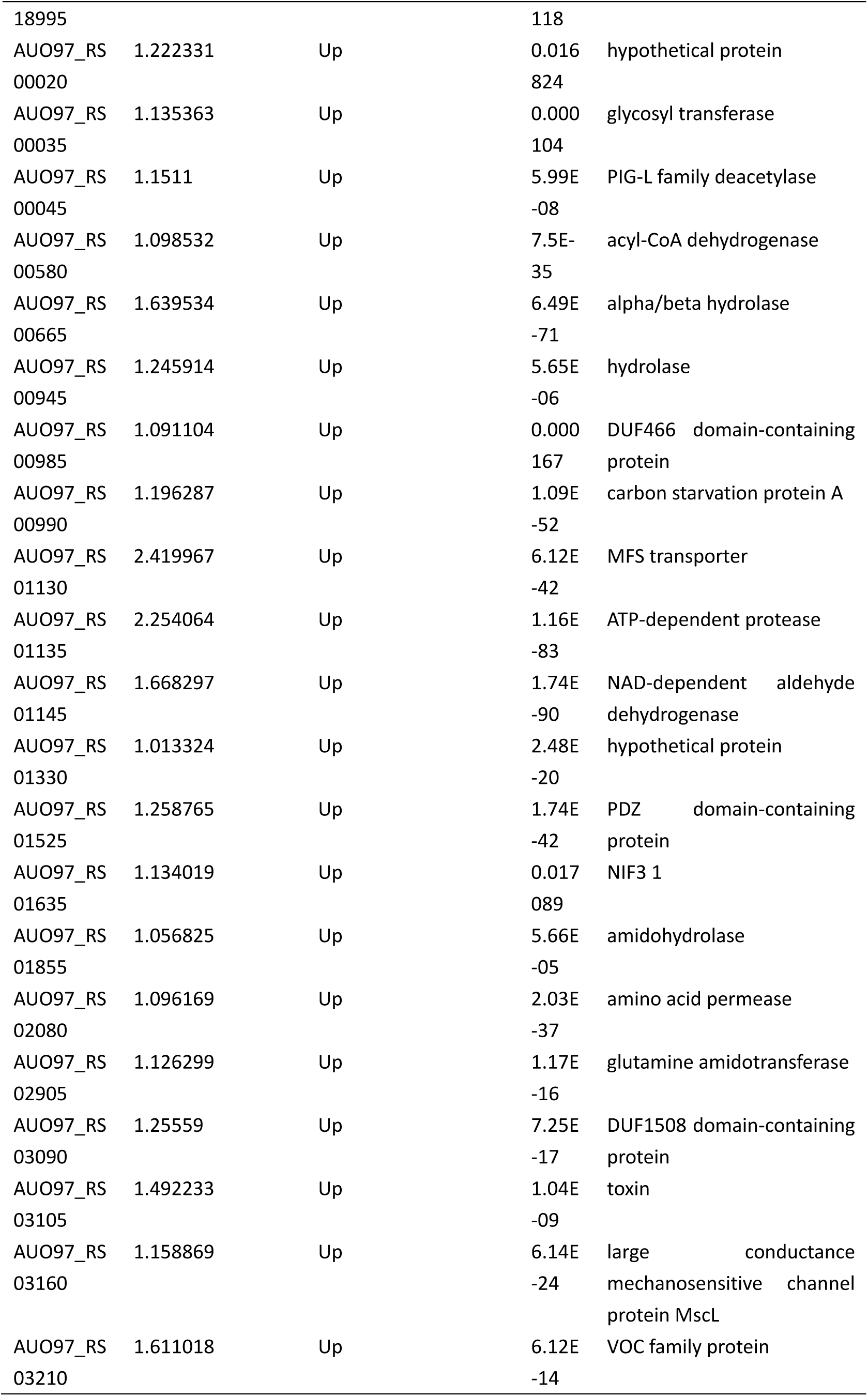
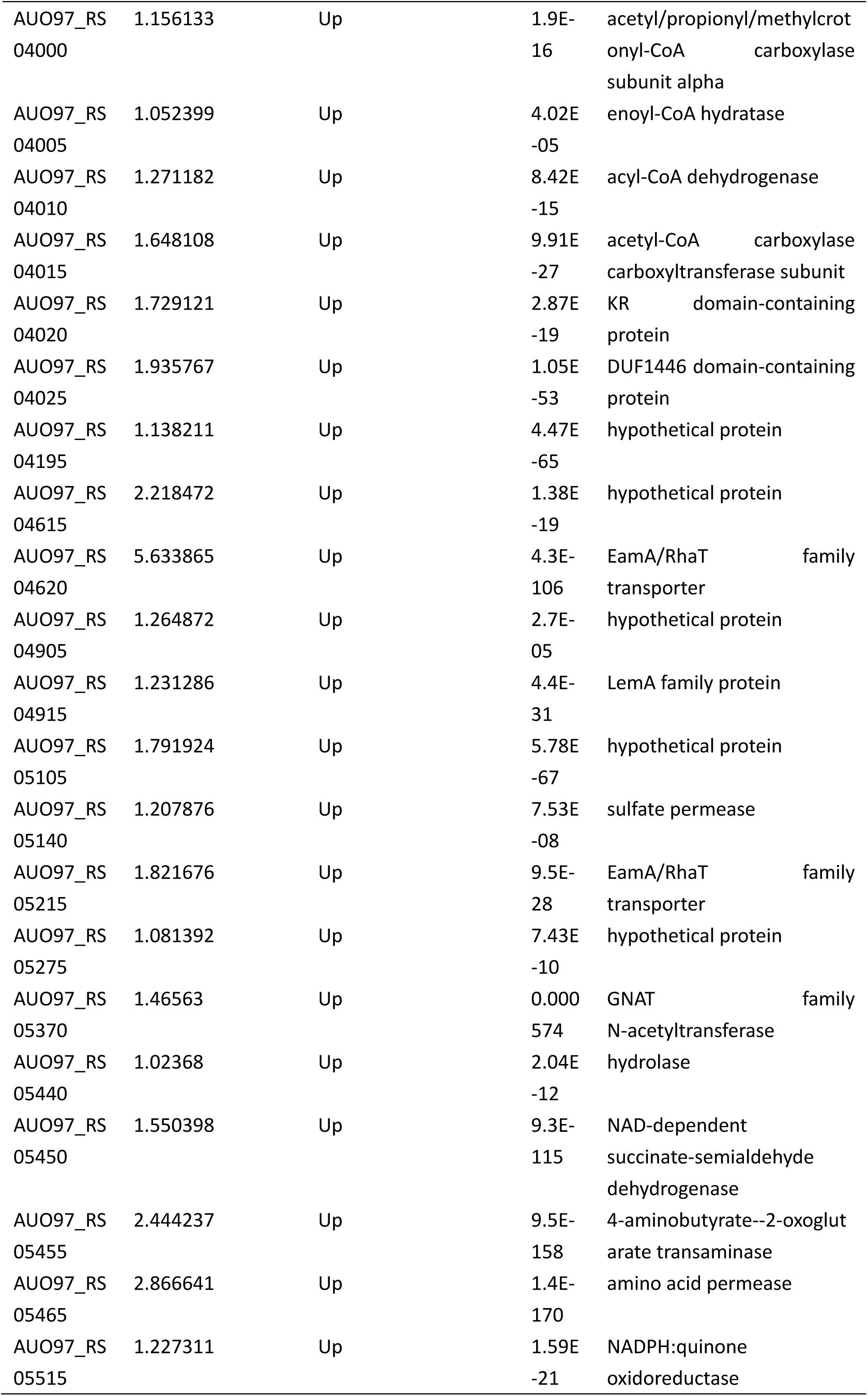
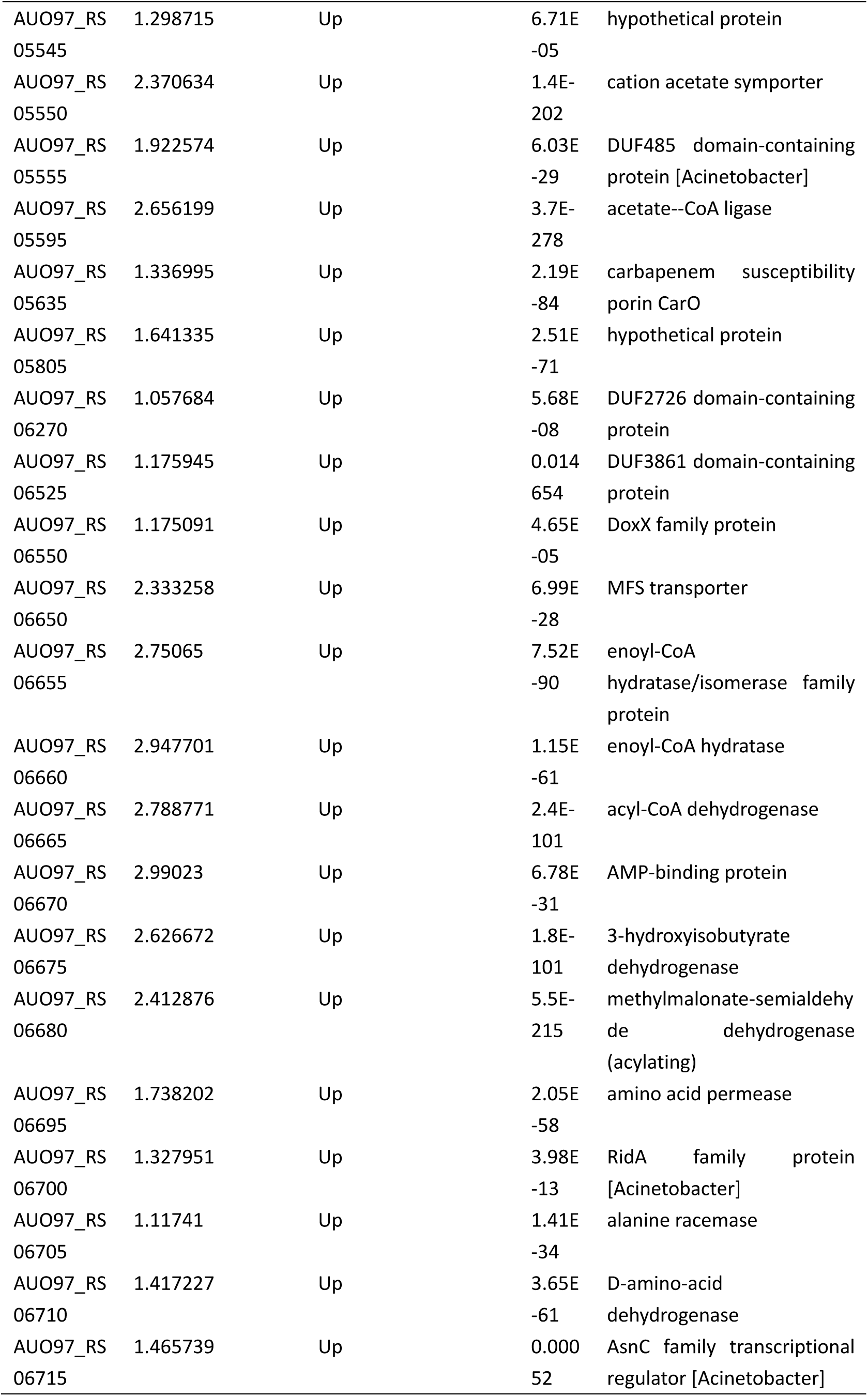
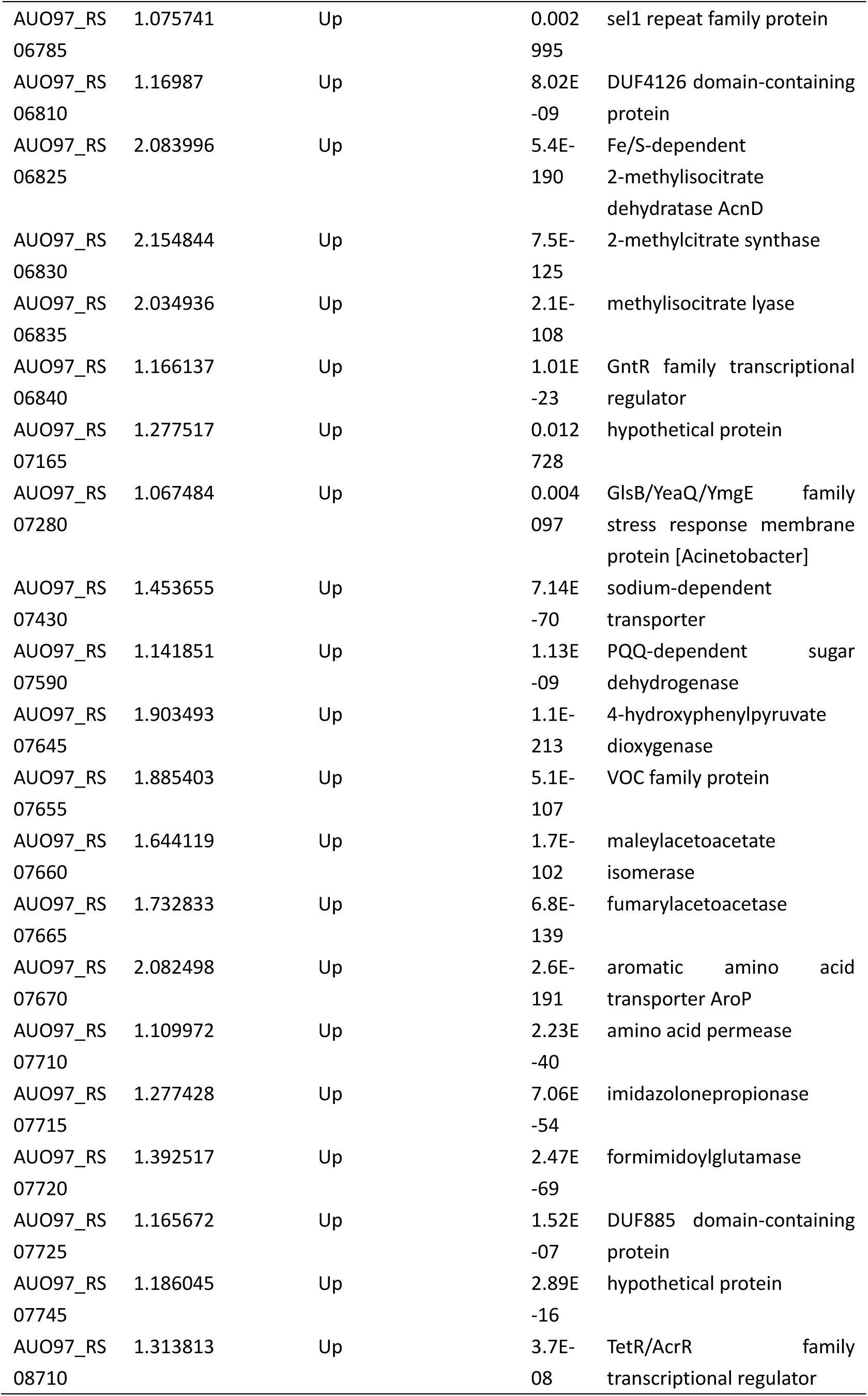
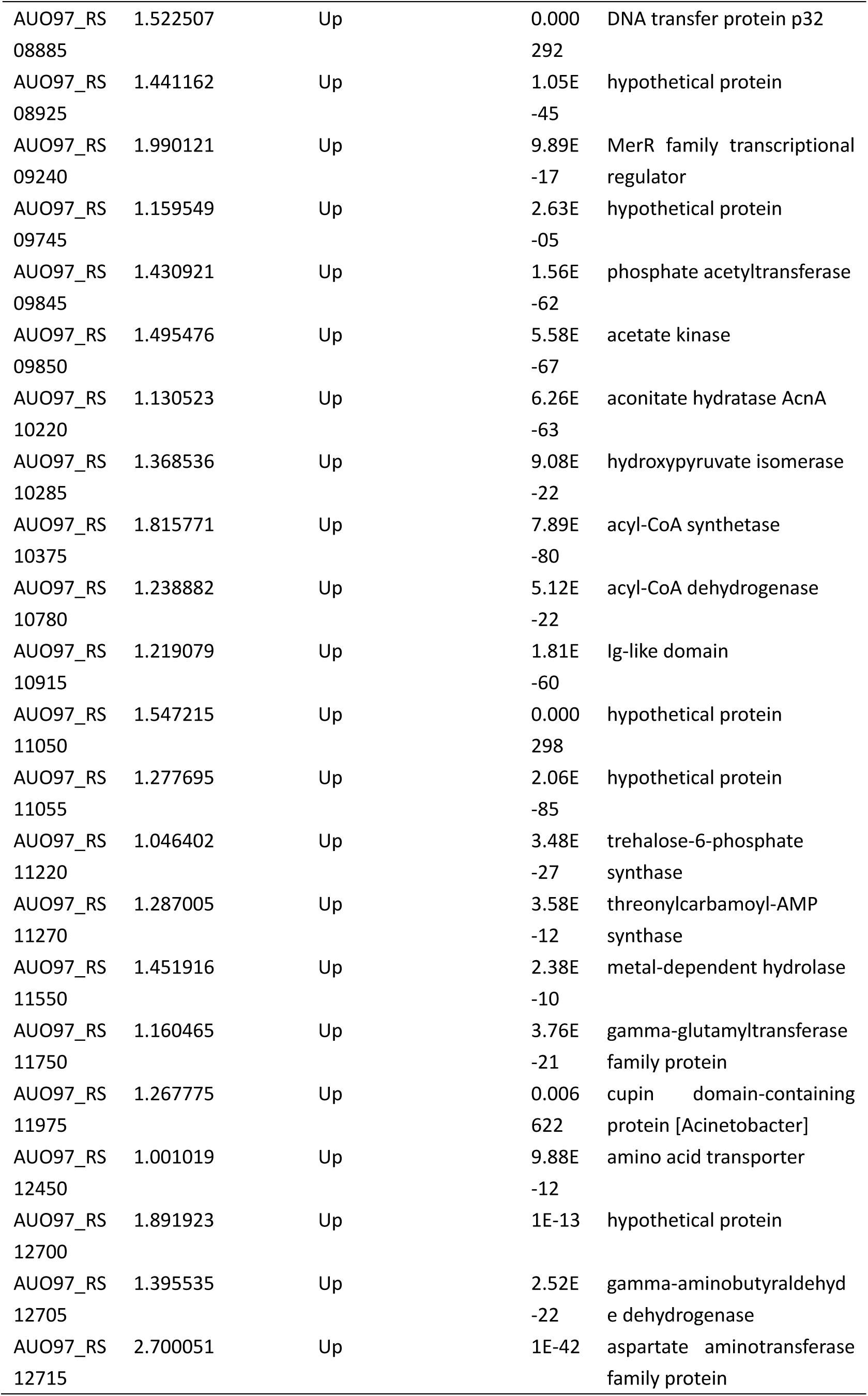
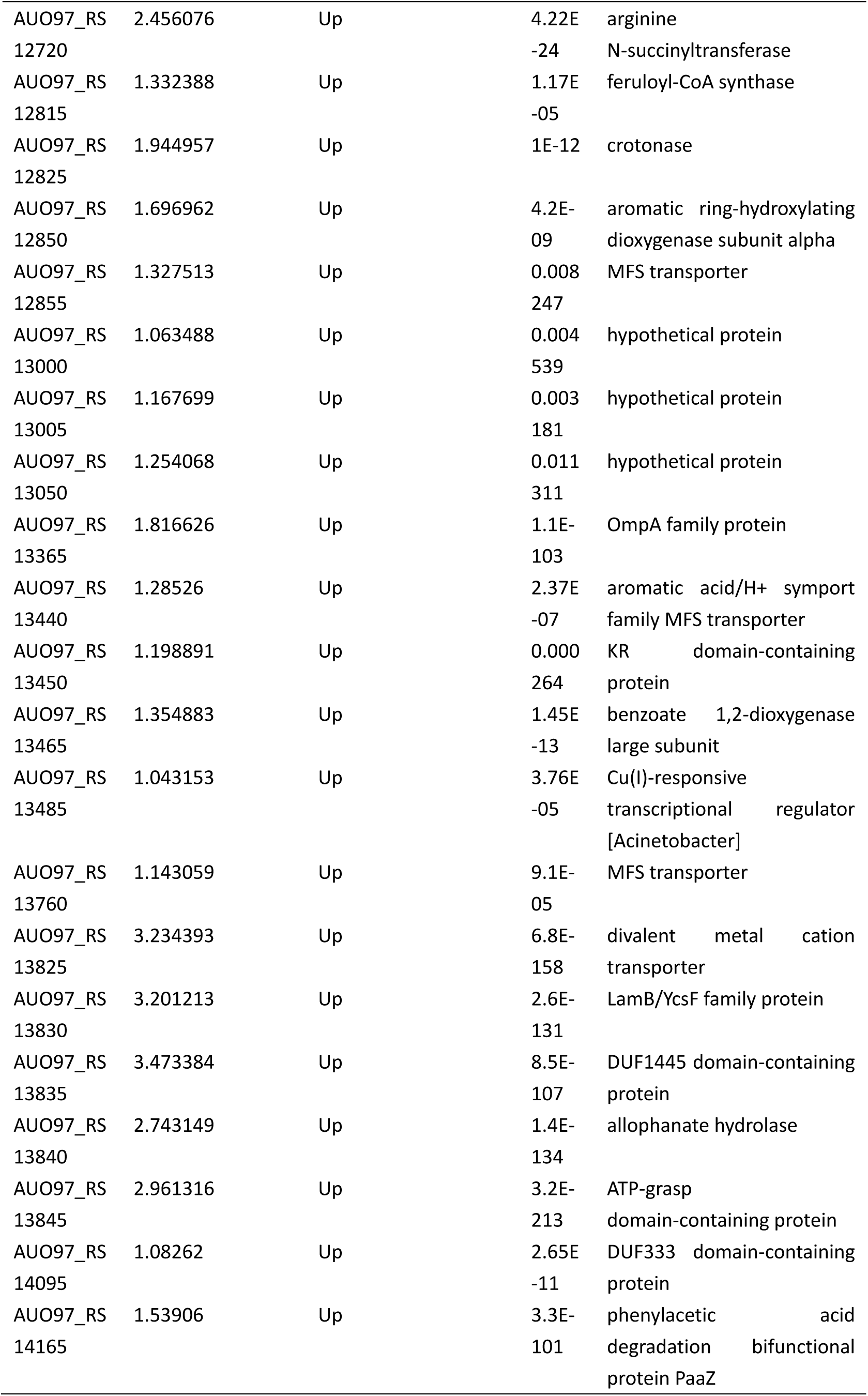
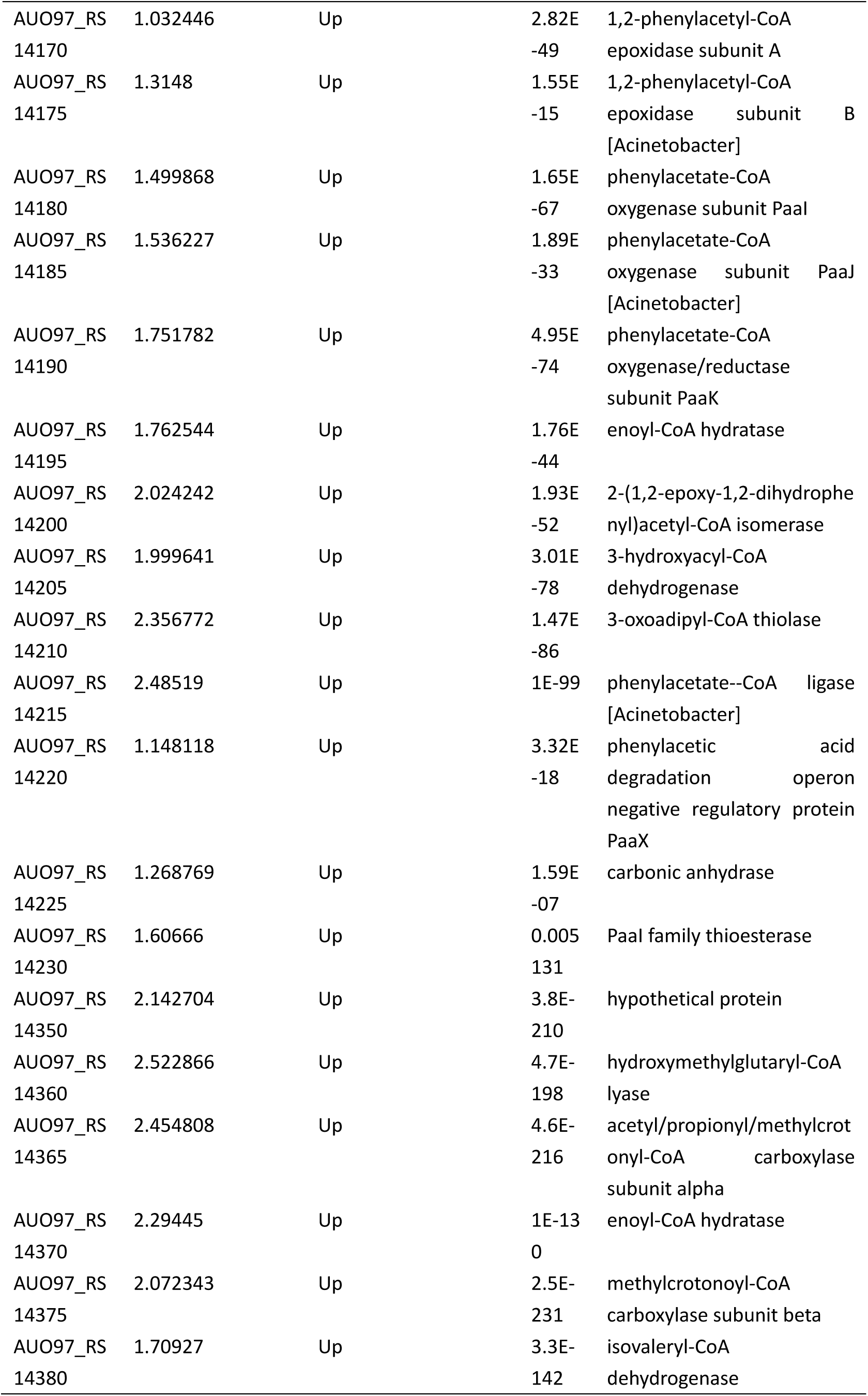
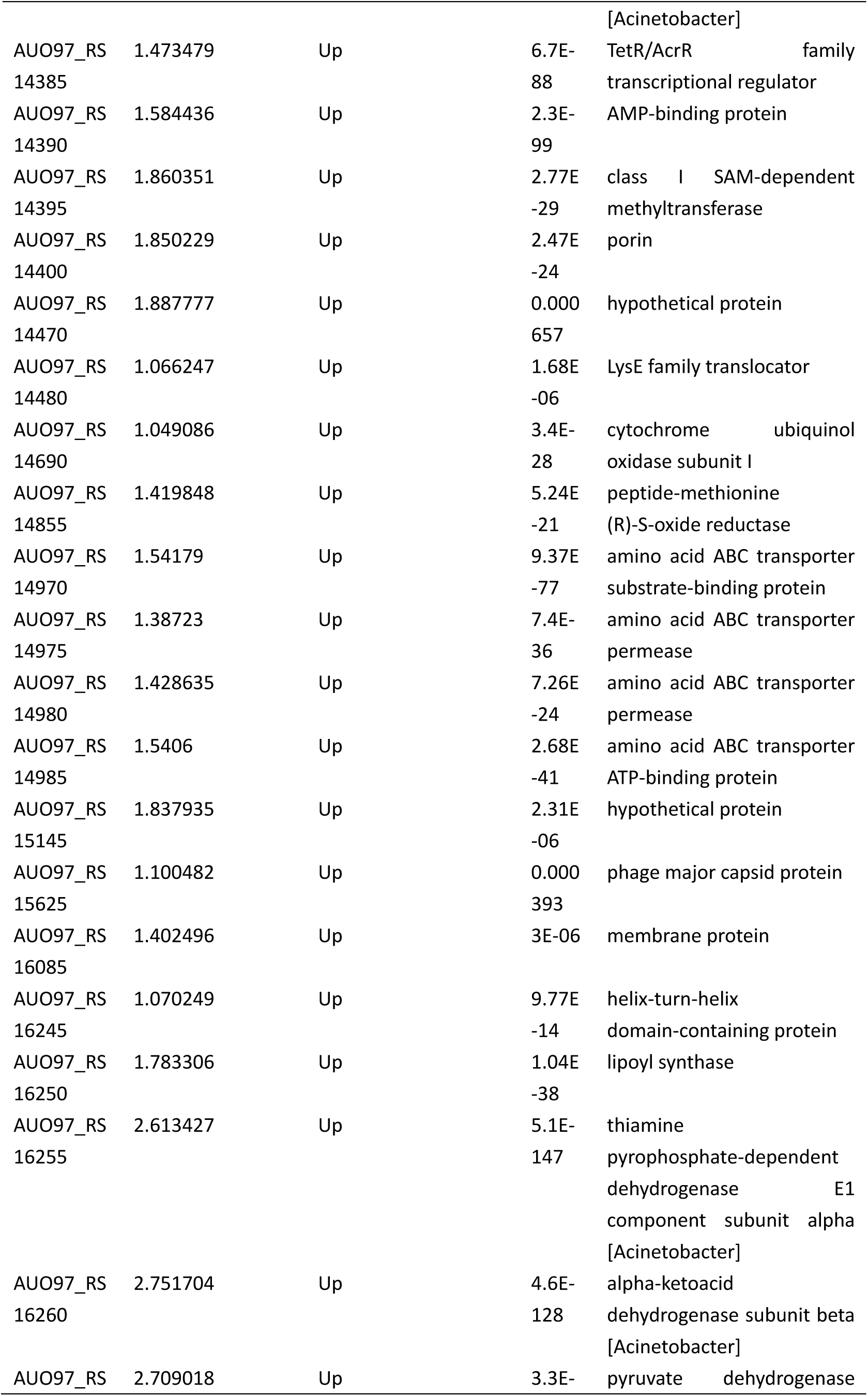
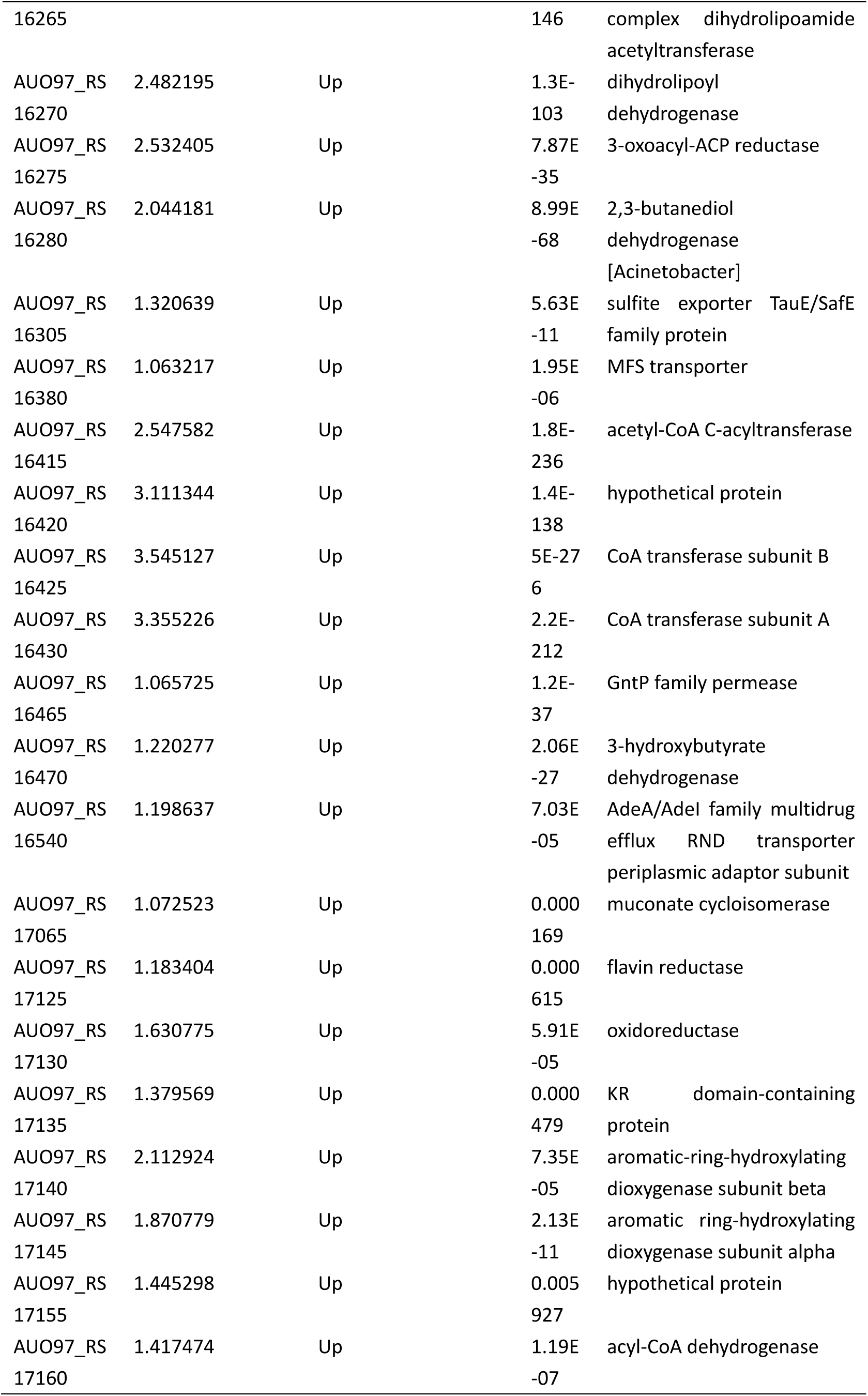
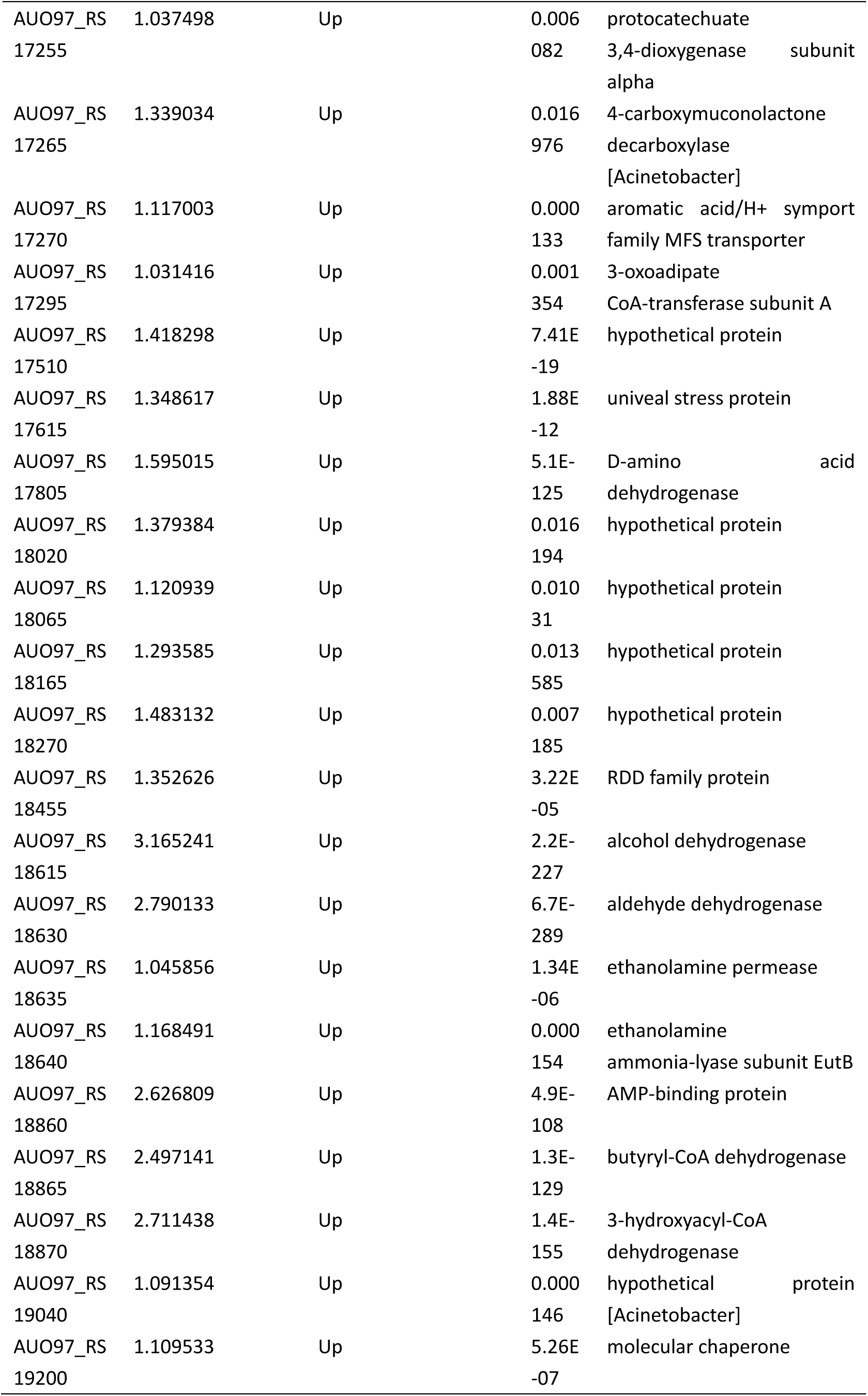
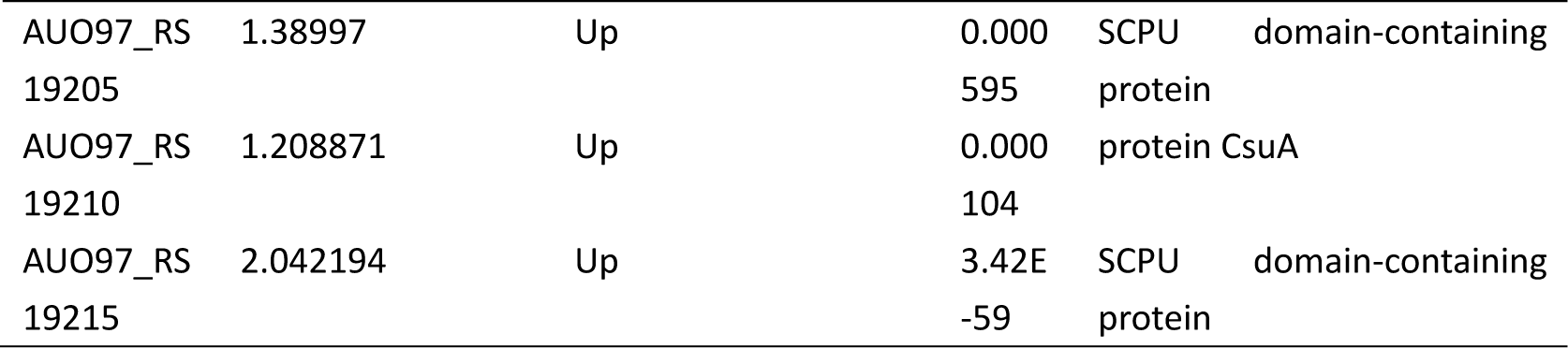
Differential expressed genes in ΔabaR strain

**Table 6.**
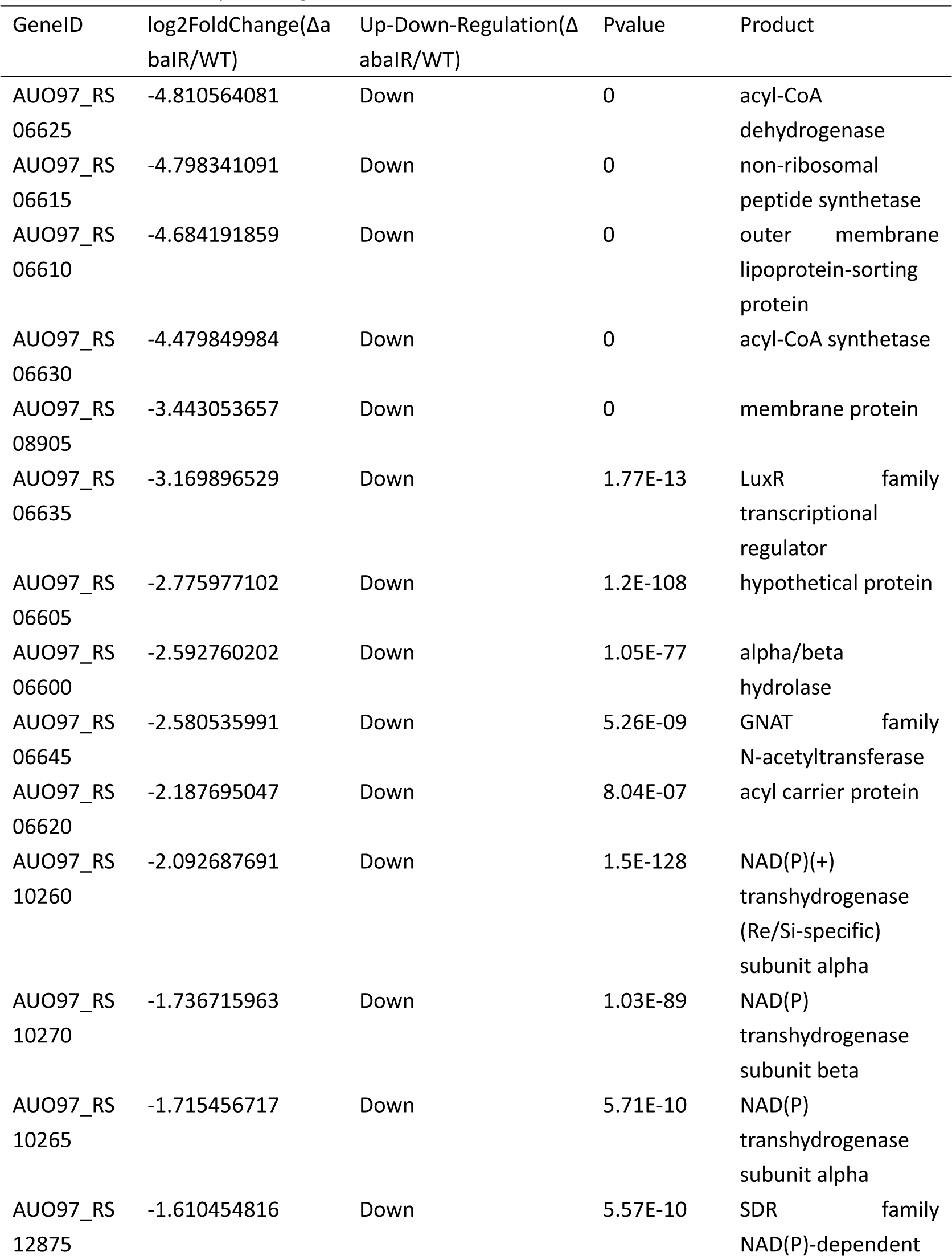
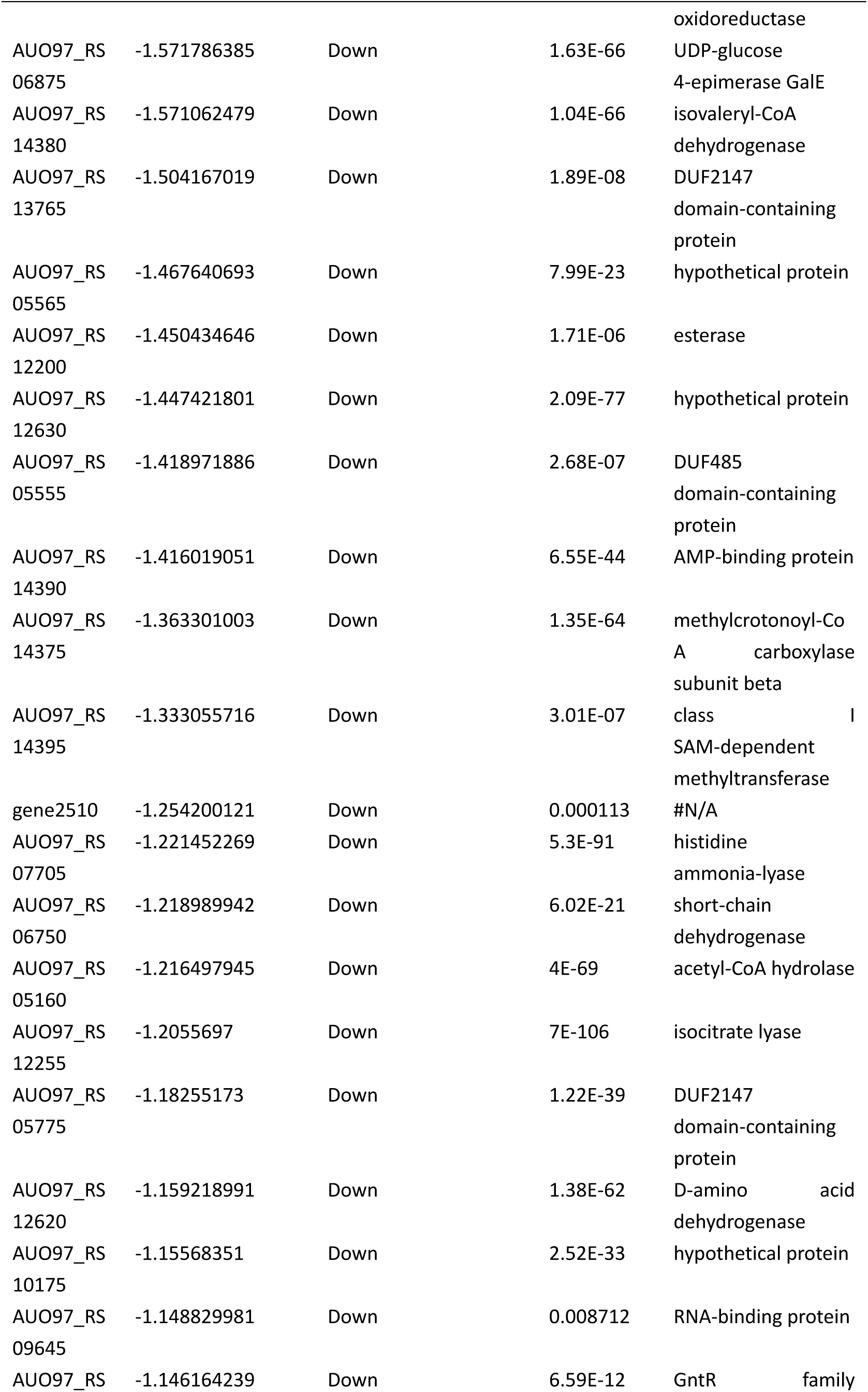
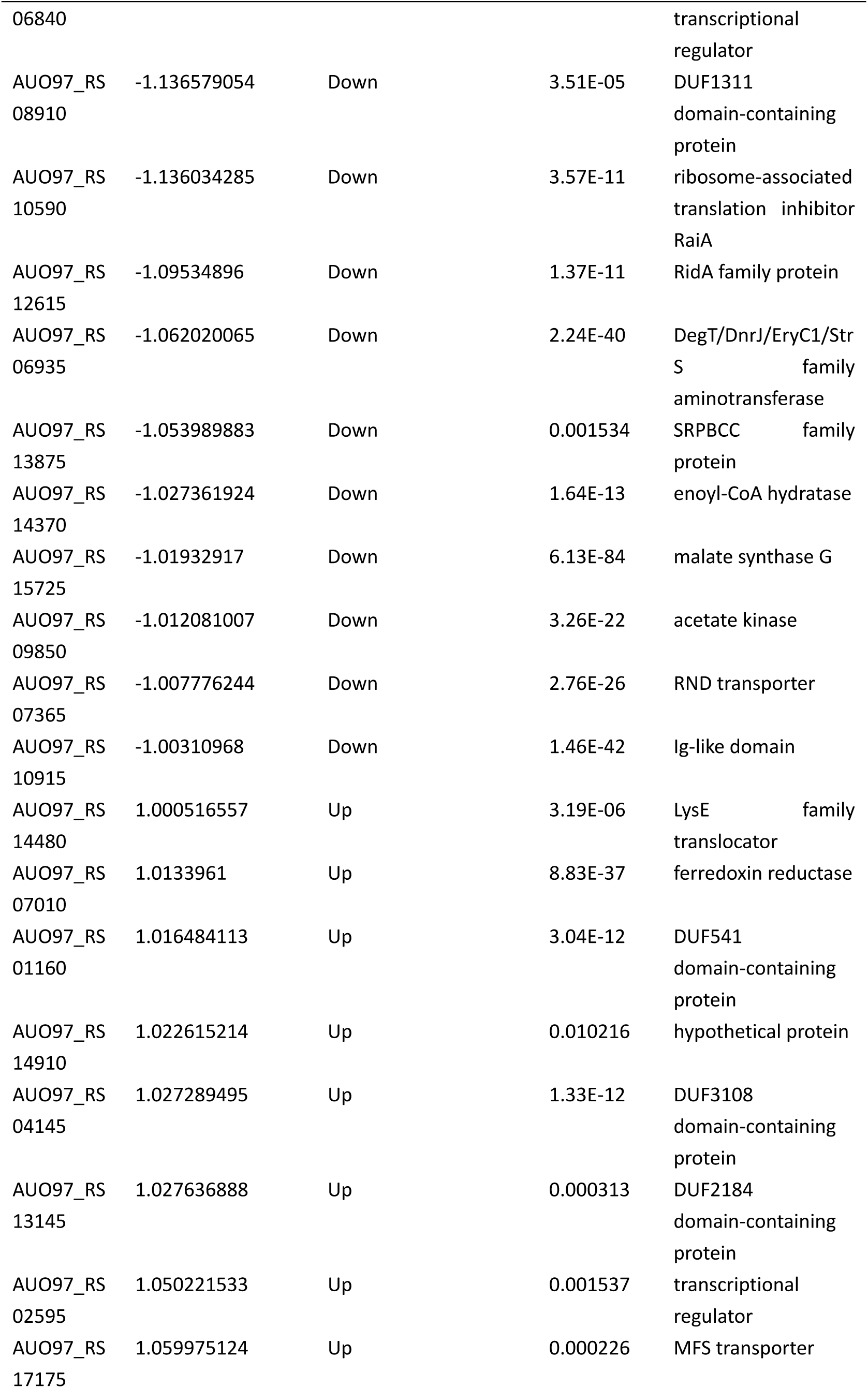
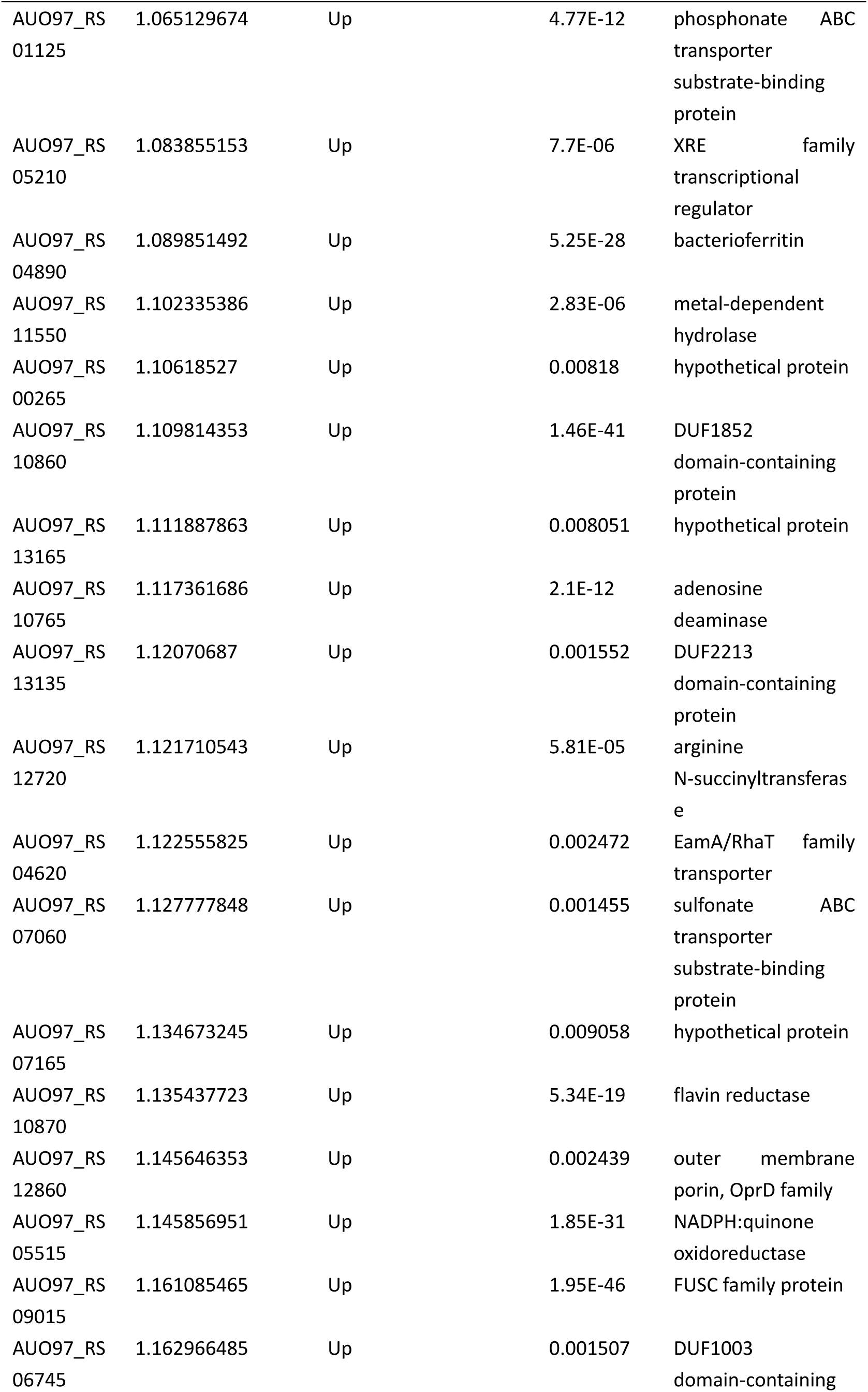
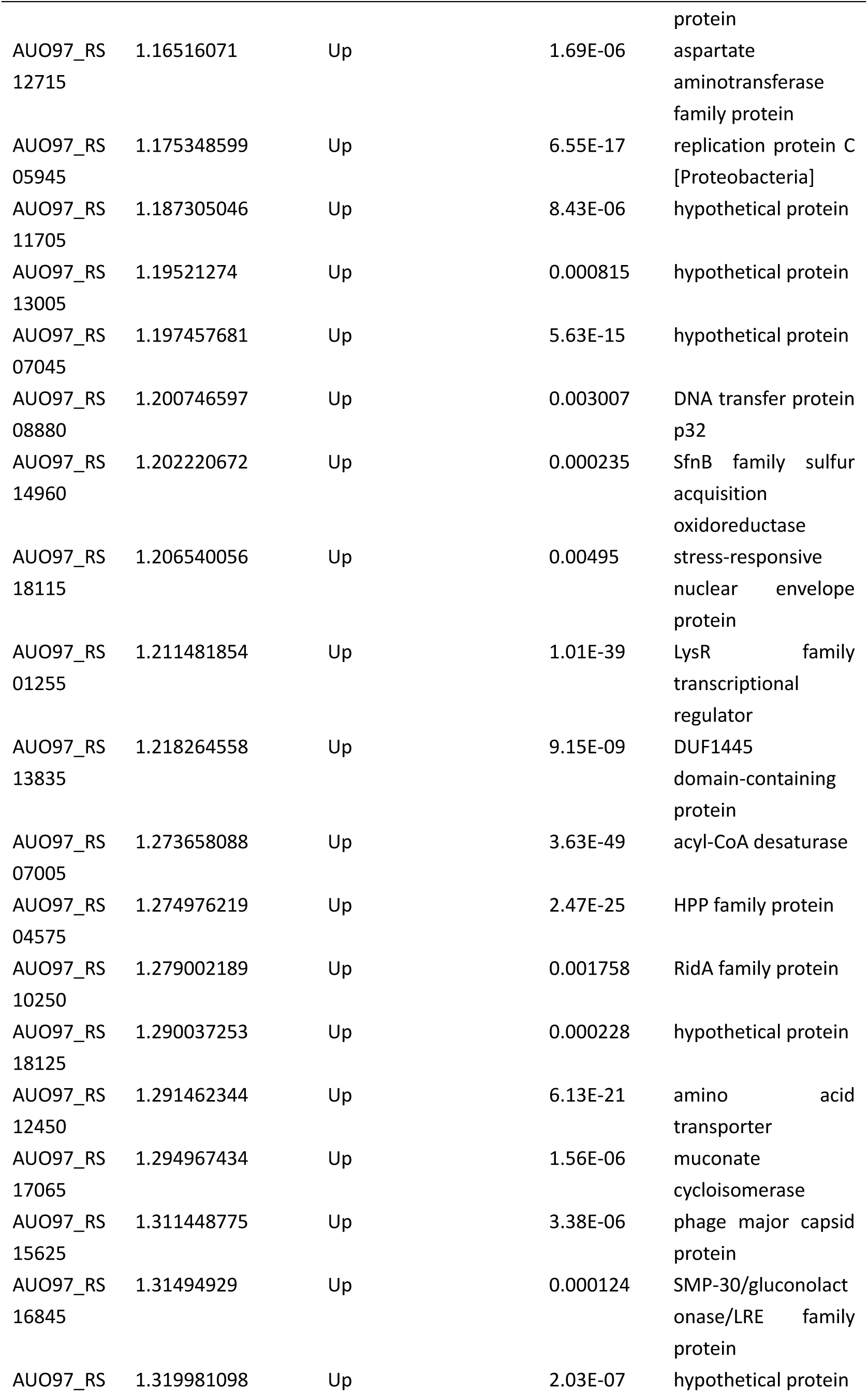
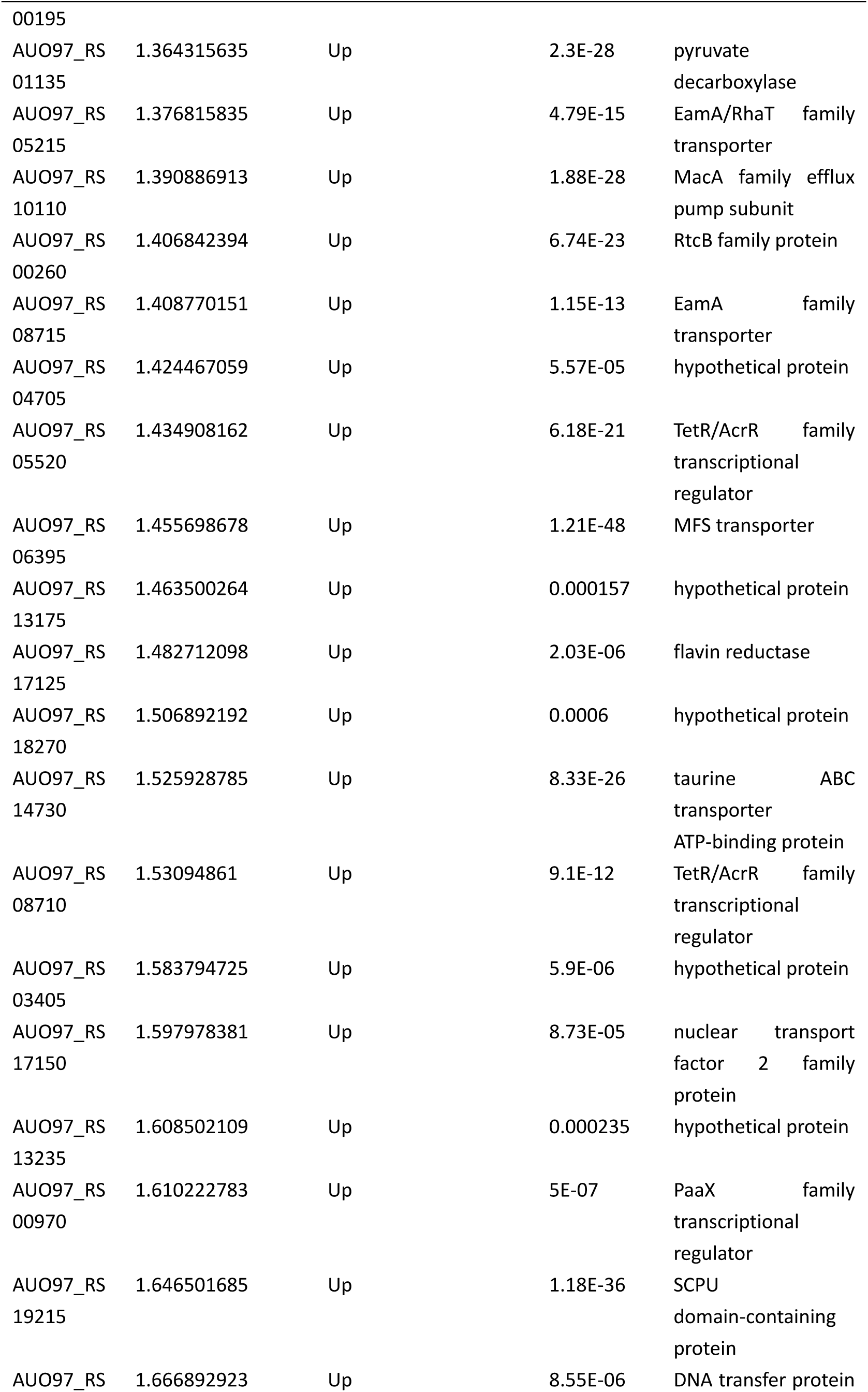
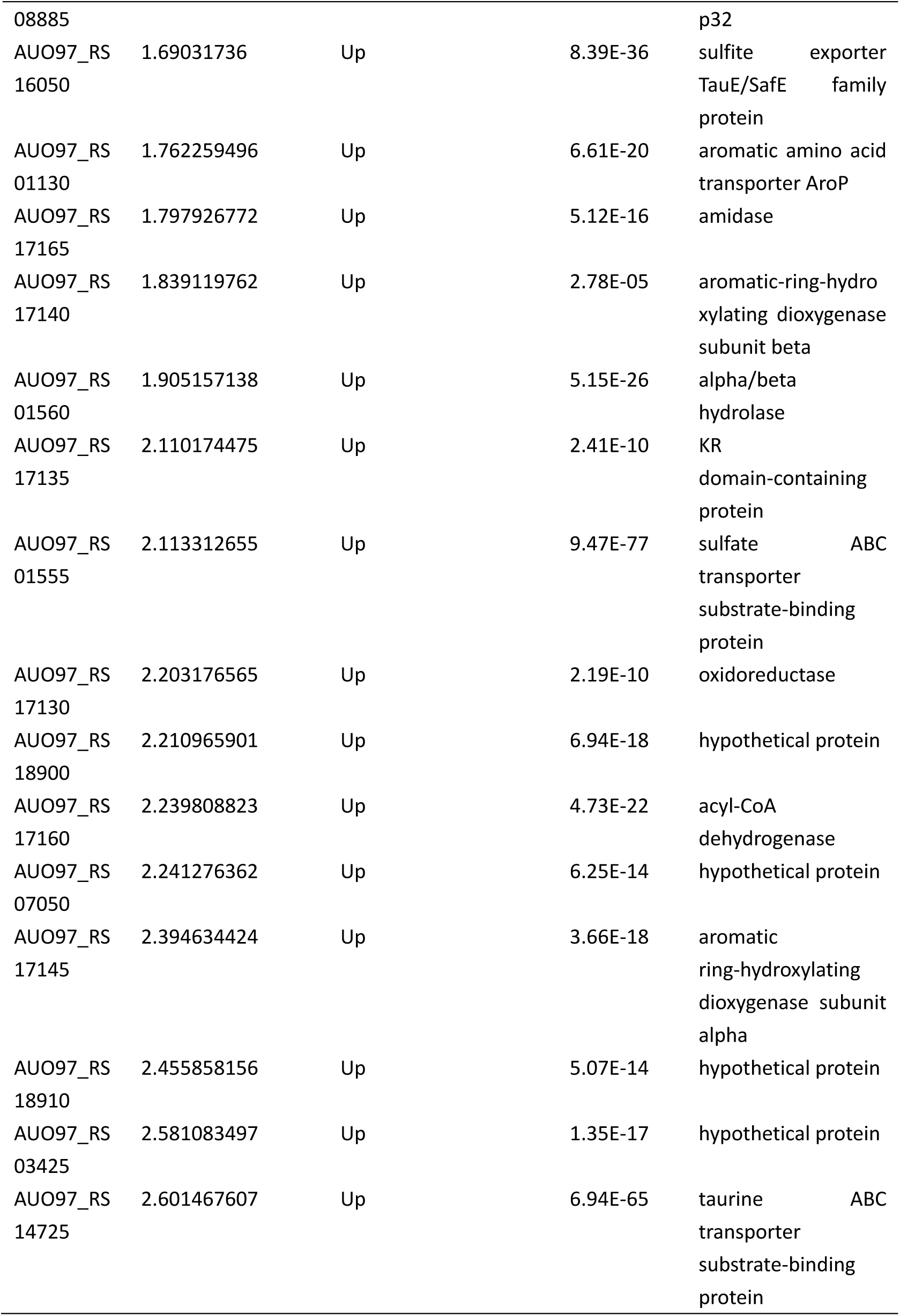
Differential expressed genes in ΔabaIR strain

